# STATegra: Multi-omics data integration - A conceptual scheme and a bioinformatics pipeline

**DOI:** 10.1101/2020.11.20.391045

**Authors:** Nuria Planell, Vincenzo Lagani, Patricia Sebastian-Leon, Frans van der Kloet, Ewoud Ewing, Nestoras Karathanasis, Arantxa Urdangarin, Imanol Arozarena, Maja Jagodic, Ioannis Tsamardinos, Sonia Tarazona, Ana Conesa, Jesper Tegner, David Gomez-Cabrero

## Abstract

Technologies for profiling samples using different omics platforms have been at the forefront since the human genome project. Large-scale multi-omics data hold the promise of deciphering different regulatory layers. Yet, while there is a myriad of bioinformatics tools, each multi-omics analysis appears to start from scratch with an arbitrary decision over which tools to use and how to combine them. It is therefore an unmet need to conceptualize how to integrate such data and to implement and validate pipelines in different cases. We have designed a conceptual framework (STATegra), aiming it to be as generic as possible for multi-omics analysis, combining machine learning component analysis, non-parametric data combination and a multi-omics exploratory analysis in a step-wise manner. While in several studies we have previously combined those integrative tools, here we provide a systematic description of the STATegra framework and its validation using two TCGA case studies. For both, the Glioblastoma and the Skin Cutaneous Melanoma cases, we demonstrate an enhanced capacity to identify features in comparison to single-omics analysis. Such an integrative multi-omics analysis framework for the identification of features and components facilitates the discovery of new biology. Finally, we provide several options for applying the STATegra framework when parametric assumptions are fulfilled, and for the case when not all the samples are profiled for all omics. The STATegra framework is built using several tools, which are being integrated step-by-step as OpenSource in the STATeg**R**a Bioconductor package https://bioconductor.org/packages/release/bioc/html/STATegra.html.

## Introduction

Computational and experimental developments have enabled the profiling of multiple layers of cell regulation: genome, transcriptome, epigenome, chromatin conformation or metabolome, among many others globally known “omics” (1,2). The development of such technologies was driven by the understanding that a single-omic does not provide enough information to allow dissecting biological mechanisms (3,4). For instance, while specific DNA variations have been linked with multiple diseases, the associated mechanisms are not fully understood (5,6). As a result, multi-omics data-sets are increasingly applied across biological domains such as cancer biology (7–11). Furthermore, single-cell multi-omics analysis (12–15) has just become a reality.

However, from the necessity of multi-omics profiling came the need for multi-omics analysis tools. Thus, integrative approaches are expected to generate significantly more comprehensive insights into the biological systems under study (SuS). There is a myriad of such tools in the literature that may be categorized and classified differently (possibly in complex ways) (3,16–21). While each of the tools is a valuable resource for any multi-omics research, combining them into a *conceptually unified framework* is key. Equally important is the fact that each framework must be as generic as possible. Thus, we introduce the STATegra framework, in which we integrate three multi-omics based approaches *into a single pipeline*: (a) Component Analysis to understand the coordination among omics data-types (22); (b) Non-Parametric Combination analysis to leverage on paired designs in order to increase statistical power (23); and (c) an integrative exploratory analysis (24). Furthermore, this framework may be extended by including additional tools such as network analysis (25,26). We incorporated most of these tools into the STATeg**R**a Bioconductor package to facilitate their use (https://bioconductor.org/packages/release/bioc/html/STATegra.html). The package is continuously being updated and developed. Furthermore, additional tools - as described in the framework - are planned to be incorporated into the Bioconductor package, e.g., the pESCA (27) for multi-omics component analysis and the GeneSetCluster (24) for multi-omics exploratory analysis.

To demonstrate the added value of the STATegra framework, we applied it to two datasets from The Cancer Genome Atlas (TCGA): the glioblastoma data-set (28) and the melanoma data-set (29). We also explored (i) the use of samples for which only a subset of omics profiles is available and (ii) the use of parametric versus non-parametric analysis.

## Materials and Methods

Additional information is included in Supplementary Information and an html-R Markdown document is provided for each data-set in Supplementary Material; each document provides a comprehensive overview of the code used in order to enhance their reproducibility.

### Downloading and preprocessing Data

We selected the Glioblastoma Multiforme (GBM) and the Skin Cutaneous Melanoma (SKCM) data-sets from the Cancer Genome Atlas (TCGA). The level 3 publicly available data for gene expression (gene expression calls), miRNA (miRNA expression calls), and DNA methylation (beta values per CpG, DNAm) were obtained per sample through the NCI’s Genomic Data Commons (GDC) portal (10). The associated metadata for each project were also obtained. Additionally, for the SKCM data-set, a curated metadata generated in a previous TCGA study was also used (29).

#### Glioblastoma multiforme

three data types were downloaded: array-based expression (mRNA) – Affymetrix Human Genome HT U133A, array-based expression (miRNA) – Agilent Microarray, and array-based DNA Methylation (DNAm) - Illumina Human Methylation 450K. The number of available samples differed depending on the omic: mRNA, miRNA, and DNAm profiles are available for 523, 518, and 95 samples, respectively (Supplementary Table 1; Supplementary Figure 2A).

#### Skin Cutaneous Melanoma

three data types were downloaded: RNA-seq-based expression (mRNA) - Illumina HiSeq 2000, miRNA-Seq-based expression (miRNA) - Illumina HiSeq 2000, and array-based DNA Methylation (DNAm) - Illumina Human Methylation 450K. The data from these three omics are available for all the individuals (n=425); however, divergences between the initial date of diagnosis (driving the metadata information) and the TCGA specimen date were identified (Supplementary Figure 1). Consequently, we decided to include only those cases for which specimens were obtained within a 1-year window from diagnosis (n=104).

Supplementary Table 1 describes the characteristics of the two data-sets and the preprocessing steps applied before starting the integrative workflow of multi-omics data. For each individual data type, we conducted an exhaustive exploration assessing the need for data normalization and/or filtering (Supplementary Methods). Metadata is available for GBM and SKCM (summarized in Supplementary Table 2 and described in Supplementary Tables 3 and 4 for a detailed description of the variables). In general, the data provided by TCGA contains information on demographic features (age, gender, race, ethnicity), tumor characteristics (age at diagnosis, primary site of the disease, stage of the neoplasm, prior glioma, ulceration in melanoma, Karnofsky score for GBM, Breslow thickness for SKCM), survival outcome (vital status, days to death, days to last follow-up) and technical processes (batch number, tissue source site - TSS, i.e., centers who collect samples and clinical metadata).

At the end of the preprocessing, numerous matrices, i.e., one matrix per every omics data-type (mRNA, miRNA, DNAm), plus one additional matrix containing the metadata of the samples, compose each data-set (GBM, SKCM). Omics data-type matrices are arranged placing measurements (a.k.a. features) on rows and samples in columns, while metadata matrices include samples as rows and metadata information (e.g., age, gender, etc.) in columns.

### Component Analysis for two data-types (omicsPCA)

To perform joint exploration of data, the two data-types must fulfil the following criteria: (i) each feature must be scaled and (ii) only samples that are common to the two data types can be analyzed. Each feature was mean-centered and then normalized to unit sum of squares (Frobenius normalization). Due to sample availability, component analysis for two data-type matrices were restricted for each analysis for common samples (Supplementary Figure 2A).

Once input data were ready, the two main omicsPCA steps were applied: model selection and subspace recovery. For model selection, we aimed to identify the correct model, which means the exact number of common (shared) components and the number of distinctive components per data-type. We investigated the following methodologies: JIVE (30) (the jive R package), PCA-GCA (31) (RegularizedSCA R package), and pESCA (27) (RpESCA and Rspectra R packages) (Supplementary Table 5 and Supplementary html-R Markdown document).

Finally, the association between metadata and the shared/individual components obtained was assessed using the Kruskal-Wallis test, Spearman’s correlation, or the Cox regression model, depending if the variable of interest was categorical, numerical or time-to-event, respectively.

All analyses were conducted in R (32).

### Non-Parametric Combination for two data-types (omicsNPC)

Non-Parametric Combination (NPC) techniques allow combining statistical evidence (p-values) across data-types to obtain a more precise characterization of the changes associated to the outcome of interest (23).

The above described approach allows to integrate data matrices defined on overlapping sets of samples. Taking advantages of this possibility, we explored the NPC following two strategies: analyzing only common samples or analyzing all available samples (including non-overlapping ones, when applicable).

Importantly, NPC methods require linking the features across data-types. To that end, the relation between mRNA and miRNA and mRNA and methylation were obtained using the *SpidermiR* R package (33) and RGmatch (34), respectively.

In the case of mRNA and miRNA mapping, different versions of annotation were found; we combined the following two: the *miRNAmeConverter* (35) and *anamiR* (36) R packages.

Finally, the NPC may be run using the *omicsNPC* function from the STATegra package using the two data-types, the mapping file (i.e., mRNA - miRNA), and the variables to include in the model (see R-code below) as inputs.

In our analyses, the outcomes of interest were survival for the GBM data-set and the primary site of tumor for the SKCM data-set. Additionally, age was included as a covariable in all the models. Depending on the nature of the outcome of interest the analysis performed during NPC differs; in GBM, the association between each molecular quantity and the time-to-event was assessed through a Cox Regression model (37), and because age is by itself a relevant factor (Supplementary Figure 5), it was treated as a time-varying factor by specifying a time-transform function (37). In SKCM associations between each molecular quantity and the primary site of the tumor were assessed through a differential expression analysis using limma (38) (highlighted lines from the R-code).

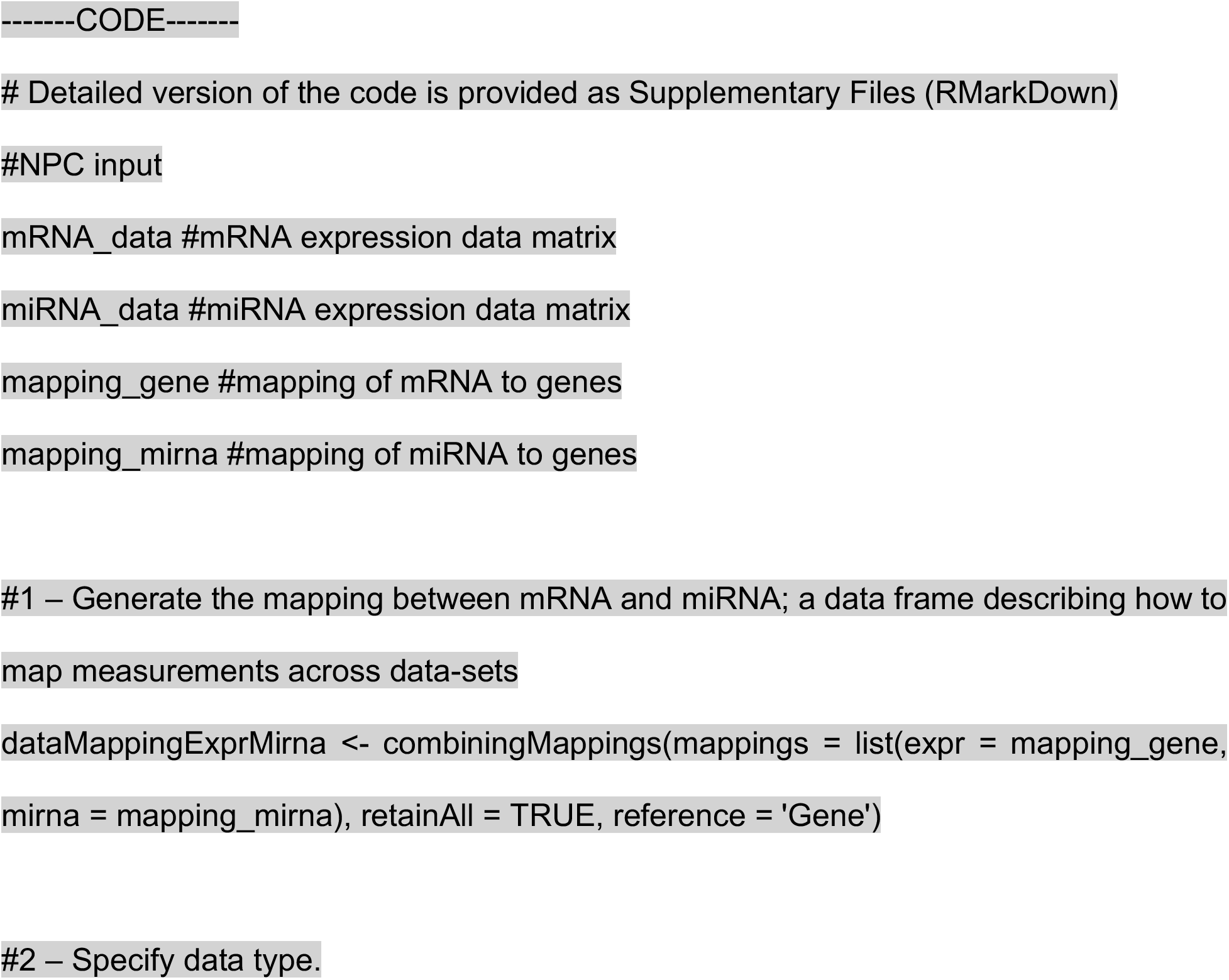
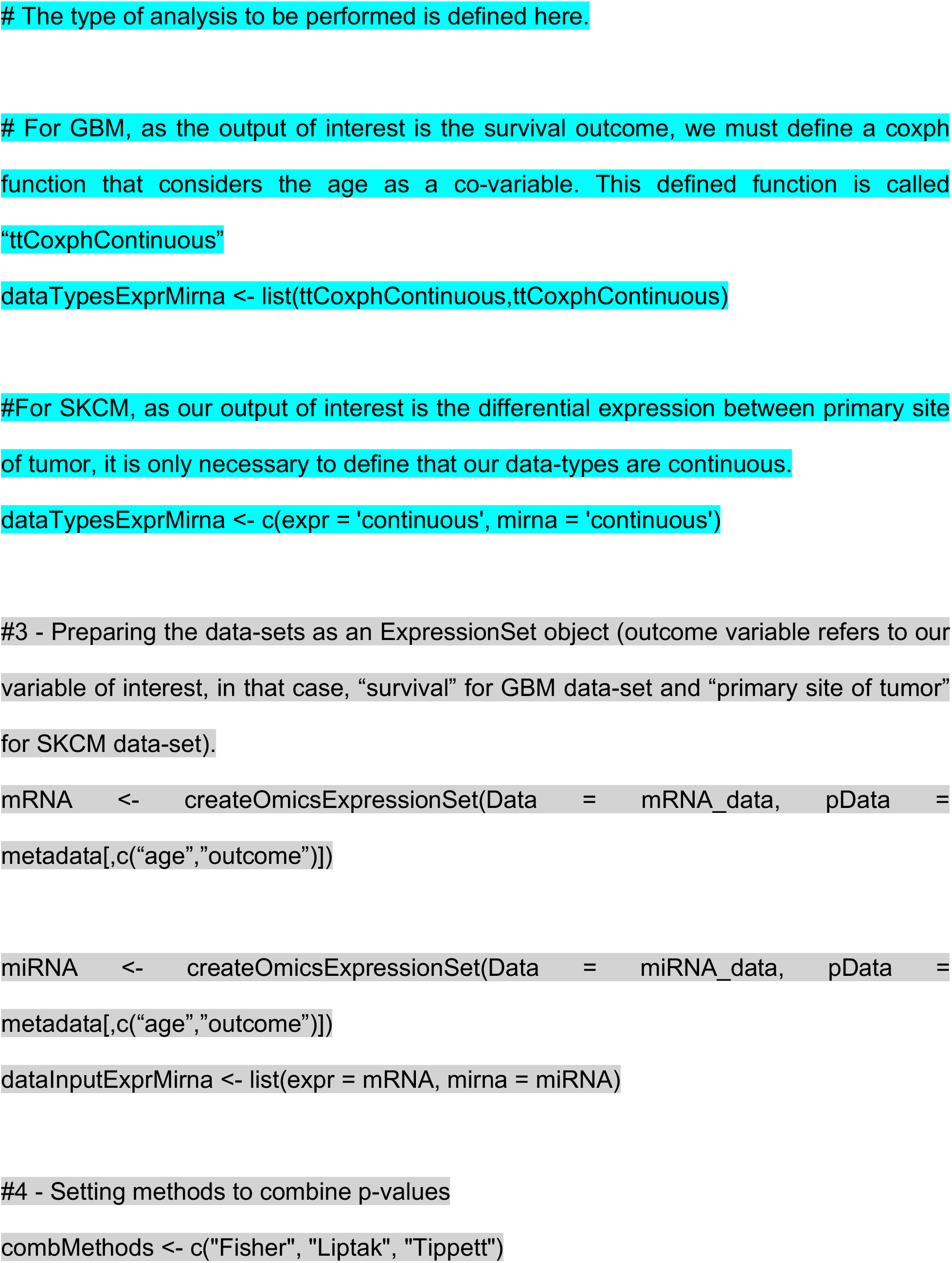
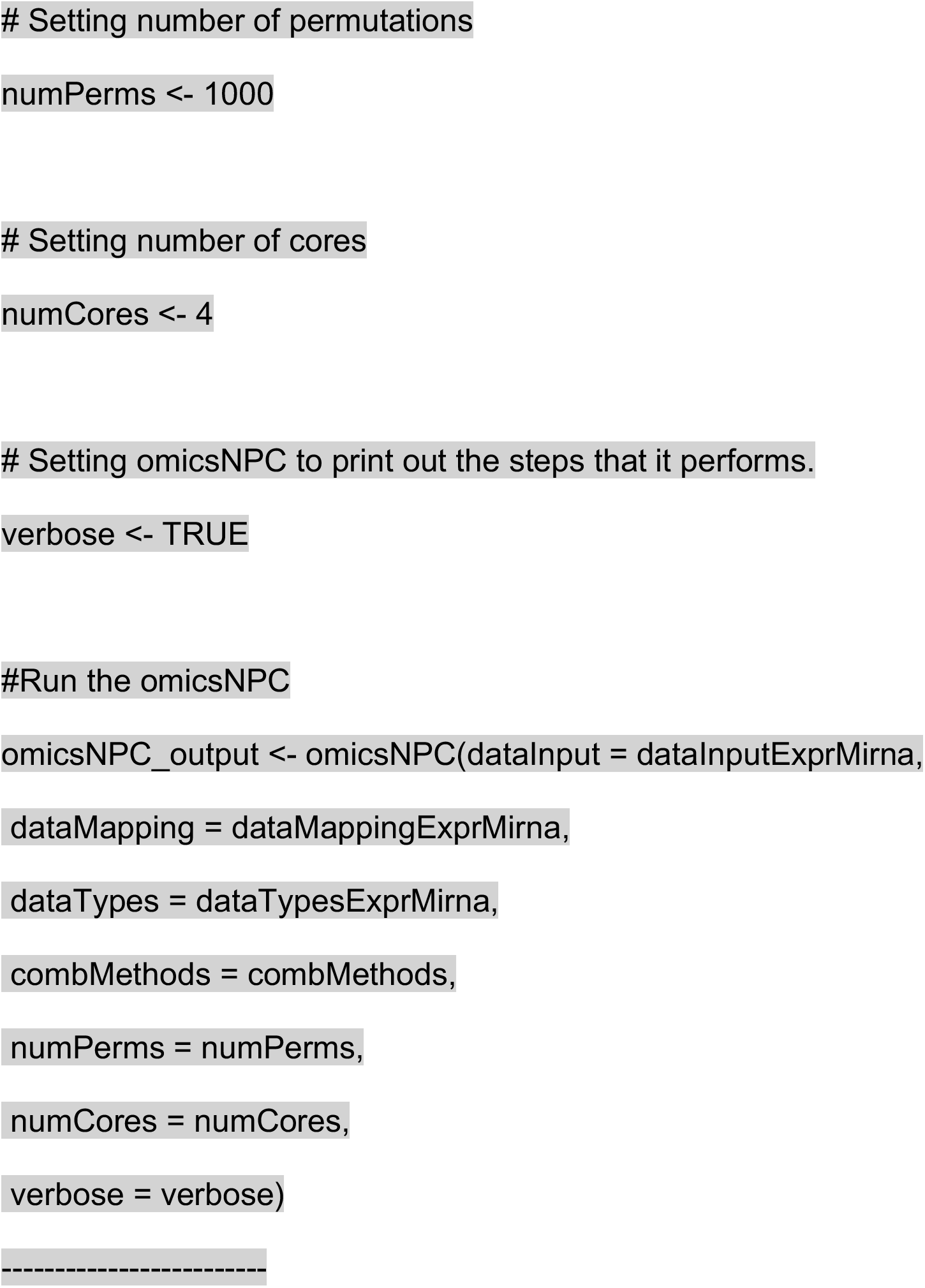

### GeneSetClustering

Significant genes from omicsNPC in the different approaches (Adj.P-value < 0.05 or Fisher p-value < 0.05 in NPC) were uploaded to the Ingenuity Pathway Analysis (IPA, (39)) database (Qiagen), and core expression analysis was performed to identify affected canonical pathways and functional annotations. Right-tailed Fisher’s exact test was used to calculate a p-value. Canonical pathways/functional annotations were clustered together using *GeneSetCluster* (24). Briefly, the gene-sets were grouped into clusters by calculating the similarity of pathways/annotations of the gene content using the relative risk (RR) of each gene-set appearing with each other. Only significant gene-sets (p-values < 0.05) with a minimum of three genes were selected for functional exploration. RR scores were clustered into groups using k-means with the optimal number of genes determined using gap statistics.

## Results

We designed the STATegra framework as a four-step analysis (Figure 1). In the first step, each data-type was analyzed separately using state-of-the-art tools for each omic. Next, in a second step, we explored the shared variability between the different data-types using unsupervised techniques such as *Joint and Individual Variation Explained* (JIVE) (30)), implemented in OmicsPCA. This analysis provided qualitative and quantitative insights into how much the different data-types (e.g., different omics) and their features were “*coordinated*”. Moreover, the analysis provided useful information for targeting specific omics combinations (2). In the third step, for those combinations of omics characterized as *coordinated*, NPC analysis allowed increasing the statistical power to identify significant features as we have recently demonstrated (40,41). For that purpose, we used the NPC within the omicsNPC function (23). In the final step, clustering tools (e.g., OmicsClustering) and gene-set enrichment analysis summarizing tools (such as GeneSetCluster (24)) allowed an integrated approach.

**Figure 1.**
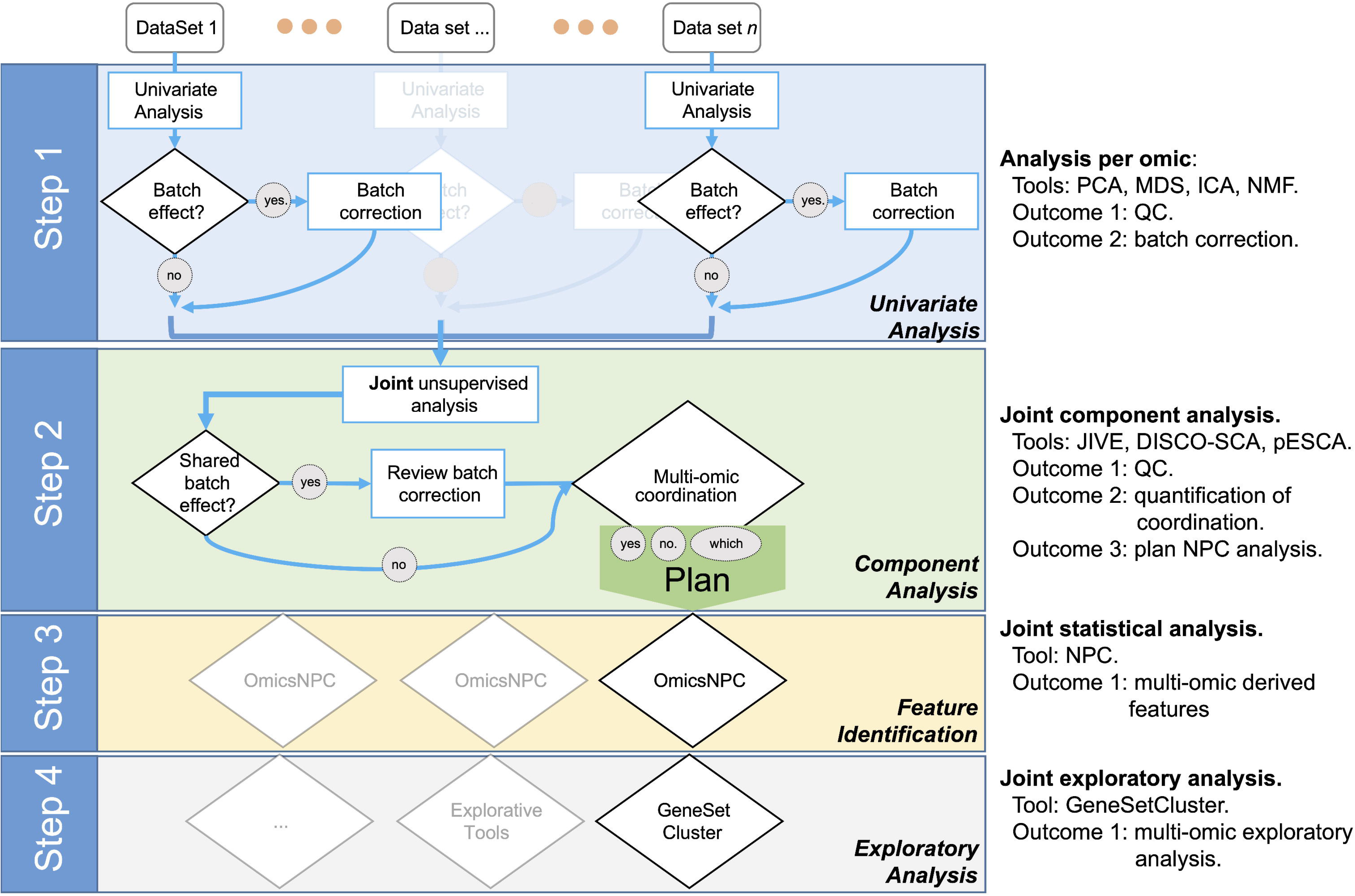
Workflow diagram of the multi-omics analysis framework.

### Selected case studies

We selected two case studies: GBM and SKCM. GBM is the first cancer studied by TCGA (42,43). The TCGA GBM data-set consists of primary tumor samples from roughly 600 cases. The data-set contains gene expression, miRNA, and DNA methylation microarrays. Based on these data, several findings have been reported, including a molecular classification of glioblastoma based on gene expression profiles (classical, proneural, neural, mesenchymal) (44). The landscape of SKCM was published by TCGA in 2015 (29). The TCGA SKCM data-set consists of melanoma samples from patients diagnosed with either primary or metastatic cutaneous melanoma or metastatic melanoma of unknown primary from ~400 cases. The data-set contains genotype information, gene expression, and methylation microarrays. Based on these data, several findings have been reported, including the genomic identification of four mutant subtypes (BRAF hotspot, NF1 mutant, RAS hotspot, and triple wild-type) and a molecular classification based on gene expression profiles (immune, keratin, and MITF-low related profiles) associated with survival time. In general, patients from both studies were Caucasian with a median age of 58-59 years and higher proportion of males (~ 60%). Mortality rate in GBM was high (78%) with a median life expectancy of around one year. For SKCM, 42% of patients died during follow-up and median life expectancy was of one year and three months (Supplementary Tables 1 and 2).

### Step 1: Independent data-type exploration and characterization

Once the data is pre-processed, we recommend conducting quality controls for each individual data-type as the first step in the STATegra framework. In our example we made use of PCA as an unsupervised exploratory analysis, but other matrix-factorization techniques may be used, e.g., Independent Component Analysis (ICA) (45) or Non-negative Matrix Factorization (NMF) (46), among others. It is important to emphasize the relevance of setting up a proper study design to avoid possible batch-effects not to be confounded with the biological effects under study: a component analysis will not overcome a wrong design.

In the case of the GBM data-set, the two first PCA components showed a limited amount of variability explained for all omics (Supplementary Figure 2B), suggesting a large per sample variability. As expected from the original TCGA publication (44), we found a significant association between the previously defined “gene expression subtypes” (44) and the first PCs of mRNA (Bonferroni adjusted p-value < 0.001; refer to Supplementary Methods). Interestingly, such association was also found for miRNA and DNAm (Supplementary Figure 2C; adjusted p-value < 0.005). Moreover, we identified several clinical variables associated with at least one of the first three main components of omics data (refer to Supplementary Methods; Bonferroni adjusted p-value < 0.05): survival outcome (mRNA, miRNA, DNAm) and TSS (mRNA).

In the case of the SKCM data-set, the two first PCA components showed a limited amount of variability explained for all omics (Supplementary Figure 3A). We identified several clinical variables associated with at least one of the first three main components of omics data (refer to Supplementary Methods; Bonferroni adjusted p-value < 0.05): primary site of disease (mRNA, miRNA), neoplasm (mRNA), and pathological stage of the disease (mRNA, miRNA).

It is worth noting that some of the clinical variables were associated with at least one of the first three components in the individual data-type exploration *for more than one omics* data type. Such results apply to both GBM and SKCM data-sets. Consequently, we hypothesize that several omics are coordinated and their analytical integration would bring more statistical power and synergistic insights. In Step 2 we investigated such assumptions.

### Step 2: Joint exploration and characterization

As previously shown, in both selected data-sets several clinical variables were associated with more than one omics data-type. Ssuch observations may indicate that some (if not all) those omics profiles are coordinated (or at least some of their features are). Therefore, the next step in the STATegra framework was to investigate and quantify a potential coordination.

Thus, instead of looking at the PCA-derived components of mRNA and miRNA separately, we investigated the existence of components (or factors) shared by both omics (47). Intuitively, while in PCA we projected using the main components per omic (refer to Supplementary Figures 2B, 3A as examples), we next aimed to identify projections where the components are informative for more than one data-type simultaneously (refer to Figures 2A, C). In summary, when analyzing the variability of data-types A and B, we aimed to identify components associated to both A and B (shared components), components associated only to A, and components associated only to B (distinctive components).

**Figure 2.**
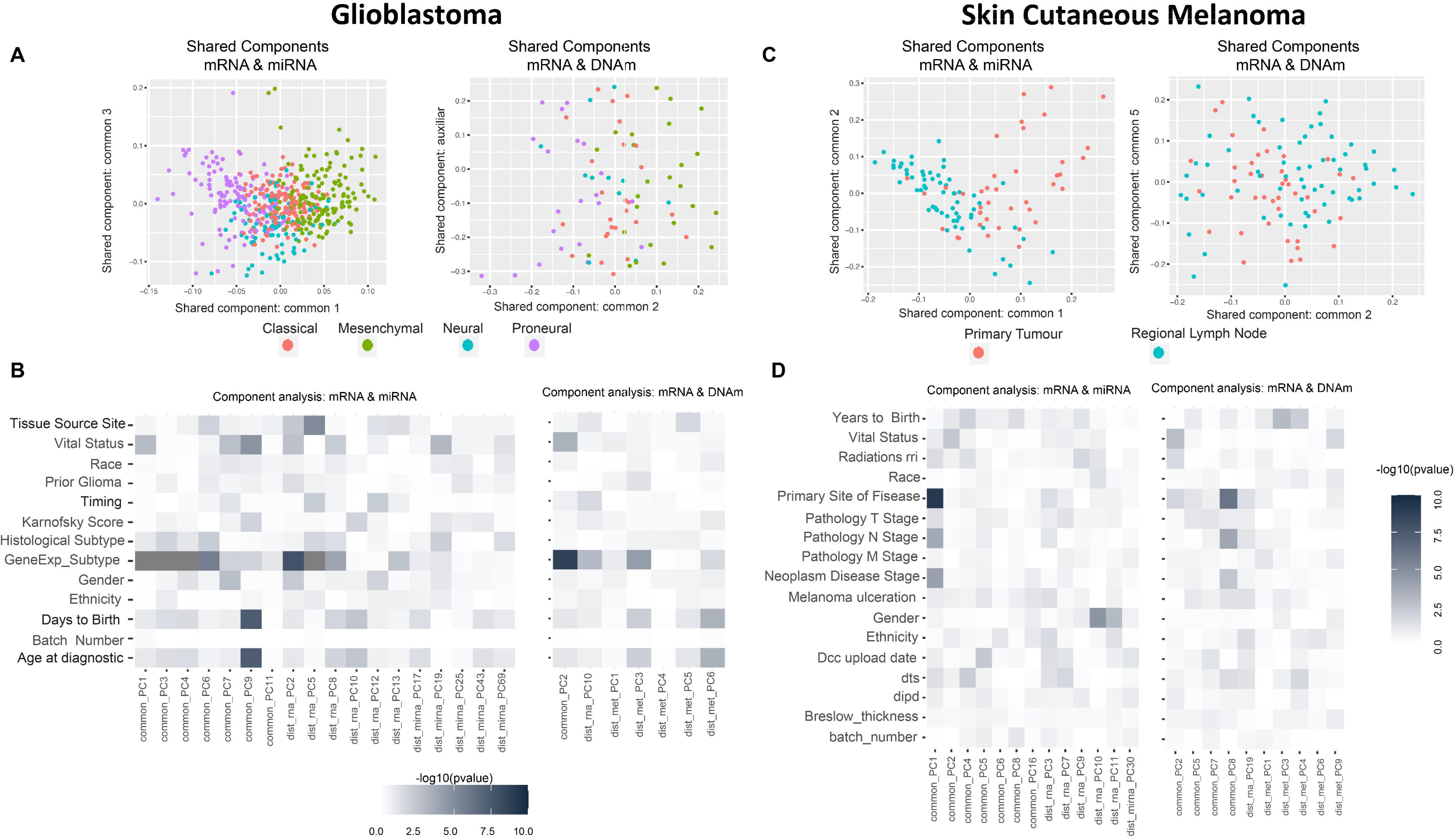
Multi-omics component analysis. (A, C) Component-based representation of GBM and SKCM joint exploration; mRNA + miRNA (left) and mRNA + methylation (right). First and second common components (or auxiliary if only one common component is found) are shown. Samples are colored based on gene expression subtype for GBM and primary site of disease for SKCM. (B, D) Heatmap representation of −log10 (p-values) of the statistical test between metadata and common and distinctive components of GBM and SKCM joint analysis; mRNA + miRNA (left) and mRNA + methylation (right). Color ranges from white to black, understood as p-values with no significance to significant p-values. Based on the nature of the variables, p-values were obtained by association, correlation, or using a survival test. *Radiations rri, dts* and *dipd* denote, respectively, “*Radiations radiation regimen indication*”, “*Days to submitted specimen dx*”, and “*Date of initial pathologic diagnosis*”. See Supplementary Tables 3 and 4 for a detailed description of the variables.

Multi-data-set component analysis methodologies have three key steps: (a) model selection, (b) subspace recovery, and (c) estimation of robustness. In (a) model selection, we aimed to identify the correct model, which means the exact number of common (*shared*) components and the number of *distinctive* components per data-type. The determination of model selection, although fundamental, remains an open question (22,48); hence, no final function has yet been included in the STATegra package. However, we explored several methods (JIVE (30), PCA-GCA (49), and pESCA (27)). Both, *common* and *distinctive* components obtained for each method are summarized in Supplementary Table 5. In our experience, the selected method depends on the nature of the data (as shown in (22)), but we also recommend the use of several methodologies in order to establish more robust insights. While the identification of the best model is *yet* an open challenge, we considered - based on the estimates - using the results from pESCA (27), specifically pESCA (1%). Once the number of *shared* and *distinctive* components were determined, the subspace recovery (identification of loads and scores for the components) should be conducted using the same methodology used for the identification of space. Finally, to address robustness estimation we refer to the method in Måge et al. (22).

In the current data-sets we were prioritizing a gene-centric analysis for both data-sets (GBM and SKCM), therefore, we posed two scenarios; the joint analysis of mRNA and miRNA, and the joint analysis of mRNA and methylation. We acknowledge that there are tools in development for integrating more than two omics; see for instance (50) and its application in Gomez-Cabrero et al. (2).

#### GBM data-set

we identified seven shared components between mRNA and miRNA and one between mRNA and DNAm (refer to Supplementary Table 5). Figure 2A shows two PC score plots; the association between components (shared and distinctive) and clinical variables is shown in Figure 2B. We observed that the shared components were mainly associated (Bonferroni adjusted p-value < 0.05) with: “gene expression subtype” derived from (44) (mRNA-miRNA, mRNA-DNAm), survival outcome (mRNA-miRNA), and age (mRNA-miRNA) (Figure 2B). No significant relationship was seen between gene expression subtype and survival outcome (Supplementary Figure 4, p-value = 0.06), although a relationship between age and survival outcome was observed (adjusted p-value < 0.05). Based on these results, we hypothesized a coordination between the mRNA and miRNA profiles, and such coordination is associated with survival. Consequently, we also considered that the integration of both data types will contribute to increase the knowledge regarding GBM survival. We identified a limited global coordination when considering the mRNA and DNAm profiles.

#### SKCM data-set

seven shared components were identified between mRNA and miRNA profiles, and four common components between mRNA and DNAm profiles (refer to Supplementary Table 5). Figure 2C shows two PC score plots, and the association between components (shared and distinctive) and clinical variables is shown in Figure 2D. The shared components identified are associated with primary site of the disease for both mRNA-miRNA and mRNA-DNAm pairs and the disease stage for the mRNA-miRNA pair (refer to Supplementary Methods; Bonferroni adjusted p-value < 0.05). Based on these results, we concluded that mRNA, miRNA and DNAm are globally coordinated, and this is mainly associated with the primary site of the disease. Therefore, the integration of the three data-types may contribute to increase the knowledge on SKCM primary site. Importantly, based on the *complexity of the data*, the joint exploration may allow datatype specific related batch effects (identified in Step 1) from batch effects associated with sample collection (which will be associated to all omics). Interestingly, more than two omics (*blocks*) can be analyzed to identify shared components (16,27,50).

The next challenge, Step 3, was to leverage the coordination identified among omics to gain statistical power in the identification of the relevant features that explain the SuS.

### Step 3: Integrative differential analysis: omicsNPC

In Step 3 we made use of NPC to increase the statistical power for the analysis of the SuS (51). Briefly, NPC non-parametrically combines p-values from associated features, such as a miRNA and one of its target genes measured on overlapping sets of samples. We used the omicsNPC (23) included in the STATegra package, specifically tailored for the characteristics of omics data.

The main advantages of the NPC include: a) high statistical power with minimal assumptions; b) wide applicability on different study designs; c) it allows integrating data modalities with different encodings, ranges, and data distributions; and d) it models the correlation structures present in the data producing unbiased/calibrated p-values, an interpretable metric (51).

OmicsNPC first analyses each data-type separately through a permutation-based scheme. Currently, omicsNPC uses the package limma or survival (coxph) for computing statistics and p-values; however, the user may also customize the functions (refer to Methods). The resulting permuted-based p-values may be combined using Tippett’s (aimed to identify findings supported by at least one omics modality), Liptak’s (aimed to identify findings supported by most omics modalities), or Fisher’s (intermediate behavior between Tippett and Liptak) combination function. Following the original NPC, omicsNPC (23) makes minimal assumptions: as permutation is employed throughout the process, no parametric form is assumed for the null distribution of the statistical tests, and the main requirement is that samples are freely exchangeable under the null-hypothesis. This frees the researcher from the need of defining and modeling between dataset dependencies. Most importantly, it provides global p-values for assessing the overall association of related features across different data modalities with the specified outcome (51).

#### GBM analysis

we aimed to investigate GBM survival through its relationship with omic features corrected for age, based on the association identified in Supplementary Figure 5. We only made use of samples profiled for all data-types (n=515 and n=83 for the mRNA-miRNA and mRNA-DNAm pairs, respectively). Table 1 (*Overlapping samples* column) presents the NPC outputs. When the NPC is applied on “mRNA and miRNA”, the integration allowed identifying 23 new genes and four new miRNAs. For “mRNA and DNAm”, the integration allowed the identification of 106 new genes and 150 new CpG sites.

**Table 1.** Non-parametric combination analysis results of two-omics data from the GBM and SKCM projects. Significance was considered for a False Discovery Rate < 0.05

|  |  | GBM |  | SKCM |
| --- | --- | --- | --- | --- |
| mRNA + miRNA |  | Overlapping samples | Whole dataset |  |
| mRNA dimension |  | 7,814 x 515 | 7,814 x 523 | 9,491 x 104 |
| mRNA significant |  | **1** | **4** | **216** |
| miRNA dimension |  | 323 x 515 | 323 x 518 | 239 x 104 |
| miRNA significant |  | **1** | **1** | **6** |
| mRNA-miRNA total pairs |  | 24,665 | 24,665 | 20,225 |
| NPC_Fisher significant pairs |  | 27 | 50 | 114 |
| New mRNA from NPC |  | **23** | **43** | **48** |
| New miRNA from NPC |  | **4** | **7** | **14** |
| mRNA + DNAm |  | Overlapping samples | Whole dataset |  |
| mRNA dimension |  | 9,620 x 83 | 9,620 x 523 | 9,564 x 104 |
| mRNA significant |  | **2** | **7** | **277** |
| Methylation dimension |  | 57,645 x 83 | 57,645 x 95 | 55,729 x 104 |
| methylation significant |  | **1** | **0** | **12** |
| mRNA-methylation total pairs |  | 57,645 | 57,645 | 55,729 |
| NPC_Fisher significant pairs |  | 150 | 332 | 432 |
| New mRNA from NPC |  | **106** | **174** | **116** |
| New methylation sites from NPC |  | **150** | **332** | **428** |

#### SKCM analysis

we explored the omics characterization associated to the primary site of the disease. When the NPC was applied on “mRNA and miRNA”, the integration allowed identifying 48 new genes and 14 new miRNAs. For “mRNA and DNAm”, the integration allowed identifying 116 new genes and 428 new CpG sites. This increase of the statistical power was expected based on the results from the joint exploration (Figure 2).

### Alternatives to Step 3

#### Including samples available for a sub-set of data-types

when doing the NPC analysis, we considered samples available for both omics. However, in the case of GBM we discarded a large number of samples. In (23,40) we modified the NPC permutation protocol to include the discarded samples. We observed that the use of all samples allowed us to identify a larger number of novel features (“mRNA and miRNA” identified 43 new genes instead of 23; for complete results refer to Table 1, Column *Whole dataset*).

#### Parametric version

The NPC requires a large number of permutations, which is time consuming. To address this, the STATegra package includes a parametric combination methodology (23,52). This parametric approach is a faster alternative to NPC, which we suggest to use in preliminary explorations. In our analyses, the parametric approach generated a larger number of significant results in comparison to the non-parametric counterpart (Supplementary Table 6), which may be explained by unaccounted interdata-sets correlations that inflate the significance of the p-values.

### Step 4: Exploratory analysis and determination of the framework’s added value

The STATegra framework provided novel genes, miRNAs, and CpG sites for the two selected cases in comparison to unimodal analyses. We investigated if such novel elements could also provide new insights at gene-set level. For this, we made use of the GeneSetCluster (24), a tool that summarizes gene-set analysis (GSA) results derived from multiple analyses. It allows identifying core-results by clustering gene-sets and posterior exploration; furthermore, it analyzes the integration of more than one gene-set (which could be derived from more than one omic) simultaneously. When investigating SKCM, we compared three GSA: (1) using genes derived from mRNA single-omic analysis, (2) using genes derived from mRNA-miRNA NPC analysis, and (3) genes in (2) not identified in (1). We observed that the set of genes in (1) identified several relevant canonical pathways, which are also identified in (2) and (3); but, especially, (2) and (3) GSA identified many additional relevant pathways as shown in Figure 3B for Canonical Pathways analysis (see box strokes on clusters). In the case of GBM, four GSAs were conducted with the following pair combinations: (a) “*considering only samples with all omics available* (OVERLAP)” or “*considering all samples* (ALL)”, and (b) “*considering all identified genes*” or “*considering genes only identified by NPC*”. We observed major differences in the summarized gene-sets between OVERLAP vs ALL; see for instance Figure 3A, when analyzing “Gene Ontology - Biological Functions” (53). The use of GeneSetCluster allowed us to demonstrate the added value of the STATegra framework. Furthermore, it is also a tool for multi-omics GSA integrative analysis that we consider as part of the STATegra framework and we plan to integrate further in the STATeg**R**a package.

**Figure 3.**
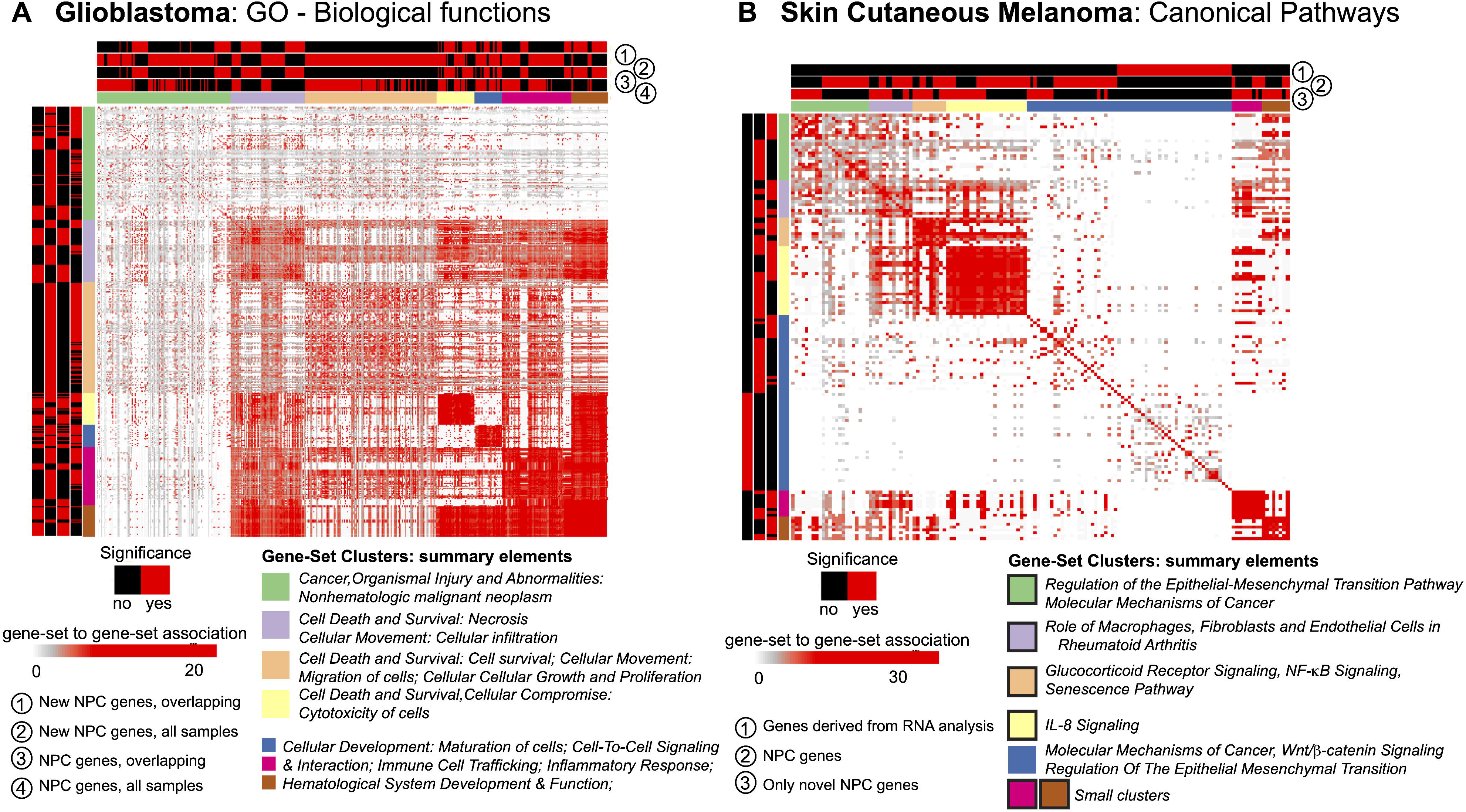
GeneSetCluster analysis. Each heatmap depicts the gene-set to gene-set RR distance (24). In each case several gene-set analyses have been conducted. Red(yes)/black(no) shows which gene-sets have been identified for each gene-set analysis. (A) GBM and (B) SKCM. In SKCM, clusters presented as black lines are those associated to discoveries through multi-omics integration.

## Discussion

There are many bioinformatics integrative tools (16,26,54–56). However, when carrying out multi-omics analysis, as a rule, researchers use a custom pipeline that combines part of the available tools. While every multi-omics data combination is different, we believe that a general framework is key to gain knowledge for an “*optimized*” integrated research analysis in the future. We here present the STATegra framework, a multi-omics integrative pipeline, the result of integrative analyses done over the last decade (23,24,40,41,57). In the two chosen case studies used to evaluate the STATegra framework, GBM and SKCM, we show through a consecutive four-step process (Figure 1), how single omics integration generates additional information. Step 2, Component Analysis, quantifies the coordination of the different data-types, a key phase to identify where omic-combination can be leveraged, and Step 3 -Non-Parametric Combination-is used to gain statistical power. In both case studies, we detect a greater number of genes. For example, the miRNAs and CpG sites are identified with our framework. Step 4 examines the added value of the biological-insights of the features identified by the integration process.

In GBM we examine the association of the omics profiles with survival. In comparison to single-omic analysis, the STATegra framework identifies new genes such as CAST, ATF5, GANAB (glycoprotein associated with GBM cancer stem cells (58)), ICAM (overexpressed in bevacizumab-resistant GBM (59)), CORO1A (upregulated in GBM (60)), LYN (*in vitro* association of enhanced survival of GBM cells (61)), MET (protooncogene) and STAT5 (enhances GBM cells migration, survival (62), and proliferation (63)), among others. Most have been previously associated with cancer and particularly to glioblastoma. We also compare the identified miRNAs with existing miRNA-derived survival signatures (64); only miR222 is identified in the single-omic analysis, while three additional miRNAs (miR31, miR221, and miR200b) are identified by STATegra.

With the analysis of GSA, STATegra identifies new gene-sets, e.g., the TREM1 signaling pathway, previously associated with GBM (65). In SKCM we investigated the omics association with the primary site of disease. In addition to the newly identified genes (refer to Table 1), the major STATegra-associated novel insights are derived from GSA analysis as shown in Figure 3B, particularly regarding the identification of the IL8 signaling, which is known to be relevant in SKCM (66,67).

It is important to point out that we are not comparing our analysis against the original publications: GBM (42,43) and SKCM (29).The idea is to compare a generic framework with single-omic approaches. Moreover, since the questions and data-sets used are different from those in the original TCGA publications, a back-to-back comparison is not justified.

The results generated by STATegra show the *added value* of a general integrative framework. Still, we acknowledge that, similarly to Operations Research there is “*no-free-lunch*” (68). Generic frameworks provide an initial approximation to any integrative analysis, and once completed, they may be further customized - and therefore further optimized - to account for the characteristics of the data and considered SuS. Still, the value of the STATegra framework is its solid integration starting point, and - after being applied in many projects -generic rules can be extracted to allow an easier and faster customization.

Frameworks as the one we present here are becoming increasingly necessary due to the amount of growing multi-omics data, particularly in the context of single-cell multi-omics (15). Further developments are required in multi-omics visualization (69), simulated data (70), or further exploitation of Component Analysis as shown in (17), among others. Thus, we consider that the STATegra framework is the starting point that will be further developed over time. The next immediate steps are the inclusion of pESCA (27) for multi-omic component analysis and GeneSetCluster (24) for multi-omic exploratory analysis within the STATeg**R**a Bioconductor package.

## Supporting information

RMarkDown and html files.

