## Supplementary Information for "STATegra: Multi-omics data integration - A conceptual scheme and a bioinformatics pipeline"

### Supplementary Material


1. Supplementary methods 2

**A) Accessing to data from the Cancer Genome Atlas** **2**

**B) Analysis per data-type** **2**

2. Supplementary Tables 4

3. Supplementary Figures 6

4. References ERROR! BOOKMARK NOT DEFINED.

### 1. Supplementary Methods

We are making the effort to implement an actionable and traceable code. To such end, most of the analyses conducted in this paper are shared in an R-MarkDown HTML format in two files, one for each case study:

- Glioblastoma Multiforme (GBM): stategra_gbm_pipeline.html
- Skin Cutaneous Melanoma (SKCM): stategra_skcm_pipeline.html

#### A) Accessing to data from the Cancer Genome Atlas

Two public data-sets from the Cancer Genome Atlas (TCGA) were used, GBM and SKCM. The level 3 public data for gene expression (expression calls for gene, per sample), miRNA (expression calls for miRNA, per sample), and methylation (beta values per CpG, per sample) were downloaded using the NCI's Genomic Data Commons portal (1). The metadata associated to each project were also obtained. Additionally, for the SKCM project, a curated metadata generated in a previous TCGA study was also used (2).

#### B) Analysis per data-type

**Normalization**

Normalized data from each individual omic was obtained and explored through density plots to assess data distribution, check data normalization, and identify if any kind of data transformation or normalization was required (i.e., logarithmic or logit). Technical replicate samples and samples associated to a unique batch were removed from the data-types. Furthermore, in methylation data-types, all probes with missing values, those related with chromosome Y or X, probes with SNPs, CpG, or SBE in target sequences, and technical control probes were removed (Supplementary Table 1).

#### Principal Component Analysis and Batch Correction

Principal component analysis (PCA) was used to explore the general variability of the data and identify outliers or batch effects. The ComBat method was used to remove batch effects from the different data-types; the parametric approach for mRNA and miRNA data-types and the "non-parametric approach" for the DNA methylation data-type (3,4). Refer to Supplementary Table 1 and Supplementary Figures 2 and 3.

#### Association between Principal Components and Clinical Variables

The association between metadata and principal components was assessed using the Kruskal-Wallis test, Spearman´s correlation, or Cox regression depending if the clinical variable was categorical, numerical, or a survival measure, respectively.

All the analysis were done in R (5)

#### 2. Supplementary Tables

**Supplementary Table 1. Omics data types description for the Glioblastoma Multiforme and Skin Cutaneous Melanoma projects from TCGA**. Type and dimension of level three data available from TCGA, pre-process steps prior integrative pipeline (filter, transformation, and batch correction), and final data dimension are shown. *Non-Parametric ComBat was applied for DNA methylation data because of the low quality model fitting obtained from the parametric ComBat approach

**Supplementary Table 2. Glioblastoma Multiform and Skin Cutaneous Melanoma metadata.** A selection of the most relevant clinical and epidemiological variables are described. Additional information regarding collected variables can be found in the Genomic Data Commons Data Portal. Absolut frequencies are shown for categorical variables and median and interquartile ranges for quantitative variables. †Cases with unclassified gene expression subtype were excluded from the analysis (n=67)

**Supplementary Table 3. GBM Metadata.**

**Supplementary Table 4. SKCM Metadata.**

**Supplementary Table 5. Model Selection Component Analysis: after batch correction.** Model Selection for dimension reduction techniques in multi-omics data. Number of common and distinctive components identified by three different methods are shown: JIVE (r.JIVE), PCA-GCA (cut-off 1% and 5% of explained variance), and pESCA (cut-off 1% and 5% of explained variance). Output for the joint analysis of mRNA and miRNA, and mRNA and methylation for both the Glioblastoma Multiform and Skin Cutaneous Melanoma projects, are reported. Missing values are obtained when no components reach the defined cut-off

**Supplementary Table 6. Parametric Combination Summary table.** Parametric Combination of two-omics data from the Glioblastoma Multiform and Skin Cutaneous Melanoma projects. Significance was considered for a False Discovery Rate < 0.05

**Supplementary Figures**

**Supplementary Figure 1. Relationship between age at initial diagnosis of the pathology (years) and age at TCGA specimen (years).** Reference line in red, where x=y

TCGA = The Cancer Genome Atlas

**Supplementary Figure 2. Glioblastoma Multiforme exploratory analysis.** (A) Venn diagram showing the overlapping cases between the evaluated GBM data types. (B) PCA representation of GBM individual data-types. First and second principal components are shown. Samples are colored according to gene expression subtype. (C) Heatmap representation of –log10 (p-values) of the statistical test between metadata and principal components of GBM data-types. Color ranges from white to black, understood as p-values with no significance to very significant p-values (max value=10). Based on the nature of the variables, p-values have been obtained through association, correlation, or a survival test

GMB = Glioblastoma Multiforme; PCA = Principal Component Analysis

**Supplementary Figure 3. Skin Cutaneous Melanoma exploratory analysis.** (A) PCA representation of SKCM individual data-sets (n=104). First and second principal components are shown. Samples are colored based on the primary site of the disease. (B) Heatmap representation of –log10 (p-values) of the statistical test between metadata and principal components of SKCM data. Color ranges from white to black, understood as p-values with no significance to very significant p-values (max value=10). Based on the nature of the variables, p-values have been obtained through association, correlation, or a survival test

SKCM = Skin Cutaneous Melanoma; PCA = Principal Component Analysis

**Supplementary Figure 4. Association of meta-data for Glioblastoma Multiforme.** Association analysis for metadata of GBM study. Heatmap representation of –log10 (p-values) of the statistical test between two variables. Color ranges from white to black, understood as p-values with no significance to very significant p-values (max value=10). Based on the nature of the variables, p-values have been obtained through association, correlation, or a survival test

GBM = Glioblastoma Multiforme

**Supplementary Figure 5**. **Glioblastoma Multiforme survival curve**. Survival curve for GBM individuals by age (young: age < 60 years old versus old: age ≥ 60 years old)

GBM = Glioblastoma Multiforme
