## Supplementary Figures for "STATegra: Multi-omics data integration - A conceptual scheme and a bioinformatics pipeline"

### Supplementary Tables

STATegRa manuscript

Supplementary Table 1.

| | Technology | Initial Data Dimension | Type | Filters | Transformation | Batch Correction | Final Data Dimension |
|---|---|---|---|---|---|---|---|
| **Glioblastoma Multiforme** | | | | | | | |
| **mRNA** | Affymetrix Human Genome HT U133A | 12,042 x 548 | Normalized log2-Expression signal | • Remove replicated samples<br>• Remove samples associated to a unique batch (1 batch – 1 samples) | No | Parametric ComBat | 12,042 x 523 |
| **miRNA** | Agilent Microarray | 534 x 571 | Normalized log2-expression signal | • Remove replicated samples<br>• Remove samples associated to a unique batch (1 batch – 1 samples) | No | Parametric ComBat | 534 x 518 |
| **Methylation** | Illumina Human Methylation 450K | 485,577 x 154 | Beta values | • Remove features with missing values<br>• Remove probes related with chromosome Y or X<br>• Remove technical control probes<br>• Remove probes with SNPs, CpG or SBE in target sequence<br>• Remove samples without metadata information | Logit-transformed (M values) | Non-Parametric ComBat * | 311,135 x 95 |
| **Skin Cutaneous Melanoma** | | | | | | | |
| **mRNA** | Illumina HiSeq 2000 | 20,531 x 428 | RSME + UQ normalization | • Remove replicated samples<br>• Remove features with 0 in all samples<br>• Remove samples taken after 1 year of diagnosis<br>• Remove cases different of “Primary Tumor” or “Regional Lymph Node” | Log 2-transformed | Parametric ComBat | 20,225 x 104 |
| **miRNA** | Illumina HiSeq 2000 | 1,046 x 428 | RPM | • Remove features with missing values<br>• Remove features with 0 in all samples<br>• Remove samples taken after 1 year of diagnosis<br>• Remove cases different of “Primary Tumor” or “Regional Lymph Node” | Log 2-transformed | Parametric ComBat | 898 x 104 |
| **Methylation** | Illumina Human Methylation 450K | 485,577 x 428 | Beta values | • Remove features with missing values<br>• Remove probes related with chromosome Y or X<br>• Remove technical control probes<br>• Remove probes with SNPs, CpG or SBE in target sequence<br>• Remove samples taken after 1 year of diagnosis<br>• Remove cases different of “Primary Tumor” or “Regional Lymph Node” | Logit-transformed (M values) | Non-Parametric ComBat * | 305,180 x 104 |

### Supplementary Table 2.

| | Glioblastoma Multiforme | | Skin Cutaneous Melanoma | |
|---|---|---|---|---|
| N | 541 | Missings | 104 | Missings |
| Gender | | | | |
| Male | 327 | | 62 (59.6) | |
| Female | 214 | | 42 (40.4) | |
| Age (years) | 59 [50-69] | | 58 [51-71] | |
| Age at diagnosis (years) | 59 [49-68] | | 57 [50-71] | |
| Race | | 20 | | |
| White | 476 | | 100 (96.2) | |
| Black or African American | 32 | | 0 (0.0) | |
| Asian | 13 | | 4 (3.8) | |
| Ethnicity | | 78 | | 2 |
| Hispanic or Latino | 12 | | 2 (1.9) | |
| Not Hispanic or Latino | 451 | | 100 (96.2) | |
| Vital Status | | | | |
| Alive | 116 | | 60 (57.7) | |
| Dead | 425 | | 44 (42.3) | |
| Days to death | 372 [165-596] | | 468 [322-738] | 61 |
| Days to last follow-up | 302 [143-513] | | 560 [293-1190] | 34 |
| Gene Expression Subtype† | | | NA | |
| Classical | 149 | | NA | |
| Mesenchymal | 164 | | NA | |
| Neural | 89 | | NA | |
| Proneural | 139 | | NA | |
| Histological Subtype | | | NA | |
| Glioblastoma Multiforme (GBM) | 6 | | NA | |
| Treated primary GBM | 20 | | NA | |
| Untreated primary (de novo) GBM | 515 | | NA | |
| Prior Glioma | | | NA | |
| Yes | 15 | | NA | |
| No | 526 | | NA | |
| Karnofsky Score | 80 [70 - 80] | | NA | |
| Timing (205 NA's) | | 205 | NA | |
| Post-Adjuvant Therapy | 66 | | NA | |
| Pre-Adjuvant Therapy | 123 | | NA | |
| Pre-Operative | 122 | | NA | |
| Other | 25 | | NA | |
| Primary site of disease | NA | | | |
| Distant Metastasis | NA | | 0 (0.0) | |
| Primary Tumor | NA | | 41 (39.4) | |
| Regional cutaneous or subcutaneous tissue | NA | | 0 (0.0) | |
| Regional lymph node | NA | | 63 (60.6) | |
| Melanoma ulceration | NA | | | 26 |
| Yes | NA | | 54 (51.9) | |
| No | NA | | 24 (23.1) | |
| Melanoma primary location know | NA | | | |
| Yes | NA | | 85 (81.7) | |
| No | NA | | 19 (18.3) | |

Supplementary Table 3.

| Variable | Description |
|---|---|
| Gene Expression Subtype | GBM molecular classification: Classical / Mesenchymal / Neural / Proneural / Unclassified |
| Gender | The collection of behaviors and attitudes that distinguish people on the basis of the societal roles expected for the two sexes. Male / Female |
| Race | A classification of humans characterized by certain heritable traits, common history, nationality, or geographic distribution.<br>White / black or african American / Asian |
| Ethnicity | A socially defined category of people based on common ancestral, cultural, biological, and social factors. Hispanic or latino / not hispanic or latino |
| Days to birth | Time interval from a person's date of birth to the date of initial pathologic diagnosis, represented as a calculated negative number of days. |
| Age at diagnostic | The age in years of the case at the initial pathological diagnosis of disease or cancer. |
| Vital status | The state of being living or deceased for cases that are part of the investigation. Alive / Dead |
| Days to death | The number of days from the date of the initial pathological diagnosis to the date of death for the case in the investigation. |
| Histological type | Glioblastoma Multiforme (GBM) / Treated primary GBM /Untreated primary (de novo) GBM |
| Prior glioma | Patient's history of prior cancer diagnosis |
| Tissue Source Site (TSS) | Centers who collects samples (tissue, cell or blood) and clinical metadata, which are then sent to a BCR. A TSS is identified by its TSS ID. Number of different TSS: 21 |
| Karnofsky Score | An index designed for classifying patients 16 years of age or older by their functional impairment. A standard way of measuring the ability of cancer patients to perform ordinary tasks. |
| Timing | A time reference for the Karnofsky score and/or the ECOG score using the defined categories.<br>Post-Adjuvant Therapy / Pre-Adjuvant Therapy / Pre-Operative / Other |
| Batch number | A set of related analytes prepared for further analysis, numbered sequentially, from the same disease. Once a Case has been assigned to a batch, subsequent shipments from that case are assigned the same batch number as the original. Number of different batches: 24 |
| Days to last follow up | The number of days from the date of the initial pathological diagnosis to the date of last follow up for the case in the investigation. |

Supplementary Table 4.

| Variable | Description |
|---|---|
| Years to birth | Numeric value to represent the calendar year in which an individual was born. |
| Vital status | The state of being living or deceased for cases that are part of the investigation. Alive / Dead |
| Days to death | The number of days from the date of the initial pathological diagnosis to the date of death for the case in the investigation. |
| Days to last follow-up | Number of days between the date used for index and the date the patient was seen or contacted at follow-up. |
| Days to submitted specimen dx | The age in years of the case at the initial pathological diagnosis of disease or cancer. |
| Primary site of disease | The anatomical site where the primary tumor is located in the organism.<br>Distant Metastasis / Primary Tumor / Regional cutaneous or subcutaneous tissue / Regional lymph node |
| Neoplasm disease stage | Stage of neoplasma. i or ii nos / stage 0 / stage i / stage ia / stage ib / stage ii / stage iia / stage iib / stage iic / stage iii / stage iiia / stage iiib / stage iiic / stage iv |
| Pathology T stage | The T category describes the original (primary) tumor. t0 / t1 / t1a / t1b / t2 / t2a / t2b / t3 / t3a / t3b / t4 / t4a / t4b / tis / tx |
| Pathology N stage | The N category describes whether or not the cancer has reached nearby lymph nodes. n0 / n1 / n1a / n1b / n2 / n2a / n2b / n2c / n3 / nx |
| Pathology M stage | The M category tells whether there are distant metastases (spread of cancer to other parts of the body). m0 / m1 / m1a / m1b / m1c |
| Melanoma ulceration | Presence of ulcers in melanoma. Yes / No |
| dcc_upload_date | Date of data upload |
| Breslow thickness | Thickness of primary tumor at initial diagnosis (mm). |
| Gender | The collection of behaviors and attitudes that distinguish people on the basis of the societal roles expected for the two sexes. Male / Female |
| Date of initial pathologic diagnosis | Date of initial pathologic diagnosis. |
| Radiations radiation regimen indication | Radiarion regime indicated. Yes / No |
| Race | A classification of humans characterized by certain heritable traits, common history, nationality, or geographic distribution.<br>White / black or african American / Asian |
| Ethnicity | A socially defined category of people based on common ancestral, cultural, biological, and social factors. Hispanic or latino / not hispanic or latino |
| Batch number | A set of related analytes prepared for further analysis, numbered sequentially, from the same disease. Once a Case has been assigned to a batch, subsequent shipments from that case are assigned the same batch number as the original. Number of different batches: 17 |

Supplementary Table 5.

| | Component | r.JIVE | PCA-GCA (1%) | PCA-GCA (5%) | pESCA (1%) | pESCA (5%) |
|---|---|---|---|---|---|---|
| **Glioblastoma** | | | | | | |
| **mRNA + miRNA** | | | | | | |
| | Common | 1 | 3 | NA | 7 | 1 |
| | Dist mRNA | 50 | 8 | NA | 6 | 4 |
| | Dist miRNA | 19 | 5 | NA | 14 | 1 |
| **mRNA + Methylation** | | | | | | |
| | Common | 3 | NA | NA | 1 | 1 |
| | Dist mRNA | 17 | NA | NA | 1 | 1 |
| | Dist methylation | 25 | NA | NA | 8 | 2 |
| **Skin Cutaneous Melanoma** | | | | | | |
| **mRNA + miRNA** | | | | | | |
| | Common | 3 | 50 | 1 | 7 | 2 |
| | Dist mRNA | 20 | 15 | 1 | 12 | 2 |
| | Dist miRNA | 17 | 13 | 2 | 2 | 3 |
| **mRNA + Methylation** | | | | | | |
| | Common | 4 | 65 | 1 | 4 | 1 |
| | Dist mRNA | 19 | 0 | 1 | 1 | 3 |
| | Dist methylation | 36 | 38 | 3 | 13 | 2 |

Supplementary Table 6.

| | GBM | | SKCM |
|---|---|---|---|
| **mRNA + miRNA** | Overlapping samples | Whole dataset | |
| mRNA dimension | 7,814 x 515 | 7,814 x 523 | 9,491 x 104 |
| mRNA significant | **1** | **4** | **216** |
| miRNA dimension | 323 x 515 | 323 x 518 | 239 x 104 |
| miRNA significant | **1** | **1** | **6** |
| mRNA-miRNA total pairs | 24,665 | 24,665 | 20,225 |
| NPC_Fisher significant pairs | 397 | 466 | 529 |
| New mRNA from NPC | **337** | **382** | **190** |
| New miRNA from NPC | **45** | **54** | **77** |
| **mRNA + methylation** | | | |
| mRNA dimension | 9,620 x 83 | 9,620 x 523 | 9,564 x 104 |
| mRNA significant | **2** | **7** | **216** |
| Methylation dimension | 57,645 x 83 | 57,645 x 95 | 55,729 x 104 |
| methylation significant | **1** | **0** | **12** |
| mRNA-methylation total pairs | 57,645 | 57,645 | 55,729 |
| NPC_Fisher significant pairs | 323 | 207 | 1,418 |
| New mRNA from NPC | **179** | **89** | **245** |
| New methylation sites from NPC | **322** | **207** | **1,406** |

### Supplementary Figures

STATegRa manuscript

Supplementary Figure 1.

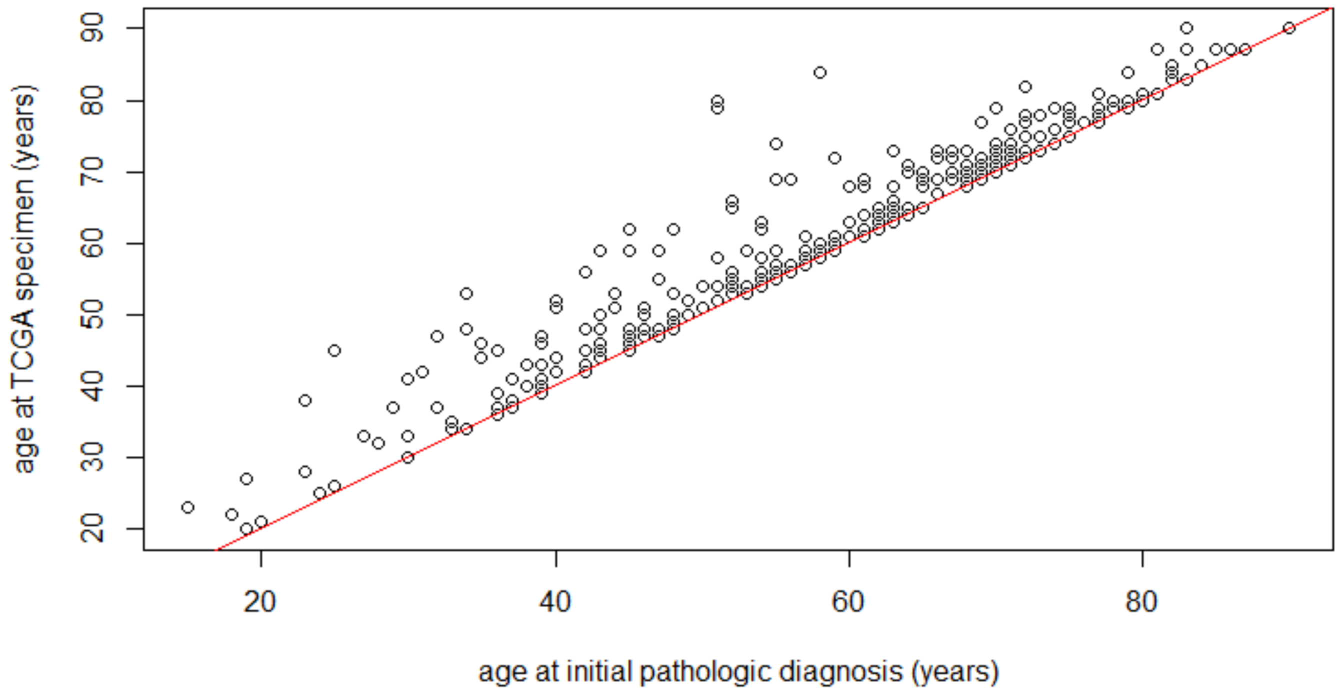

Supplementary Fig 2.

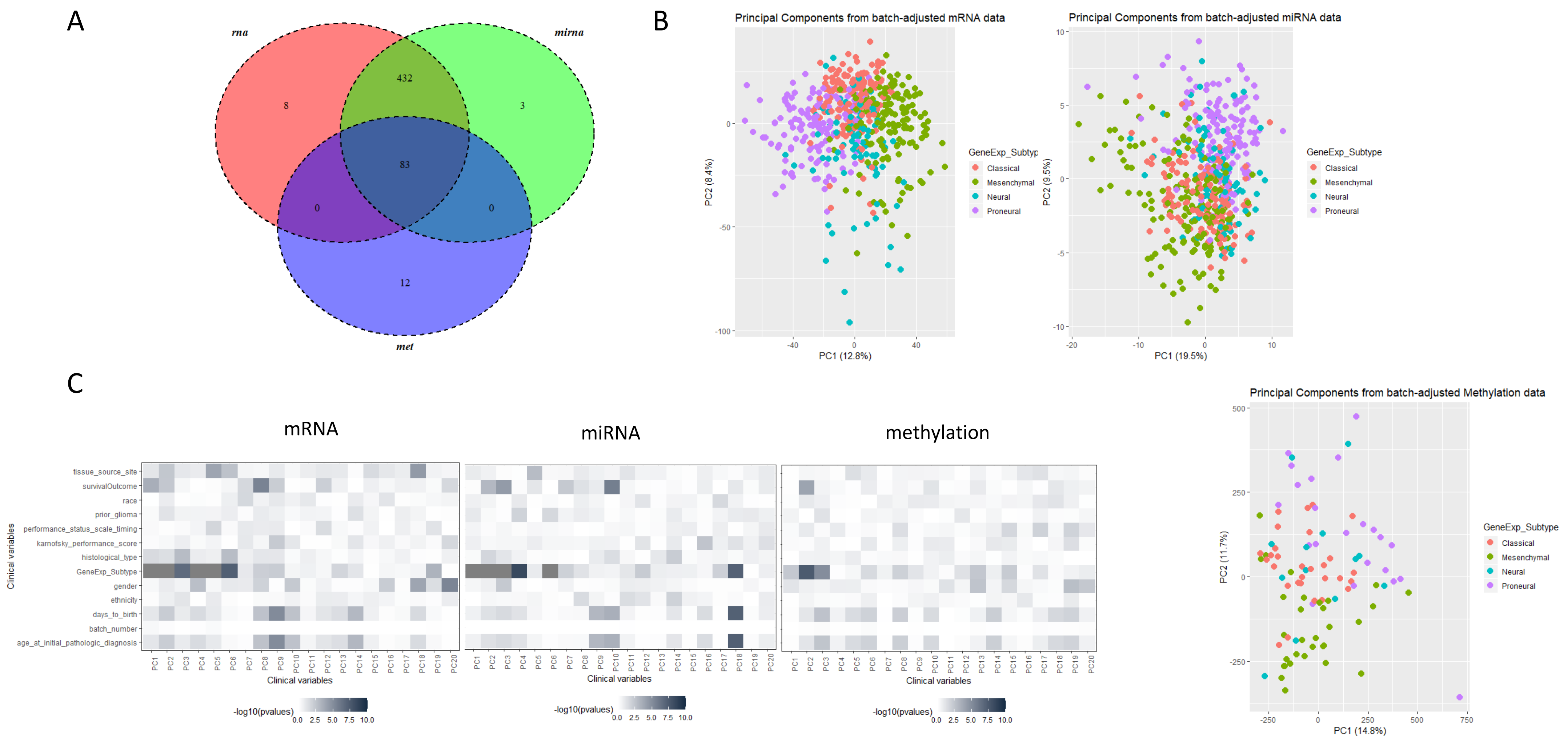

Supplementary Fig 3.

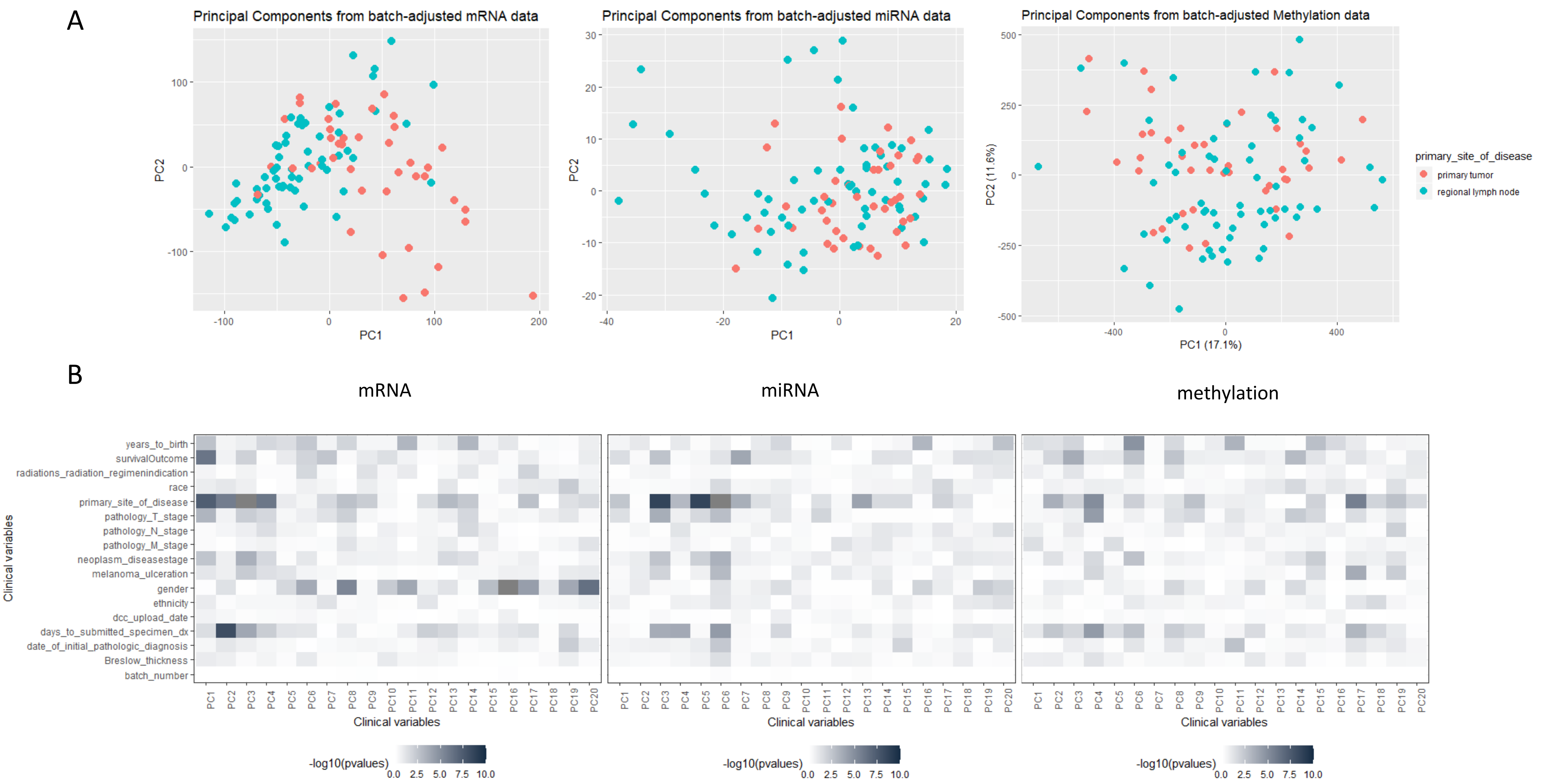

Supplementary Figure 4.

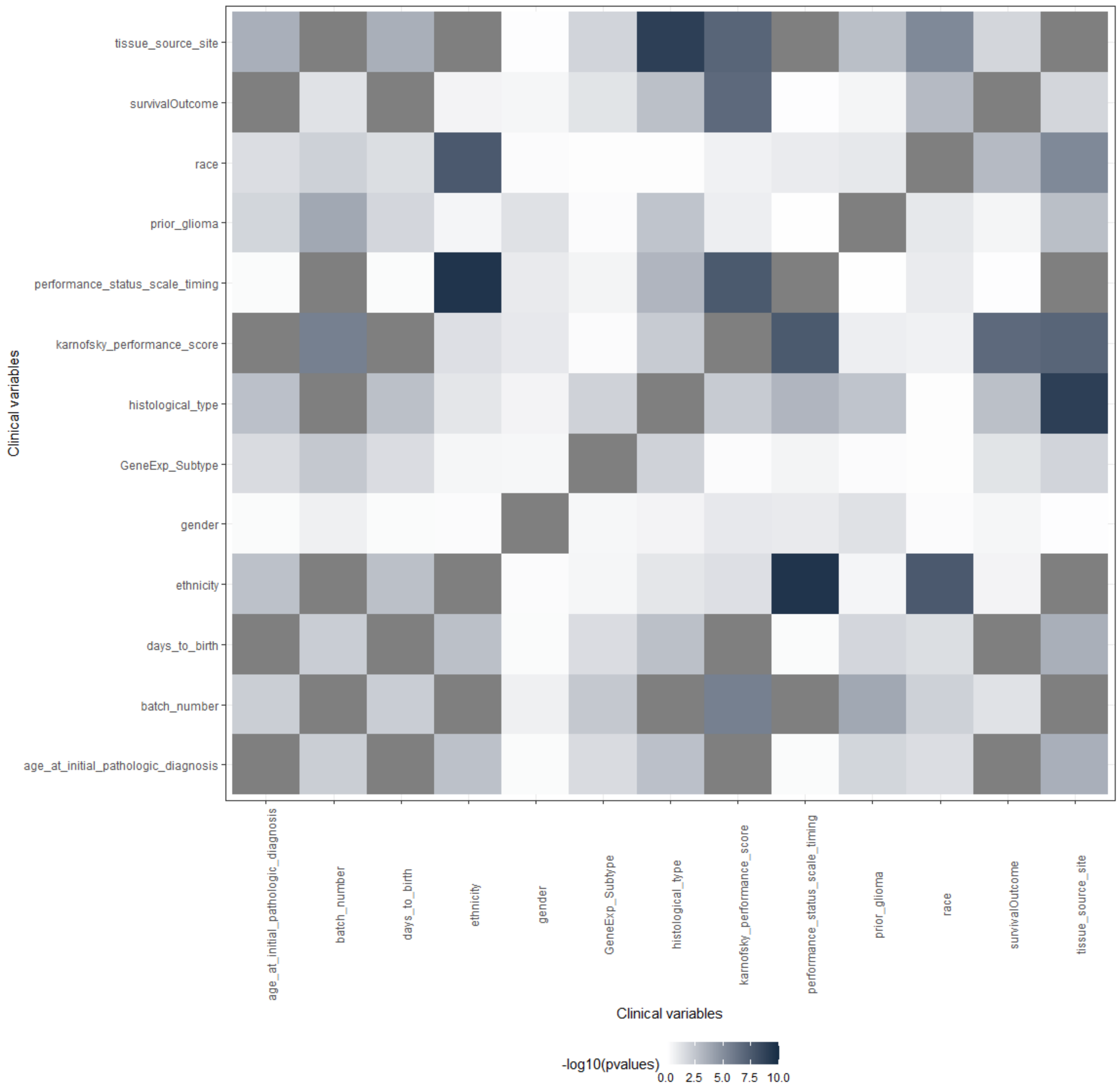

**Supplementary Figure 5.**

### Age in GBM

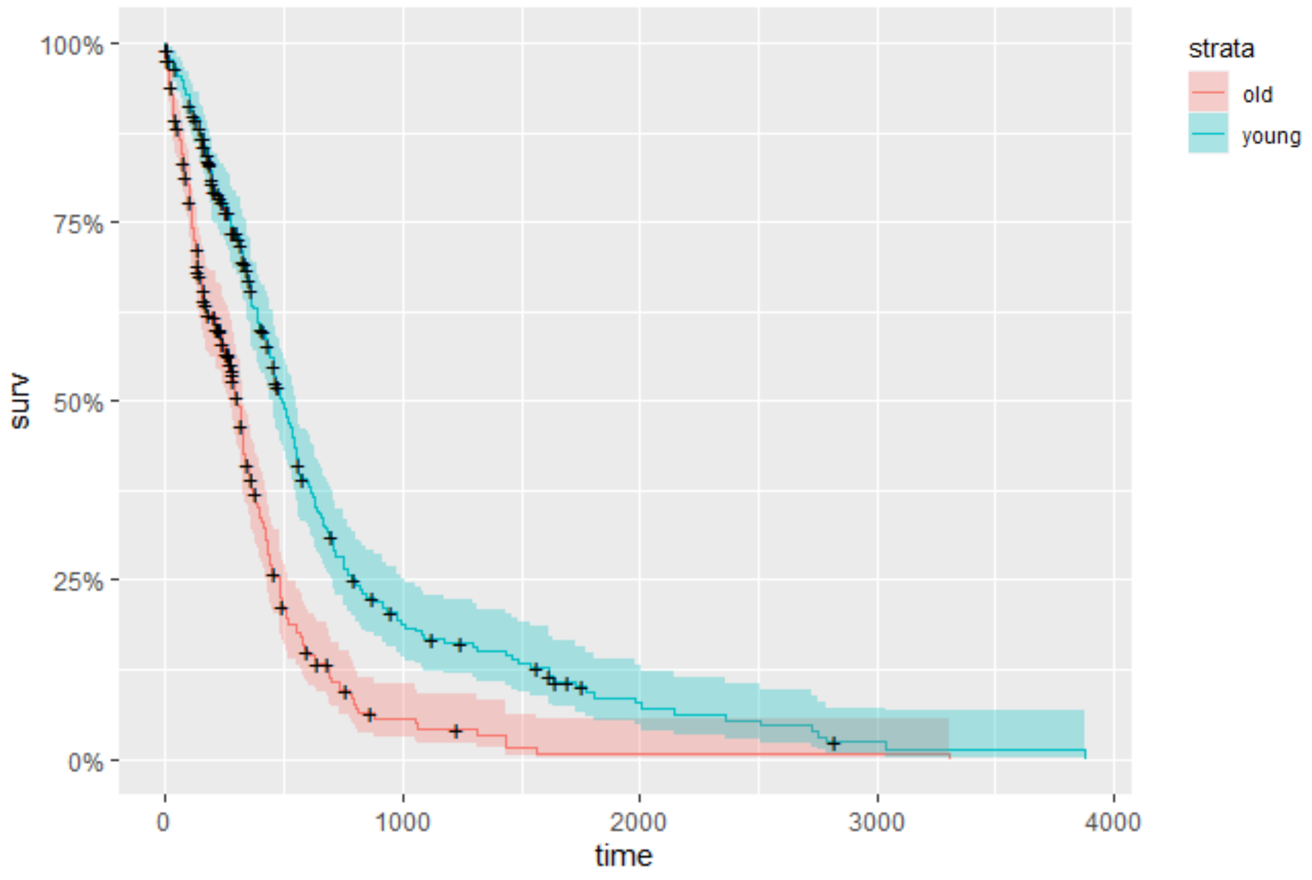
