## Supplementary material for "STATegra: Multi-omics data integration - A conceptual scheme and a bioinformatics pipeline": RMarkDown and html files.: stategra_gbm_pipeline.html

Omics integraive pipeline: STATegRa


### Omics integraive pipeline: STATegRa

Translational Bioinformatics Unit (Navarrabiomed)

#### 2020-11-02

### Contents

- 1 Introduction - Glioblastoma Multiforme
- 2 Initial data exploration and pre-process
  - 2.1 Metadata exploration
  - 2.2 Individual omics exploration
    - 2.2.1 Expression data (`rna_gbm`)
    - 2.2.2 miRNA data (`mirna_gbm`)
    - 2.2.3 Methylation data (`met_gbm`)
  - 2.3 Joint omics exploration
    - 2.3.1 Common samples
    - 2.3.2 OmicsPCA exploration
      - 2.3.2.1 Data scaling
      - 2.3.2.2 Model selection
        - 2.3.2.2.1 JIVE
        - 2.3.2.2.2 PCA-GCA
        - 2.3.2.2.3 pESCA
      - 2.3.2.3 Sub-space recovery
  - 2.4 Batch correction
    - 2.4.1 Expression data (`rna_gbm_corrected`)
    - 2.4.2 miRNA data (`mirna_gbm_corrected`)
    - 2.4.3 Methylation data (`logit_met_gbm`)
  - 2.5 Characterization of the data: Individual exploration
  - 2.6 Characterization of the data: Joint exploration
    - 2.6.1 Common samples
    - 2.6.2 OmicsPCA exploration
      - 2.6.2.1 Data scaling
      - 2.6.2.2 Model selection
        - 2.6.2.2.1 JIVE
        - 2.6.2.2.2 PCA-GCA
        - 2.6.2.2.3 pESCA
      - 2.6.2.3 Sub-space recovery
  - 2.7 Integrative differential analysis by NPC
    - 2.7.1 mRNA + miRNA
      - 2.7.1.1 Mapping file
      - 2.7.1.2 Overlaping samples
        - 2.7.1.2.1 Parametric Combination
        - 2.7.1.2.2 Non Parametric Combination
      - 2.7.1.3 All samples
        - 2.7.1.3.1 Parametric Combination
        - 2.7.1.3.2 Non Parametric Combination
      - 2.7.1.4 Integration strategies comparison
    - 2.7.2 mRNA + Methylation
      - 2.7.2.1 Mapping file
      - 2.7.2.2 Overlaping samples
        - 2.7.2.2.1 Parametric Combination
        - 2.7.2.2.2 Non Parametric Combination
      - 2.7.2.3 All samples
        - 2.7.2.3.1 Parametric Combination
        - 2.7.2.3.2 Non Parametric Combination
      - 2.7.2.4 Integration strategies comparison

### 1 Introduction - Glioblastoma Multiforme

Following the example above, we aim to provide a basic integrative workflow of omics data by combining existing tools implemented in the `STATegRa` R-package.

To this aim, the Glioblastoma Multiforme (GBM) dataset from The Cancer Genome Atlas (TCGA) is explored.

This dataset consists of primary tumor brain samples from roughly 600 patients including different omics data and metadata.

A description of the variables collected as metadata within the TCGA context is found in the NCBI’s GDC Documentation - Data Dictionary.

The level 3 public data for gene expression (mRNA), miRNA and methylation were obtained through the NCI’s Genomic Data Commons (GDC) portal (Tomczak, Czerwinska, & Wiznerowicz, 2015) and stored in an R-enviroment.

To start the analysis we are going to upload the data in our R session.

```
#Load the data to R session
load('20180613_1_gbm_raw_data.RData', verbose=TRUE)
```

```
## Loading objects:
##   rna_gbm
##   met_gbm
##   mirna_gbm
##   clinical_gbm
```

After upload the public data (omics + metadata): mRNA (`rna_gbm`), miRNA (`mirna_gbm`), methylation (`met_gbm`) and metadata (`clinical_gbm`; we can explore and check the dimension of these data-types.

```
dim(rna_gbm)
```

```
## [1] 12042   548
```

```
dim(met_gbm)
```

```
## [1] 485577    154
```

```
dim(mirna_gbm)
```

```
## [1] 534 571
```

```
dim(clinical_gbm)
```

```
## [1] 608  52
```

We can observed that the dimension of the omics differs, pointing that not all the omics are available for all cases, specially in the case of methylation data.

### 2 Initial data exploration and pre-process

Before start the integration analysis approach, an exploration of each omic data and metadata is required / recommended.

#### 2.1 Metadata exploration

Metadata provided by TCGA contains information about demographic features (age, gender, race, ethnicity), tumour characteristics (age at diagnosis, prior glioma, karnofsky score, gene expression subtype) and technical processes (batch, TSS, project, disease).

Exploring metadata we realized that there are 67 cases with unclassified gene expression subtype and we decide to remove all these cases from the analysis.

```
summary(clinical_gbm$GeneExp_Subtype)
```

```
##    Classical  Mesenchymal       Neural    Proneural Unclassified 
##          149          164           89          139           67
```

```
#Remove unclassified samples
samples_to_remove <- which(clinical_gbm$GeneExp_Subtype=='Unclassified') 
clinical_gbm <- clinical_gbm[-samples_to_remove,] 

# Drop unused levels of GeneExp_Subtype
clinical_gbm$GeneExp_Subtype <- droplevels(clinical_gbm$GeneExp_Subtype,
                                           "Unclassified")
```

Based on information provided we calculate the survival curve and define the survival variable `survivalOutcome` and check the age effect on survival.

```
# Load require packages
library(survival)
library(ggplot2)
library(ggfortify)

clinical_gbm$survivalTime <- pmin(clinical_gbm$days_to_death,
                                  clinical_gbm$days_to_last_followup, 
                                  na.rm=TRUE)

clinical_gbm$survivalOutcome <- Surv(time=clinical_gbm$survivalTime,
                                     event=clinical_gbm$vital_status=='Dead')

# Plot survival curve
my.fit <- survfit(clinical_gbm$survivalOutcome ~ 1)
autoplot(my.fit, surv.colour='orange', censor.colour='red',
         main='Kaplan-Meier estimate with 95% confidence bounds\n
         Glioblastoma',
         xlab='Time (days)')
```

```
## Check by age (in days from birth)
summary(clinical_gbm$days_to_birth)
```

```
##    Min. 1st Qu.  Median    Mean 3rd Qu.    Max. 
##  -32612  -25085  -21698  -21264  -18219   -3982
```

```
# Show in years
summary(-1*clinical_gbm$days_to_birth/365)
```

```
##    Min. 1st Qu.  Median    Mean 3rd Qu.    Max. 
##   10.91   49.92   59.45   58.26   68.73   89.35
```

```
# Set a cut-off arround 60 years
clinical_gbm$grouping_age <- rep("young",nrow(clinical_gbm))
clinical_gbm$grouping_age[clinical_gbm$days_to_birth < -21698] <- "old"
my.fit <- survfit(clinical_gbm$survivalOutcome ~ as.factor(clinical_gbm$grouping_age))
autoplot(my.fit)
```

```
survdiff(clinical_gbm$survivalOutcome ~ as.factor(clinical_gbm$grouping_age))
```

```
## Call:
## survdiff(formula = clinical_gbm$survivalOutcome ~ as.factor(clinical_gbm$grouping_age))
## 
##                                              N Observed Expected (O-E)^2/E
## as.factor(clinical_gbm$grouping_age)=old   270      216      149      30.3
## as.factor(clinical_gbm$grouping_age)=young 271      209      276      16.3
##                                            (O-E)^2/V
## as.factor(clinical_gbm$grouping_age)=old        49.2
## as.factor(clinical_gbm$grouping_age)=young      49.2
## 
##  Chisq= 49.2  on 1 degrees of freedom, p= 2e-12
```

We can clearly see a strong effect of age on survival curve. Young individuals (<60 years old) show higher survival than old individuals (>=60 years old).

Then, we define a function to explore the association between the clinical variables called `clinCorrelation`.

```
clin_var <- c('GeneExp_Subtype',
              'gender',
              'batch_number',
              'race',
              'ethnicity',
              'histological_type',
              'survivalOutcome',
              'age_at_initial_pathologic_diagnosis',
              'days_to_birth',
              'prior_glioma',
              'tissue_source_site',
              'karnofsky_performance_score',
              'performance_status_scale_timing')

clinCorrelation(clinical=clinical_gbm, clin_var=clin_var, maxlogPvalue = 10)
```

The resulting plot shows the -log10(pvalue) for each association test. High association is observed between:

- `batch_number`, `tissue_source_site` and `histological_type`
- `age_at_initial_pathologic_diagnosis`, `survivalOutcome` and `days_to_birth` (as the majority of cases included in the study are newly diagnosed) and `karnofsky_performance_score`.

#### 2.2 Individual omics exploration

Now, previous to any omics integrative approach, we should know which type of data do we have. Depending on the initial data characteristics we should apply some normalization, batch correction or transformation process.

##### 2.2.1 Expression data (`rna_gbm`)

Expression data from GBM is generated under microarray technology (Affymetrix Human Genome HT U133A), so we will have data from fluorescence intensity signal. As we are working with level 3 data, the data has been already normalized under the RMA approach (Cancer Genome Atlas Research Network.Nature 455.7216 (2008): 1061).

Let’s explore the distribution of the data.

```
summary(rna_gbm[,1:3])
```

```
##  TCGA-02-0432-01  TCGA-08-0245-01  TCGA-06-0137-01_rep2
##  Min.   : 3.271   Min.   : 3.290   Min.   : 3.299      
##  1st Qu.: 4.471   1st Qu.: 4.462   1st Qu.: 4.617      
##  Median : 5.753   Median : 5.832   Median : 5.377      
##  Mean   : 6.174   Mean   : 6.205   Mean   : 5.808      
##  3rd Qu.: 7.489   3rd Qu.: 7.541   3rd Qu.: 6.528      
##  Max.   :13.560   Max.   :13.713   Max.   :13.118
```

```
# Prepare data for distribution plot using ggplot
sampleNames = vector()
intensities = vector()
for (i in 1:ncol(rna_gbm)){
  sampleNames = c(sampleNames,rep(colnames(rna_gbm)[i],nrow(rna_gbm)))
  intensities = c(intensities,rna_gbm[,i])
}

arrayData <- data.frame(intensities,sampleNames)

g = ggplot(arrayData, aes(intensities, color=sampleNames))

g + geom_density() + 
  theme(legend.position='none') +
  labs(title='Gene Expression Density Curve',
       x='Microarray intensity Signal',
       y = 'Density')
```

Density plot shows a bimodal distribution, not usually expected in microarray data, but data is already normalized, so we will work with data provided without any further modification.

Checking the samples, we identify 10 cases with several replicated samples.

```
grep('rep',colnames(rna_gbm), value=TRUE)
```

```
##  [1] "TCGA-06-0137-01_rep2" "TCGA-06-0138-01_rep2" "TCGA-06-0156-01_rep3"
##  [4] "TCGA-06-0168-01_rep2" "TCGA-06-0148-01_rep1" "TCGA-06-0145-01_rep5"
##  [7] "TCGA-06-0138-01_rep1" "TCGA-06-0156-01_rep1" "TCGA-06-0148-01_rep3"
## [10] "TCGA-06-0137-01_rep3" "TCGA-06-0137-01_rep1" "TCGA-06-0137-01_rep4"
## [13] "TCGA-06-0145-01_rep3" "TCGA-06-0148-01_rep2" "TCGA-06-0208-01_rep1"
## [16] "TCGA-06-0145-01_rep6" "TCGA-06-0145-01_rep1" "TCGA-06-0211-01_rep1"
## [19] "TCGA-06-0148-01_rep4" "TCGA-06-0145-01_rep4" "TCGA-06-0208-01_rep2"
## [22] "TCGA-06-0211-01_rep2" "TCGA-06-0154-01_rep1" "TCGA-06-0156-01_rep2"
## [25] "TCGA-06-0176-01_rep1" "TCGA-06-0168-01_rep1" "TCGA-06-0154-01_rep2"
## [28] "TCGA-06-0176-01_rep2" "TCGA-06-0145-01_rep2"
```

We decide to keep the first replicate of each one of these cases and discard the rest.

```
#Keep just first replicate
colnames(rna_gbm) <- gsub('_rep1', '', colnames(rna_gbm))
rna_gbm <- rna_gbm[,colnames(rna_gbm)%in%rownames(clinical_gbm)]
```

Finally, the input data for gene expression of GBM consist in a normalized matrix with 12,042 features per 525 samples.

Let’s explore a little bit more this omic-type alone, using an unsupervised dimension reduction methodology such as Principal Component analysis (PCA), in order to see if there’s some batch-effect.

```
# Compute PCA
pca_rna <- prcomp(t(rna_gbm))
var_prcomp <- pca_rna$sdev^2

# Plot PCs variance
pcvar <- data.frame(var=var_prcomp/sum(var_prcomp), 
                    pc=c(1:length(var_prcomp)))

ggplot(pcvar[1:15,], aes(x = pc)) + geom_line(aes(y=var)) + 
  labs(x='Principal Component', y='Explained Variance') + 
  geom_point(aes(y=var)) + 
  ggtitle('RNA: Principal Components variance')
```

```
# Plot PCs scores
df <- data.frame(PC1=pca_rna$x[,1], PC2=pca_rna$x[,2], 
                 clinical_gbm[colnames(rna_gbm),
                              c('GeneExp_Subtype','batch_number')])

ggplot(df) +
  geom_point(aes(x=PC1,y=PC2,color=factor(GeneExp_Subtype)),
             size=5,
             shape=20)+
  guides(color=guide_legend('GeneExp_Subtype'),
         fill=guide_legend('GeneExp_Subtype')) +
  labs(x='PC1 (16.6%)', y='PC2 (10.2%)') +
  ggtitle('Principal Components for RNA data by Gene Expression Subtype')+
  theme(legend.position="bottom")
```

```
ggplot(df) +
  geom_point(aes(x=PC1, y=PC2, color=factor(batch_number)),
             size=5, 
             shape=20)+
  guides(color=guide_legend('batch_number'),
         fill=guide_legend('batch_number'))+
  labs(x='PC1 (16.6%)', y='PC2 (10.2%)') +
  ggtitle('Principal Components for RNA data by Batch')+
  theme(legend.position="bottom")
```

The first component explains only the 16.6% of variability, that in combination with second component seems that relates with gene expression subtypes, however, looking at batch level, it seems that there’s some batch effect in our data.

A function to explore the association between the clinical variables and principal components called `pcaCorrelation` is defined.

```
pcaCorrelation(clinical=clinical_gbm[colnames(rna_gbm),],
               clin_var=clin_var, pca_data=pca_rna$x, maxlogPvalue = 10)
```

A high association of `batch_number` and `tissue_source_site` with first principal component is found. Moreover, almost the rest of the components explored show a significant association with `batch_number`.
In the other side, `GenExp_Subtype` shows high association with the first six principal components.

These results highlight a batch effect that must be corrected.

##### 2.2.2 miRNA data (`mirna_gbm`)

miRNA data from GBM is generated under microarray technology (Agilent H-miRNA 8x15K), so we will have data from fluorescence intensity signal. As we are working with level 3 data, data is already normalized under quantil approach, log2-transformed and a Distance Weighted Discrimination (DWD) method has been applied for batch correction (Cancer Genome Atlas Research Network. Nature 455.7216 (2008): 1061.).

Let’s explore the distribution of the data.

```
summary(mirna_gbm[,1:4])
```

```
##  TCGA-02-0432-01  TCGA-08-0245-01  TCGA-28-1750-01  TCGA-06-1084-01 
##  Min.   : 5.676   Min.   : 5.654   Min.   : 5.614   Min.   : 5.503  
##  1st Qu.: 5.813   1st Qu.: 5.793   1st Qu.: 5.785   1st Qu.: 5.810  
##  Median : 6.128   Median : 6.056   Median : 6.077   Median : 6.112  
##  Mean   : 6.962   Mean   : 6.944   Mean   : 6.942   Mean   : 6.968  
##  3rd Qu.: 7.432   3rd Qu.: 7.464   3rd Qu.: 7.504   3rd Qu.: 7.584  
##  Max.   :14.998   Max.   :14.721   Max.   :14.992   Max.   :14.987
```

```
# Prepare data for ggplot
sampleNames = vector()
intensities = vector()
for (i in 1:ncol(mirna_gbm)){
  sampleNames = c(sampleNames,rep(colnames(mirna_gbm)[i],nrow(mirna_gbm)))
  intensities = c(intensities,mirna_gbm[,i])
}

arrayData <- data.frame(intensities,sampleNames)

g = ggplot(arrayData, aes(intensities, color=sampleNames))
g + geom_density() + theme(legend.position='none') +
  labs(title='miRNA Density Curve', 
       x='Microarray intensity Signal', 
       y='Density')
```

Density plot shows a normal shape for miRNA microarray data.

Checking the samples, we identify 5 cases with replicated samples.

```
grep("rep",colnames(mirna_gbm), value=TRUE)
```

```
##  [1] "TCGA-28-1751-01_rep1" "TCGA-26-1799-01_rep2" "TCGA-19-1788-01_rep1"
##  [4] "TCGA-28-1751-01_rep2" "TCGA-27-1833-01_rep1" "TCGA-14-1825-01_rep1"
##  [7] "TCGA-14-1825-01_rep2" "TCGA-19-1788-01_rep2" "TCGA-27-1833-01_rep2"
## [10] "TCGA-26-1799-01_rep1"
```

We decide to keep the first replicate of each one of these cases and discard the rest.

```
#Keep just first replicate
colnames(mirna_gbm) <- gsub('_rep1', '', colnames(mirna_gbm))
mirna_gbm <- mirna_gbm[,colnames(mirna_gbm)%in%rownames(clinical_gbm)]
```

Finally, the input data for miRNA expression of GBM consist in a normalized matrix with 534 features per 520 samples.

Let’s explore a little bit more this omic-type alone, using an unsupervised dimension reduction methodology such as Principal Component analysis (PCA), in order to see if there’s some batch-effect.

```
# Compute PCA
pca_mirna <- prcomp(t(mirna_gbm))
var_prcomp <- pca_mirna$sdev^2

# Plot PCs variance
pcvar <- data.frame(var=var_prcomp/sum(var_prcomp), 
                    pc=c(1:length(var_prcomp)))

ggplot(pcvar[1:15,], aes(x = pc)) + geom_line(aes(y=var)) + 
  labs(x='Principal Component', y='Explained Variance') + 
  geom_point(aes(y=var)) + 
  ggtitle('miRNA: Principal Components variance')
```

```
# Plot PCs scores
df <- data.frame(PC1=pca_mirna$x[,1], PC2=pca_mirna$x[,2],
                 clinical_gbm[colnames(mirna_gbm),
                              c('GeneExp_Subtype','batch_number')])

ggplot(df) +
  geom_point(aes(x=PC1, y=PC2, color=factor(GeneExp_Subtype)),
             size=5, 
             shape=20) +
  guides(color=guide_legend('GeneExp_Subtype'),
         fill=guide_legend('GeneExp_Subtype')) +
  labs(x='PC1 (19.4%)', y='PC2 (9.4%)') +
  ggtitle('Principal Components for miRNA data by Gene Expression Subtype')+
  theme(legend.position="bottom")
```

```
ggplot(df) +
  geom_point(aes(x=PC1, y=PC2, color=factor(batch_number)),
             size=5, 
             shape=20) +
  guides(color=guide_legend('batch_number'),
         fill=guide_legend('batch_number')) +
  labs(x='PC1 (19.4%)', y='PC2 (9.4%)') +
  ggtitle('Principal Components for miRNA data by Batch')+
  theme(legend.position="bottom")
```

The first component explains the 19.4% of variability, that in combination with second component seems that relates with gene expression subtypes, no effect of batch is observed in this data type.

The association between the clinical variables and principal components was explored using the `pcaCorrelation` function.

```
pcaCorrelation(clinical=clinical_gbm[colnames(mirna_gbm),],
               clin_var=clin_var, pca_data=pca_mirna$x, maxlogPvalue = 10)
```

In this case, no batch effect is detected. Instead, `GenExp_Subtype` shows significant association with the first six components.

##### 2.2.3 Methylation data (`met_gbm`)

Methylation data from GBM is generated under microarray technology (Illumina Infinium Human Methylation 450K). As we are working with level 3 data, the data has been already processes and the available data corresponds to beta values.

```
summary(met_gbm[,1:4])
```

```
##  TCGA-76-6661-01 TCGA-74-6577-01 TCGA-14-1034-02 TCGA-76-6282-01
##  Min.   :0.01    Min.   :0.01    Min.   :0.01    Min.   :0.01   
##  1st Qu.:0.07    1st Qu.:0.05    1st Qu.:0.06    1st Qu.:0.04   
##  Median :0.47    Median :0.41    Median :0.60    Median :0.46   
##  Mean   :0.47    Mean   :0.44    Mean   :0.51    Mean   :0.45   
##  3rd Qu.:0.87    3rd Qu.:0.81    3rd Qu.:0.92    3rd Qu.:0.85   
##  Max.   :1.00    Max.   :0.99    Max.   :0.99    Max.   :0.99   
##  NA's   :89836   NA's   :89583   NA's   :91295   NA's   :89572
```

As we can see in the summary of the first 4 samples there are a large number of missings

```
# Calculate the number of NAs by row (feature)
number_NAs <- rowSums(is.na(met_gbm))
# Features with missing values
sum(number_NAs!=0) #103,125 features with at least one NA
```

```
## [1] 103125
```

```
# Features with missings in all cases
sum(number_NAs==ncol(met_gbm)) #89,512 features with all NAs
```

```
## [1] 89512
```

There are 89,512 features with missing values in all cases, moreover there are 13,613 features with missings in at least one case.
We decide to remove all features that have at least one missing value, resulting in a dataset with 382,452. features.

```
met_gbm <- met_gbm[number_NAs==0,] 
# New data dimension:
dim(met_gbm)
```

```
## [1] 382452    154
```

Moreover, additional filters are applied based on probes annotation. Gender related probes (chr Y and X) and probes with SNPs, CpG or SBE in target sequence are removed.

```
# Load require packages
library(IlluminaHumanMethylation450kanno.ilmn12.hg19)
```

```
# Get annotation data from methylation
ann450k <- getAnnotation(IlluminaHumanMethylation450kanno.ilmn12.hg19)
ann450k[1:4,1:5]
```

```
## DataFrame with 4 rows and 5 columns
##                    chr       pos      strand        Name    AddressA
##            <character> <integer> <character> <character> <character>
## cg00050873        chrY   9363356           -  cg00050873    32735311
## cg00212031        chrY  21239348           -  cg00212031    29674443
## cg00213748        chrY   8148233           -  cg00213748    30703409
## cg00214611        chrY  15815688           -  cg00214611    69792329
```

```
# Gender related probes
xy_linked <- which(ann450k$chr=='chrY' | ann450k$chr=='chrX') 
#11648 features

# SNPs, CpG and SBE in target sequence
target_snp <- which(!is.na(ann450k$Probe_rs)) #87018 features
target_cpg <- which(!is.na(ann450k$CpG_rs)) #16998 features
target_sbe <- which(!is.na(ann450k$SBE_rs)) #7876 features

# Filter dataset
rm_probes <- unique(c(xy_linked,target_snp, target_sbe, target_cpg)) 
#114614 features

met_gbm <- met_gbm[rownames(met_gbm)%in%rownames(ann450k)[-rm_probes],] 
#New dimension: 311135 x 154
```

Also, there are 59 samples in methylation dataset without information in metadata.

```
setdiff(colnames(met_gbm),rownames(clinical_gbm))
```

```
##  [1] "TCGA-76-6661-01" "TCGA-74-6577-01" "TCGA-06-6388-01" "TCGA-06-6693-01"
##  [5] "TCGA-06-A7TL-01" "TCGA-RR-A6KC-01" "TCGA-19-4065-02" "TCGA-14-0862-01"
##  [9] "TCGA-26-A7UX-01" "TCGA-19-A6J5-01" "TCGA-28-2501-01" "TCGA-74-6578-01"
## [13] "TCGA-06-6698-01" "TCGA-19-A60I-01" "TCGA-06-1806-01" "TCGA-06-A7TK-01"
## [17] "TCGA-06-6700-01" "TCGA-06-6695-01" "TCGA-28-5211-01" "TCGA-41-6646-01"
## [21] "TCGA-OX-A56R-01" "TCGA-06-6694-01" "TCGA-14-1450-01" "TCGA-76-6286-01"
## [25] "TCGA-74-6573-01" "TCGA-06-6699-01" "TCGA-74-6581-01" "TCGA-19-4065-01"
## [29] "TCGA-76-6660-01" "TCGA-06-6701-01" "TCGA-19-5953-01" "TCGA-06-A5U1-01"
## [33] "TCGA-81-5911-01" "TCGA-06-A6S0-01" "TCGA-26-6174-01" "TCGA-RR-A6KA-01"
## [37] "TCGA-4W-AA9S-01" "TCGA-76-6663-01" "TCGA-74-6573-11" "TCGA-76-6283-01"
## [41] "TCGA-06-A6S1-01" "TCGA-14-1043-01" "TCGA-19-A6J4-01" "TCGA-06-A5U0-01"
## [45] "TCGA-76-6662-01" "TCGA-76-6657-01" "TCGA-26-6173-01" "TCGA-74-6575-01"
## [49] "TCGA-76-6656-01" "TCGA-74-6584-01" "TCGA-RR-A6KB-01" "TCGA-4W-AA9R-01"
## [53] "TCGA-06-6697-01" "TCGA-76-6280-01" "TCGA-28-2510-01" "TCGA-76-6664-01"
## [57] "TCGA-14-0740-01" "TCGA-14-1395-01" "TCGA-4W-AA9T-01"
```

These cases are removed from the study.

```
met_gbm <- met_gbm[,colnames(met_gbm)%in%rownames(clinical_gbm)] 
#New datset dimension:
dim(met_gbm)
```

```
## [1] 311135     95
```

Finally, we transform the beta values to M values.

```
# Load require packages
library(lumi)

# Transform to M values
logit_met_gbm <- beta2m(met_gbm) 

summary(logit_met_gbm[,1:3])
```

```
##  TCGA-14-1034-02    TCGA-76-6282-01   TCGA-19-0957-01  
##  Min.   :-7.39091   Min.   :-7.5284   Min.   :-6.8585  
##  1st Qu.:-3.93945   1st Qu.:-4.4331   1st Qu.:-3.9460  
##  Median : 0.59805   Median :-0.2235   Median :-0.4586  
##  Mean   :-0.05774   Mean   :-0.7481   Mean   :-0.7780  
##  3rd Qu.: 3.56892   3rd Qu.: 2.5226   3rd Qu.: 1.9586  
##  Max.   : 7.41139   Max.   : 7.5606   Max.   : 7.1144
```

The resulting input GBM data for methylation consists in a M values matrix with 311,135 features per 95 samples.

Let’s explore a little bit more this omic-type alone, using an unsupervised dimension reduction methodology such as Principal Component analysis (PCA), in order to see if there’s some batch-effect.

```
# Compute PCA
pca_met <- prcomp(t(logit_met_gbm))
var_prcomp <- pca_met$sdev^2

# Plot PCs variance
pcvar <- data.frame(var=var_prcomp/sum(var_prcomp), 
                    pc=c(1:length(var_prcomp)))

ggplot(pcvar[1:15,], aes(x = pc)) + geom_line(aes(y=var)) + 
  labs(x='Principal Component', y='Explained Variance') + 
  geom_point(aes(y=var)) + 
  ggtitle('Methylation: Principal Components variance')
```

```
# Plot PCs scores
df <- data.frame(PC1=pca_met$x[,1], PC2=pca_met$x[,2], 
                 clinical_gbm[colnames(logit_met_gbm),
                              c('GeneExp_Subtype','batch_number')])

ggplot(df) +
  geom_point(aes(x=PC1, y=PC2, color=factor(GeneExp_Subtype)),
             size=5,
             shape=20) +
  guides(color=guide_legend('GeneExp_Subtype'),
         fill=guide_legend('GeneExp_Subtype')) +
  labs(x='PC1 (14.6%)', y='PC2 (10.8%)') +
  ggtitle('Principal Components for Methylation data by 
          Gene Expression Subtype') +
  theme(legend.position="bottom")
```

```
ggplot(df) +
  geom_point(aes(x=PC1, y=PC2, color=factor(batch_number)), 
             size=5, 
             shape=20) +
  guides(color=guide_legend('batch_number'),
         fill=guide_legend('batch_number')) +
  labs(x='PC1 (14.6%)', y='PC2 (10.8%)') +
  ggtitle('Principal Components for Methylation data by Batch') +
  theme(legend.position="bottom")
```

No cluster of samples according gene expression subtype or batch is apreciated.

Let’s explore the association between clinical variables and principal components by the `pcaCorrelation` function.

```
pcaCorrelation(clinical=clinical_gbm[colnames(logit_met_gbm),], 
               clin_var=clin_var, pca_data=pca_met$x, maxlogPvalue = 10)
```

`GenExp_Subtype` shows significant association with the first three components. Also, high association with `survivalOutcome` is found with second component. Worth noting, several components, mainly PC10 and PC11 show significant association with `batch_number`, and association of `days_to_birth` and `age_at_initial_pathologic_diagnosis` with PC3, PC6 and PC10 is identified.

These results highlight a batch effect in methylation dataset that must be corrected.

#### 2.3 Joint omics exploration

In order to identify if there are share components between the omics or say in other words, see how much the different data-types are “explaining the samples in the same way”, we will perform what we define as join- or omics- PCA.

##### 2.3.1 Common samples

As we have different number of samples in each omic-type, we need to explore the shared samples between omics.

```
# Load require packages
library(VennDiagram)
library(plotrix)
```

```
# Venn Diagram
m <- list(rna=colnames(rna_gbm), 
          mirna=colnames(mirna_gbm), 
          met=colnames(logit_met_gbm))

vennPlot <- venn.diagram(m, NULL, fill=c('red','green','blue'), 
                         cat.fontface=4, lty=2, cex=1.2, cat.cex=1.2)
grid.newpage()
grid.draw(vennPlot)
```

We have 517 samples (almost 100%) with both mRNA and miRNA data, however, just 83 samples have mRNA, miRNA and methylation data.

##### 2.3.2 OmicsPCA exploration

The exploration of each omic-type independently have been shown a batch effect that must be corrected at least in mRNA and methylation data-types.

Let’s explore what happens if we perform a joint exploration using multi-omics PCA appraoch.

```
# Load require packages
library(STATegRa)
library(MASS)
library(gridExtra)
```

###### 2.3.2.1 Data scaling

Before start the multi-omic approach, mean centred of each feature and the data set normalization to unit sum of squares (forbenious normalization) is recommended to make more comparabable the different omics data-sets.

```
#Calculate Forbenious normalization
frobenius_rna <- norm(as.matrix(rna_gbm), type="F")
frobenius_mirna <- norm(as.matrix(mirna_gbm), type="F")
frobenius_met <- norm(as.matrix(logit_met_gbm), type="F")

#Mean-centring and division by Frobenius normalization factor
rna_gbm_norm <- t(scale(t(rna_gbm), scale=FALSE))/frobenius_rna
mirna_gbm_norm <- t(scale(t(mirna_gbm), scale=FALSE))/frobenius_mirna
met_gbm_norm <- t(scale(t(logit_met_gbm), scale=FALSE))/frobenius_met

##Mean-centring without Frobenius normalization
rna_gbm_norm2 <- t(scale(t(rna_gbm), scale=FALSE))
mirna_gbm_norm2 <- t(scale(t(mirna_gbm), scale=FALSE))
met_gbm_norm2 <- t(scale(t(logit_met_gbm), scale=FALSE))
```

Then, we need to keep the common samples between the different data-types.

```
#Common samples between mRNA and miRNA
ids_rna_mirna <- intersect(colnames(rna_gbm), colnames(mirna_gbm)) 
length(ids_rna_mirna)
```

```
## [1] 517
```

```
#Common samples between mRNA and Methylation
ids_rna_met <- intersect(colnames(rna_gbm), colnames(logit_met_gbm))
length(ids_rna_met)
```

```
## [1] 83
```

###### 2.3.2.2 Model selection

Now, we are ready to explore if there are share components between data-types.

The first step is to determine the number of share/common and individual components between omics.

How to determine these numbers is still an open question nowadays, so the user can explore different methods like jive, pca-gca and pESCA, between others. Following, we show how to perform the model selection based on these three methods.

###### 2.3.2.2.1 JIVE

```
# Load require packages
library(r.jive)

#Using r.jive

####################################################
## mRNA + miRNA (already centered and normalized) ##
####################################################
Data <- list(mRNA=rna_gbm_norm[,ids_rna_mirna],
             miRNA=mirna_gbm_norm[,ids_rna_mirna])
rjive_rna_mirna <- jive(Data, scale = FALSE, center = FALSE) 


##########################################################
## mRNA + methylation (already centered and normalized) ##
##########################################################
Data <- list(mRNA=rna_gbm_norm[,ids_rna_met],
              met=met_gbm_norm[,ids_rna_met])
rjive_rna_met <- jive(Data, scale = FALSE, center = FALSE)
```

###### 2.3.2.2.2 PCA-GCA

```
# Load require packages
library(RegularizedSCA)

#Using pca-gca

####################################################
## mRNA + miRNA (already centered and normalized) ##
####################################################
data <- cbind(t(rna_gbm_norm[,ids_rna_mirna]), 
              t(mirna_gbm_norm[,ids_rna_mirna]))

# Number of variables
Jk <- c(nrow(rna_gbm_norm), nrow(mirna_gbm_norm))

# Select components
pca_gca(data, Jk, cor_min = .7)

##########################################################
## mRNA + methylation (already centered and normalized) ##
##########################################################
data <- cbind(t(rna_gbm_norm[,ids_rna_met]), 
              t(met_gbm_norm[,ids_rna_met]))

# Number of variables
Jk <- c(nrow(rna_gbm_norm), nrow(met_gbm_norm))

# Select components
pca_gca(data, Jk, cor_min = .7)
```

###### 2.3.2.2.3 pESCA

```
# Load require packages
library(RpESCA)
library(RSpectra)

#Using RpESCA

################################################
## mRNA + miRNA (already centered)            ##
################################################

dataSets <- list(rna=t(rna_gbm_norm2[,ids_rna_mirna]), 
                 mirna=t(mirna_gbm_norm2[,ids_rna_mirna]))

dataTypes <- 'GG'

# used parameters
opts <- list()
opts$tol_obj <- 1E-4
opts$quiet <- 1

# save the estimated alphas, selected number of PCs and cvErrors
alphas <- rep(NA,2)
R_selected_list <- as.list(1:2)
cvErrors_list <- as.list(1:2)

# alpha estimation for each data set
for (i in 1:length(dataSets)){
  # index ith data set
  X <- dataSets[[i]]
  
  # alpha estimation procedure
  alpha_est <- alpha_estimation(X,K=3,Rs = 5:20,opts=opts)
  alphas[i] <- alpha_est$alphas_mean
  R_selected_list[[i]] <- alpha_est$R_CV
  cvErrors_list[[i]] <- alpha_est$cvErrors
}
names(alphas) <- paste0('alpha', 1:2)

## Setting the parameters for the pESCA model

# concave function and its hyper-parameter
fun_concave <- 'gdp'; gamma <- 1;
penalty = 'L2' # concave L2 norm penalty

# Parameters of a pESCA with concave L2norm penalty model
opts <- list()
opts$gamma <- gamma  # hyper-parameter for the used penalty
opts$rand_start <- 0
opts$tol_obj <- 1e-6 # stopping criteria
opts$maxit   <- 500
opts$alphas  <- alphas
#opts$R # components used Default 0.5*min(I,J) - Minimum number of variables.
opts$thr_path <- 0 # generaint thresholding path or not
opts$quiet <- 1

# Model selection pESCA conave L2 norm penalty
nTries <- 15
lambdas_CV <- log10_seq(from=1, to=500, length.out=nTries)

result_CV <- pESCA_CV(dataSets, dataTypes,
                      lambdas_CV, 
                      penalty=penalty, 
                      fun_concave=fun_concave, 
                      opts=opts)


# select the model with minimum CV error
index_min_cv <- which.min(result_CV$cvErrors_mat[,1])

# cvErrors during the model selection process
result_CV$cvErrors_mat

# selected value of lambda
lambdas_CV[index_min_cv]

# fit the final model
lambdas_opt <- rep(lambdas_CV[index_min_cv],length(dataSets))
opts_opt <- result_CV$inits[[index_min_cv]]
opts_opt$tol_obj <- 1E-8 # using high precision model


pESCA_L2 <- pESCA(dataSets = dataSets,
                          dataTypes = dataTypes,
                          lambdas = lambdas_opt,
                          penalty = penalty,
                          fun_concave= fun_concave,
                          opts = opts_opt)


mu <- pESCA_L2$mu
A <- pESCA_L2$A
B <- pESCA_L2$B
S <- pESCA_L2$S

# estimated variation explained ratios for each data set or the full data set
pESCA_L2$varExpTotals

# estimated variation explained ratios of each PC for each data set or the full data set
pESCA_L2$varExpPCs

#Cut-off 1% 
sel1 <- which(pESCA_L2$varExpPCs[1,]>1) #14
sel2 <- which(pESCA_L2$varExpPCs[2,]>1) #23
intersect(sel1,sel2)
# 8 common components
# 6 dist mRNA + 15 miRNA
# Total: 29 components

#Cut-off 5%
sel1 <- which(pESCA_L2$varExpPCs[1,]>5) #3
sel2 <- which(pESCA_L2$varExpPCs[2,]>5) #2
intersect(sel1,sel2)
# 1 common components
# 2 dist mRNA + 1 miRNA
# Total: 4 components


################################################
## mRNA + Methylation (already centered)      ##
################################################

dataSets <- list(rna=t(rna_gbm_norm2[,ids_rna_met]), 
                 met=t(met_gbm_norm2[,ids_rna_met]))

dataTypes <- 'GG'

# used parameters
opts <- list()
opts$tol_obj <- 1E-4
opts$quiet <- 1

# save the estimated alphas, selected number of PCs and cvErrors
alphas <- rep(NA,2)
R_selected_list <- as.list(1:2)
cvErrors_list <- as.list(1:2)

# alpha estimation for each data set
for (i in 1:length(dataSets)){
  # index ith data set
  X <- dataSets[[i]]
  
  # alpha estimation procedure
  alpha_est <- alpha_estimation(X,K=3,Rs = 5:20,opts=opts)
  alphas[i] <- alpha_est$alphas_mean
  R_selected_list[[i]] <- alpha_est$R_CV
  cvErrors_list[[i]] <- alpha_est$cvErrors
}
names(alphas) <- paste0('alpha', 1:2)

## Setting the parameters for the pESCA model

# concave function and its hyper-parameter
fun_concave <- 'gdp'; gamma <- 1;
penalty = 'L2' # concave L2 norm penalty

# Parameters of a pESCA with concave L2norm penalty model
opts <- list()
opts$gamma <- gamma  # hyper-parameter for the used penalty
opts$rand_start <- 0
opts$tol_obj <- 1e-6 # stopping criteria
opts$maxit   <- 500
opts$alphas  <- alphas
#opts$R  # components used Default 0.5*min(I,J) - Minimum number of variables.
opts$thr_path <- 0 # generaint thresholding path or not
opts$quiet <- 1

# Model selection pESCA conave L2 norm penalty
nTries <- 15
lambdas_CV <- log10_seq(from=1, to=500, length.out=nTries)

result_CV <- pESCA_CV(dataSets, dataTypes,
                      lambdas_CV, 
                      penalty=penalty, 
                      fun_concave=fun_concave, 
                      opts=opts)


# select the model with minimum CV error
index_min_cv <- which.min(result_CV$cvErrors_mat[,1])

# cvErrors during the model selection process
result_CV$cvErrors_mat

# selected value of lambda
lambdas_CV[index_min_cv]

# fit the final model
lambdas_opt <- rep(lambdas_CV[index_min_cv],length(dataSets))
opts_opt <- result_CV$inits[[index_min_cv]]
opts_opt$tol_obj <- 1E-8 # using high precision model


pESCA_L2 <- pESCA(dataSets = dataSets,
                          dataTypes = dataTypes,
                          lambdas = lambdas_opt,
                          penalty = penalty,
                          fun_concave= fun_concave,
                          opts = opts_opt)

mu <- pESCA_L2$mu
A <- pESCA_L2$A
B <- pESCA_L2$B
S <- pESCA_L2$S

# estimated variation explained ratios for each data set or the full data set
pESCA_L2$varExpTotals

# estimated variation explained ratios of each PC for each data set or the full data set
pESCA_L2$varExpPCs

#Cut-off 1%
sel1 <- which(pESCA_L2$varExpPCs[1,]>1) #3
sel2 <- which(pESCA_L2$varExpPCs[2,]>1) #16
intersect(sel1,sel2)
# 2 common components
# 1 dist mRNA + 14 methylation
# Total: 17 components

#Cut-off 5%
sel1 <- which(pESCA_L2$varExpPCs[1,]>5) #3
sel2 <- which(pESCA_L2$varExpPCs[2,]>5) #3
intersect(sel1,sel2)
# 1 common components
# 2 dist mRNA + 2 methylation
# Total: 5 components
```

Here, there is the table summarizing the output of the different methods.

|  | Component | r.JIVE | PCA-GCA (1%) | PCA-GCA (5%) | pESCA(1%) | pESCA (5%) |
| --- | --- | --- | --- | --- | --- | --- |
| mRNA + miRNA |  |  |  |  |  |  |
|  | Common | 3 | 3 | NA | 8 | 1 |
|  | Dist. mRNA | 46 | 11 | NA | 6 | 2 |
|  | Dist. miRNA | 24 | 6 | NA | 15 | 1 |
| mRNA + met |  |  |  |  |  |  |
|  | Common | 0 | 3 | NA | 2 | 1 |
|  | Dist. mRNA | 12 | 0 | NA | 1 | 2 |
|  | Dist. miRNA | 14 | 79 | NA | 14 | 2 |

As we can observe, the results from the different methods diverge. We are not able to recommend a method but, in general, we can say that it seems that there’s a small part of data variation that is share between omics. To follow with the analysis, we will explore the omicsPCA based on the output from pESCA (1%).

###### 2.3.2.3 Sub-space recovery

Based on the number of components identified (pESCA) we can recover the subspace.

It is interesting to take a look on the common and distinctive components identified through this method.

```
################################################
## mRNA + miRNA (already centered)            ##
################################################

# Explore the common and distinctive components and those variation explained
# Common components
cc <- intersect(sel1,sel2)
pESCA_L2$varExpPCs[,cc]
```

```
##              PC2      PC3      PC4      PC6      PC7      PC8      PC9     PC13
## X_1    10.246012 8.896854 4.711449 3.272001 1.845306 1.449171 1.562691 1.105669
## X_2     7.834064 1.539726 3.200150 3.044305 3.105618 1.127440 1.152379 2.376831
## X_full 10.110927 8.484807 4.626807 3.259248 1.915891 1.431152 1.539711 1.176863
```

```
# Distinctive components
dist_rna <- sel1[!sel1%in%cc] #rna
pESCA_L2$varExpPCs[,dist_rna]
```

```
##               PC1       PC5     PC10     PC11      PC12      PC14
## X_1    16.1078901 4.3493992 1.494169 1.535083 1.1826284 1.0305920
## X_2     0.4011646 0.3775905 0.000000 0.000000 0.4774752 0.0000000
## X_full 15.2282114 4.1269521 1.410486 1.449108 1.1431353 0.9728721
```

```
dist_mirna <- sel2[!sel2%in%cc] #mirna
pESCA_L2$varExpPCs[,dist_mirna]
```

```
##             PC23      PC32       PC37      PC38      PC61      PC67       PC94
## X_1    0.5756060 0.4479649  0.0000000 0.3558212 0.2303761 0.2340943 0.00000000
## X_2    3.6881272 1.1809512 12.2061008 1.2900009 3.9495539 1.0380562 1.19710506
## X_full 0.7499274 0.4890169  0.6836209 0.4081413 0.4386742 0.2791213 0.06704566
##             PC99     PC112     PC113      PC133     PC137      PC138     PC142
## X_1    0.0000000 0.0000000 0.0000000 0.00000000 0.0000000 0.00000000 0.0000000
## X_2    1.8600263 3.5963533 2.1699062 1.17380760 1.8502620 1.17134159 2.3753909
## X_full 0.1041736 0.2014191 0.1215288 0.06574085 0.1036267 0.06560274 0.1330373
##             PC143
## X_1    0.00000000
## X_2    1.11051782
## X_full 0.06219621
```

```
# Keep de common and distinctive components
common_pc <- pESCA_L2$A[,cc]
colnames(common_pc) <- colnames(pESCA_L2$varExpPCs[,cc])

dist_rna_pc <- pESCA_L2$A[,dist_rna]
colnames(dist_rna_pc) <- names(dist_rna)

dist_mirna_pc <- pESCA_L2$A[,dist_mirna]
colnames(dist_mirna_pc) <- names(dist_mirna)


# metadata for common samples
clin <- clinical_gbm[ids_rna_mirna,]

# PCA plot
require(ggplot2)

df <- as.data.frame(common_pc)
df$group <- clin$GeneExp_Subtype
df$batch <- clin$batch_number

#plot of first common components by gene expression subtype
p <- ggplot(df,aes_string("PC2","PC3",color="group"))
p + geom_point(size=5, shape=20) + ggtitle("PCA by Gene Expression Subtype") + theme(legend.position="bottom")
```

```
#plot of first common components by batch
p <- ggplot(df,aes_string("PC2","PC3",color="batch"))
p + geom_point(size=5, shape=20) + ggtitle("PCA by batch") + theme(legend.position="bottom")
```

Let’s explore how relate these common and distinctive components with the clinical variables.

```
df <- data.frame(common_pc,dist_rna_pc,dist_mirna_pc)
colnames(df) <- c(paste("common_",colnames(common_pc),sep=""),
                  paste("dist_rna_",colnames(dist_rna_pc),sep=""),
                  paste("dist_mirna_",colnames(dist_mirna_pc),sep=""))

pcaCorrelation(clinical=clin, clin_var, df, maxlogPvalue =10, maxPC=ncol(df), pc.names=TRUE)
```

The main common variability is associated with `GeneExp_Subtype`, `survivalOutcome`,`batch_number` and `tissue_source_cite`. These analysis highlight a strong batch effect, identified in common components but also in the distinct variables of mRNA.

Now, let’s explore what happens with the joint exploration of RNA and DNA methylation.

```
################################################
## mRNA + methylation (already centered)      ##
################################################

# Explore the common and distinctive components and those variation explained
# Common components
cc <- intersect(sel1,sel2)
pESCA_L2$varExpPCs[,cc]
```

```
##              PC2       PC7
## X_1     5.473947 24.818060
## X_2    10.571043  1.603196
## X_full 10.296246  2.854765
```

```
# Distinctive components
dist_rna <- sel1[!sel1%in%cc] #rna
pESCA_L2$varExpPCs[,dist_rna]
```

```
##        X_1        X_2     X_full 
## 13.0520328  0.0000000  0.7036662
```

```
dist_met <- sel2[!sel2%in%cc] #met
pESCA_L2$varExpPCs[,dist_met]
```

```
##             PC1      PC3      PC4      PC5      PC6      PC8      PC9     PC10
## X_1     0.00000 0.000000 0.000000 0.000000 0.000000 0.000000 0.000000 0.000000
## X_2    14.47574 7.594365 3.588316 3.122772 2.770845 2.131806 1.783550 2.249335
## X_full 13.69532 7.184935 3.394861 2.954416 2.621462 2.016875 1.687395 2.128068
##            PC11     PC12     PC14     PC15      PC16     PC17
## X_1    0.000000 0.000000 0.000000 0.000000 0.0000000 0.000000
## X_2    1.767639 1.432827 1.124813 1.099665 1.0454081 1.264661
## X_full 1.672341 1.355580 1.064172 1.040379 0.9890477 1.196480
```

```
# Keep de common and distinctive components
common_pc <- pESCA_L2$A[,cc]
colnames(common_pc) <- colnames(pESCA_L2$varExpPCs[,cc])

dist_rna_pc <- data.frame(pESCA_L2$A[,dist_rna])
colnames(dist_rna_pc) <- names(dist_rna)

dist_met_pc <- pESCA_L2$A[,dist_met]
colnames(dist_met_pc) <- names(dist_met)


# metadata for common samples
clin <- clinical_gbm[ids_rna_met,]

# PCA plot
require(ggplot2)

df <- as.data.frame(common_pc)
df$group <- clin$GeneExp_Subtype
df$batch <- clin$batch_number

#plot of first common components by gene expression subtype
p <- ggplot(df,aes_string("PC2","PC7",color="group"))
p + geom_point(size=5, shape=20) + ggtitle("PCA by Gene Expression Subtype") + theme(legend.position="bottom")
```

```
#plot of first common components by batch
p <- ggplot(df,aes_string("PC2","PC7",color="batch"))
p + geom_point(size=5, shape=20) + ggtitle("PCA by batch") + theme(legend.position="bottom")
```

A clear batch effect is observed in the principal common components representation.

Let’s explore how relate these common and distinctive components with the clinical variables.

```
df <- data.frame(common_pc,dist_rna_pc,dist_met_pc)
colnames(df) <- c(paste("common_",colnames(common_pc),sep=""),
                  paste("dist_rna_",colnames(dist_rna_pc),sep=""),
                  paste("dist_met_",colnames(dist_met_pc),sep=""))

pcaCorrelation(clinical=clin, clin_var, df, maxlogPvalue =10, maxPC=ncol(df), pc.names=TRUE)
```

The main common variability is associated with `GeneExp_Subtype`, `survivalOutcome`,`batch_number` and `tissue_source_cite`. Again, a strong batch effect is identified in both data-types.

#### 2.4 Batch correction

In order to remove batch effect from different omics data we applied the ComBat method

```
# Load require package
library(sva)
```

Before to apply ComBat, it’s important to note that there are two batches with only one sample

```
summary(as.factor(as.character(clinical_gbm[,'batch_number'])))
```

```
##   1.83.0  10.71.0 111.51.0  16.74.0   2.83.0  20.71.0  26.41.0  26.62.0 
##       25       28       37       50       17       48        1       44 
##   3.82.0  38.61.0   4.87.0   5.77.0   6.81.0  62.42.0  62.58.0   7.81.0 
##       19       29       40       63       35        1       24       21 
##  79.53.0   8.80.0 
##       35       24
```

```
clinical_gbm[which(clinical_gbm$batch_number%in%c('26.41.0','62.42.0')),c(1,4)]
```

```
##                         name batch_number
## TCGA-27-1836-01 TCGA-27-1836      26.41.0
## TCGA-32-2495-01 TCGA-32-2495      62.42.0
```

These two samples must be removed to apply the batch correction. They are present in mRNA and miRNA data-types, so we remove it from both.

```
rm_samples_batch <- c('TCGA-27-1836-01', 'TCGA-32-2495-01')

rna_gbm_corrected <- rna_gbm[,!colnames(rna_gbm)%in%rm_samples_batch] 
#New dimension: 12042 x 523

mirna_gbm_corrected <- mirna_gbm[,!colnames(mirna_gbm)%in%rm_samples_batch] 
#New dimension: 534 x 518
```

Now, we are ready to apply the batch correction for each omic data-type.

##### 2.4.1 Expression data (`rna_gbm_corrected`)

```
mod <- model.matrix(~gender+GeneExp_Subtype+days_to_birth, 
                    data=clinical_gbm[colnames(rna_gbm_corrected),])

batch <- as.factor(as.character(clinical_gbm[colnames(rna_gbm_corrected),
                                             'batch_number']))

rna_gbm_corrected <- ComBat(as.matrix(rna_gbm_corrected),
                            batch=batch,
                            mod=mod, 
                            prior.plot=TRUE)
```

Exploring the PCA from corrected data we can see that the batch effect have been removed.

```
pca_rna <- prcomp(t(rna_gbm_corrected))
var_prcomp <- pca_rna$sdev^2

# Plot PCs variance
pcvar <- data.frame(var=var_prcomp/sum(var_prcomp), pc=c(1:length(var_prcomp)))
ggplot(pcvar[1:15,], aes(x = pc)) + geom_line(aes(y=var)) + 
  labs(x='Principal Component', y='Explained Variance') + 
  geom_point(aes(y=var)) + 
  ggtitle('mRNA: Principal Components variance')
```

```
# Plot PCs scores
df <- data.frame(PC1=pca_rna$x[,1], PC2=pca_rna$x[,2],
                 clinical_gbm[colnames(rna_gbm_corrected),
                              c("GeneExp_Subtype","gender","batch_number")])

ggplot(df) +
  geom_point(aes(x=PC1, y=PC2,color=factor(GeneExp_Subtype)),size=5,shape=20) +
  guides(color=guide_legend("GeneExp_Subtype"), 
         fill=guide_legend("GeneExp_Subtype")) +
  labs(x='PC1 (12.8%)', y='PC2 (8.4%)') +
  ggtitle("Principal Components from batch-adjusted mRNA data") +
  theme(legend.position="bottom")
```

```
ggplot(df) +
  geom_point(aes(x=PC1, y=PC2, color=factor(batch_number)), size=5, shape=20) +
  guides(color=guide_legend("batch_number"),
         fill=guide_legend("batch_number")) +
  labs(x='PC1 (12.8%)', y='PC2 (8.4%)') +
  ggtitle("Principal Components from batch-adjusted mRNA data") +
  theme(legend.position="bottom")
```

Now, the association analysis shows how the batch effect is clearly removed from our data and the main variability is related with gene expression subtype.

```
pcaCorrelation(clinical=clinical_gbm[colnames(rna_gbm_corrected),],
               clin_var=clin_var, pca_data=pca_rna$x, maxlogPvalue = 10)
```

##### 2.4.2 miRNA data (`mirna_gbm_corrected`)

```
mod <- model.matrix(~gender+GeneExp_Subtype+days_to_birth, 
                    data=clinical_gbm[colnames(mirna_gbm_corrected),])

batch <- as.factor(as.character(clinical_gbm[colnames(mirna_gbm_corrected),
                                             'batch_number']))

mirna_gbm_corrected <- ComBat(as.matrix(mirna_gbm_corrected),
                              batch=batch,
                              mod=mod, 
                              prior.plot=TRUE)
```

Exploring the PCA from corrected data we can see that the batch effect have been removed. However, in the case of miRNA, the results are equivalent to before batch adjustment, as no batch-effect was detected in the initial data set.

```
#miRNA
pca_mirna <- prcomp(t(mirna_gbm_corrected))
var_prcomp <- pca_mirna$sdev^2


# Plot PCs variance
pcvar <- data.frame(var=var_prcomp/sum(var_prcomp), pc=c(1:length(var_prcomp)))
ggplot(pcvar[1:15,], aes(x = pc)) + geom_line(aes(y=var)) + 
  labs(x='Principal Component', y='Explained Variance') + 
  geom_point(aes(y=var)) + 
  ggtitle('miRNA: Principal Components variance')
```

```
# Plot PCs scores
df <- data.frame(PC1=pca_mirna$x[,1], PC2=pca_mirna$x[,2],
                 clinical_gbm[colnames(mirna_gbm_corrected),
                              c("GeneExp_Subtype","gender","batch_number")])

ggplot(df) +
  geom_point(aes(x=PC1, y=PC2, color=factor(GeneExp_Subtype)),size=5,shape=20) +
  guides(color=guide_legend("GeneExp_Subtype"),
         fill=guide_legend("GeneExp_Subtype")) +
  labs(x='PC1 (19.5%)', y='PC2 (9.5%)') +
  ggtitle("Principal Components from batch-adjusted miRNA data") +
  theme(legend.position="bottom")
```

```
ggplot(df) +
  geom_point(aes(x=PC1, y=PC2, color=factor(batch_number)), size=5, shape=20) +
  guides(color=guide_legend("batch_number"),
         fill=guide_legend("batch_number")) +
  labs(x='PC1 (19.5%)', y='PC2 (9.5%)') +
  ggtitle("Principal Components from batch-adjusted miRNA data") +
  theme(legend.position="bottom")
```

```
pcaCorrelation(clinical=clinical_gbm[colnames(mirna_gbm_corrected),],
               clin_var=clin_var, pca_data=pca_mirna$x, maxlogPvalue = 10)
```

##### 2.4.3 Methylation data (`logit_met_gbm`)

As methylation data not follows a gausian distribution, a non-parametric ComBat is applied.

```
mod <- model.matrix(~gender+GeneExp_Subtype+days_to_birth, 
                    data=clinical_gbm[colnames(logit_met_gbm),])

batch <- as.factor(as.character(clinical_gbm[colnames(logit_met_gbm),
                                             'batch_number']))
  
logit_met_gbm_corrected <- ComBat(logit_met_gbm,
                                  batch=batch, 
                                  mod=mod, 
                                  prior.plot=TRUE, 
                                  par.prior=FALSE, 
                                  BPPARAM=MulticoreParam())
```

Note: the non-parametric ComBat spends long time (>4days with our resources) to compute the corrected matrix.

Exploring the PCA from corrected data we can see that the batch effect have been removed.

```
pca_met <- prcomp(t(logit_met_gbm_corrected))
var_prcomp <- pca_met$sdev^2

# Plot PCs variance
pcvar <- data.frame(var=var_prcomp/sum(var_prcomp), pc=c(1:length(var_prcomp)))
ggplot(pcvar[1:15,], aes(x = pc)) + geom_line(aes(y=var)) + 
  labs(x='Principal Component', y='Explained Variance') + 
  geom_point(aes(y=var)) + 
  ggtitle('Methylation: Principal Components variance')
```

```
# Plot PCs scores
df <- data.frame(PC1=pca_met$x[,1], PC2=pca_met$x[,2],
                 clinical_gbm[colnames(logit_met_gbm_corrected),
                              c("GeneExp_Subtype","gender","batch_number")])

ggplot(df) +
  geom_point(aes(x=PC1, y=PC2,color=factor(GeneExp_Subtype)),size=5,shape=20) +
  guides(color=guide_legend("GeneExp_Subtype"),
         fill=guide_legend("GeneExp_Subtype")) +
  labs(x='PC1 (14.8%)', y='PC2 (11.7%)') +
  ggtitle("Principal Components from batch-adjusted Methylation data") +
  theme(legend.position="bottom")
```

```
ggplot(df) +
  geom_point(aes(x=PC1, y=PC2, color=factor(batch_number)), size=5, shape=20) +
  guides(color=guide_legend("batch_number"),
         fill=guide_legend("batch_number")) +
  labs(x='PC1 (14.8%)', y='PC2 (11.7%)') +
  ggtitle("Principal Components from batch-adjusted Methylation data") +
  theme(legend.position="bottom")
```

Now, the association analysis shows how the batch effect is clearly removed from our data and the main variability is related with gene expression subtype and survival outcome.

```
pcaCorrelation(clinical=clinical_gbm[colnames(logit_met_gbm_corrected),],
               clin_var=clin_var, pca_data=pca_met$x, maxlogPvalue = 10)
```

#### 2.5 Characterization of the data: Individual exploration

After data pre-process we are ready to work with the different omics.
Batch effect has been corrected in each data-type and the appropiate transformation has been applied according the data-type.

As seen in the previous exploration after batch correction (see previous section), the graphical representation of explained variance for the first fifteen components show that variance explained for the first components is not quite high, moving between 10 and 20%. This fact highlights the data heterogeneity and the missing of a driving variable explaining the most variability of our data. As expected, high association with the gene expression subtypes, previously defined, is found with the first PCs of mRNA, but also in miRNA and methylation. Relevant variables associated with first components are survival outcome, histological type, prior glioma, age and gender.

#### 2.6 Characterization of the data: Joint exploration

##### 2.6.1 Common samples

As we have different number of samples in each omic dataset, we need to determine the share samples between omics.

```
# Load require packages
library(VennDiagram)
library(plotrix)
```

```
# Venn Diagram
m <- list(rna=colnames(rna_gbm_corrected), 
          mirna=colnames(mirna_gbm_corrected), 
          met=colnames(logit_met_gbm_corrected))

vennPlot <- venn.diagram(m, NULL, fill=c('red','green','blue'), 
                         cat.fontface=4, lty=2, cex=1.2, cat.cex=1.2)
grid.newpage()
grid.draw(vennPlot)
```

We have 515 samples (almost 100%) with both mRNA and miRNA data, however, just 83 samples have mRNA, miRNA and methylation data.

##### 2.6.2 OmicsPCA exploration

Based on first data exploration and pre-process, we know that the most rellevant variables in our omics data-types after batch correction are gene expression subtype, survival outcome and age.

But, how much the different data-types are “explaining the samples in the same way”? To be more specific: are there share components or not? How many?

Let’s perform a joint exploration using the `omicsCompAnalysis` function from `STATegRa` package.

```
# Load require packages
library(STATegRa)
library(MASS)
library(gridExtra)
```

###### 2.6.2.1 Data scaling

Before start the multi-omic approach, mean centred of each feature and the data set normalization to unit sum of squares (forbenious normalization) is recommended to make more comparabable the different omics data-sets.

```
#Calculate Forbenious normalization
frobenius_rna <- norm(as.matrix(rna_gbm_corrected), type="F")
frobenius_mirna <- norm(as.matrix(mirna_gbm_corrected), type="F")
frobenius_met <- norm(as.matrix(logit_met_gbm_corrected), type="F")

#Mean-centring and division by Frobenius normalization factor
rna_gbm_norm <- t(scale(t(rna_gbm_corrected), scale=FALSE))/frobenius_rna
mirna_gbm_norm <- t(scale(t(mirna_gbm_corrected), scale=FALSE))/frobenius_mirna
met_gbm_norm <- t(scale(t(logit_met_gbm_corrected), scale=FALSE))/frobenius_met

##Mean-centring without Frobenius normalization
rna_gbm_norm2 <- t(scale(t(rna_gbm_corrected), scale=FALSE))
mirna_gbm_norm2 <- t(scale(t(mirna_gbm_corrected), scale=FALSE))
met_gbm_norm2 <- t(scale(t(logit_met_gbm_corrected), scale=FALSE))
```

Then, we need to keep the common samples between the different data sets.

```
#Common samples between mRNA and miRNA
ids_rna_mirna <- intersect(colnames(rna_gbm_corrected), colnames(mirna_gbm_corrected)) 
length(ids_rna_mirna)
```

```
## [1] 515
```

```
#Common samples between mRNA and Methylation
ids_rna_met <- intersect(colnames(rna_gbm_corrected), colnames(logit_met_gbm_corrected))
length(ids_rna_met)
```

```
## [1] 83
```

###### 2.6.2.2 Model selection

Now, we are ready to explore if there are share components between data-types.

The first step is to determine the number of share/common and individual components between omics.

How to determine these numbers is still an open question nowadays, so the user can explore different methods like jive, pca-gca and pESCA, between others. Following, we show how to perform the model selection based on these three methods.

###### 2.6.2.2.1 JIVE

```
# Load require packages
library(r.jive)
```

```
#Using r.jive

################################################
## mRNA + miRNA (already centered)            ##
################################################
Data <- list(mRNA=rna_gbm_norm[,ids_rna_mirna],
             miRNA=mirna_gbm_norm[,ids_rna_mirna])
rjive_rna_mirna <- jive(Data, scale = FALSE, center = FALSE) 


######################################################
## mRNA + methylation (already centered)            ##
######################################################
Data <- list(mRNA=rna_gbm_norm[,ids_rna_met],
              met=met_gbm_norm[,ids_rna_met])
rjive_rna_met <- jive(Data, scale = FALSE, center = FALSE)
```

###### 2.6.2.2.2 PCA-GCA

```
# Load require packages
library(RegularizedSCA)

#Using pca-gca

################################################
## mRNA + miRNA (already centered and scaled) ##
################################################
data <- cbind(t(rna_gbm_norm[,ids_rna_mirna]), 
              t(mirna_gbm_norm[,ids_rna_mirna]))

# Number of variables
Jk <- c(nrow(rna_gbm_norm), nrow(mirna_gbm_norm))

# Select components
pca_gca(data, Jk, cor_min = .7)

######################################################
## mRNA + methylation (already centered and scaled) ##
######################################################
data <- cbind(t(rna_gbm_norm[,ids_rna_met]), 
              t(met_gbm_norm[,ids_rna_met]))

# Number of variables
Jk <- c(nrow(rna_gbm_norm), nrow(met_gbm_norm))

# Select components
pca_gca(data, Jk, cor_min = .7)
```

###### 2.6.2.2.3 pESCA

```
# Load require packages
library(RpESCA)
library(RSpectra)

#Using RpESCA

################################################
## mRNA + miRNA (already centered)            ##
################################################

dataSets <- list(rna=t(rna_gbm_norm2[,ids_rna_mirna]), 
                 mirna=t(mirna_gbm_norm2[,ids_rna_mirna]))

dataTypes <- 'GG'

# used parameters
opts <- list()
opts$tol_obj <- 1E-4
opts$quiet <- 1

# save the estimated alphas, selected number of PCs and cvErrors
alphas <- rep(NA,2)
R_selected_list <- as.list(1:2)
cvErrors_list <- as.list(1:2)

# alpha estimation for each data set
for (i in 1:length(dataSets)){
  # index ith data set
  X <- dataSets[[i]]
  
  # alpha estimation procedure
  alpha_est <- alpha_estimation(X,K=3,Rs = 5:20,opts=opts)
  alphas[i] <- alpha_est$alphas_mean
  R_selected_list[[i]] <- alpha_est$R_CV
  cvErrors_list[[i]] <- alpha_est$cvErrors
}
names(alphas) <- paste0('alpha', 1:2)

## Setting the parameters for the pESCA model

# concave function and its hyper-parameter
fun_concave <- 'gdp'; gamma <- 1;
penalty = 'L2' # concave L2 norm penalty

# Parameters of a pESCA with concave L2norm penalty model
opts <- list()
opts$gamma <- gamma  # hyper-parameter for the used penalty
opts$rand_start <- 0
opts$tol_obj <- 1e-6 # stopping criteria
opts$maxit   <- 500
opts$alphas  <- alphas
#opts$R  # components used Default 0.5*min(I,J) - Minimum number of variables.
opts$thr_path <- 0 # generaint thresholding path or not
opts$quiet <- 1

# Model selection pESCA conave L2 norm penalty
nTries <- 15
lambdas_CV <- log10_seq(from=1, to=500, length.out=nTries)

result_CV <- pESCA_CV(dataSets, dataTypes,
                      lambdas_CV, 
                      penalty=penalty, 
                      fun_concave=fun_concave, 
                      opts=opts)


# select the model with minimum CV error
index_min_cv <- which.min(result_CV$cvErrors_mat[,1])

# cvErrors during the model selection process
result_CV$cvErrors_mat

# selected value of lambda
lambdas_CV[index_min_cv]

# fit the final model
lambdas_opt <- rep(lambdas_CV[index_min_cv],length(dataSets))
opts_opt <- result_CV$inits[[index_min_cv]]
opts_opt$tol_obj <- 1E-8 # using high precision model


pESCA_L2 <- pESCA(dataSets = dataSets,
                          dataTypes = dataTypes,
                          lambdas = lambdas_opt,
                          penalty = penalty,
                          fun_concave= fun_concave,
                          opts = opts_opt)


mu <- pESCA_L2$mu
A <- pESCA_L2$A
B <- pESCA_L2$B
S <- pESCA_L2$S

# estimated variation explained ratios for each data set or the full data set
pESCA_L2$varExpTotals

# estimated variation explained ratios of each PC for each data set or the full data set
pESCA_L2$varExpPCs

#Cut-off 1% 
sel1 <- which(pESCA_L2$varExpPCs[1,]>1) #13
sel2 <- which(pESCA_L2$varExpPCs[2,]>1) #21
intersect(sel1,sel2)
# 7 common components
# 6 dist mRNA + 14 miRNA
# Total: 27 components

# Cut-off 5%
sel1 <- which(pESCA_L2$varExpPCs[1,]>5) #5
sel2 <- which(pESCA_L2$varExpPCs[2,]>5) #2
intersect(sel1,sel2)
# 1 common components
# 4 dist mRNA + 1 miRNA
# Total: 6 components


################################################
## mRNA + Methylation (already centered)      ##
################################################

dataSets <- list(rna=t(rna_gbm_norm2[,ids_rna_met]), 
                 met=t(met_gbm_norm2[,ids_rna_met]))

dataTypes <- 'GG'

# used parameters
opts <- list()
opts$tol_obj <- 1E-4
opts$quiet <- 1

# save the estimated alphas, selected number of PCs and cvErrors
alphas <- rep(NA,2)
R_selected_list <- as.list(1:2)
cvErrors_list <- as.list(1:2)

# alpha estimation for each data set
for (i in 1:length(dataSets)){
  # index ith data set
  X <- dataSets[[i]]
  
  # alpha estimation procedure
  alpha_est <- alpha_estimation(X,K=3,Rs = 5:20,opts=opts)
  alphas[i] <- alpha_est$alphas_mean
  R_selected_list[[i]] <- alpha_est$R_CV
  cvErrors_list[[i]] <- alpha_est$cvErrors
}
names(alphas) <- paste0('alpha', 1:2)

## Setting the parameters for the pESCA model

# concave function and its hyper-parameter
fun_concave <- 'gdp'; gamma <- 1;
penalty = 'L2' # concave L2 norm penalty

# Parameters of a pESCA with concave L2norm penalty model
opts <- list()
opts$gamma <- gamma  # hyper-parameter for the used penalty
opts$rand_start <- 0
opts$tol_obj <- 1e-6 # stopping criteria
opts$maxit   <- 500
opts$alphas  <- alphas
#opts$R # components used Default 0.5*min(I,J) - Minimum number of variables.
opts$thr_path <- 0 # generaint thresholding path or not
opts$quiet <- 1

# Model selection pESCA conave L2 norm penalty
nTries <- 15
lambdas_CV <- log10_seq(from=1, to=500, length.out=nTries)

result_CV <- pESCA_CV(dataSets, dataTypes,
                      lambdas_CV, 
                      penalty=penalty, 
                      fun_concave=fun_concave, 
                      opts=opts)


# select the model with minimum CV error
index_min_cv <- which.min(result_CV$cvErrors_mat[,1])

# cvErrors during the model selection process
result_CV$cvErrors_mat

# selected value of lambda
lambdas_CV[index_min_cv]

# fit the final model
lambdas_opt <- rep(lambdas_CV[index_min_cv],length(dataSets))
opts_opt <- result_CV$inits[[index_min_cv]]
opts_opt$tol_obj <- 1E-8 # using high precision model


pESCA_L2 <- pESCA(dataSets = dataSets,
                          dataTypes = dataTypes,
                          lambdas = lambdas_opt,
                          penalty = penalty,
                          fun_concave= fun_concave,
                          opts = opts_opt)

mu <- pESCA_L2$mu
A <- pESCA_L2$A
B <- pESCA_L2$B
S <- pESCA_L2$S

# estimated variation explained ratios for each data set or the full data set
pESCA_L2$varExpTotals

# estimated variation explained ratios of each PC for each data set or the full data set
pESCA_L2$varExpPCs

#Cut-off 1%
sel1 <- which(pESCA_L2$varExpPCs[1,]>1) #2
sel2 <- which(pESCA_L2$varExpPCs[2,]>1) #9
intersect(sel1,sel2)
# 1 common components
# 1 dist mRNA + 8 methylation
# Total: 10 components

#Cut-off 5%
sel1 <- which(pESCA_L2$varExpPCs[1,]>5) #2
sel2 <- which(pESCA_L2$varExpPCs[2,]>5) #3
intersect(sel1,sel2)
# 1 common components
# 1 dist mRNA + 2 methylation
# Total: 4 components
```

Here, there is the table summarizing the output of the different methods.

|  | Component | r.JIVE | PCA-GCA (1%) | PCA-GCA (5%) | pESCA(1%) | pESCA (5%) |
| --- | --- | --- | --- | --- | --- | --- |
| mRNA + miRNA |  |  |  |  |  |  |
|  | Common | 1 | 3 | NA | 7 | 1 |
|  | Dist. mRNA | 50 | 8 | NA | 6 | 4 |
|  | Dist. miRNA | 19 | 5 | NA | 14 | 1 |
| mRNA + met |  |  |  |  |  |  |
|  | Common | 3 | NA | NA | 1 | 1 |
|  | Dist. mRNA | 17 | NA | NA | 1 | 1 |
|  | Dist. miRNA | 25 | NA | NA | 8 | 2 |

Again, we can observe that the results from different methods have relevant divergencies. Mainly, we can say that share variation between data is not too large, but it seems that there’s something explained in the same way. To follow with the analysis, we will explore the omicsPCA based on the output of pESCA (1%).

###### 2.6.2.3 Sub-space recovery

Based on the number of components identified (pESCA) we can recover the subspace.

It is interesting to take a look on the common and distinctive components identified thorugh this method.

```
################################################
## mRNA + miRNA (already centered)            ##
################################################

# Explore the common and distinctive components and those variation explained
# Common components
cc <- intersect(sel1,sel2)
pESCA_L2$varExpPCs[,cc]
```

```
##             PC1      PC3      PC4      PC6      PC7      PC9     PC11
## X_1    12.31097 6.648208 5.547511 2.958637 2.419148 1.833125 1.282475
## X_2    10.02265 4.273318 3.941567 1.445866 3.060845 1.418305 1.353780
## X_full 12.16291 6.494543 5.443599 2.860755 2.460668 1.806284 1.287089
```

```
# Distinctive components
dist_rna <- sel1[!sel1%in%cc] #rna
pESCA_L2$varExpPCs[,dist_rna]
```

```
##              PC2       PC5       PC8      PC10     PC12     PC13
## X_1    8.0683705 5.3879052 2.0484025 1.4743292 1.195001 1.091788
## X_2    0.5333225 0.3595895 0.6113683 0.8597992 0.000000 0.000000
## X_full 7.5808210 5.0625519 1.9554203 1.4345665 1.117680 1.021145
```

```
dist_mirna <- sel2[!sel2%in%cc] #mirna
pESCA_L2$varExpPCs[,dist_mirna]
```

```
##             PC17      PC19       PC25      PC43      PC69      PC74     PC113
## X_1    0.6366036 0.6955921  0.0000000 0.2801706 0.1707055 0.2450968 0.0000000
## X_2    2.0762611 1.3458879 11.3074793 4.6778476 1.0128559 1.3851853 2.4789441
## X_full 0.7297555 0.7376690  0.7316418 0.5647189 0.2251962 0.3188653 0.1603982
##             PC118     PC129     PC130     PC134     PC136      PC139      PC147
## X_1    0.00000000 0.0000000 0.0000000 0.0000000 0.0000000 0.00000000 0.00000000
## X_2    1.17557417 2.1242175 3.1881778 1.6015304 1.6993565 1.27223736 1.16930411
## X_full 0.07606463 0.1374459 0.2062886 0.1036258 0.1099556 0.08231915 0.07565893
```

```
# Keep de common and distinctive components
common_pc <- pESCA_L2$A[,cc]
colnames(common_pc) <- colnames(pESCA_L2$varExpPCs[,cc])

dist_rna_pc <- pESCA_L2$A[,dist_rna]
colnames(dist_rna_pc) <- names(dist_rna)

dist_mirna_pc <- pESCA_L2$A[,dist_mirna]
colnames(dist_mirna_pc) <- names(dist_mirna)


# metadata for common samples
clin <- clinical_gbm[ids_rna_mirna,]

# PCA plot
require(ggplot2)

df <- as.data.frame(common_pc)
df$group <- clin$GeneExp_Subtype
df$batch <- clin$batch_number

#plot of first common components by gene expression subtype
p <- ggplot(df,aes_string("PC1","PC3",color="group"))
p + geom_point(size=5, shape=20) + ggtitle("PCA by Gene Expression Subtype") + theme(legend.position="bottom")
```

```
#plot of first common components by batch
p <- ggplot(df,aes_string("PC1","PC3",color="batch"))
p + geom_point(size=5, shape=20) + ggtitle("PCA by batch") + theme(legend.position="bottom")
```

Let’s explore how relate these common and distinctive components with the clinical variables.

```
df <- data.frame(common_pc,dist_rna_pc,dist_mirna_pc)
colnames(df) <- c(paste("common_",colnames(common_pc),sep=""),
                  paste("dist_rna_",colnames(dist_rna_pc),sep=""),
                  paste("dist_mirna_",colnames(dist_mirna_pc),sep=""))

pcaCorrelation(clinical=clin, clin_var, df, maxlogPvalue =10, maxPC=ncol(df), pc.names=TRUE)
```

The main common variability for mRNA + miRNA is associated with `GeneExp_Subtype`, `survivalOutcome`, `days_to_birth` and `age_at_initial_pathologic_diagnosis`. No batch effect is identified. Moreover, the main variability explained by mRNA distinctive components are associated with `GeneExp_Subtype`, too.

```
################################################
## mRNA + methylation (already centered)      ##
################################################

# Explore the common and distinctive components and those variation explained
# Common components
cc <- intersect(sel1,sel2)
pESCA_L2$varExpPCs[,cc]
```

```
##       X_1       X_2    X_full 
##  8.126533 11.062577 10.928992
```

```
# Distinctive components
dist_rna <- sel1[!sel1%in%cc] #rna
pESCA_L2$varExpPCs[,dist_rna]
```

```
##       X_1       X_2    X_full 
## 12.012788  0.000000  0.546561
```

```
dist_met <- sel2[!sel2%in%cc] #met
pESCA_L2$varExpPCs[,dist_met]
```

```
##             PC1      PC3      PC4      PC5      PC6      PC7      PC8      PC9
## X_1     0.00000 0.000000 0.000000 0.000000 0.000000 0.000000 0.000000 0.000000
## X_2    14.81974 8.101900 3.433565 2.998602 2.848740 2.132701 1.956806 1.764833
## X_full 14.14547 7.733277 3.277344 2.862171 2.719128 2.035667 1.867774 1.684536
```

```
# Keep de common and distinctive components
common_pc <- data.frame(pESCA_L2$A[,cc])
colnames(common_pc) <- paste("PC", cc, sep="")

dist_rna_pc <- data.frame(pESCA_L2$A[,dist_rna])
colnames(dist_rna_pc) <- names(dist_rna)

dist_met_pc <- pESCA_L2$A[,dist_met]
colnames(dist_met_pc) <- names(dist_met)


# metadata for common samples
clin <- clinical_gbm[ids_rna_met,]

# PCA plot
require(ggplot2)

df <- as.data.frame(common_pc)
df$group <- clin$GeneExp_Subtype
df$batch <- clin$batch_number

# As only one common component is found, an auxiliar varaible is defined in order to visualize the first components.
df$auxiliar <- runif(nrow(df),min(df$PC2, na.rm=TRUE),max(df$PC2, na.rm=TRUE))

#plot of first common components by gene expression subtype
p <- ggplot(df,aes_string("PC2","auxiliar",color="group"))
p + geom_point(size=5, shape=20) + ggtitle("PCA by Gene Expression Subtype") + theme(legend.position="bottom")
```

```
#plot of first common components by batch
p <- ggplot(df,aes_string("PC2","auxiliar",color="batch"))
p + geom_point(size=5, shape=20) + ggtitle("PCA by batch") + theme(legend.position="bottom")
```

Let’s explore how relate these common and distinctive components with the clinical variables.

```
df <- data.frame(common_pc,dist_rna_pc,dist_met_pc)
colnames(df) <- c(paste("common_",colnames(common_pc),sep=""),
                  paste("dist_rna_",colnames(dist_rna_pc),sep=""),
                  paste("dist_met_",colnames(dist_met_pc),sep=""))

pcaCorrelation(clinical=clin, clin_var, df, maxlogPvalue =10, maxPC=ncol(df), pc.names=TRUE)
```

Again, the main common variability for mRNA + methylation is associated with `GeneExp_Subtype` and `survivalOutcome`. Moreover, distinctive components for methylation are associated with `days_to_birth` and `age_at_initial_pathologic_diagnosis`.

#### 2.7 Integrative differential analysis by NPC

Now, we want to identify genes that, according to all modalities considered as a whole (mRNA, miRNA, methylation, etc.), are either deregulated or associated to an outcome of interest.

From the previous steps (individual omics exploration and joint exploration) we will be considering (1) which co-variates are necessary to include and (2) which analysis do we want to perform.

Let’s explore the combination of two omics, then NPC will provide two outcomes:

- The features that are differentiated in both omics, therefore we are looking into coordinated effects.
- The features that are differentiated in only one of the omics.

These kind of approach allows different number of samples between omics, so we will explore both escenarios.

##### 2.7.1 mRNA + miRNA

First of all we need to have a mapping file mathcing omics.

###### 2.7.1.1 Mapping file

```
#Load the mapping info
#load('hsa-vtm-gene.Rdata')
#the mapping can be obtained using SpidermiR package

#Identify the miRBase version of mapping file and data
library(miRNAmeConverter)

miRNAs <- rownames(mirna_gbm_corrected)
miRNAs_mapping <- names(id)
nc <- MiRNANameConverter()
assessVersion(nc, miRNAs, verbose = FALSE) #version 9.2
```

```
##    version frequency
## 1      9.2       534
## 2      9.1       534
## 3      9.0       533
## 4      8.2       516
## 5      8.1       515
## 6     10.0       430
## 7     10.1       424
## 8     11.0       422
## 9     14.0       420
## 10    13.0       420
## 11    12.0       420
## 12    15.0       419
## 13    16.0       416
## 14    17.0       415
## 15     8.0       360
## 16     7.1       340
## 17    18.0       245
## 18    19.0       218
## 19     6.0       208
## 20    20.0       174
## 21    21.0       173
## 22    22.0       165
```

```
assessVersion(nc, miRNAs_mapping, verbose = FALSE) #version 20
```

```
##    version frequency
## 1     20.0       573
## 2     21.0       569
## 3     22.0       560
## 4     19.0       546
## 5     18.0       533
## 6     17.0       380
## 7     16.0       372
## 8     15.0       359
## 9     14.0       341
## 10    13.0       339
## 11    12.0       338
## 12    11.0       337
## 13    10.1       321
## 14    10.0       314
## 15     9.2       228
## 16     9.1       228
## 17     9.0       227
## 18     8.2       220
## 19     8.1       220
## 20     8.0       181
## 21     7.1       178
## 22     6.0       145
```

```
#convert miRNA annotation from our miRNA matrix from v.9.2 to v.20
library(anamiR)
new_mirna_matrix <- miR_converter(mirna_gbm_corrected, remove_old = TRUE, original_version=9.2, latest_version = 20)
#Original data 534 miRNAs, now 527 miRNAs


#restricting mapping info to measured miRNA
id_mapping <- id[intersect(names(id), rownames(new_mirna_matrix))] 
#from the initial 527 miRNA features we get 332 miRNA shared with mapping info

#creating the miRNA association tables
mirna2GeneMap <- stack(id_mapping) #convert in a data.frame. Each line has a gene and one mirna. 
names(mirna2GeneMap) <- c('Gene', 'miRNA')
#Dimension: 33324 x 2 (33,324 pairs)

head(mirna2GeneMap)
```

```
##     Gene         miRNA
## 1  ACAD8 hsa-let-7a-5p
## 2   ACTB hsa-let-7a-5p
## 3 ACVR1B hsa-let-7a-5p
## 4   ADSL hsa-let-7a-5p
## 5   AGO1 hsa-let-7a-5p
## 6   AGO2 hsa-let-7a-5p
```

```
#creating the mRNA association tables
expr2GeneMap <- data.frame(measurement = rownames(rna_gbm_corrected), 
                           Gene = rownames(rna_gbm_corrected))
#Dimension: 12,042 x 2

head(expr2GeneMap)
```

```
##   measurement    Gene
## 1       FSTL1   FSTL1
## 2        AACS    AACS
## 3       RPS11   RPS11
## 4     CREB3L1 CREB3L1
## 5       ELMO2   ELMO2
## 6       PNMA1   PNMA1
```

```
#creating the data mapping
dataMappingExprMirna <- combiningMappings(mappings = list(expr = expr2GeneMap, 
                                                          mirna = mirna2GeneMap),
                                          retainAll = TRUE, reference = 'Gene'); 
#Dimension: 24,665 x 3

head(dataMappingExprMirna)
```

```
##      expr           mirna   Gene
## 4     A2M  hsa-miR-122-5p    A2M
## 5     A2M   hsa-miR-98-5p    A2M
## 6     A2M  hsa-miR-128-3p    A2M
## 7  A4GALT hsa-miR-193b-3p A4GALT
## 9    AAAS   hsa-miR-16-5p   AAAS
## 10   AAAS hsa-miR-193b-3p   AAAS
```

We can see that based on our mRNA and miRNA analysis, we have 24,665 pairs in total, it corresponds to 7,814 unique genes (64.9% from the whole mRNA dataset) and 323 unique miRNAs (61.3% from teh whole miRNA dataset).

Then, we define the relevant clinical variables in each scenario. Our variable of interest in the case of GBM is the `survivalOtucome`. Moreover, `days_to_birth` is included as a confounder.

```
clinicalVariables <- list(expr = c('days_to_birth', 'survivalOutcome'), 
                          met = c('days_to_birth', 'survivalOutcome'),
                          mirna = c('days_to_birth', 'survivalOutcome'));
```

Then, we can explore the combination of these two omics using a parametric or non parametric combination (PC or NPC). Moreover, we can perform the analysis on all available samples or just for those that are common between datasets.
So, these are the different scenarios that we can have:

- Overlapping samples

  - Parametric Combination of p-values (`omicsPC`)
  - Non Parametric Combination of p-values (`omicsNPC`)
- All samples

  - Parametric Combination of p-values (`omicsPC`)
  - Non Parametric Combination of p-values (`omicsNPC`)

As first approach, we will start to explore the combination just for the overlapping samples, followed by the all samples approach.

###### 2.7.1.2 Overlaping samples

```
#restrict the datasets to the elements of the data mapping and those samples that are common between omics
exprTMP <- rna_gbm_corrected[unique(na.omit(dataMappingExprMirna$expr)), ids_rna_mirna]; #7,814 x 515

mirnaTMP <- new_mirna_matrix[unique(na.omit(dataMappingExprMirna$mirna)), ids_rna_mirna]; #323 x 515

#specifying the data types.
dataTypesExprMirna <- list(ttCoxphContinuous,ttCoxphContinuous)

#preparing the datasets
exprTMP <- createOmicsExpressionSet(Data = as.matrix(exprTMP), 
                                    pData = clinical_gbm[ids_rna_mirna,clinicalVariables[['expr']]])
names(pData(exprTMP)) <- c("age","outcome")

mirnaTMP <- createOmicsExpressionSet(Data = as.matrix(mirnaTMP), 
                                     pData = clinical_gbm[ids_rna_mirna,clinicalVariables[['mirna']]])
names(pData(mirnaTMP)) <- c("age","outcome")

dataInputExprMirna <- list(expr = exprTMP, mirna = mirnaTMP)
```

###### 2.7.1.2.1 Parametric Combination

```
set.seed(12345)
omicsPCRes_cPC <- omicsPC(dataInput = dataInputExprMirna,
                       dataMapping = dataMappingExprMirna,
                       dataTypes = dataTypesExprMirna,
                       verbose = TRUE)
```

The output of `omicsPC` provides information regarding p-value and adjusted p-value from both individual and combined analysis.

```
res <- omicsPCRes_cPC
str(res)
```

```
## List of 4
##  $ expr     : num [1:7814, 1:6] 0.1343 -0.2055 -0.0694 0.2895 0.0531 ...
##   ..- attr(*, "dimnames")=List of 2
##   .. ..$ : chr [1:7814] "A2M" "A4GALT" "AAAS" "AACS" ...
##   .. ..$ : chr [1:6] "coef" "exp(coef)" "se(coef)" "z" ...
##  $ mirna    : num [1:323, 1:6] 1.0453 -0.0639 -0.0494 0.0683 -0.0126 ...
##   ..- attr(*, "dimnames")=List of 2
##   .. ..$ : chr [1:323] "hsa-miR-122-5p" "hsa-miR-98-5p" "hsa-miR-128-3p" "hsa-miR-193b-3p" ...
##   .. ..$ : chr [1:6] "coef" "exp(coef)" "se(coef)" "z" ...
##  $ pvaluesPC:'data.frame':   24665 obs. of  8 variables:
##   ..$ expr     : chr [1:24665] "A2M" "A2M" "A2M" "A4GALT" ...
##   ..$ mirna    : chr [1:24665] "hsa-miR-122-5p" "hsa-miR-98-5p" "hsa-miR-128-3p" "hsa-miR-193b-3p" ...
##   ..$ Fisher   : num [1:24665] 0.022 0.0457 0.0495 0.4351 0.8912 ...
##   ..$ Liptak   : num [1:24665] 0.0179 0.0626 0.0743 0.3432 0.8693 ...
##   ..$ Tippett  : num [1:24665] 0.0317 0.0317 0.0317 0.775 1 ...
##   ..$ Benjamini: num [1:24665] 0.0317 0.0317 0.0317 0.3878 0.8893 ...
##   ..$ Simes    : num [1:24665] 0.0317 0.0317 0.0317 0.3878 0.8893 ...
##   ..$ Sidak    : num [1:24665] 0.0315 0.0315 0.0315 0.6249 0.8722 ...
##  $ qvaluesPC:'data.frame':   24665 obs. of  8 variables:
##   ..$ expr     : chr [1:24665] "A2M" "A2M" "A2M" "A4GALT" ...
##   ..$ mirna    : chr [1:24665] "hsa-miR-122-5p" "hsa-miR-98-5p" "hsa-miR-128-3p" "hsa-miR-193b-3p" ...
##   ..$ Fisher   : num [1:24665] 0.251 0.31 0.319 0.755 0.974 ...
##   ..$ Liptak   : num [1:24665] 0.249 0.391 0.414 0.728 0.963 ...
##   ..$ Tippett  : num [1:24665] 0.252 0.252 0.252 1 1 ...
##   ..$ Benjamini: num [1:24665] 0.251 0.251 0.251 0.729 0.964 ...
##   ..$ Simes    : num [1:24665] 0.251 0.251 0.251 0.729 0.964 ...
##   ..$ Sidak    : num [1:24665] 0.25 0.25 0.25 0.855 0.967 ...
```

```
#Nominal p-values plot individual omics
data_plot <- data.frame(pval=c(res$expr[,"Pr(>|z|)"],res$mirna[,"Pr(>|z|)"]), 
                        group=c(rep(names(res)[1], nrow(res$expr)), 
                                rep(names(res)[2], nrow(res$mirna))))

p1 <- ggplot(data_plot, aes(x=group, y=pval, fill=group)) +
  geom_violin(trim=FALSE)+
  labs(title='Plot of individual omics p-value from overlapping samples')

#FDR plot individual omics
data_plot <- data.frame(pval=c(res$expr[,"adj.P.Val"],res$mirna[,"adj.P.Val"]), 
                        group=c(rep(names(res)[1], nrow(res$expr)),
                                rep(names(res)[2], nrow(res$mirna))))

p2 <- ggplot(data_plot, aes(x=group, y=pval, fill=group)) +
  geom_violin(trim=FALSE)+
  labs(title='Plot of individual omics FDR from overlapping samples')


#Nominal p-value plot PC
data_plot <- data.frame(pval=unlist(res$pvaluesPC[,-c(1,2)]), 
                        combination_method=rep(colnames(res$pvaluesPC[,-c(1,2)]),
                                               each=nrow(res$pvaluesPC)))

p3 <- ggplot(data_plot, aes(x=combination_method, y=pval, 
                            fill=combination_method)) + 
  geom_violin(trim=FALSE)+
  labs(title='Plot of p-values in overlapping omicsPC')


#FDR plot PC
data_plot <- data.frame(pval=unlist(res$qvaluesPC[,-c(1,2)]), 
                        combination_method=rep(colnames(res$qvaluesPC[,-c(1,2)]),
                                               each=nrow(res$qvaluesPC)))

p4 <- ggplot(data_plot, aes(x=combination_method, y=pval,
                            fill=combination_method)) +
  geom_violin(trim=FALSE)+
  labs(title='Plot of FDR in overlapping omicsPC')


grid.arrange(p1, p2, p3, p4, ncol=2, nrow=2)
```

Now, based on FDR results we can check if there is an increase of statistical power when both omics are jointly analyzed.

Note: here we consider significant an FDR<0.05.

```
#significant genes
(sig_expr_cPC <- rownames(res$expr)[which(res$expr[,"adj.P.Val"]<0.05)])
```

```
## [1] "FAM46A"
```

```
# 1 genes

#significant miRNA
(sig_mirna_cPC <- rownames(res$mirna)[which(res$mirna[,"adj.P.Val"]<0.05)])
```

```
## [1] "hsa-miR-222-3p"
```

```
# 1 miRNA 

#Significant genes from mRNA + miRNA parametric combination (Fisher, FDR<0.05)
sig_expr_mirna_cPC <- res$qvaluesPC[which(res$qvaluesPC$Fisher<0.05),1:2] 
#397 pairs

#venn diagram
m1 <- list(rna=sig_expr_cPC, rna_fisher=unique(sig_expr_mirna_cPC$expr))

vennPlot1 <- venn.diagram(m1, NULL, fill=c('red','blue'), 
                         cat.fontface=4, lty=2, cex=1.2, cat.cex=1.2, 
                         main='Significant genes identified with individual (rna) or integration (rna_fisher)\n omics analysis: PC')
grid.newpage()
grid.draw(vennPlot1)
```

```
m2 <- list(mirna=sig_mirna_cPC, mirna_fisher=unique(sig_expr_mirna_cPC$mirna))

vennPlot2 <- venn.diagram(m2, NULL, fill=c('green','blue'), 
                         cat.fontface=4, lty=2, cex=1.2, cat.cex=1.2,
                         main='Significant miRNA identified with individual (mirna) or integration (mirna_fisher)\n omics analysis: PC')
grid.newpage()
grid.draw(vennPlot2)
```

When we perform the combination analysis we identify 337 new genes related with GBM survival and 45 miRNAs.

Here we can see the list of some of the new genes (n=337) identified based on Fisher combination.

```
setdiff(sig_expr_mirna_cPC$expr,sig_expr_cPC)[1:50]
```

```
##  [1] "ACACA"    "ACSL3"    "ACTB"     "ACTG1"    "AHSA1"    "ALG3"    
##  [7] "AMELX"    "ANXA2"    "AP1B1"    "AP3B1"    "APLP2"    "APOBEC3C"
## [13] "APOL2"    "APP"      "ARL4C"    "ATF5"     "ATP11B"   "ATP1A1"  
## [19] "ATP2A2"   "ATP7B"    "AUTS2"    "BAG3"     "BBC3"     "BICD1"   
## [25] "BPTF"     "C11orf57" "C8orf33"  "CABYR"    "CADM1"    "CANT1"   
## [31] "CANX"     "CAPRIN1"  "CASC3"    "CASKIN2"  "CASP3"    "CAST"    
## [37] "CBX2"     "CCDC47"   "CCR1"     "CCT3"     "CD44"     "CDKL5"   
## [43] "CDKN1B"   "CDKN1C"   "CEBPB"    "CENPT"    "CEP250"   "CFLAR"   
## [49] "CHL1"     "CKAP2"
```

And the new miRNAs (n=45) identified based on Fisher combination.

```
setdiff(sig_expr_mirna_cPC$mirna,sig_mirna_cPC)
```

```
##  [1] "hsa-miR-221-3p"  "hsa-miR-148a-3p" "hsa-miR-155-5p"  "hsa-miR-132-3p" 
##  [5] "hsa-miR-7-5p"    "hsa-miR-92a-3p"  "hsa-miR-26a-5p"  "hsa-miR-192-5p" 
##  [9] "hsa-miR-18a-3p"  "hsa-miR-21-5p"   "hsa-miR-34a-5p"  "hsa-miR-31-5p"  
## [13] "hsa-miR-124-3p"  "hsa-miR-335-5p"  "hsa-miR-181a-5p" "hsa-miR-27a-3p" 
## [17] "hsa-miR-27b-3p"  "hsa-miR-346"     "hsa-miR-128-3p"  "hsa-miR-218-5p" 
## [21] "hsa-miR-484"     "hsa-miR-9-5p"    "hsa-miR-19b-3p"  "hsa-miR-16-5p"  
## [25] "hsa-miR-143-3p"  "hsa-miR-26b-5p"  "hsa-miR-378a-3p" "hsa-miR-215-5p" 
## [29] "hsa-miR-25-3p"   "hsa-miR-98-5p"   "hsa-miR-30a-5p"  "hsa-miR-129-5p" 
## [33] "hsa-miR-1"       "hsa-miR-199a-3p" "hsa-miR-145-5p"  "hsa-miR-200b-3p"
## [37] "hsa-miR-671-5p"  "hsa-miR-596"     "hsa-miR-93-5p"   "hsa-let-7e-5p"  
## [41] "hsa-miR-203a"    "hsa-miR-196a-5p" "hsa-miR-299-5p"  "hsa-miR-331-3p" 
## [45] "hsa-miR-197-3p"
```

###### 2.7.1.2.2 Non Parametric Combination

```
set.seed(12345)

# Setting methods to combine pvalues
combMethods <- c("Fisher", "Liptak", "Tippett")
# Setting number of permutations
numPerms <- 1000
# Setting number of cores
numCores <- 4
# Setting omicsNPC to print out the steps that it performs.
verbose <- TRUE

omicsNPCRes_cNPC_rna_mirna <- omicsNPC(dataInput = dataInputExprMirna,
                         dataMapping = dataMappingExprMirna,
                         dataTypes = dataTypesExprMirna,
                         combMethods = combMethods, 
                         numPerms = numPerms,
                         numCores = numCores,
                         verbose = verbose)
```

The output of `omicsNPC` provides information regarding p-value and adjusted p-value from both individual and combined analysis.

```
res <- omicsNPCRes_cNPC_rna_mirna

#Nominal p-values plot individual omics
data_plot <- data.frame(pval=c(res$expr[,"Pr(>|z|)"],res$mirna[,"Pr(>|z|)"]), 
                        group=c(rep(names(res)[1], nrow(res$expr)), 
                                rep(names(res)[2], nrow(res$mirna))))

p1 <- ggplot(data_plot, aes(x=group, y=pval, fill=group)) +
  geom_violin(trim=FALSE)+
  labs(title='Plot of individual omics p-value from overlapping samples')

#FDR plot individual omics
data_plot <- data.frame(pval=c(res$expr[,"adj.P.Val"],res$mirna[,"adj.P.Val"]), 
                        group=c(rep(names(res)[1], nrow(res$expr)),
                                rep(names(res)[2], nrow(res$mirna))))

p2 <- ggplot(data_plot, aes(x=group, y=pval, fill=group)) +
  geom_violin(trim=FALSE)+
  labs(title='Plot of individual omics FDR from overlapping samples')


#Nominal p-value plot NPC
data_plot <- data.frame(pval=unlist(res$pvaluesNPC[,-c(1:4)]), 
                        combination_method=rep(colnames(res$pvaluesNPC[,-c(1:4)])
                                               , each=nrow(res$pvaluesNPC)))

p3 <- ggplot(data_plot, aes(x=combination_method, y=pval, fill=combination_method)) +
  geom_violin(trim=FALSE)+
  labs(title='Plot of p-values in overlapping omicsNPC')


#FDR plot NPC
data_plot <- data.frame(pval=unlist(res$qvaluesNPC[,-c(1:4)]), 
                        combination_method=rep(colnames(res$qvaluesNPC[,-c(1:4)])
                                               , each=nrow(res$qvaluesNPC)))

p4 <- ggplot(data_plot, aes(x=combination_method, y=pval, 
                            fill=combination_method)) +
  geom_violin(trim=FALSE)+
  labs(title='Plot of FDR in overlapping omicsNPC')


grid.arrange(p1, p2, p3, p4, ncol=2, nrow=2)
```

Now, based on FDR results we can check if there is an increase of statistical power when both omics are jointly analyzed.

Note: here we consider significant an FDR<0.05.

```
#Significant genes or miRNA from individual omics (FDR<0.05)

#significant genes
(sig_expr_cNPC <- rownames(res$expr)[which(res$expr[,"adj.P.Val"]<0.05)])
```

```
## [1] "FAM46A"
```

```
# 1 genes

#significant miRNA
(sig_mirna_cNPC <- rownames(res$mirna)[which(res$mirna[,"adj.P.Val"]<0.05)])
```

```
## [1] "hsa-miR-222-3p"
```

```
# 1 miRNA

#Significant genes from mRNA + miRNA non parametric combination (Fisher, FDR<0.05)
(sig_expr_mirna_cNPC <- res$qvaluesNPC[which(res$qvaluesNPC[,'Fisher']<0.05),1:2])
```

```
##           expr          mirna
## 2961      CAST hsa-miR-222-3p
## 4533    CORO1A hsa-miR-222-3p
## 4535    CORO1A hsa-miR-221-3p
## 7198    FAM46A hsa-miR-34a-5p
## 7199    FAM46A  hsa-miR-21-5p
## 8004     GANAB hsa-miR-222-3p
## 8175      GDI2 hsa-miR-222-3p
## 8300      GLO1 hsa-miR-222-3p
## 8663     GRB10 hsa-miR-222-3p
## 9307  HIST3H2A hsa-miR-34a-5p
## 9864     ICAM1 hsa-miR-221-3p
## 9870     ICAM1 hsa-miR-222-3p
## 11406 LGALS3BP hsa-miR-222-3p
## 11423   LHFPL2 hsa-miR-221-3p
## 11790      LYN hsa-miR-222-3p
## 11841     MAFG hsa-miR-218-5p
## 11894    MAP1B hsa-miR-222-3p
## 12570   MGAT4B hsa-miR-222-3p
## 13368     NAIP hsa-miR-221-3p
## 14202    NSUN5 hsa-miR-222-3p
## 15529     PKP2 hsa-miR-222-3p
## 17947    RNF10 hsa-miR-222-3p
## 20848   STAT5A hsa-miR-222-3p
## 21240   TATDN2 hsa-miR-222-3p
## 21980    TMEM2  hsa-miR-21-5p
## 22130    TNIP1 hsa-miR-221-3p
## 22530   TRIM44 hsa-miR-222-3p
```

```
# 27 pairs

#venn diagram
m1 <- list(rna=sig_expr_cNPC, rna_fisher=unique(sig_expr_mirna_cNPC$expr))

vennPlot1 <- venn.diagram(m1, NULL, fill=c('red','blue'), 
                         cat.fontface=4, lty=2, cex=1.2, cat.cex=1.2, main='Significant genes identified with individual (rna) or integration (rna_fisher)\n omics analysis: NPC')
grid.newpage()
grid.draw(vennPlot1)
```

```
m2 <- list(mirna=sig_mirna_cNPC, mirna_fisher=unique(sig_expr_mirna_cNPC$mirna))

vennPlot2 <- venn.diagram(m2, NULL, fill=c('green','blue'), 
                         cat.fontface=4, lty=2, cex=1.2, cat.cex=1.2, main='Significant miRNA identified with individual (mirna) or integration (mirna_fisher)\n omics analysis: NPC')
grid.newpage()
grid.draw(vennPlot2)
```

When we perform the combination analysis we identify 23 new genes related with GBM survival and 1 miRNAs.

Here we can see part of the list of new genes identified based on Fisher combination.

```
setdiff(sig_expr_mirna_cNPC$expr,sig_expr_cNPC)
```

```
##  [1] "CAST"     "CORO1A"   "GANAB"    "GDI2"     "GLO1"     "GRB10"   
##  [7] "HIST3H2A" "ICAM1"    "LGALS3BP" "LHFPL2"   "LYN"      "MAFG"    
## [13] "MAP1B"    "MGAT4B"   "NAIP"     "NSUN5"    "PKP2"     "RNF10"   
## [19] "STAT5A"   "TATDN2"   "TMEM2"    "TNIP1"    "TRIM44"
```

And the 4 new miRNA identified based on Fisher combination.

```
setdiff(sig_expr_mirna_cNPC$mirna,sig_mirna_cNPC)
```

```
## [1] "hsa-miR-221-3p" "hsa-miR-34a-5p" "hsa-miR-21-5p"  "hsa-miR-218-5p"
```

###### 2.7.1.3 All samples

```
#restrict the datasets to the elements of the data mapping
exprTMP <- rna_gbm_corrected[unique(na.omit(dataMappingExprMirna$expr)),]; #7,814 x 523

mirnaTMP <- new_mirna_matrix[unique(na.omit(dataMappingExprMirna$mirna)),]; #323 x 518

#specifying the data types.
dataTypesExprMirna <-  list(ttCoxphContinuous,ttCoxphContinuous)

#preparing the datasets
exprTMP <- createOmicsExpressionSet(Data = as.matrix(exprTMP), 
                                    pData = clinical_gbm[colnames(rna_gbm_corrected),clinicalVariables[['expr']]])
names(pData(exprTMP)) <- c("age","outcome")

mirnaTMP <- createOmicsExpressionSet(Data = as.matrix(mirnaTMP), 
                                     pData = clinical_gbm[colnames(mirna_gbm_corrected),clinicalVariables[['mirna']]])
names(pData(mirnaTMP)) <- c("age","outcome")

dataInputExprMirna <- list(expr = exprTMP, mirna = mirnaTMP)
```

###### 2.7.1.3.1 Parametric Combination

```
set.seed(12345)
omicsPCRes_aPC <- omicsPC(dataInput = dataInputExprMirna,
                       dataMapping = dataMappingExprMirna,
                       dataTypes = dataTypesExprMirna,
                       verbose = TRUE)
```

The output of `omicsPC` provides information regarding p-value and adjusted p-value from both individual and combined analysis.

```
res <- omicsPCRes_aPC

#Nominal p-values plot individual omics
data_plot <- data.frame(pval=c(res$expr[,"Pr(>|z|)"],res$mirna[,"Pr(>|z|)"]), 
                        group=c(rep(names(res)[1], nrow(res$expr)), 
                                rep(names(res)[2], nrow(res$mirna))))

p1 <- ggplot(data_plot, aes(x=group, y=pval, fill=group)) +
  geom_violin(trim=FALSE)+
  labs(title='Plot of individual omics p-value from all samples')

#FDR plot individual omics
data_plot <- data.frame(pval=c(res$expr[,"adj.P.Val"],res$mirna[,"adj.P.Val"]), 
                        group=c(rep(names(res)[1], nrow(res$expr)),
                                rep(names(res)[2], nrow(res$mirna))))

p2 <- ggplot(data_plot, aes(x=group, y=pval, fill=group)) +
  geom_violin(trim=FALSE)+
  labs(title='Plot of individual omics FDR from all samples')


#Nominal p-value plot PC
data_plot <- data.frame(pval=unlist(res$pvaluesPC[,-c(1,2)]), 
                        combination_method=rep(colnames(res$pvaluesPC[,-c(1,2)]),
                                               each=nrow(res$pvaluesPC)))

p3 <- ggplot(data_plot, aes(x=combination_method, y=pval, 
                            fill=combination_method)) +
  geom_violin(trim=FALSE)+
  labs(title='Plot of p-values from omicsPC (all samples)')


#FDR plot PC
data_plot <- data.frame(pval=unlist(res$qvaluesPC[,-c(1,2)]), 
                        combination_method=rep(colnames(res$qvaluesPC[,-c(1,2)]),
                                               each=nrow(res$qvaluesPC)))

p4 <- ggplot(data_plot, aes(x=combination_method, y=pval, 
                            fill=combination_method)) +
  geom_violin(trim=FALSE)+
  labs(title='Plot of FDR from omicsPC (all samples)')


grid.arrange(p1, p2, p3, p4, ncol=2, nrow=2)
```

Now, based on FDR results we can check if there are an increase of statistical power when both omics are jointly analyzed.

Note: here we consider significant an FDR<0.05.

```
#Significant genes or miRNA from individual omics (FDR<0.05)

#significant genes
(sig_expr_aPC <- rownames(res$expr)[which(res$expr[,"adj.P.Val"]<0.05)])
```

```
## [1] "FAM46A" "FUT4"   "FZD7"   "SNX10"
```

```
# 4 genes

#significant miRNA
(sig_mirna_aPC <- rownames(res$mirna)[which(res$mirna[,"adj.P.Val"]<0.05)])
```

```
## [1] "hsa-miR-222-3p"
```

```
# 1 miRNA

#Significant genes from mRNA + miRNA parametric combination (Fisher, FDR<0.05)
sig_expr_mirna_aPC <- res$qvaluesPC[which(res$qvaluesPC[,'Fisher']<0.05),1:2]
# 466 pairs

#venn diagram
m1 <- list(rna=sig_expr_aPC, rna_fisher=unique(sig_expr_mirna_aPC$expr))

vennPlot1 <- venn.diagram(m1, NULL, fill=c('red','blue'), 
                         cat.fontface=4, lty=2, cex=1.2, cat.cex=1.2, main='Significant genes identified with individual (rna) or integration (rna_fisher)\n omics analysis: PC (all samples)')
grid.newpage()
grid.draw(vennPlot1)
```

```
m2 <- list(mirna=sig_mirna_aPC, mirna_fisher=unique(sig_expr_mirna_aPC$mirna))

vennPlot2 <- venn.diagram(m2, NULL, fill=c('green','blue'), 
                         cat.fontface=4, lty=2, cex=1.2, cat.cex=1.2, main='Significant miRNA identified with individual (mirna) or integration (mirna_fisher)\n omics analysis: PC (all samples)')
grid.newpage()
grid.draw(vennPlot2)
```

When we perform the combination analysis we identify 382 new genes related with GBM survival and 54 miRNAs.

Here we can see part of the list of new genes (n=382) identified based on Fisher combination.

```
setdiff(sig_expr_mirna_aPC$expr,sig_expr_aPC)[1:50]
```

```
##  [1] "ABHD3"    "ACACA"    "ACSL3"    "ACTB"     "ACTG1"    "AHSA1"   
##  [7] "ALG3"     "AMELX"    "AMOT"     "ANXA2"    "AP1B1"    "AP3B1"   
## [13] "APLP2"    "APOBEC3C" "APOL2"    "APP"      "ARL4C"    "ATF5"    
## [19] "ATP11B"   "ATP1A1"   "ATP2A2"   "ATP7B"    "AUTS2"    "BAG3"    
## [25] "BBC3"     "BCL2L11"  "BICD1"    "BPTF"     "C11orf57" "C8orf33" 
## [31] "CABYR"    "CANT1"    "CANX"     "CAPG"     "CAPRIN1"  "CASC3"   
## [37] "CASKIN2"  "CASP3"    "CAST"     "CBX2"     "CCDC47"   "CCR1"    
## [43] "CCT3"     "CD44"     "CDC25C"   "CDKL5"    "CDKN1B"   "CDKN1C"  
## [49] "CEBPB"    "CENPT"
```

And the new miRNAs (n=54) identified based on Fisher combination.

```
setdiff(sig_expr_mirna_aPC$mirna,sig_mirna_aPC)
```

```
##  [1] "hsa-miR-221-3p"  "hsa-miR-148a-3p" "hsa-miR-155-5p"  "hsa-miR-132-3p" 
##  [5] "hsa-miR-7-5p"    "hsa-miR-92a-3p"  "hsa-miR-26a-5p"  "hsa-miR-192-5p" 
##  [9] "hsa-miR-18a-3p"  "hsa-miR-34a-5p"  "hsa-miR-21-5p"   "hsa-miR-31-5p"  
## [13] "hsa-miR-124-3p"  "hsa-miR-335-5p"  "hsa-miR-181a-5p" "hsa-miR-27a-3p" 
## [17] "hsa-miR-27b-3p"  "hsa-miR-346"     "hsa-miR-128-3p"  "hsa-miR-204-5p" 
## [21] "hsa-miR-20b-5p"  "hsa-miR-218-5p"  "hsa-miR-215-5p"  "hsa-miR-484"    
## [25] "hsa-miR-9-5p"    "hsa-miR-19b-3p"  "hsa-miR-331-3p"  "hsa-miR-16-5p"  
## [29] "hsa-miR-143-3p"  "hsa-miR-26b-5p"  "hsa-miR-378a-3p" "hsa-miR-25-3p"  
## [33] "hsa-miR-98-5p"   "hsa-miR-30a-5p"  "hsa-miR-129-5p"  "hsa-miR-1"      
## [37] "hsa-miR-199a-3p" "hsa-miR-145-5p"  "hsa-miR-200b-3p" "hsa-miR-122-5p" 
## [41] "hsa-miR-126-3p"  "hsa-miR-671-5p"  "hsa-miR-596"     "hsa-miR-93-5p"  
## [45] "hsa-let-7e-5p"   "hsa-let-7a-5p"   "hsa-miR-32-5p"   "hsa-miR-375"    
## [49] "hsa-miR-203a"    "hsa-miR-769-3p"  "hsa-miR-196a-5p" "hsa-miR-299-5p" 
## [53] "hsa-miR-146a-5p" "hsa-miR-197-3p"
```

###### 2.7.1.3.2 Non Parametric Combination

```
set.seed(12345)

# Setting methods to combine pvalues
combMethods <- c("Fisher", "Liptak", "Tippett")
# Setting number of permutations
numPerms <- 1000
# Setting number of cores
numCores <- 4
# Setting omicsNPC to print out the steps that it performs.
verbose <- TRUE

omicsNPCRes_aNPC_rna_mirna <- omicsNPC(dataInput = dataInputExprMirna,
                         dataMapping = dataMappingExprMirna,
                         dataTypes = dataTypesExprMirna,
                         combMethods = combMethods, 
                         numPerms = numPerms,
                         numCores = numCores,
                         verbose = verbose)
```

The output of `omicsNPC` provides information regarding p-value and adjusted p-value from both individual and combined analysis.

```
res <- omicsNPCRes_aNPC_rna_mirna

#Nominal p-values plot individual omics
data_plot <- data.frame(pval=c(res$expr[,"Pr(>|z|)"],res$mirna[,"Pr(>|z|)"]), 
                        group=c(rep(names(res)[1], nrow(res$expr)), 
                                rep(names(res)[2], nrow(res$mirna))))

p1 <- ggplot(data_plot, aes(x=group, y=pval, fill=group)) +
  geom_violin(trim=FALSE)+
  labs(title='Plot of individual omics p-value from overlapping samples')

#FDR plot individual omics
data_plot <- data.frame(pval=c(res$expr[,"adj.P.Val"],res$mirna[,"adj.P.Val"]), 
                        group=c(rep(names(res)[1], nrow(res$expr)),
                                rep(names(res)[2], nrow(res$mirna))))

p2 <- ggplot(data_plot, aes(x=group, y=pval, fill=group)) +
  geom_violin(trim=FALSE)+
  labs(title='Plot of individual omics FDR from overlapping samples')


#Nominal p-value plot NPC
data_plot <- data.frame(pval=unlist(res$pvaluesNPC[,-c(1:4)]), 
                        combination_method=rep(colnames(res$pvaluesNPC[,-c(1:4)])
                                               , each=nrow(res$pvaluesNPC)))

p3 <- ggplot(data_plot, aes(x=combination_method, y=pval, 
                            fill=combination_method)) +
  geom_violin(trim=FALSE)+
  labs(title='Plot of p-values in overlapping omicsNPC')


#FDR plot PC
data_plot <- data.frame(pval=unlist(res$qvaluesNPC[,-c(1:4)]), 
                        combination_method=rep(colnames(res$qvaluesNPC[,-c(1:4)])
                                               , each=nrow(res$qvaluesNPC)))

p4 <- ggplot(data_plot, aes(x=combination_method, y=pval, fill=combination_method)) +
  geom_violin(trim=FALSE)+
  labs(title='Plot of FDR in overlapping omicsNPC')


grid.arrange(p1, p2, p3, p4, ncol=2, nrow=2)
```

Now, based on FDR results we can check if there are an increase of statistical power when both omics are jointly analyzed.

Note: here we consider significant an FDR<0.05.

```
#Significant genes or miRNA from individual omics (FDR<0.05)

#significant genes
(sig_expr_aNPC <- rownames(res$expr)[which(res$expr[,"adj.P.Val"]<0.05)])
```

```
## [1] "FAM46A" "FUT4"   "FZD7"   "SNX10"
```

```
# 4 genes

#significant miRNA
(sig_mirna_aNPC <- rownames(res$mirna)[which(res$mirna[,"adj.P.Val"]<0.05)])
```

```
## [1] "hsa-miR-222-3p"
```

```
# 1 miRNA

#Significant genes from mRNA + miRNA non parametric combination (Fisher, FDR<0.05)
sig_expr_mirna_aNPC <- res$qvaluesNPC[which(res$qvaluesNPC[,'Fisher']<0.05),1:2]
# 50 pairs

#venn diagram
m1 <- list(rna=sig_expr_aNPC, rna_fisher=unique(sig_expr_mirna_aNPC$expr))

vennPlot1 <- venn.diagram(m1, NULL, fill=c('red','blue'), 
                         cat.fontface=4, lty=2, cex=1.2, cat.cex=1.2, main='Significant genes identified with individual (rna) or integration (rna_fisher)\n omics analysis: NPC (all samples)')
grid.newpage()
grid.draw(vennPlot1)
```

```
m2 <- list(mirna=sig_mirna_aNPC, mirna_fisher=unique(sig_expr_mirna_aNPC$mirna))

vennPlot2 <- venn.diagram(m2, NULL, fill=c('green','blue'), 
                         cat.fontface=4, lty=2, cex=1.2, cat.cex=1.2, main='Significant miRNA identified with individual (mirna) or integration (mirna_fisher)\n omics analysis: NPC (all samples)')
grid.newpage()
grid.draw(vennPlot2)
```

When we perform the combination analysis we identify 43 new genes related with GBM survival and 7 miRNAs.

Here we can see the list of new genes (n=43) identified based on Fisher combination.

```
setdiff(sig_expr_mirna_aNPC$expr,sig_expr_aNPC)
```

```
##  [1] "ATF5"     "CAST"     "CLIC1"    "CORO1A"   "CUTA"     "EFNB2"   
##  [7] "ENO1"     "EXOC3"    "FNDC3B"   "GANAB"    "GATA4"    "GDI2"    
## [13] "GLO1"     "GRB10"    "HIST3H2A" "ICAM1"    "LGALS3BP" "LHFPL2"  
## [19] "LRRFIP1"  "LYN"      "MAP1B"    "MET"      "MGAT4B"   "MYD88"   
## [25] "MYO1C"    "NAIP"     "NSUN5"    "OLIG2"    "OSBPL10"  "PKP2"    
## [31] "PRDM16"   "RNF10"    "SAP30L"   "SH3BGRL3" "STAT5A"   "TATDN2"  
## [37] "TBK1"     "TMEM2"    "TNFSF10"  "TNIP1"    "TRIM44"   "TUB"     
## [43] "ZIC3"
```

And the new miRNAs (n=7) identified based on Fisher combination.

```
setdiff(sig_expr_mirna_aNPC$mirna,sig_mirna_aNPC)
```

```
## [1] "hsa-miR-221-3p"  "hsa-miR-155-5p"  "hsa-miR-218-5p"  "hsa-miR-34a-5p" 
## [5] "hsa-miR-21-5p"   "hsa-miR-200b-3p" "hsa-miR-31-5p"
```

###### 2.7.1.4 Integration strategies comparison

Finally, we can compare the divergences between parametric and non-parametric approaches and between the overlapping or whole cohort escenarios.

```
#Significant pairs from different approaches

#parmetric combination with overlapping samples (n=397)
oPC <- paste(sig_expr_mirna_cPC$expr,sig_expr_mirna_cPC$mirna, sep=":")
#parmetric combination with all samples (n=466)
aPC <- paste(sig_expr_mirna_aPC$expr,sig_expr_mirna_aPC$mirna, sep=":")
#non parmetric combination with overlapping samples (n=27)
oNPC <- paste(sig_expr_mirna_cNPC$expr,sig_expr_mirna_cNPC$mirna, sep=":")
#non parmetric combination with all samples (n=50)
aNPC <- paste(sig_expr_mirna_aNPC$expr,sig_expr_mirna_aNPC$mirna, sep=":")

res_all <- list(oPC=oPC, aPC=aPC, oNPC=oNPC, aNPC=aNPC)

vennPlot_m1 <- venn.diagram(res_all, NULL, fill=c("green","blue","orange","pink"),
                            cat.fontface=4, lty=2, cex=1.2, cat.cex=1.2,
                            main="Significant pairs for the different escenarios \n FDR (cut-off=0.05)")

grid.newpage()
grid.draw(vennPlot_m1)
```

If we look in the significant pairs identified in each case (FDR<0.05), in total there are 472 pairs identified by the different approaches; being 5.5% (26/472) common between different methods.
The use of whole data allows the identification of a largest number of signifcant pairs in both parametric and non parametric approach. Importantly, the main differences between identified pairs are found between parametric and non parametric approaches. PC approach identifies a large number of significant pairs in comparison to NPC approach.

##### 2.7.2 mRNA + Methylation

First of all we need to have a mapping file mathcing omics.

###### 2.7.2.1 Mapping file

```
#Load the mapping info
#load('methy2GeneMap.RData')
methy2GeneMap <- methy2GeneMap[-40902, ]; #repetition

#restricting mapping info to methylation info
methy2GeneMap_1 <- methy2GeneMap[methy2GeneMap[,1]%in%rownames(logit_met_gbm_corrected),] #from the initial 311,135 methyl features we get 57,645 in common

head(methy2GeneMap_1)
```

```
##          name   Gene
## 15 cg02725885 PPAP2C
## 16 cg09908879 PPAP2C
## 17 cg16691033 PPAP2C
## 18 cg02935627 PPAP2C
## 19 cg12272837 PPAP2C
## 20 cg25697727 PPAP2C
```

```
#creating the mRNA association tables has been previously created
head(expr2GeneMap)
```

```
##   measurement    Gene
## 1       FSTL1   FSTL1
## 2        AACS    AACS
## 3       RPS11   RPS11
## 4     CREB3L1 CREB3L1
## 5       ELMO2   ELMO2
## 6       PNMA1   PNMA1
```

```
#creating the data mapping (57,645 x 3)
dataMappingExprMet <- combiningMappings(mappings = list(expr = expr2GeneMap, 
                                                        met = methy2GeneMap_1), 
                                          retainAll = TRUE, reference = 'Gene'); 

head(dataMappingExprMet)
```

```
##     expr        met   Gene
## 2    A2M cg00134295    A2M
## 3 A4GALT cg01268859 A4GALT
## 4 A4GALT cg07393322 A4GALT
## 5 A4GALT cg26705472 A4GALT
## 6 A4GALT cg20706071 A4GALT
## 7  A4GNT cg23669440  A4GNT
```

We can see that based on our mRNA and methylation analysis, we have 57,645 pairs in total, it corresponds to 9,620 unique genes (80% from the whole mRNA dataset) and 57,645 unique methylation sites (18% from teh whole methylation dataset).

Then, we define the relevant clinical variables in each escenario. The variable of interest in the case of GBM is the `survivalOtucome`. Moreover, Moreover, `days_to_birth` is included as a confounder.

```
clinicalVariables <- list(expr = c('days_to_birth', 'survivalOutcome'), 
                          met = c('days_to_birth', 
                                  'survivalOutcome'),
                          mirna = c('days_to_birth', 'survivalOutcome'));
```

###### 2.7.2.2 Overlaping samples

```
#restrict the datasets to the elements of the data mapping
exprTMP <- rna_gbm_corrected[unique(na.omit(dataMappingExprMet$expr)),
                             ids_rna_met]
#Dimension: 9,620 x 83

metTMP <-logit_met_gbm_corrected[unique(na.omit(dataMappingExprMet$met)),
                                 ids_rna_met]
#Dimension: 57,645 x 83

#specifying the data types.
dataTypesExprMet <- list(ttCoxphContinuous,ttCoxphContinuous)

#preparing the datasets
exprTMP <- createOmicsExpressionSet(Data = as.matrix(exprTMP), 
                                    pData = clinical_gbm[ids_rna_met,
                                                         clinicalVariables[['expr']]])
names(pData(exprTMP)) <- c("age","outcome")

metTMP <- createOmicsExpressionSet(Data = as.matrix(metTMP), 
                                     pData = clinical_gbm[ids_rna_met,
                                                          clinicalVariables[['met']]])
names(pData(metTMP)) <- c("age","outcome")

dataInputExprMet <- list(expr = exprTMP, met = metTMP)
```

###### 2.7.2.2.1 Parametric Combination

```
set.seed(12345)
omicsPCRes_cPC <- omicsPC(dataInput = dataInputExprMet,
                       dataMapping = dataMappingExprMet,
                       dataTypes = dataTypesExprMet,
                       verbose = TRUE)
```

The output of `omicsPC` provides information regarding p-value and adjusted p-value from both individual and combined analysis.

```
res <- omicsPCRes_cPC

#Nominal p-values plot individual omics
data_plot <- data.frame(pval=c(res$expr[,"Pr(>|z|)"],res$met[,"Pr(>|z|)"]), 
                        group=c(rep(names(res)[1], nrow(res$expr)), 
                                rep(names(res)[2], nrow(res$met))))

p1 <- ggplot(data_plot, aes(x=group, y=pval, fill=group)) +
  geom_violin(trim=FALSE)+
  labs(title='Plot of individual omics p-value from overlapping samples')

#FDR plot individual omics
data_plot <- data.frame(pval=c(res$expr[,"adj.P.Val"],res$met[,"adj.P.Val"]), 
                        group=c(rep(names(res)[1], nrow(res$expr)),
                                rep(names(res)[2], nrow(res$met))))

p2 <- ggplot(data_plot, aes(x=group, y=pval, fill=group)) +
  geom_violin(trim=FALSE)+
  labs(title='Plot of individual omics FDR from overlapping samples')


#Nominal p-value plot PC
data_plot <- data.frame(pval=unlist(res$pvaluesPC[,-c(1,2)]), 
                        combination_method=rep(colnames(res$pvaluesPC[,-c(1,2)]),
                                               each=nrow(res$pvaluesPC)))

p3 <- ggplot(data_plot, aes(x=combination_method, y=pval, 
                            fill=combination_method)) + 
  geom_violin(trim=FALSE)+
  labs(title='Plot of p-values in overlapping omicsPC')


#FDR plot PC
data_plot <- data.frame(pval=unlist(res$qvaluesPC[,-c(1,2)]), 
                        combination_method=rep(colnames(res$qvaluesPC[,-c(1,2)]),
                                               each=nrow(res$qvaluesPC)))

p4 <- ggplot(data_plot, aes(x=combination_method, y=pval,
                            fill=combination_method)) +
  geom_violin(trim=FALSE)+
  labs(title='Plot of FDR in overlapping omicsPC')


grid.arrange(p1, p2, p3, p4, ncol=2, nrow=2)
```

Now, based on FDR results we can check if there are an increase of statistical power when both omics are jointly analyzed.

Note: here we consider significant an FDR<0.05.

```
#significant genes
(sig_expr_cPC <- rownames(res$expr)[which(res$expr[,"adj.P.Val"]<0.05)])
```

```
## [1] "CCDC41" "IGFBP6"
```

```
# 2 genes

#significant methyl
(sig_met_cPC <- rownames(res$met)[which(res$met[,"adj.P.Val"]<0.05)])
```

```
## [1] "cg24113409"
```

```
# 1 methylation site

#Significant genes from mRNA + miRNA non parametric combination (Fisher, FDR<0.05)
sig_expr_met_cPC <- res$qvaluesPC[which(res$qvaluesPC[,'Fisher']<0.05),1:2]
#323 pairs

#venn diagram
m1 <- list(rna=sig_expr_cPC, rna_fisher=unique(sig_expr_met_cPC$expr))

vennPlot1 <- venn.diagram(m1, NULL, fill=c('red','blue'), 
                         cat.fontface=4, lty=2, cex=1.2, cat.cex=1.2, 
                         main='Significant genes identified with individual (rna) or integration (rna_fisher)\n omics analysis: PC')
grid.newpage()
grid.draw(vennPlot1)
```

```
m2 <- list(met=sig_met_cPC, mirna_fisher=unique(sig_expr_met_cPC$met))

vennPlot2 <- venn.diagram(m2, NULL, fill=c('green','blue'), 
                         cat.fontface=4, lty=2, cex=1.2, cat.cex=1.2,
                         main='Significant methylation identified with individual (met) or integration (met_fisher)\n omics analysis: PC')
grid.newpage()
grid.draw(vennPlot2)
```

When we perform the combination analysis we identify 179 new genes related with GBM survival and 322 methylation sites.

Here we can see part of the new genes (n=179) identified based on Fisher combination.

```
setdiff(sig_expr_met_cPC$expr,sig_expr_cPC)[1:50]
```

```
##  [1] "ABCG1"     "ADAM22"    "ADCYAP1R1" "ADORA3"    "AIM1L"     "AKAP1"    
##  [7] "ALDH3B1"   "ALDH3B2"   "ALX1"      "AMPD3"     "ANGPTL4"   "ANK1"     
## [13] "ANXA11"    "APOBEC3C"  "ARPC1B"    "ARSJ"      "AVPI1"     "B3GAT3"   
## [19] "BANK1"     "BCAN"      "BCMO1"     "C1orf61"   "C20orf43"  "CAMTA2"   
## [25] "CCR1"      "CDCP1"     "CDKN1A"    "CDYL"      "CEACAM4"   "CHMP6"    
## [31] "CHN2"      "CLIP4"     "COL14A1"   "COL16A1"   "CPM"       "CREBBP"   
## [37] "CREM"      "CRISPLD2"  "CSNK1D"    "CTSB"      "CXCR4"     "CYB5R2"   
## [43] "DDR1"      "DENND2D"   "DLEU2"     "DSN1"      "DUS1L"     "DYNC2H1"  
## [49] "ECHS1"     "EGR2"
```

And part of the new methylation sites (n=322) identified based on Fisher combination.

```
setdiff(sig_expr_met_cPC$met,sig_met_cPC)[1:50]
```

```
##  [1] "cg01289965" "cg27220091" "cg27076139" "cg17617930" "cg05855208"
##  [6] "cg26915889" "cg12813724" "cg10667386" "cg05620821" "cg19933414"
## [11] "cg09982224" "cg15322932" "cg14871333" "cg24390938" "cg18229767"
## [16] "cg02494246" "cg20333014" "cg14996220" "cg12069276" "cg19132462"
## [21] "cg05647503" "cg18010131" "cg25968076" "cg07268332" "cg10811474"
## [26] "cg19585816" "cg04153489" "cg11647108" "cg06523744" "cg17140191"
## [31] "cg07634959" "cg12724221" "cg25217772" "cg05935311" "cg12664038"
## [36] "cg25217710" "cg19321979" "cg26781524" "cg26827725" "cg01518279"
## [41] "cg02232839" "cg14177914" "cg10945917" "cg26339180" "cg09578319"
## [46] "cg12509435" "cg05135942" "cg12762432" "cg17644311" "cg07118243"
```

###### 2.7.2.2.2 Non Parametric Combination

```
set.seed(12345)

# Setting methods to combine pvalues
combMethods <- c("Fisher", "Liptak", "Tippett")
# Setting number of permutations
numPerms <- 1000
# Setting number of cores
numCores <- 1
# Setting omicsNPC to print out the steps that it performs.
verbose <- TRUE

omicsNPCRes_cNPC <- omicsNPC(dataInput = dataInputExprMet,
                         dataMapping = dataMappingExprMet,
                         dataTypes = dataTypesExprMet,
                         combMethods = combMethods, 
                         numPerms = numPerms,
                         numCores = numCores,
                         verbose = verbose)
```

The output of `omicsNPC` provides information regarding p-value and adjusted p-value from both individual and combined analysis.

```
res <- omicsNPCRes_cNPC_rna_met

#Nominal p-values plot individual omics
data_plot <- data.frame(pval=c(res$expr[,"Pr(>|z|)"],res$met[,"Pr(>|z|)"]), 
                        group=c(rep(names(res)[1], nrow(res$expr)), 
                                rep(names(res)[2], nrow(res$met))))

p1 <- ggplot(data_plot, aes(x=group, y=pval, fill=group)) +
  geom_violin(trim=FALSE)+
  labs(title='Plot of individual omics p-value from overlapping samples')

#FDR plot individual omics
data_plot <- data.frame(pval=c(res$expr[,"adj.P.Val"],res$met[,"adj.P.Val"]),
                        group=c(rep(names(res)[1], nrow(res$expr)),
                                rep(names(res)[2], nrow(res$met))))

p2 <- ggplot(data_plot, aes(x=group, y=pval, fill=group)) +
  geom_violin(trim=FALSE)+
  labs(title='Plot of individual omics FDR from overlapping samples')


#Nominal p-value plot NPC
data_plot <- data.frame(pval=unlist(res$pvaluesNPC[,-c(1:4)]), 
                        combination_method=rep(colnames(res$pvaluesNPC[,-c(1:4)])
                                               , each=nrow(res$pvaluesNPC)))

p3 <- ggplot(data_plot, aes(x=combination_method, y=pval,
                            fill=combination_method)) +
  geom_violin(trim=FALSE)+
  labs(title='Plot of p-values in overlapping omicsNPC')


#FDR plot NPC
data_plot <- data.frame(pval=unlist(res$qvaluesNPC[,-c(1:4)]), 
                        combination_method=rep(colnames(res$qvaluesNPC[,-c(1:4)])
                                               , each=nrow(res$qvaluesNPC)))

p4 <- ggplot(data_plot, aes(x=combination_method, y=pval,
                            fill=combination_method)) +
  geom_violin(trim=FALSE)+
  labs(title='Plot of FDR in overlapping omicsNPC')


grid.arrange(p1, p2, p3, p4, ncol=2, nrow=2)
```

Now, based on FDR results we can check if there are an increase of statistical power when both omics are jointly analyzed.

Note: here we consider significant an FDR<0.05.

```
#Significant genes or miRNA from individual omics (FDR<0.05)

#significant genes
(sig_expr_cNPC <- rownames(res$expr)[which(res$expr[,"adj.P.Val"]<0.05)])
```

```
## [1] "CCDC41" "IGFBP6"
```

```
# 2 genes

#significant met
(sig_met_cNPC <- rownames(res$met)[which(res$met[,"adj.P.Val"]<0.05)])
```

```
## [1] "cg24113409"
```

```
# 1 methylation site

#Significant genes from mRNA + methylation non parametric combination (Fisher, FDR<0.05)
sig_expr_met_cNPC <- res$qvaluesNPC[which(res$qvaluesNPC[,'Fisher']<0.05),1:2]
# 150 pairs

#venn diagram
m1 <- list(rna=sig_expr_cNPC, rna_fisher=unique(sig_expr_met_cNPC$expr))

vennPlot1 <- venn.diagram(m1, NULL, fill=c('red','blue'), 
                         cat.fontface=4, lty=2, cex=1.2, cat.cex=1.2, 
                         main='Significant genes identified with individual (rna) or integration (rna_fisher)\n omics analysis: PC')
grid.newpage()
grid.draw(vennPlot1)
```

```
m2 <- list(met=sig_met_cNPC, met_fisher=unique(sig_expr_met_cNPC$met))

vennPlot2 <- venn.diagram(m2, NULL, fill=c('green','blue'), 
                         cat.fontface=4, lty=2, cex=1.2, cat.cex=1.2,
                         main='Significant methylation identified with individual (met) or integration (met_fisher)\n omics analysis: PC')
grid.newpage()
grid.draw(vennPlot2)
```

When we perform the combination analysis we identify 106 new genes related with GBM survival and 150 methylation sites.

Here we can see part of the new genes (n=106) identified based on Fisher combination.

```
setdiff(sig_expr_met_cNPC$expr,sig_expr_cNPC)[1:50]
```

```
##  [1] "ABCG1"    "ADAM22"   "ADORA3"   "ALDH3B2"  "AMPD3"    "ANXA11"  
##  [7] "APOBEC3C" "ARHGEF4"  "ARID5A"   "ARPC1B"   "AVPI1"    "BCAN"    
## [13] "BCMO1"    "CCR1"     "CDCP1"    "CDKN1A"   "CDYL"     "CEACAM4" 
## [19] "CHI3L1"   "CHMP6"    "CHST12"   "CPM"      "CYB5R2"   "DENND2D" 
## [25] "EDNRB"    "EGR2"     "ELL2"     "ENOSF1"   "EPB41L3"  "EXPH5"   
## [31] "FHL2"     "GAA"      "GCNT1"    "GPT"      "GRN"      "HCK"     
## [37] "HK1"      "HLA-DMA"  "HOXA3"    "HPCAL1"   "HRAS"     "HTATIP2" 
## [43] "ICAM1"    "IDH2"     "IFNGR2"   "IGFBP3"   "IL1RAP"   "IL4R"    
## [49] "IL6R"     "ILF3"
```

And part of the new methylation sites (n=150) identified based on Fisher combination.

```
setdiff(sig_expr_met_cNPC$met,sig_met_cNPC)[1:50]
```

```
##  [1] "cg01289965" "cg27220091" "cg26915889" "cg20333014" "cg12069276"
##  [6] "cg18010131" "cg25968076" "cg07268332" "cg11647108" "cg06523744"
## [11] "cg12910906" "cg16388829" "cg17140191" "cg12724221" "cg12664038"
## [16] "cg25217710" "cg19321979" "cg26781524" "cg11589536" "cg13722619"
## [21] "cg05389236" "cg24425727" "cg17526952" "cg24816460" "cg08249424"
## [26] "cg21529807" "cg05891548" "cg02097014" "cg00691948" "cg05483875"
## [31] "cg21211357" "cg21433768" "cg24575083" "cg16281276" "cg25722212"
## [36] "cg12847373" "cg20018723" "cg11571761" "cg20415170" "cg07100532"
## [41] "cg23564700" "cg16622495" "cg06550462" "cg12967001" "cg10022155"
## [46] "cg26344233" "cg01154930" "cg22289360" "cg26572973" "cg09957864"
```

###### 2.7.2.3 All samples

```
#restrict the datasets to the elements of the data mapping
exprTMP <- rna_gbm_corrected[unique(na.omit(dataMappingExprMet$expr)),]; 
#Dimension: 9,620 x 523

metTMP <- logit_met_gbm_corrected[unique(na.omit(dataMappingExprMet$met)),]; 
#Dimension: 57,645 x 95

#specifying the data types.
dataTypesExprMet <- list(ttCoxphContinuous,ttCoxphContinuous)

#preparing the datasets
exprTMP <- createOmicsExpressionSet(Data = as.matrix(exprTMP), 
                                    pData = clinical_gbm[colnames(rna_gbm_corrected),clinicalVariables[['expr']]])

names(pData(exprTMP)) <- c("age","outcome")

metTMP <- createOmicsExpressionSet(Data = as.matrix(metTMP), 
                                     pData = clinical_gbm[colnames(logit_met_gbm_corrected),clinicalVariables[['met']]])

names(pData(metTMP)) <- c("age","outcome")

dataInputExprMet <- list(expr = exprTMP, met = metTMP)
```

###### 2.7.2.3.1 Parametric Combination

```
set.seed(12345)
omicsPCRes_aPC <- omicsPC(dataInput = dataInputExprMet,
                       dataMapping = dataMappingExprMet,
                       dataTypes = dataTypesExprMet,
                       verbose = TRUE)
```

The output of `omicsPC` provides information regarding p-value and adjusted p-value from both individual and combined analysis.

```
res <- omicsPCRes_aPC

#Nominal p-values plot individual omics
data_plot <- data.frame(pval=c(res$expr[,"Pr(>|z|)"],res$met[,"Pr(>|z|)"]), 
                        group=c(rep(names(res)[1], nrow(res$expr)), 
                                rep(names(res)[2], nrow(res$met))))

p1 <- ggplot(data_plot, aes(x=group, y=pval, fill=group)) +
  geom_violin(trim=FALSE)+
  labs(title='Plot of individual omics p-value from all samples')

#FDR plot individual omics
data_plot <- data.frame(pval=c(res$expr[,"adj.P.Val"],res$met[,"adj.P.Val"]), 
                        group=c(rep(names(res)[1], nrow(res$expr)),
                                rep(names(res)[2], nrow(res$met))))

p2 <- ggplot(data_plot, aes(x=group, y=pval, fill=group)) +
  geom_violin(trim=FALSE)+
  labs(title='Plot of individual omics FDR from all samples')


#Nominal p-value plot PC
data_plot <- data.frame(pval=unlist(res$pvaluesPC[,-c(1,2)]), 
                        combination_method=rep(colnames(res$pvaluesPC[,-c(1,2)]),
                                               each=nrow(res$pvaluesPC)))

p3 <- ggplot(data_plot, aes(x=combination_method, y=pval, 
                            fill=combination_method)) +
  geom_violin(trim=FALSE)+
  labs(title='Plot of p-values from omicsPC (all samples)')


#FDR plot PC
data_plot <- data.frame(pval=unlist(res$qvaluesPC[,-c(1,2)]), 
                        combination_method=rep(colnames(res$qvaluesPC[,-c(1,2)]),
                                               each=nrow(res$qvaluesPC)))

p4 <- ggplot(data_plot, aes(x=combination_method, y=pval, 
                            fill=combination_method)) +
  geom_violin(trim=FALSE)+
  labs(title='Plot of FDR from omicsPC (all samples)')


grid.arrange(p1, p2, p3, p4, ncol=2, nrow=2)
```

Now, based on FDR results we can check if there are an increase of statistical power when both omics are jointly analyzed.

Note: here we consider significant an FDR<0.05.

```
#Significant genes or methylation from individual omics (FDR<0.05)

#significant genes
(sig_expr_aPC <- rownames(res$expr)[which(res$expr[,"adj.P.Val"]<0.05)])
```

```
## [1] "FAM46A"  "FNDC3B"  "FUT4"    "FZD7"    "PRM2"    "SIGLEC9" "SNX10"
```

```
# 7 genes

#significant methylation sites
(sig_met_aPC <- rownames(res$met)[which(res$met[,"adj.P.Val"]<0.05)])
```

```
## character(0)
```

```
# 0 methylation sites

#Significant genes from mRNA + methylation non parametric combination (Fisher, FDR<0.05)
sig_expr_met_aPC <- res$qvaluesPC[which(res$qvaluesPC[,'Fisher']<0.05),1:2]
# 207 pairs

#venn diagram
m1 <- list(rna=sig_expr_aPC, rna_fisher=unique(sig_expr_met_aPC$expr))

vennPlot1 <- venn.diagram(m1, NULL, fill=c('red','blue'), 
                         cat.fontface=4, lty=2, cex=1.2, cat.cex=1.2, main='Significant genes identified with individual (rna) or integration (rna_fisher)\n omics analysis: PC (all samples)')
grid.newpage()
grid.draw(vennPlot1)
```

```
m2 <- list(met=sig_met_aPC, met_fisher=unique(sig_expr_met_aPC$met))

vennPlot2 <- venn.diagram(m2, NULL, fill=c('green','blue'), 
                         cat.fontface=4, lty=2, cex=1.2, cat.cex=1.2, main='Significant methylation sites identified with individual (met) or integration (met_fisher)\n omics analysis: PC (all samples)')
grid.newpage()
grid.draw(vennPlot2)
```

When we perform the combination analysis we identify 89 new genes related with GBM survival and 207 methylation sites.

Here we can see part of new genes (n=89) identified based on Fisher combination.

```
setdiff(sig_expr_met_aPC$expr,sig_expr_aPC)
```

```
##  [1] "ALX1"     "AMPD3"    "ANXA11"   "APOBEC3C" "ARHGAP25" "ARPC1B"  
##  [7] "ASCC1"    "ASCL2"    "ATP1A1"   "BICD1"    "BMP4"     "BRF1"    
## [13] "BST2"     "BTNL2"    "C13orf15" "C2"       "C6orf48"  "CCR1"    
## [19] "CHI3L1"   "CLIP4"    "CPD"      "CTSZ"     "CYB5R2"   "DENND2D" 
## [25] "DGKZ"     "DKK3"     "DUS1L"    "DUSP5"    "EPB41L3"  "GCM2"    
## [31] "GJA3"     "GRB10"    "HEXB"     "HIST3H2A" "HIVEP3"   "HOMER1"  
## [37] "HPCAL1"   "HRAS"     "HS3ST3A1" "ICAM1"    "IFNGR2"   "IGFBP2"  
## [43] "IGFBP3"   "IL17RA"   "LGALS3"   "LGALS3BP" "LGALS8"   "LHFPL2"  
## [49] "LITAF"    "LRRK1"    "LZTS1"    "MKNK1"    "MLNR"     "MYD88"   
## [55] "NCKAP1L"  "NDRG2"    "NFE2L3"   "NKX6-1"   "NUAK1"    "OSMR"    
## [61] "PARVB"    "PAX9"     "PCSK1"    "PDGFA"    "PHF11"    "PLEKHG3" 
## [67] "PLK2"     "PRKAG2"   "RAP2B"    "RGS16"    "ROBO4"    "SH2B3"   
## [73] "SHANK1"   "SLC16A3"  "SLC20A1"  "SLC43A3"  "SOX1"     "SOX21"   
## [79] "SREBF1"   "SV2B"     "TBX4"     "TCIRG1"   "TLR2"     "TLX3"    
## [85] "TMEM2"    "TRPC6"    "UGP2"     "UPK2"     "ZNF208"
```

And the new methylation sites (n=207) identified based on Fisher combination (first 20 shown).

```
setdiff(sig_expr_met_aPC$met,sig_met_aPC)[1:20]
```

```
##  [1] "cg14996220" "cg25800765" "cg19132462" "cg25968076" "cg11647108"
##  [6] "cg06523744" "cg15258980" "cg17140191" "cg19577016" "cg27252164"
## [11] "cg11342277" "cg15990192" "cg13079047" "cg21587861" "cg09229893"
## [16] "cg04269188" "cg16313343" "cg01254505" "cg17049418" "cg19281363"
```

###### 2.7.2.3.2 Non Parametric Combination

```
set.seed(12345)

# Setting methods to combine pvalues
combMethods <- c("Fisher", "Liptak", "Tippett")
# Setting number of permutations
numPerms <- 1000
# Setting number of cores
numCores <- 4
# Setting omicsNPC to print out the steps that it performs.
verbose <- TRUE

omicsNPCRes_aNPC_rna_met <- omicsNPC(dataInput = dataInputExprMet,
                         dataMapping = dataMappingExprMet,
                         dataTypes = dataTypesExprMet,
                         combMethods = combMethods, 
                         numPerms = numPerms,
                         numCores = numCores,
                         verbose = verbose)
```

The output of `omicsNPC` provides information regarding p-value and adjusted p-value from both individual and combined analysis.

```
res <- omicsNPCRes_aNPC_rna_met

#Nominal p-values plot individual omics
data_plot <- data.frame(pval=c(res$expr[,"Pr(>|z|)"],res$met[,"Pr(>|z|)"]), 
                        group=c(rep(names(res)[1], nrow(res$expr)), 
                                rep(names(res)[2], nrow(res$met))))

p1 <- ggplot(data_plot, aes(x=group, y=pval, fill=group)) +
  geom_violin(trim=FALSE)+
  labs(title='Plot of individual omics p-value from overlapping samples')

#FDR plot individual omics
data_plot <- data.frame(pval=c(res$expr[,"adj.P.Val"],res$met[,"adj.P.Val"]), 
                        group=c(rep(names(res)[1], nrow(res$expr)),
                                rep(names(res)[2], nrow(res$met))))

p2 <- ggplot(data_plot, aes(x=group, y=pval, fill=group)) +
  geom_violin(trim=FALSE)+
  labs(title='Plot of individual omics FDR from overlapping samples')


#Nominal p-value plot NPC
data_plot <- data.frame(pval=unlist(res$pvaluesNPC[,-c(1:4)]), 
                        combination_method=rep(colnames(res$pvaluesNPC[,-c(1:4)])
                                               , each=nrow(res$pvaluesNPC)))

p3 <- ggplot(data_plot, aes(x=combination_method, y=pval, 
                            fill=combination_method)) +
  geom_violin(trim=FALSE)+
  labs(title='Plot of p-values in overlapping omicsNPC')


#FDR plot PC
data_plot <- data.frame(pval=unlist(res$qvaluesNPC[,-c(1:4)]), 
                        combination_method=rep(colnames(res$qvaluesNPC[,-c(1:4)])
                                               , each=nrow(res$qvaluesNPC)))

p4 <- ggplot(data_plot, aes(x=combination_method, y=pval, 
                            fill=combination_method)) +
  geom_violin(trim=FALSE)+
  labs(title='Plot of FDR in overlapping omicsNPC')


grid.arrange(p1, p2, p3, p4, ncol=2, nrow=2)
```

Now, based on FDR results we can check if there are an increase of statistical power when both omics are jointly analyzed.

Note: here we consider significant an FDR<0.05.

```
#Significant genes or methylation sites from individual omics (FDR<0.05)

#significant genes
(sig_expr_aNPC <- rownames(res$expr)[which(res$expr[,"adj.P.Val"]<0.05)])
```

```
## [1] "FAM46A"  "FNDC3B"  "FUT4"    "FZD7"    "PRM2"    "SIGLEC9" "SNX10"
```

```
#7 genes

#significant methylation sites
(sig_met_aNPC <- rownames(res$met)[which(res$met[,"adj.P.Val"]<0.05)])
```

```
## character(0)
```

```
#0 methylation sites

#Significant genes from mRNA + met non parametric combination (Fisher, FDR<0.05)
sig_expr_met_aNPC <- res$qvaluesNPC[which(res$qvaluesNPC[,'Fisher']<0.05),1:2]
# 332 pairs

#venn diagram
m1 <- list(rna=sig_expr_aNPC, rna_fisher=unique(sig_expr_met_aNPC$expr))

vennPlot1 <- venn.diagram(m1, NULL, fill=c('red','blue'), 
                         cat.fontface=4, lty=2, cex=1.2, cat.cex=1.2, main='Significant genes identified with individual (rna) or integration (rna_fisher)\n omics analysis: NPC (all samples)')
grid.newpage()
grid.draw(vennPlot1)
```

```
m2 <- list(met=sig_met_aNPC, met_fisher=unique(sig_expr_met_aNPC$met))

vennPlot2 <- venn.diagram(m2, NULL, fill=c('green','blue'), 
                         cat.fontface=4, lty=2, cex=1.2, cat.cex=1.2, main='Significant methylation sites identified with individual (met) or integration (met_fisher)\n omics analysis: NPC (all samples)')

grid.newpage()
grid.draw(vennPlot2)
```

When we perform the combination analysis we identify 174 genes related with GBM survival and 332 methylation site.

Here we can see a subset of the new genes (n=174) identified based on Fisher combination.

```
setdiff(sig_expr_met_aNPC$expr,sig_expr_aNPC)[1:50]
```

```
##  [1] "ABCB9"    "ADK"      "AHNAK2"   "AIM1"     "ALX1"     "AMPD3"   
##  [7] "ANXA11"   "APOBEC3C" "ARHGAP25" "ARPC1B"   "ASCC1"    "ASCL2"   
## [13] "ASPSCR1"  "ATP1A1"   "ATP2A1"   "AVIL"     "B3GAT3"   "BCAN"    
## [19] "BCL3"     "BMP4"     "BRF1"     "BST2"     "BTNL2"    "C13orf15"
## [25] "C2"       "C6orf48"  "CALCR"    "CAMTA2"   "CCR1"     "CDCP1"   
## [31] "CDYL"     "CHI3L1"   "CHST2"    "COL16A1"  "CPD"      "CRHR2"   
## [37] "CSNK1D"   "CTSZ"     "CUTA"     "CYB5R2"   "DENND2D"  "DGKZ"    
## [43] "DKK3"     "DLC1"     "DUS1L"    "ECHDC2"   "EMILIN1"  "EPB41L3" 
## [49] "ETFA"     "F3"
```

And the new methylation sites (n=332) identified based on Fisher combination.

```
setdiff(sig_expr_met_aNPC$met,sig_met_aNPC)[1:50]
```

```
##  [1] "cg03328639" "cg16088251" "cg02719508" "cg23413349" "cg02409351"
##  [6] "cg14116122" "cg14996220" "cg25800765" "cg10794058" "cg19132462"
## [11] "cg18010131" "cg25968076" "cg11647108" "cg06523744" "cg15258980"
## [16] "cg17140191" "cg19577016" "cg27252164" "cg05309948" "cg04571951"
## [21] "cg21437208" "cg11342277" "cg15990192" "cg13079047" "cg10442913"
## [26] "cg00474713" "cg04046364" "cg25217772" "cg25217710" "cg17740327"
## [31] "cg09229893" "cg16389901" "cg04269188" "cg16313343" "cg01254505"
## [36] "cg17049418" "cg19281363" "cg00025981" "cg07249939" "cg12484688"
## [41] "cg14734916" "cg17243044" "cg13127825" "cg25110523" "cg06636203"
## [46] "cg25013586" "cg00058449" "cg06804823" "cg16617910" "cg09578319"
```

###### 2.7.2.4 Integration strategies comparison

Finally, we can compare the divergences between parametric and non-parametric approaches and the overlapping or whole cohort escenarios.

```
#Significant pairs from different approaches

#parmetric combination with overlapping samples
oPC <- paste(sig_expr_met_cPC$expr,sig_expr_met_cPC$met, sep=":") #323
#parmetric combination with all samples
aPC <- paste(sig_expr_met_aPC$expr,sig_expr_met_aPC$met, sep=":") #207
#non parmetric combination with overlapping samples
oNPC <- paste(sig_expr_met_cNPC$expr,sig_expr_met_cNPC$met, sep=":") #150
#non parmetric combination with all samples
aNPC <- paste(sig_expr_met_aNPC$expr,sig_expr_met_aNPC$met, sep=":") #332

res_all <- list(oPC=oPC, aPC=aPC, oNPC=oNPC, aNPC=aNPC)

vennPlot_m1 <- venn.diagram(res_all, NULL, fill=c("green","blue","orange","pink"), 
                            cat.fontface=4, lty=2, cex=1.2, cat.cex=1.2, 
                          main="Significant pairs for the different escenarios \n FDR (cut-off=0.05)")

grid.newpage()
grid.draw(vennPlot_m1)
```

In this approach we can see that the results differ considerably depending on the type of approach used.

Finally, you can see the R session information used to run all the code.

```
sessionInfo()
```

```
## R version 4.0.3 (2020-10-10)
## Platform: x86_64-w64-mingw32/x64 (64-bit)
## Running under: Windows 10 x64 (build 16299)
## 
## Matrix products: default
## 
## Random number generation:
##  RNG:     Mersenne-Twister 
##  Normal:  Inversion 
##  Sample:  Rounding 
##  
## locale:
## [1] LC_COLLATE=Spanish_Spain.1252  LC_CTYPE=Spanish_Spain.1252   
## [3] LC_MONETARY=Spanish_Spain.1252 LC_NUMERIC=C                  
## [5] LC_TIME=Spanish_Spain.1252    
## 
## attached base packages:
##  [1] grid      stats4    parallel  stats     graphics  grDevices utils    
##  [8] datasets  methods   base     
## 
## other attached packages:
##  [1] data.table_1.13.0                                 
##  [2] anamiR_1.13.0                                     
##  [3] miRNAmeConverter_1.16.0                           
##  [4] miRBaseVersions.db_1.1.0                          
##  [5] r.jive_2.1                                        
##  [6] sva_3.36.0                                        
##  [7] BiocParallel_1.22.0                               
##  [8] genefilter_1.70.0                                 
##  [9] mgcv_1.8-33                                       
## [10] nlme_3.1-149                                      
## [11] gridExtra_2.3                                     
## [12] MASS_7.3-53                                       
## [13] STATegRa_1.17.1                                   
## [14] plotrix_3.7-8                                     
## [15] VennDiagram_1.6.20                                
## [16] futile.logger_1.4.3                               
## [17] lumi_2.40.0                                       
## [18] IlluminaHumanMethylation450kanno.ilmn12.hg19_0.6.0
## [19] minfi_1.34.0                                      
## [20] bumphunter_1.30.0                                 
## [21] locfit_1.5-9.4                                    
## [22] iterators_1.0.13                                  
## [23] foreach_1.5.1                                     
## [24] Biostrings_2.56.0                                 
## [25] XVector_0.28.0                                    
## [26] SummarizedExperiment_1.18.2                       
## [27] DelayedArray_0.14.1                               
## [28] matrixStats_0.57.0                                
## [29] Biobase_2.48.0                                    
## [30] GenomicRanges_1.40.0                              
## [31] GenomeInfoDb_1.24.2                               
## [32] IRanges_2.22.2                                    
## [33] S4Vectors_0.26.1                                  
## [34] BiocGenerics_0.34.0                               
## [35] ggfortify_0.4.11                                  
## [36] ggplot2_3.3.2                                     
## [37] survival_3.2-7                                    
## [38] reshape_0.8.8                                     
## [39] BiocStyle_2.16.1                                  
## 
## loaded via a namespace (and not attached):
##   [1] questionr_0.7.3           tidyselect_1.1.0         
##   [3] RSQLite_2.2.1             AnnotationDbi_1.50.3     
##   [5] combinat_0.0-8            munsell_0.5.0            
##   [7] codetools_0.2-16          preprocessCore_1.50.0    
##   [9] nleqslv_3.3.2             miniUI_0.1.1.1           
##  [11] withr_2.3.0               colorspace_1.4-1         
##  [13] AlgDesign_1.2.0           highr_0.8                
##  [15] knitr_1.30                rstudioapi_0.11          
##  [17] labeling_0.4.2            GenomeInfoDbData_1.2.3   
##  [19] bit64_4.0.5               farver_2.0.3             
##  [21] rhdf5_2.32.4              vctrs_0.3.4              
##  [23] generics_0.0.2            lambda.r_1.2.4           
##  [25] xfun_0.18                 BiocFileCache_1.12.1     
##  [27] R6_2.4.1                  illuminaio_0.30.0        
##  [29] bitops_1.0-6              assertthat_0.2.1         
##  [31] promises_1.1.1            scales_1.1.1             
##  [33] gtable_0.3.0              affy_1.66.0              
##  [35] methylumi_2.34.0          rlang_0.4.8              
##  [37] calibrate_1.7.7           splines_4.0.3            
##  [39] rtracklayer_1.48.0        GEOquery_2.56.0          
##  [41] BiocManager_1.30.10       yaml_2.2.1               
##  [43] abind_1.4-5               GenomicFeatures_1.40.1   
##  [45] httpuv_1.5.4              RMySQL_0.10.20           
##  [47] SpatioTemporal_1.1.9.1    tools_4.0.3              
##  [49] bookdown_0.21             nor1mix_1.3-0            
##  [51] affyio_1.58.0             ellipsis_0.3.1           
##  [53] gplots_3.1.0              RColorBrewer_1.1-2       
##  [55] siggenes_1.62.0           Rcpp_1.0.5               
##  [57] plyr_1.8.6                progress_1.2.2           
##  [59] zlibbioc_1.34.0           purrr_0.3.4              
##  [61] RCurl_1.98-1.2            prettyunits_1.1.1        
##  [63] openssl_1.4.3             cluster_2.1.0            
##  [65] haven_2.3.1               magrittr_1.5             
##  [67] futile.options_1.0.1      hms_0.5.3                
##  [69] mime_0.9                  evaluate_0.14            
##  [71] xtable_1.8-4              klaR_0.6-15              
##  [73] XML_3.99-0.5              mclust_5.4.6             
##  [75] compiler_4.0.3            biomaRt_2.44.4           
##  [77] gage_2.38.3               tibble_3.0.4             
##  [79] KernSmooth_2.23-17        crayon_1.3.4             
##  [81] agricolae_1.3-3           htmltools_0.5.0          
##  [83] later_1.1.0.1             geneplotter_1.66.0       
##  [85] tidyr_1.1.2               DBI_1.1.0                
##  [87] formatR_1.7               dbplyr_1.4.4             
##  [89] rappdirs_0.3.1            Matrix_1.2-18            
##  [91] readr_1.4.0               quadprog_1.5-8           
##  [93] forcats_0.5.0             pkgconfig_2.0.3          
##  [95] GenomicAlignments_1.24.0  xml2_1.3.2               
##  [97] annotate_1.66.0           rngtools_1.5             
##  [99] multtest_2.44.0           beanplot_1.2             
## [101] doRNG_1.8.2               scrime_1.3.5             
## [103] stringr_1.4.0             digest_0.6.25            
## [105] graph_1.66.0              rmarkdown_2.4            
## [107] base64_2.0                edgeR_3.30.3             
## [109] DelayedMatrixStats_1.10.1 curl_4.3                 
## [111] shiny_1.5.0               Rsamtools_2.4.0          
## [113] gtools_3.8.2              lifecycle_0.2.0          
## [115] Rhdf5lib_1.10.1           askpass_1.1              
## [117] limma_3.44.3              labelled_2.7.0           
## [119] pillar_1.4.6              lattice_0.20-41          
## [121] KEGGREST_1.28.0           GO.db_3.11.4             
## [123] fastmap_1.0.1             httr_1.4.2               
## [125] glue_1.4.2                png_0.1-7                
## [127] bit_4.0.4                 stringi_1.5.3            
## [129] HDF5Array_1.16.1          blob_1.2.1               
## [131] DESeq2_1.28.1             caTools_1.18.0           
## [133] memoise_1.1.0             dplyr_1.0.2
```
