## Supplementary material for "STATegra: Multi-omics data integration - A conceptual scheme and a bioinformatics pipeline": RMarkDown and html files.: stategra_GeneSetCluster.html


### Stategra\_GeneSetCluster

###### Ewoud Ewing

#### 11/13/2020

- 0.1 Introduction
- 0.2 Analysis of Canonical Pathways
  - 0.2.1 RNA.Met.skcm.Can
  - 0.2.2 RNA.Mir.skcm.Can
  - 0.2.3 RNA.Met.GBM.Can
  - 0.2.4 RNA.Mir.GBM.Can
- 0.3 Analysis of Disease and Functions
  - 0.3.1 RNA.Met.skcm.Func
  - 0.3.2 RNA.Mir.skcm.Func
  - 0.3.3 RNA.Met.GBM.Func
  - 0.3.4 RNA.Mir.GBM.Func
- 0.4 Exporting the results

#### 0.1 Introduction

This is an R Markdown document of the GeneSet. For more details on using GeneSetCluster see https://github.com/TranslationalBioinformaticsUnit/GeneSetCluster.

Cite: Ewing, E., Planell-Picola, N., Jagodic, M. et al. GeneSetCluster: a tool for summarizing and integrating gene-set analysis results. BMC Bioinformatics 21, 443 (2020). https://doi.org/10.1186/s12859-020-03784-z

```
require(GeneSetCluster)
```

```
## Loading required package: GeneSetCluster
```

```
##
```

```
##
```

```
##
```

```
require(ggplot2)
```

```
## Loading required package: ggplot2
```

```
RNA.Mir.GBM <- list.files(path = "~/Stategra/Stategra_GSC/gbm/rna_mirna/", full.names = T, recursive = T)
RNA.Met.GBM <- list.files(path = "~/Stategra/Stategra_GSC/gbm/rna_met/", full.names = T, recursive = T)
RNA.Mir.skcm <- list.files(path = "~/Stategra/Stategra_GSC/skcm/rna_mirna/", full.names = T, recursive = T)
RNA.Met.skcm <- list.files(path = "~/Stategra/Stategra_GSC/skcm/rna_met/", full.names = T, recursive = T)

RNA.Mir.GBM.Can <- RNA.Mir.GBM[grep("Canonical", RNA.Mir.GBM)]
RNA.Met.GBM.Can <- RNA.Met.GBM[grep("Canonical", RNA.Met.GBM)]
RNA.Mir.skcm.Can <- RNA.Mir.skcm[grep("Canonical", RNA.Mir.skcm)]
RNA.Met.skcm.Can <- RNA.Met.skcm[grep("Canonical", RNA.Met.skcm)]
RNA.Mir.GBM.Func <- RNA.Mir.GBM[grep("Disease_functions", RNA.Mir.GBM)]
RNA.Met.GBM.Func <- RNA.Met.GBM[grep("Disease_functions", RNA.Met.GBM)]
RNA.Mir.skcm.Func <- RNA.Mir.skcm[grep("Disease_functions", RNA.Mir.skcm)]
RNA.Met.skcm.Func <- RNA.Met.skcm[grep("Disease_functions", RNA.Met.skcm)]
```

#### 0.2 Analysis of Canonical Pathways

IPA give the user a lot of output, we focused on the Canonical Pathways and disease and functions, which we analysed seperatly.

##### 0.2.1 RNA.Met.skcm.Can

We make an PathwayObject for every group seperate for Canonical Pathways and disease and functions. GeneSetCluster could combine them but then it wouldnt be as clear what the additional functional informaton comes from the OmicsNPC.

```
groups <- limma::strsplit2(RNA.Met.skcm.Can, split="/")[,11]
groups <- gsub(pattern = "20200921_", replacement = "", groups)

RNA.Met.skcm.Can.Obj2<- LoadGeneSets(file_location = RNA.Met.skcm.Can[2:4], 
                                     groupnames = groups[2:4], P.cutoff = 1.31, Mol.cutoff =3, 
                                     Source = "IPA", type = "Canonical_Pathways", 
                                     structure = "SYMBOL", Organism = "org.Hs.eg.db", seperator = ",")
```

```
## [=========================================================]
```

```
## [<<<<            LoadGeneSets START                  >>>>>]
```

```
## -----------------------------------------------------------
```

```
## Loading data from /mnt/hel/home/ewoewi/Stategra/Stategra_GSC/skcm/rna_met//20200921_npc_rna_met_sig_new_genes_skcm_corrected/Canonical_pathways.xls
```

```
## Loading data from /mnt/hel/home/ewoewi/Stategra/Stategra_GSC/skcm/rna_met//20200921_npc_rna_met_skcm/Canonical_pathways.xls
```

```
## Loading data from /mnt/hel/home/ewoewi/Stategra/Stategra_GSC/skcm/rna_met//20200921_single_genes_rna_met_skcm/Canonical_pathways.xls
```

```
## Loading IPA Canonical_Pathways
```

```
## -----------------------------------------------------------
```

```
## [<<<<<               LoadGeneSets END               >>>>>>]
```

```
## [=========================================================]
```

```
## [You may want to process CombineGeneSets next.            ]
```

```
## [or merge objects using MergeObjects                     ]
```

```
## [or select certain types from objects using ManageGeneSets]
```

```
#There is a build in loader for IPA and GREAT datasets

RNA.Met.skcm.Can.Obj2 <- CombineGeneSets(RNA.Met.skcm.Can.Obj2,combineMethod ="Standard",  display = "Expanded")
```

```
## [=========================================================]
```

```
## [<<<<            CombineGeneSets START               >>>>>]
```

```
## -----------------------------------------------------------
```

```
## transforming all genes to upper case, make sure this doesnt change the data
```

```
## raw data has 124 genes
```

```
## Transformed data has 124 genes
```

```
## Combining experiments
```

```
## preparing expanded display
```

```
## calulating RR
```

```
## -----------------------------------------------------------
```

```
## [<<<<<             CombineGeneSets END              >>>>>>]
```

```
## [=========================================================]
```

```
## [You may want to process ClusterGeneSets next.            ]
```

```
#Here we calculate the relative risk between the different Gene Sets

#OptimalGeneSets(RNA.Met.skcm.Can.Obj, method = "gap",cluster_method = "kmeans", max_cluster = 10, main="RNA.Met.skcm.Can.Obj")
#Calculate the optimal number of clusters

RNA.Met.skcm.Can.Obj2 <- ClusterGeneSets(RNA.Met.skcm.Can.Obj2, clusters = 6, method = "kmeans", order = "cluster")
```

```
## [=========================================================]
```

```
## [<<<<            ClusterGeneSets START               >>>>>]
```

```
## -----------------------------------------------------------
```

```
## Using all Gene sets
```

```
## Running kmeans
```

```
## Ordering pathway clusters
```

```
## -----------------------------------------------------------
```

```
## [<<<<<             ClusterGeneSets END              >>>>>>]
```

```
## [=========================================================]
```

```
## [You may want to process HighlightGeneSets next.            ]
```

```
## [You may want to plot the results using PlotGeneSets next. ]
```

```
#Here we cluster using Kmeans in the 6 clusters determined by OptimalGeneSets


#Because of how we want the plots to look, we remove the group information so its just the pathways where they appear.
x <- RNA.Met.skcm.Can.Obj2@plot$aka2
x <- x[,!colnames(x) == "Group"]
RNA.Met.skcm.Can.Obj2@plot$aka2 <- x

x <- RNA.Met.skcm.Can.Obj2@plot$aka3
x <- x[2:length(x)]
RNA.Met.skcm.Can.Obj2@plot$aka3 <- x
```

```
#Generating the plot
PlotGeneSets(Object = RNA.Met.skcm.Can.Obj2, main = "RNA.Met.skcm.Can.Obj", RR.max = 50)
```

##### 0.2.2 RNA.Mir.skcm.Can

```
groups <- limma::strsplit2(RNA.Mir.skcm.Can, split="/")[,11]
groups <- gsub(pattern = "20200921_", replacement = "", groups)
RNA.Mir.skcm.Can.Obj2 <- LoadGeneSets(file_location = RNA.Mir.skcm.Can[2:4], 
                                     groupnames = groups[2:4], P.cutoff = 1.31, Mol.cutoff =3, 
                                     Source = "IPA", type = "Canonical_Pathways", 
                                     structure = "SYMBOL", Organism = "org.Hs.eg.db", seperator = ",")
```

```
## [=========================================================]
```

```
## [<<<<            LoadGeneSets START                  >>>>>]
```

```
## -----------------------------------------------------------
```

```
## Loading data from /mnt/hel/home/ewoewi/Stategra/Stategra_GSC/skcm/rna_mirna//20200921_npc_rna_mirna_sig_new_genes_skcm_corrected/Canonical_pathways.xls
```

```
## Loading data from /mnt/hel/home/ewoewi/Stategra/Stategra_GSC/skcm/rna_mirna//20200921_npc_rna_mirna_skcm/Canonical_pathways.xls
```

```
## Loading data from /mnt/hel/home/ewoewi/Stategra/Stategra_GSC/skcm/rna_mirna//20200921_single_genes_rna_mirna_skcm/Canonical_pathways.xls
```

```
## Loading IPA Canonical_Pathways
```

```
## -----------------------------------------------------------
```

```
## [<<<<<               LoadGeneSets END               >>>>>>]
```

```
## [=========================================================]
```

```
## [You may want to process CombineGeneSets next.            ]
```

```
## [or merge objects using MergeObjects                     ]
```

```
## [or select certain types from objects using ManageGeneSets]
```

```
RNA.Mir.skcm.Can.Obj2 <- CombineGeneSets(RNA.Mir.skcm.Can.Obj2, display = "Expanded")
```

```
## [=========================================================]
```

```
## [<<<<            CombineGeneSets START               >>>>>]
```

```
## -----------------------------------------------------------
```

```
## transforming all genes to upper case, make sure this doesnt change the data
```

```
## raw data has 79 genes
```

```
## Transformed data has 79 genes
```

```
## Combining experiments
```

```
## preparing expanded display
```

```
## calulating RR
```

```
## -----------------------------------------------------------
```

```
## [<<<<<             CombineGeneSets END              >>>>>>]
```

```
## [=========================================================]
```

```
## [You may want to process ClusterGeneSets next.            ]
```

```
#OptimalGeneSets(RNA.Mir.skcm.Can.Obj, method = "gap",cluster_method = "kmeans", max_cluster = 10, main="RNA.Mir.skcm.Can.Obj")

RNA.Mir.skcm.Can.Obj2 <- ClusterGeneSets(RNA.Mir.skcm.Can.Obj2, clusters = 7, method = "kmeans", order = "cluster")
```

```
## [=========================================================]
```

```
## [<<<<            ClusterGeneSets START               >>>>>]
```

```
## -----------------------------------------------------------
```

```
## Using all Gene sets
```

```
## Running kmeans
```

```
## Ordering pathway clusters
```

```
## -----------------------------------------------------------
```

```
## [<<<<<             ClusterGeneSets END              >>>>>>]
```

```
## [=========================================================]
```

```
## [You may want to process HighlightGeneSets next.            ]
```

```
## [You may want to plot the results using PlotGeneSets next. ]
```

```
x <- RNA.Mir.skcm.Can.Obj2@plot$aka2
x <- x[,!colnames(x) == "Group"]
RNA.Mir.skcm.Can.Obj2@plot$aka2 <- x

x <- RNA.Mir.skcm.Can.Obj2@plot$aka3
x <- x[2:length(x)]
RNA.Mir.skcm.Can.Obj2@plot$aka3 <- x

PlotGeneSets(Object = RNA.Mir.skcm.Can.Obj2, main = "RNA.Mir.skcm.Can.Obj", RR.max = 50)
```

```
#Generating the plot
PlotGeneSets(Object = RNA.Mir.skcm.Can.Obj2, main = "RNA.Mir.skcm.Can.Obj", RR.max = 50)
```

##### 0.2.3 RNA.Met.GBM.Can

```
groups <- limma::strsplit2(RNA.Met.GBM.Can, split="/")[,11]
groups <- gsub(pattern = "20200921_", replacement = "", groups)
groups <- gsub(pattern = "_Correct", replacement = "", groups)

RNA.Met.GBM.Can.Obj2 <- LoadGeneSets(file_location = RNA.Met.GBM.Can[c(1,2,4,6)], 
                                    groupnames = groups[c(1,2,4,6)], P.cutoff = 1.31, Mol.cutoff =3, 
                                    Source = "IPA", type = "Canonical_Pathways", 
                                    structure = "SYMBOL", Organism = "org.Hs.eg.db", seperator = ",")
```

```
## [=========================================================]
```

```
## [<<<<            LoadGeneSets START                  >>>>>]
```

```
## -----------------------------------------------------------
```

```
## Loading data from /mnt/hel/home/ewoewi/Stategra/Stategra_GSC/gbm/rna_met//20200921_npc_rna_met_all_gbm/Canonical_pathways.xls
```

```
## Loading data from /mnt/hel/home/ewoewi/Stategra/Stategra_GSC/gbm/rna_met//20200921_npc_rna_met_common_gbm_Correct/Canonical_pathways.xls
```

```
## Loading data from /mnt/hel/home/ewoewi/Stategra/Stategra_GSC/gbm/rna_met//20200921_npc_rna_met_sig_new_genes_all_gbm_corrected/Canonical_pathways.xls
```

```
## Loading data from /mnt/hel/home/ewoewi/Stategra/Stategra_GSC/gbm/rna_met//20200921_npc_rna_met_sig_new_genes_common_gbm_corrected/Canonical_pathways.xls
```

```
## Loading IPA Canonical_Pathways
```

```
## -----------------------------------------------------------
```

```
## [<<<<<               LoadGeneSets END               >>>>>>]
```

```
## [=========================================================]
```

```
## [You may want to process CombineGeneSets next.            ]
```

```
## [or merge objects using MergeObjects                     ]
```

```
## [or select certain types from objects using ManageGeneSets]
```

```
RNA.Met.GBM.Can.Obj2 <-CombineGeneSets(RNA.Met.GBM.Can.Obj2, display = "Expanded")
```

```
## [=========================================================]
```

```
## [<<<<            CombineGeneSets START               >>>>>]
```

```
## -----------------------------------------------------------
```

```
## transforming all genes to upper case, make sure this doesnt change the data
```

```
## raw data has 37 genes
```

```
## Transformed data has 37 genes
```

```
## Combining experiments
```

```
## preparing expanded display
```

```
## calulating RR
```

```
## -----------------------------------------------------------
```

```
## [<<<<<             CombineGeneSets END              >>>>>>]
```

```
## [=========================================================]
```

```
## [You may want to process ClusterGeneSets next.            ]
```

```
RNA.Met.GBM.Can.Obj2 <- ClusterGeneSets(RNA.Met.GBM.Can.Obj2, clusters = 4, method = "kmeans", order = "cluster")
```

```
## [=========================================================]
```

```
## [<<<<            ClusterGeneSets START               >>>>>]
```

```
## -----------------------------------------------------------
```

```
## Using all Gene sets
```

```
## Running kmeans
```

```
## Ordering pathway clusters
```

```
## -----------------------------------------------------------
```

```
## [<<<<<             ClusterGeneSets END              >>>>>>]
```

```
## [=========================================================]
```

```
## [You may want to process HighlightGeneSets next.            ]
```

```
## [You may want to plot the results using PlotGeneSets next. ]
```

```
x <- RNA.Met.GBM.Can.Obj2@plot$aka2
x <- x[,!colnames(x) == "Group"]
RNA.Met.GBM.Can.Obj2@plot$aka2 <- x

x <- RNA.Met.GBM.Can.Obj2@plot$aka3
x <- x[2:length(x)]
RNA.Met.GBM.Can.Obj2@plot$aka3 <- x
```

```
#Generating the plot
PlotGeneSets(Object = RNA.Met.GBM.Can.Obj2, main = "RNA.Met.GBM.Can.Obj", RR.max = 50)
```

##### 0.2.4 RNA.Mir.GBM.Can

```
groups <- limma::strsplit2(RNA.Mir.GBM.Can, split="/")[,11]
groups <- gsub(pattern = "20200921_", replacement = "", groups)
#groups <- gsub(pattern = "_correct", replacement = "", groups)

idx <- c(1,4)
RNA.Mir.GBM.Can.Obj2 <- LoadGeneSets(file_location = RNA.Mir.GBM.Can[idx], 
                                    groupnames = groups[idx], P.cutoff = 1.31, Mol.cutoff =3, 
                                    Source = "IPA", type = "Canonical_Pathways", 
                                    structure = "SYMBOL", Organism = "org.Hs.eg.db", seperator = ",")
```

```
## [=========================================================]
```

```
## [<<<<            LoadGeneSets START                  >>>>>]
```

```
## -----------------------------------------------------------
```

```
## Loading data from /mnt/hel/home/ewoewi/Stategra/Stategra_GSC/gbm/rna_mirna//20200921_npc_rna_mirna_all_gbm_correct/Canonical_pathways.xls
```

```
## Loading data from /mnt/hel/home/ewoewi/Stategra/Stategra_GSC/gbm/rna_mirna//20200921_npc_rna_mirna_sig_new_genes_all_gbm_corrected/Canonical_pathways.xls
```

```
## Loading IPA Canonical_Pathways
```

```
## -----------------------------------------------------------
```

```
## [<<<<<               LoadGeneSets END               >>>>>>]
```

```
## [=========================================================]
```

```
## [You may want to process CombineGeneSets next.            ]
```

```
## [or merge objects using MergeObjects                     ]
```

```
## [or select certain types from objects using ManageGeneSets]
```

```
RNA.Mir.GBM.Can.Obj2 <- CombineGeneSets(RNA.Mir.GBM.Can.Obj2, display = "Expanded")
```

```
## [=========================================================]
```

```
## [<<<<            CombineGeneSets START               >>>>>]
```

```
## -----------------------------------------------------------
```

```
## transforming all genes to upper case, make sure this doesnt change the data
```

```
## raw data has 9 genes
```

```
## Transformed data has 9 genes
```

```
## Combining experiments
```

```
## preparing expanded display
```

```
## calulating RR
```

```
## -----------------------------------------------------------
```

```
## [<<<<<             CombineGeneSets END              >>>>>>]
```

```
## [=========================================================]
```

```
## [You may want to process ClusterGeneSets next.            ]
```

```
RNA.Mir.GBM.Can.Obj2 <- ClusterGeneSets(RNA.Mir.GBM.Can.Obj2, clusters = 2, method = "kmeans", order = "cluster")
```

```
## [=========================================================]
```

```
## [<<<<            ClusterGeneSets START               >>>>>]
```

```
## -----------------------------------------------------------
```

```
## Using all Gene sets
```

```
## Running kmeans
```

```
## Ordering pathway clusters
```

```
## -----------------------------------------------------------
```

```
## [<<<<<             ClusterGeneSets END              >>>>>>]
```

```
## [=========================================================]
```

```
## [You may want to process HighlightGeneSets next.            ]
```

```
## [You may want to plot the results using PlotGeneSets next. ]
```

```
x <- RNA.Mir.GBM.Can.Obj2@plot$aka2
x <- x[,!colnames(x) == "Group"]
RNA.Mir.GBM.Can.Obj2@plot$aka2 <- x

x <- RNA.Mir.GBM.Can.Obj2@plot$aka3
x <- x[2:length(x)]
RNA.Mir.GBM.Can.Obj2@plot$aka3 <- x


PlotGeneSets(Object = RNA.Mir.GBM.Can.Obj2, main = "RNA.Mir.GBM.Can.Obj", RR.max = 50)
```

```
#Generating the plot
PlotGeneSets(Object = RNA.Mir.GBM.Can.Obj2, main = "RNA.Mir.GBM.Can.Obj", RR.max = 50)
```

#### 0.3 Analysis of Disease and Functions

Here we are analysing the Disease and Functions output from IPA.

##### 0.3.1 RNA.Met.skcm.Func

```
groups <- limma::strsplit2(RNA.Met.skcm.Func, split="/")[,11]
groups <- gsub(pattern = "20200921_", replacement = "", groups)

idx <- c(1,3,4,5)

RNA.Met.skcm.Func.Obj2 <- LoadGeneSets(file_location = RNA.Met.skcm.Func[idx], 
                                      groupnames = groups[idx], P.cutoff = 0.05, Mol.cutoff =3, 
                                      Source = "IPA", type = "Functional_annotations", 
                                      structure = "SYMBOL", Organism = "org.Hs.eg.db", seperator = ",")
```

```
## [=========================================================]
```

```
## [<<<<            LoadGeneSets START                  >>>>>]
```

```
## -----------------------------------------------------------
```

```
## Loading data from /mnt/hel/home/ewoewi/Stategra/Stategra_GSC/skcm/rna_met//20200921_npc_rna_met_sig_new_genes_skcm_corrected/Disease_functions.xls
```

```
## Loading data from /mnt/hel/home/ewoewi/Stategra/Stategra_GSC/skcm/rna_met//20200921_npc_rna_met_skcm/Disease_functions.xls
```

```
## Loading data from /mnt/hel/home/ewoewi/Stategra/Stategra_GSC/skcm/rna_met//20200921_single_genes_rna_met_skcm/Disease_functions.xls
```

```
## Loading data from /mnt/hel/home/ewoewi/Stategra/Stategra_GSC/skcm/rna_met//20200921_single_met_rna_met_skcm/Disease_functions.xls
```

```
## Loading IPA Functional_annotations
```

```
## -----------------------------------------------------------
```

```
## [<<<<<               LoadGeneSets END               >>>>>>]
```

```
## [=========================================================]
```

```
## [You may want to process CombineGeneSets next.            ]
```

```
## [or merge objects using MergeObjects                     ]
```

```
## [or select certain types from objects using ManageGeneSets]
```

```
RNA.Met.skcm.Func.Obj2 <- CombineGeneSets(RNA.Met.skcm.Func.Obj2, display = "Expanded")
```

```
## [=========================================================]
```

```
## [<<<<            CombineGeneSets START               >>>>>]
```

```
## -----------------------------------------------------------
```

```
## transforming all genes to upper case, make sure this doesnt change the data
```

```
## raw data has 387 genes
```

```
## Transformed data has 387 genes
```

```
## Combining experiments
```

```
## preparing expanded display
```

```
## calulating RR
```

```
## -----------------------------------------------------------
```

```
## [<<<<<             CombineGeneSets END              >>>>>>]
```

```
## [=========================================================]
```

```
## [You may want to process ClusterGeneSets next.            ]
```

```
RNA.Met.skcm.Func.Obj2 <- ClusterGeneSets(RNA.Met.skcm.Func.Obj2, clusters = 4, method = "kmeans", order = "cluster")
```

```
## [=========================================================]
```

```
## [<<<<            ClusterGeneSets START               >>>>>]
```

```
## -----------------------------------------------------------
```

```
## Using all Gene sets
```

```
## Running kmeans
```

```
## Ordering pathway clusters
```

```
## -----------------------------------------------------------
```

```
## [<<<<<             ClusterGeneSets END              >>>>>>]
```

```
## [=========================================================]
```

```
## [You may want to process HighlightGeneSets next.            ]
```

```
## [You may want to plot the results using PlotGeneSets next. ]
```

```
x <- RNA.Met.skcm.Func.Obj2@plot$aka2
x <- x[,!colnames(x) == "Group"]
RNA.Met.skcm.Func.Obj2@plot$aka2 <- x

x <- RNA.Met.skcm.Func.Obj2@plot$aka3
x <- x[2:length(x)]
RNA.Met.skcm.Func.Obj2@plot$aka3 <- x
```

```
#Generating the plot
PlotGeneSets(Object = RNA.Met.skcm.Func.Obj2, main = "RNA.Met.skcm.Func.Obj")
```

##### 0.3.2 RNA.Mir.skcm.Func

```
groups <- limma::strsplit2(RNA.Mir.skcm.Func, split="/")[,11]
groups <- gsub(pattern = "20200921_", replacement = "", groups)

idx <- c(1,3:5)

RNA.Mir.skcm.Func.Obj2 <- LoadGeneSets(file_location = RNA.Mir.skcm.Func[idx], 
                                      groupnames = groups[idx], P.cutoff = 0.05, Mol.cutoff =3, 
                                      Source = "IPA", type = "Functional_annotations", 
                                      structure = "SYMBOL", Organism = "org.Hs.eg.db", seperator = ",")
```

```
## [=========================================================]
```

```
## [<<<<            LoadGeneSets START                  >>>>>]
```

```
## -----------------------------------------------------------
```

```
## Loading data from /mnt/hel/home/ewoewi/Stategra/Stategra_GSC/skcm/rna_mirna//20200921_npc_rna_mirna_sig_new_genes_skcm_corrected/Disease_functions.xls
```

```
## Loading data from /mnt/hel/home/ewoewi/Stategra/Stategra_GSC/skcm/rna_mirna//20200921_npc_rna_mirna_skcm/Disease_functions.xls
```

```
## Loading data from /mnt/hel/home/ewoewi/Stategra/Stategra_GSC/skcm/rna_mirna//20200921_single_genes_rna_mirna_skcm/Disease_functions.xls
```

```
## Loading data from /mnt/hel/home/ewoewi/Stategra/Stategra_GSC/skcm/rna_mirna//20200921_single_mirna_rna_mirna_skcm/Disease_functions.xls
```

```
## Loading IPA Functional_annotations
```

```
## -----------------------------------------------------------
```

```
## [<<<<<               LoadGeneSets END               >>>>>>]
```

```
## [=========================================================]
```

```
## [You may want to process CombineGeneSets next.            ]
```

```
## [or merge objects using MergeObjects                     ]
```

```
## [or select certain types from objects using ManageGeneSets]
```

```
RNA.Mir.skcm.Func.Obj2 <- CombineGeneSets(RNA.Mir.skcm.Func.Obj2, display = "Expanded")
```

```
## [=========================================================]
```

```
## [<<<<            CombineGeneSets START               >>>>>]
```

```
## -----------------------------------------------------------
```

```
## transforming all genes to upper case, make sure this doesnt change the data
```

```
## raw data has 267 genes
```

```
## Transformed data has 267 genes
```

```
## Combining experiments
```

```
## preparing expanded display
```

```
## calulating RR
```

```
## -----------------------------------------------------------
```

```
## [<<<<<             CombineGeneSets END              >>>>>>]
```

```
## [=========================================================]
```

```
## [You may want to process ClusterGeneSets next.            ]
```

```
RNA.Mir.skcm.Func.Obj2 <- ClusterGeneSets(RNA.Mir.skcm.Func.Obj2, clusters = 7, method = "kmeans", order = "cluster")
```

```
## [=========================================================]
```

```
## [<<<<            ClusterGeneSets START               >>>>>]
```

```
## -----------------------------------------------------------
```

```
## Using all Gene sets
```

```
## Running kmeans
```

```
## Ordering pathway clusters
```

```
## -----------------------------------------------------------
```

```
## [<<<<<             ClusterGeneSets END              >>>>>>]
```

```
## [=========================================================]
```

```
## [You may want to process HighlightGeneSets next.            ]
```

```
## [You may want to plot the results using PlotGeneSets next. ]
```

```
x <- RNA.Mir.skcm.Func.Obj2@plot$aka2
x <- x[,!colnames(x) == "Group"]
RNA.Mir.skcm.Func.Obj2@plot$aka2 <- x

x <- RNA.Mir.skcm.Func.Obj2@plot$aka3
x <- x[2:length(x)]
RNA.Mir.skcm.Func.Obj2@plot$aka3 <- x
```

```
#Generating the plot
PlotGeneSets(Object = RNA.Mir.skcm.Func.Obj2, main = "RNA.Mir.skcm.Func.Obj", RR.max = 75)
```

##### 0.3.3 RNA.Met.GBM.Func

```
groups <- limma::strsplit2(RNA.Met.GBM.Func, split="/")[,11]
groups <- gsub(pattern = "20200921_", replacement = "", groups)
groups <- gsub(pattern = "_Correct", replacement = "", groups)

idx <- c(1,2,3,5)

RNA.Met.GBM.Func.Obj2 <- LoadGeneSets(file_location = RNA.Met.GBM.Func[idx], 
                                     groupnames = groups[idx], P.cutoff = 0.05, Mol.cutoff =3, 
                                     Source = "IPA", type = "Functional_annotations", 
                                     structure = "SYMBOL", Organism = "org.Hs.eg.db", seperator = ",")
```

```
## [=========================================================]
```

```
## [<<<<            LoadGeneSets START                  >>>>>]
```

```
## -----------------------------------------------------------
```

```
## Loading data from /mnt/hel/home/ewoewi/Stategra/Stategra_GSC/gbm/rna_met//20200921_npc_rna_met_all_gbm/Disease_functions.xls
```

```
## Loading data from /mnt/hel/home/ewoewi/Stategra/Stategra_GSC/gbm/rna_met//20200921_npc_rna_met_common_gbm_Correct/Disease_functions.xls
```

```
## Loading data from /mnt/hel/home/ewoewi/Stategra/Stategra_GSC/gbm/rna_met//20200921_npc_rna_met_sig_new_genes_all_gbm_corrected/Disease_functions.xls
```

```
## Loading data from /mnt/hel/home/ewoewi/Stategra/Stategra_GSC/gbm/rna_met//20200921_npc_rna_met_sig_new_genes_common_gbm_corrected/Disease_functions.xls
```

```
## Loading IPA Functional_annotations
```

```
## -----------------------------------------------------------
```

```
## [<<<<<               LoadGeneSets END               >>>>>>]
```

```
## [=========================================================]
```

```
## [You may want to process CombineGeneSets next.            ]
```

```
## [or merge objects using MergeObjects                     ]
```

```
## [or select certain types from objects using ManageGeneSets]
```

```
RNA.Met.GBM.Func.Obj2 <- CombineGeneSets(RNA.Met.GBM.Func.Obj2, display = "Expanded")
```

```
## [=========================================================]
```

```
## [<<<<            CombineGeneSets START               >>>>>]
```

```
## -----------------------------------------------------------
```

```
## transforming all genes to upper case, make sure this doesnt change the data
```

```
## raw data has 104 genes
```

```
## Transformed data has 104 genes
```

```
## Combining experiments
```

```
## preparing expanded display
```

```
## calulating RR
```

```
## -----------------------------------------------------------
```

```
## [<<<<<             CombineGeneSets END              >>>>>>]
```

```
## [=========================================================]
```

```
## [You may want to process ClusterGeneSets next.            ]
```

```
#RNA.Met.GBM.Func.Obj2 <- ClusterGeneSets(RNA.Met.GBM.Func.Obj2, clusters = 5, method = "kmeans")

RNA.Met.GBM.Func.Obj2 <- ClusterGeneSets(RNA.Met.GBM.Func.Obj2, clusters = 5, method = "kmeans", order = "cluster")
```

```
## [=========================================================]
```

```
## [<<<<            ClusterGeneSets START               >>>>>]
```

```
## -----------------------------------------------------------
```

```
## Using all Gene sets
```

```
## Running kmeans
```

```
## Ordering pathway clusters
```

```
## -----------------------------------------------------------
```

```
## [<<<<<             ClusterGeneSets END              >>>>>>]
```

```
## [=========================================================]
```

```
## [You may want to process HighlightGeneSets next.            ]
```

```
## [You may want to plot the results using PlotGeneSets next. ]
```

```
x <- RNA.Met.GBM.Func.Obj2@plot$aka2
x <- x[,!colnames(x) == "Group"]
RNA.Met.GBM.Func.Obj2@plot$aka2 <- x

x <- RNA.Met.GBM.Func.Obj2@plot$aka3
x <- x[2:length(x)]
RNA.Met.GBM.Func.Obj2@plot$aka3 <- x
```

```
#Generating the plot
PlotGeneSets(Object = RNA.Met.GBM.Func.Obj2, main = "RNA.Met.GBM.Func.Obj", RR.max = 50)
```

##### 0.3.4 RNA.Mir.GBM.Func

```
groups <- limma::strsplit2(RNA.Mir.GBM.Func, split="/")[,11]
groups <- gsub(pattern = "20200921_", replacement = "", groups)
groups <- gsub(pattern = "_Correct", replacement = "", groups)

idx <- c(1,2,4,5,7)

RNA.Mir.GBM.Func.Obj2 <- LoadGeneSets(file_location = RNA.Mir.GBM.Func[idx], 
                                     groupnames = groups[idx], P.cutoff = 0.05, Mol.cutoff =3, 
                                     Source = "IPA", type = "Functional_annotations", 
                                     structure = "SYMBOL", Organism = "org.Hs.eg.db", seperator = ",")
```

```
## [=========================================================]
```

```
## [<<<<            LoadGeneSets START                  >>>>>]
```

```
## -----------------------------------------------------------
```

```
## Loading data from /mnt/hel/home/ewoewi/Stategra/Stategra_GSC/gbm/rna_mirna//20200921_npc_rna_mirna_all_gbm_correct/Disease_functions.xls
```

```
## Loading data from /mnt/hel/home/ewoewi/Stategra/Stategra_GSC/gbm/rna_mirna//20200921_npc_rna_mirna_common_gbm/Disease_functions.xls
```

```
## Loading data from /mnt/hel/home/ewoewi/Stategra/Stategra_GSC/gbm/rna_mirna//20200921_npc_rna_mirna_sig_new_genes_all_gbm/Disease_functions.xls
```

```
## Loading data from /mnt/hel/home/ewoewi/Stategra/Stategra_GSC/gbm/rna_mirna//20200921_npc_rna_mirna_sig_new_genes_gbm_corrected/Disease_functions.xls
```

```
## Loading data from /mnt/hel/home/ewoewi/Stategra/Stategra_GSC/gbm/rna_mirna//20200921_single_genes_rna_mirna_all_gbm_correct/Disease_functions.xls
```

```
## Loading IPA Functional_annotations
```

```
## -----------------------------------------------------------
```

```
## [<<<<<               LoadGeneSets END               >>>>>>]
```

```
## [=========================================================]
```

```
## [You may want to process CombineGeneSets next.            ]
```

```
## [or merge objects using MergeObjects                     ]
```

```
## [or select certain types from objects using ManageGeneSets]
```

```
RNA.Mir.GBM.Func.Obj2 <- CombineGeneSets(RNA.Mir.GBM.Func.Obj2, display = "Expanded")
```

```
## [=========================================================]
```

```
## [<<<<            CombineGeneSets START               >>>>>]
```

```
## -----------------------------------------------------------
```

```
## transforming all genes to upper case, make sure this doesnt change the data
```

```
## raw data has 46 genes
```

```
## Transformed data has 46 genes
```

```
## Combining experiments
```

```
## preparing expanded display
```

```
## calulating RR
```

```
## -----------------------------------------------------------
```

```
## [<<<<<             CombineGeneSets END              >>>>>>]
```

```
## [=========================================================]
```

```
## [You may want to process ClusterGeneSets next.            ]
```

```
RNA.Mir.GBM.Func.Obj2 <- ClusterGeneSets(RNA.Mir.GBM.Func.Obj2, clusters = 7, method = "kmeans", order = "cluster")
```

```
## [=========================================================]
```

```
## [<<<<            ClusterGeneSets START               >>>>>]
```

```
## -----------------------------------------------------------
```

```
## Using all Gene sets
```

```
## Running kmeans
```

```
## Ordering pathway clusters
```

```
## -----------------------------------------------------------
```

```
## [<<<<<             ClusterGeneSets END              >>>>>>]
```

```
## [=========================================================]
```

```
## [You may want to process HighlightGeneSets next.            ]
```

```
## [You may want to plot the results using PlotGeneSets next. ]
```

```
x <- RNA.Mir.GBM.Func.Obj2@plot$aka2
x <- x[,!colnames(x) == "Group"]
RNA.Mir.GBM.Func.Obj2@plot$aka2 <- x

x <- RNA.Mir.GBM.Func.Obj2@plot$aka3
x <- x[2:length(x)]
RNA.Mir.GBM.Func.Obj2@plot$aka3 <- x
```

```
#Generating the plot
PlotGeneSets(Object = RNA.Mir.GBM.Func.Obj2, main = "RNA.Mir.GBM.Func.Obj", RR.max = 100)
```

#### 0.4 Exporting the results

```
WriteGeneSets(Object = RNA.Met.skcm.Can.Obj2, file_location ="~/Stategra/Stategra_GSC/Final_analysis/", name = "RNA.Met.skcm.Can.Obj", write = "Data")
WriteGeneSets(Object = RNA.Mir.GBM.Can.Obj2, file_location ="~/Stategra/Stategra_GSC/Final_analysis/", name = "RNA.Mir.GBM.Can.Obj", write = "Data")
WriteGeneSets(Object = RNA.Mir.skcm.Can.Obj2, file_location ="~/Stategra/Stategra_GSC/Final_analysis/", name = "RNA.Mir.skcm.Can.Obj", write = "Data")
WriteGeneSets(Object = RNA.Met.GBM.Can.Obj2, file_location ="~/Stategra/Stategra_GSC/Final_analysis/", name = "RNA.Met.GBM.Can.Obj", write = "Data")

WriteGeneSets(Object = RNA.Met.skcm.Func.Obj2, file_location ="~/Stategra/Stategra_GSC/Final_analysis/", name = "RNA.Met.skcm.Func.Obj", write = "Data")
WriteGeneSets(Object = RNA.Mir.GBM.Func.Obj2, file_location ="~/Stategra/Stategra_GSC/Final_analysis/", name = "RNA.Mir.GBM.Func.Obj", write = "Data")
WriteGeneSets(Object = RNA.Mir.skcm.Func.Obj2, file_location ="~/Stategra/Stategra_GSC/Final_analysis/", name = "RNA.Mir.skcm.Func.Obj", write = "Data")
WriteGeneSets(Object = RNA.Met.GBM.Func.Obj2, file_location ="~/Stategra/Stategra_GSC/Final_analysis/", name = "RNA.Met.GBM.Func.Obj", write = "Data")
```
