## Supplementary material for "STATegra: Multi-omics data integration - A conceptual scheme and a bioinformatics pipeline": RMarkDown and html files.: stategra_GeneSetCluster.Rmd

````
---
title: "Stategra_GeneSetCluster"
author: "Ewoud Ewing"
date: "11/13/2020"
output: 
  html_document: 
    number_sections: yes
    toc: yes
---

```{r setup, include=FALSE}
knitr::opts_chunk$set(echo = TRUE, warning=FALSE, eval=T,results = "hold")
```

## Introduction

This is an R Markdown document of the GeneSet. For more details on using GeneSetCluster see <https://github.com/TranslationalBioinformaticsUnit/GeneSetCluster>. 

Cite: Ewing, E., Planell-Picola, N., Jagodic, M. et al. GeneSetCluster: a tool for summarizing and integrating gene-set analysis results. BMC Bioinformatics 21, 443 (2020). https://doi.org/10.1186/s12859-020-03784-z


```{r Loading}

require(GeneSetCluster)
require(ggplot2)

RNA.Mir.GBM <- list.files(path = "~/Stategra/Stategra_GSC/gbm/rna_mirna/", full.names = T, recursive = T)
RNA.Met.GBM <- list.files(path = "~/Stategra/Stategra_GSC/gbm/rna_met/", full.names = T, recursive = T)
RNA.Mir.skcm <- list.files(path = "~/Stategra/Stategra_GSC/skcm/rna_mirna/", full.names = T, recursive = T)
RNA.Met.skcm <- list.files(path = "~/Stategra/Stategra_GSC/skcm/rna_met/", full.names = T, recursive = T)

RNA.Mir.GBM.Can <- RNA.Mir.GBM[grep("Canonical", RNA.Mir.GBM)]
RNA.Met.GBM.Can <- RNA.Met.GBM[grep("Canonical", RNA.Met.GBM)]
RNA.Mir.skcm.Can <- RNA.Mir.skcm[grep("Canonical", RNA.Mir.skcm)]
RNA.Met.skcm.Can <- RNA.Met.skcm[grep("Canonical", RNA.Met.skcm)]
RNA.Mir.GBM.Func <- RNA.Mir.GBM[grep("Disease_functions", RNA.Mir.GBM)]
RNA.Met.GBM.Func <- RNA.Met.GBM[grep("Disease_functions", RNA.Met.GBM)]
RNA.Mir.skcm.Func <- RNA.Mir.skcm[grep("Disease_functions", RNA.Mir.skcm)]
RNA.Met.skcm.Func <- RNA.Met.skcm[grep("Disease_functions", RNA.Met.skcm)]
```

## Analysis of Canonical Pathways
IPA give the user a lot of output, we focused on the Canonical Pathways and disease and functions, which we analysed seperatly. 

### RNA.Met.skcm.Can
We make an PathwayObject for every group seperate for Canonical Pathways and disease and functions. GeneSetCluster could combine them but then it wouldnt be as clear what the additional functional informaton comes from the OmicsNPC. 

```{r RNA.Met.skcm.Can}
groups <- limma::strsplit2(RNA.Met.skcm.Can, split="/")[,11]
groups <- gsub(pattern = "20200921_", replacement = "", groups)

RNA.Met.skcm.Can.Obj2<- LoadGeneSets(file_location = RNA.Met.skcm.Can[2:4], 
                                     groupnames = groups[2:4], P.cutoff = 1.31, Mol.cutoff =3, 
                                     Source = "IPA", type = "Canonical_Pathways", 
                                     structure = "SYMBOL", Organism = "org.Hs.eg.db", seperator = ",")
#There is a build in loader for IPA and GREAT datasets

RNA.Met.skcm.Can.Obj2 <- CombineGeneSets(RNA.Met.skcm.Can.Obj2,combineMethod ="Standard",  display = "Expanded")
#Here we calculate the relative risk between the different Gene Sets

#OptimalGeneSets(RNA.Met.skcm.Can.Obj, method = "gap",cluster_method = "kmeans", max_cluster = 10, main="RNA.Met.skcm.Can.Obj")
#Calculate the optimal number of clusters

RNA.Met.skcm.Can.Obj2 <- ClusterGeneSets(RNA.Met.skcm.Can.Obj2, clusters = 6, method = "kmeans", order = "cluster")
#Here we cluster using Kmeans in the 6 clusters determined by OptimalGeneSets


#Because of how we want the plots to look, we remove the group information so its just the pathways where they appear.
x <- RNA.Met.skcm.Can.Obj2@plot$aka2
x <- x[,!colnames(x) == "Group"]
RNA.Met.skcm.Can.Obj2@plot$aka2 <- x

x <- RNA.Met.skcm.Can.Obj2@plot$aka3
x <- x[2:length(x)]
RNA.Met.skcm.Can.Obj2@plot$aka3 <- x

```

```{r fig1, fig.height = 10, fig.width = 15, fig.align = "center"}
#Generating the plot
PlotGeneSets(Object = RNA.Met.skcm.Can.Obj2, main = "RNA.Met.skcm.Can.Obj", RR.max = 50)
```


### RNA.Mir.skcm.Can


```{r RNA.Mir.skcm.Can}
groups <- limma::strsplit2(RNA.Mir.skcm.Can, split="/")[,11]
groups <- gsub(pattern = "20200921_", replacement = "", groups)
RNA.Mir.skcm.Can.Obj2 <- LoadGeneSets(file_location = RNA.Mir.skcm.Can[2:4], 
                                     groupnames = groups[2:4], P.cutoff = 1.31, Mol.cutoff =3, 
                                     Source = "IPA", type = "Canonical_Pathways", 
                                     structure = "SYMBOL", Organism = "org.Hs.eg.db", seperator = ",")

RNA.Mir.skcm.Can.Obj2 <- CombineGeneSets(RNA.Mir.skcm.Can.Obj2, display = "Expanded")
#OptimalGeneSets(RNA.Mir.skcm.Can.Obj, method = "gap",cluster_method = "kmeans", max_cluster = 10, main="RNA.Mir.skcm.Can.Obj")

RNA.Mir.skcm.Can.Obj2 <- ClusterGeneSets(RNA.Mir.skcm.Can.Obj2, clusters = 7, method = "kmeans", order = "cluster")

x <- RNA.Mir.skcm.Can.Obj2@plot$aka2
x <- x[,!colnames(x) == "Group"]
RNA.Mir.skcm.Can.Obj2@plot$aka2 <- x

x <- RNA.Mir.skcm.Can.Obj2@plot$aka3
x <- x[2:length(x)]
RNA.Mir.skcm.Can.Obj2@plot$aka3 <- x

PlotGeneSets(Object = RNA.Mir.skcm.Can.Obj2, main = "RNA.Mir.skcm.Can.Obj", RR.max = 50)
```

```{r fig2, fig.height = 10, fig.width = 15, fig.align = "center"}
#Generating the plot
PlotGeneSets(Object = RNA.Mir.skcm.Can.Obj2, main = "RNA.Mir.skcm.Can.Obj", RR.max = 50)
```


### RNA.Met.GBM.Can


```{r RNA.Met.GBM.Can}
groups <- limma::strsplit2(RNA.Met.GBM.Can, split="/")[,11]
groups <- gsub(pattern = "20200921_", replacement = "", groups)
groups <- gsub(pattern = "_Correct", replacement = "", groups)

RNA.Met.GBM.Can.Obj2 <- LoadGeneSets(file_location = RNA.Met.GBM.Can[c(1,2,4,6)], 
                                    groupnames = groups[c(1,2,4,6)], P.cutoff = 1.31, Mol.cutoff =3, 
                                    Source = "IPA", type = "Canonical_Pathways", 
                                    structure = "SYMBOL", Organism = "org.Hs.eg.db", seperator = ",")

RNA.Met.GBM.Can.Obj2 <-CombineGeneSets(RNA.Met.GBM.Can.Obj2, display = "Expanded")

RNA.Met.GBM.Can.Obj2 <- ClusterGeneSets(RNA.Met.GBM.Can.Obj2, clusters = 4, method = "kmeans", order = "cluster")

x <- RNA.Met.GBM.Can.Obj2@plot$aka2
x <- x[,!colnames(x) == "Group"]
RNA.Met.GBM.Can.Obj2@plot$aka2 <- x

x <- RNA.Met.GBM.Can.Obj2@plot$aka3
x <- x[2:length(x)]
RNA.Met.GBM.Can.Obj2@plot$aka3 <- x


```

```{r fig3, fig.height = 10, fig.width = 15, fig.align = "center"}
#Generating the plot
PlotGeneSets(Object = RNA.Met.GBM.Can.Obj2, main = "RNA.Met.GBM.Can.Obj", RR.max = 50)
```


### RNA.Mir.GBM.Can


```{r RNA.Mir.GBM.Can}
groups <- limma::strsplit2(RNA.Mir.GBM.Can, split="/")[,11]
groups <- gsub(pattern = "20200921_", replacement = "", groups)
#groups <- gsub(pattern = "_correct", replacement = "", groups)

idx <- c(1,4)
RNA.Mir.GBM.Can.Obj2 <- LoadGeneSets(file_location = RNA.Mir.GBM.Can[idx], 
                                    groupnames = groups[idx], P.cutoff = 1.31, Mol.cutoff =3, 
                                    Source = "IPA", type = "Canonical_Pathways", 
                                    structure = "SYMBOL", Organism = "org.Hs.eg.db", seperator = ",")

RNA.Mir.GBM.Can.Obj2 <- CombineGeneSets(RNA.Mir.GBM.Can.Obj2, display = "Expanded")

RNA.Mir.GBM.Can.Obj2 <- ClusterGeneSets(RNA.Mir.GBM.Can.Obj2, clusters = 2, method = "kmeans", order = "cluster")

x <- RNA.Mir.GBM.Can.Obj2@plot$aka2
x <- x[,!colnames(x) == "Group"]
RNA.Mir.GBM.Can.Obj2@plot$aka2 <- x

x <- RNA.Mir.GBM.Can.Obj2@plot$aka3
x <- x[2:length(x)]
RNA.Mir.GBM.Can.Obj2@plot$aka3 <- x


PlotGeneSets(Object = RNA.Mir.GBM.Can.Obj2, main = "RNA.Mir.GBM.Can.Obj", RR.max = 50)

```

```{r fig4, fig.height = 10, fig.width = 15, fig.align = "center"}
#Generating the plot
PlotGeneSets(Object = RNA.Mir.GBM.Can.Obj2, main = "RNA.Mir.GBM.Can.Obj", RR.max = 50)
```

## Analysis of Disease and Functions
Here we are analysing the Disease and Functions output from IPA.


### RNA.Met.skcm.Func


```{r RNA.Met.skcm.Func}
groups <- limma::strsplit2(RNA.Met.skcm.Func, split="/")[,11]
groups <- gsub(pattern = "20200921_", replacement = "", groups)

idx <- c(1,3,4,5)

RNA.Met.skcm.Func.Obj2 <- LoadGeneSets(file_location = RNA.Met.skcm.Func[idx], 
                                      groupnames = groups[idx], P.cutoff = 0.05, Mol.cutoff =3, 
                                      Source = "IPA", type = "Functional_annotations", 
                                      structure = "SYMBOL", Organism = "org.Hs.eg.db", seperator = ",")

RNA.Met.skcm.Func.Obj2 <- CombineGeneSets(RNA.Met.skcm.Func.Obj2, display = "Expanded")
RNA.Met.skcm.Func.Obj2 <- ClusterGeneSets(RNA.Met.skcm.Func.Obj2, clusters = 4, method = "kmeans", order = "cluster")

x <- RNA.Met.skcm.Func.Obj2@plot$aka2
x <- x[,!colnames(x) == "Group"]
RNA.Met.skcm.Func.Obj2@plot$aka2 <- x

x <- RNA.Met.skcm.Func.Obj2@plot$aka3
x <- x[2:length(x)]
RNA.Met.skcm.Func.Obj2@plot$aka3 <- x

```

```{r fig5, fig.height = 10, fig.width = 15, fig.align = "center"}
#Generating the plot
PlotGeneSets(Object = RNA.Met.skcm.Func.Obj2, main = "RNA.Met.skcm.Func.Obj")
```

### RNA.Mir.skcm.Func


```{r RNA.Mir.skcm.Func}
groups <- limma::strsplit2(RNA.Mir.skcm.Func, split="/")[,11]
groups <- gsub(pattern = "20200921_", replacement = "", groups)

idx <- c(1,3:5)

RNA.Mir.skcm.Func.Obj2 <- LoadGeneSets(file_location = RNA.Mir.skcm.Func[idx], 
                                      groupnames = groups[idx], P.cutoff = 0.05, Mol.cutoff =3, 
                                      Source = "IPA", type = "Functional_annotations", 
                                      structure = "SYMBOL", Organism = "org.Hs.eg.db", seperator = ",")

RNA.Mir.skcm.Func.Obj2 <- CombineGeneSets(RNA.Mir.skcm.Func.Obj2, display = "Expanded")


RNA.Mir.skcm.Func.Obj2 <- ClusterGeneSets(RNA.Mir.skcm.Func.Obj2, clusters = 7, method = "kmeans", order = "cluster")

x <- RNA.Mir.skcm.Func.Obj2@plot$aka2
x <- x[,!colnames(x) == "Group"]
RNA.Mir.skcm.Func.Obj2@plot$aka2 <- x

x <- RNA.Mir.skcm.Func.Obj2@plot$aka3
x <- x[2:length(x)]
RNA.Mir.skcm.Func.Obj2@plot$aka3 <- x

```

```{r fig6, fig.height = 10, fig.width = 15, fig.align = "center"}
#Generating the plot
PlotGeneSets(Object = RNA.Mir.skcm.Func.Obj2, main = "RNA.Mir.skcm.Func.Obj", RR.max = 75)
```

### RNA.Met.GBM.Func


```{r RNA.Met.GBM.Func}
groups <- limma::strsplit2(RNA.Met.GBM.Func, split="/")[,11]
groups <- gsub(pattern = "20200921_", replacement = "", groups)
groups <- gsub(pattern = "_Correct", replacement = "", groups)

idx <- c(1,2,3,5)

RNA.Met.GBM.Func.Obj2 <- LoadGeneSets(file_location = RNA.Met.GBM.Func[idx], 
                                     groupnames = groups[idx], P.cutoff = 0.05, Mol.cutoff =3, 
                                     Source = "IPA", type = "Functional_annotations", 
                                     structure = "SYMBOL", Organism = "org.Hs.eg.db", seperator = ",")

RNA.Met.GBM.Func.Obj2 <- CombineGeneSets(RNA.Met.GBM.Func.Obj2, display = "Expanded")

#RNA.Met.GBM.Func.Obj2 <- ClusterGeneSets(RNA.Met.GBM.Func.Obj2, clusters = 5, method = "kmeans")

RNA.Met.GBM.Func.Obj2 <- ClusterGeneSets(RNA.Met.GBM.Func.Obj2, clusters = 5, method = "kmeans", order = "cluster")

x <- RNA.Met.GBM.Func.Obj2@plot$aka2
x <- x[,!colnames(x) == "Group"]
RNA.Met.GBM.Func.Obj2@plot$aka2 <- x

x <- RNA.Met.GBM.Func.Obj2@plot$aka3
x <- x[2:length(x)]
RNA.Met.GBM.Func.Obj2@plot$aka3 <- x


```

```{r fig7, fig.height = 10, fig.width = 15, fig.align = "center"}
#Generating the plot
PlotGeneSets(Object = RNA.Met.GBM.Func.Obj2, main = "RNA.Met.GBM.Func.Obj", RR.max = 50)
```

### RNA.Mir.GBM.Func


```{r RNA.Mir.GBM.Func}
groups <- limma::strsplit2(RNA.Mir.GBM.Func, split="/")[,11]
groups <- gsub(pattern = "20200921_", replacement = "", groups)
groups <- gsub(pattern = "_Correct", replacement = "", groups)

idx <- c(1,2,4,5,7)

RNA.Mir.GBM.Func.Obj2 <- LoadGeneSets(file_location = RNA.Mir.GBM.Func[idx], 
                                     groupnames = groups[idx], P.cutoff = 0.05, Mol.cutoff =3, 
                                     Source = "IPA", type = "Functional_annotations", 
                                     structure = "SYMBOL", Organism = "org.Hs.eg.db", seperator = ",")

RNA.Mir.GBM.Func.Obj2 <- CombineGeneSets(RNA.Mir.GBM.Func.Obj2, display = "Expanded")

RNA.Mir.GBM.Func.Obj2 <- ClusterGeneSets(RNA.Mir.GBM.Func.Obj2, clusters = 7, method = "kmeans", order = "cluster")

x <- RNA.Mir.GBM.Func.Obj2@plot$aka2
x <- x[,!colnames(x) == "Group"]
RNA.Mir.GBM.Func.Obj2@plot$aka2 <- x

x <- RNA.Mir.GBM.Func.Obj2@plot$aka3
x <- x[2:length(x)]
RNA.Mir.GBM.Func.Obj2@plot$aka3 <- x


```

```{r fig8, fig.height = 10, fig.width = 15, fig.align = "center"}
#Generating the plot
PlotGeneSets(Object = RNA.Mir.GBM.Func.Obj2, main = "RNA.Mir.GBM.Func.Obj", RR.max = 100)
```


##Exporting the results

```{r exporting the data}

WriteGeneSets(Object = RNA.Met.skcm.Can.Obj2, file_location ="~/Stategra/Stategra_GSC/Final_analysis/", name = "RNA.Met.skcm.Can.Obj", write = "Data")
WriteGeneSets(Object = RNA.Mir.GBM.Can.Obj2, file_location ="~/Stategra/Stategra_GSC/Final_analysis/", name = "RNA.Mir.GBM.Can.Obj", write = "Data")
WriteGeneSets(Object = RNA.Mir.skcm.Can.Obj2, file_location ="~/Stategra/Stategra_GSC/Final_analysis/", name = "RNA.Mir.skcm.Can.Obj", write = "Data")
WriteGeneSets(Object = RNA.Met.GBM.Can.Obj2, file_location ="~/Stategra/Stategra_GSC/Final_analysis/", name = "RNA.Met.GBM.Can.Obj", write = "Data")

WriteGeneSets(Object = RNA.Met.skcm.Func.Obj2, file_location ="~/Stategra/Stategra_GSC/Final_analysis/", name = "RNA.Met.skcm.Func.Obj", write = "Data")
WriteGeneSets(Object = RNA.Mir.GBM.Func.Obj2, file_location ="~/Stategra/Stategra_GSC/Final_analysis/", name = "RNA.Mir.GBM.Func.Obj", write = "Data")
WriteGeneSets(Object = RNA.Mir.skcm.Func.Obj2, file_location ="~/Stategra/Stategra_GSC/Final_analysis/", name = "RNA.Mir.skcm.Func.Obj", write = "Data")
WriteGeneSets(Object = RNA.Met.GBM.Func.Obj2, file_location ="~/Stategra/Stategra_GSC/Final_analysis/", name = "RNA.Met.GBM.Func.Obj", write = "Data")


```
````
