## Supplementary material for "STATegra: Multi-omics data integration - A conceptual scheme and a bioinformatics pipeline": RMarkDown and html files.: stategra_skcm_pipeline.html

Omics integraive pipeline: STATegRa


### Omics integraive pipeline: STATegRa

Translational Bioinformatics Unit (Navarrabiomed)

#### 2020-11-02

### Contents

- 1 Introduction - Skin Cutaneous Melanoma
- 2 Skin Cutaneous Melanoma
  - 2.1 Initial data exploration and pre-process
    - 2.1.1 Metadata exploration
    - 2.1.2 Individual omics data-type exploration
      - 2.1.2.1 Expression data (`rna_skcm`)
      - 2.1.2.2 miRNA data (`mirna_skcm`)
      - 2.1.2.3 Methylation data (`met_skcm`)
    - 2.1.3 Joint omics exploration
      - 2.1.3.1 Common samples
      - 2.1.3.2 OmicsPCA exploration
      - 2.1.3.3 Model selection
        - 2.1.3.3.1 JIVE
        - 2.1.3.3.2 PCA-GCA
        - 2.1.3.3.3 pESCA
      - 2.1.3.4 Sub-space recovery
    - 2.1.4 Batch correction
      - 2.1.4.1 Expression data (`log2_rna_skcm_corrected`)
      - 2.1.4.2 miRNA data (`log2_mirna_skcm_corrected`)
      - 2.1.4.3 Methylation data (`logit_met_skcm`)
  - 2.2 Characterization of the data: Individual exploration.
    - 2.2.1 Expression data (`subset_log2_rna_skcm_corrected`)
    - 2.2.2 miRNA data (`subset_log2_mirna_skcm_corrected`)
    - 2.2.3 Methylation data (`subset_logit_met_skcm_corrected`)
  - 2.3 Characterization of the data: Joint exploration.
    - 2.3.1 Data scaling
    - 2.3.2 Model selection
      - 2.3.2.1 JIVE
      - 2.3.2.2 PCA-GCA
      - 2.3.2.3 pESCA
    - 2.3.3 Sub-space recovery
  - 2.4 Integrative differential analysis by NPC
    - 2.4.1 mRNA + miRNA
      - 2.4.1.1 Mapping file
      - 2.4.1.2 Parametric Combination
      - 2.4.1.3 Non Parametric Combination
      - 2.4.1.4 Integration strategies comparison
    - 2.4.2 mRNA + Methylation
      - 2.4.2.1 Mapping file
      - 2.4.2.2 Parametric Combination
      - 2.4.2.3 Non Parametric Combination
      - 2.4.2.4 Integration strategies comparison

### 1 Introduction - Skin Cutaneous Melanoma

Following the examples above we aim to provide an integrative workflow of omics data by combining existing tools implemented in the `STATegRa` package.

To this aim, the Skin Cutaneous Melanoma (SKCM) dataset from The Cancer Genome Atlas (TCGA) was explored.

The level 3 public data for gene expression, miRNA and methylation were obtained through the NCI’s Genomic Data Commons (GDC) portal (Tomczak, Czerwinska, & Wiznerowicz, 2015).

```
load('20190108_1_skcm_raw_data.RData', verbose=TRUE)
```

```
## Loading objects:
##   clinical_skcm
##   met_skcm
##   mirna_skcm
##   publication_metadata
##   rna_skcm
```

### 2 Skin Cutaneous Melanoma

The TCGA-SKCM dataset consists in melanoma samples from patients diagnoses of either primary or metastatic cutaneous melanoma or metastatic melanoma of unknown primary from roughly 400 cases.

The metadata associated to this project is stored in the `clinical_skcm` R object.

```
dim(clinical_skcm)
```

```
## [1] 428  22
```

A description of the variables collected as metadata within the TCGA context is found in the NCBI’s GDC Documentation - Data Dictionary.

Different omics were produced from the collected samples. For this example we will focuse on three: mRNA (`rna_skcm`), miRNA (`mirna_skcm`) and Methylation (`met_skcm`).

```
dim(rna_skcm)
```

```
## [1] 20531   428
```

```
dim(mirna_skcm)
```

```
## [1] 1046  428
```

```
dim(met_skcm)
```

```
## [1] 485577    428
```

Moreover, we will used the curated metadata from the melanoma publication in Cell (Akbani, Rehan, et al., 2015).

```
dim(publication_metadata)
```

```
## [1] 333 114
```

Worth highlighting that the number of cases used in previous publication are different from the actual number of cases available in the TCGA (333 vs 428).

#### 2.1 Initial data exploration and pre-process

Before start the integration analysis approach, an exploration of each omic data-type and metadata is required.

##### 2.1.1 Metadata exploration

Metadata provided by TCGA contains information about demographic features (age, gender, race, ethnicity,etc.), tumour characteristics (age at diagnosis, primary site of disease, neoplasm disease stage, ulceration in melanoma, Breslow thickness,etc.) and technical processes (such as batch).

```
names(clinical_skcm)
```

```
##  [1] "Composite Element REF"                 
##  [2] "years_to_birth"                        
##  [3] "vital_status"                          
##  [4] "days_to_death"                         
##  [5] "days_to_last_followup"                 
##  [6] "days_to_submitted_specimen_dx"         
##  [7] "primary_site_of_disease"               
##  [8] "neoplasm_diseasestage"                 
##  [9] "pathology_T_stage"                     
## [10] "pathology_N_stage"                     
## [11] "pathology_M_stage"                     
## [12] "melanoma_ulceration"                   
## [13] "melanoma_primary_known"                
## [14] "Breslow_thickness"                     
## [15] "dcc_upload_date"                       
## [16] "gender"                                
## [17] "date_of_initial_pathologic_diagnosis"  
## [18] "radiation_therapy"                     
## [19] "radiations_radiation_regimenindication"
## [20] "race"                                  
## [21] "ethnicity"                             
## [22] "batch_number"
```

Additionally we define the survival outcome:

```
clinical_skcm$survivalTime <- pmin(clinical_skcm$days_to_death,
                                  clinical_skcm$days_to_last_followup, na.rm = TRUE)

clinical_skcm$survivalOutcome <- Surv(time = clinical_skcm$survivalTime,
                                     event = clinical_skcm$vital_status == '1')
```

Exploring the data three replicates were identified and removed

```
rep_samples <- grep("\\.1",rownames(clinical_skcm))
rownames(clinical_skcm)[rep_samples]
```

```
## [1] "TCGA-ER-A2NF.1" "TCGA-ER-A19T.1" "TCGA-GN-A4U8.1"
```

```
clinical_skcm <- clinical_skcm[-rep_samples,]
```

Within the curated metadata from the TCGA Cell publication on Melanoma Project, additional information for a subset of 333 cases is available.

```
names(publication_metadata)
```

```
##   [1] "Name"                                                                                    
##   [2] "Name2"                                                                                   
##   [3] "ALL_SAMPLES"                                                                             
##   [4] "MUTATIONSUBTYPES"                                                                        
##   [5] "ALL_PRIMARY_VS_METASTATIC"                                                               
##   [6] "REGIONAL_VS_PRIMARY"                                                                     
##   [7] "UV.signature"                                                                            
##   [8] "RNASEQ.CLUSTER_CONSENHIER"                                                               
##   [9] "MethTypes.201408"                                                                        
##  [10] "MIRCluster"                                                                              
##  [11] "ProteinCluster"                                                                          
##  [12] "OncoSignCluster"                                                                         
##  [13] "CDKN2A.cg13601799._meth"                                                                 
##  [14] "KIT.cg10087973._meth"                                                                    
##  [15] "BRAF_cna"                                                                                
##  [16] "NRAS_cna"                                                                                
##  [17] "MITF_cna"                                                                                
##  [18] "KIT_cna"                                                                                 
##  [19] "PDGFRA_cna"                                                                              
##  [20] "KDR_cna"                                                                                 
##  [21] "MYC_cna"                                                                                 
##  [22] "CCND1_cna"                                                                               
##  [23] "CDK4_cna"                                                                                
##  [24] "MDM2_cna"                                                                                
##  [25] "TERT_cna"                                                                                
##  [26] "EP300_cna"                                                                               
##  [27] "NOTCH2_cna"                                                                              
##  [28] "CDKN2A_cna"                                                                              
##  [29] "PTEN_cna"                                                                                
##  [30] "TP53_cna"                                                                                
##  [31] "PPP6C_cna"                                                                               
##  [32] "ARID2_cna"                                                                               
##  [33] "IDH1_cna"                                                                                
##  [34] "MAP2K1_cna"                                                                              
##  [35] "DDX3X_cna"                                                                               
##  [36] "RAC1_cna"                                                                                
##  [37] "NF1_cna"                                                                                 
##  [38] "CASP8_cna"                                                                               
##  [39] "CTNNB1_cna"                                                                              
##  [40] "PCDHGA1_cna"                                                                             
##  [41] "SERPINB3_cna"                                                                            
##  [42] "IRF7_cna"                                                                                
##  [43] "HRAS_cna"                                                                                
##  [44] "PTPN11_cna"                                                                              
##  [45] "KRAS_cna"                                                                                
##  [46] "GNA11_cna"                                                                               
##  [47] "GNAQ_cna"                                                                                
##  [48] "AKT1_cna"                                                                                
##  [49] "APC_cna"                                                                                 
##  [50] "EZH2_cna"                                                                                
##  [51] "FBXW7_cna"                                                                               
##  [52] "GNAS_cna"                                                                                
##  [53] "PIK3CA_cna"                                                                              
##  [54] "ARID2_mut"                                                                               
##  [55] "BRAF_mut"                                                                                
##  [56] "NRAS_mut"                                                                                
##  [57] "KRAS_mut"                                                                                
##  [58] "HRAS_mut"                                                                                
##  [59] "NF1_mut"                                                                                 
##  [60] "IDH1_mut"                                                                                
##  [61] "GNA11_mut"                                                                               
##  [62] "GNAQ_mut"                                                                                
##  [63] "CDK4_mut"                                                                                
##  [64] "CDKN2A_mut"                                                                              
##  [65] "DDX3X_mut"                                                                               
##  [66] "KIT_mut"                                                                                 
##  [67] "MAP2K1_mut"                                                                              
##  [68] "PPP6C_mut"                                                                               
##  [69] "PTEN_mut"                                                                                
##  [70] "RAC1_mut"                                                                                
##  [71] "RB1_mut"                                                                                 
##  [72] "TP53_mut"                                                                                
##  [73] "TERT_MUT"                                                                                
##  [74] "rnaseq_TERT.RSEM.NORM_DATA"                                                              
##  [75] "rppa_LCK.LCK.R.V"                                                                        
##  [76] "rppa_SYK.SYK.M.V"                                                                        
##  [77] "PIGMENT.SCORE"                                                                           
##  [78] "NECROSIS"                                                                                
##  [79] "Tumour.content........nuceli.that.are.tumour.cells......0.100.."                         
##  [80] "LYMPHOCYTE.DISTRIBUTION"                                                                 
##  [81] "LYMPHOCYTE.DENSITY"                                                                      
##  [82] "LYMPHOCYTE.SCORE"                                                                        
##  [83] "PURITY..ABSOLUTE."                                                                       
##  [84] "PLOIDY..ABSOLUTE."                                                                       
##  [85] "TOTAL.MUTATIONS"                                                                         
##  [86] "UV.RATE"                                                                                 
##  [87] "CURATED_AGE_AT_INITIAL_PATHOLOGIC_DIAGNOSIS"                                             
##  [88] "CURATED_AGE_AT_TCGA_SPECIMEN"                                                            
##  [89] "GENDER"                                                                                  
##  [90] "CURATED_KNOWN_PRIMARY"                                                                   
##  [91] "CURATED_SITE_OF_PRIMARY_TUMOR_KNOWN_PRIMARY_ONLY"                                        
##  [92] "CURATED_SITE_OF_PRIMARY_TUMOR_INCLUDING_UNKNOWN"                                         
##  [93] "CURATED_BRESLOW"                                                                         
##  [94] "CURATED_ULCERATION"                                                                      
##  [95] "CURATED_T_STAGE_AT_DIAGNOSIS_COMPLEX"                                                    
##  [96] "CURATED_T_STAGE_AT_DIAGNOSIS_SIMPLE"                                                     
##  [97] "CURATED_N_STAGE_AT_DIAGNOSIS_COMPLEX"                                                    
##  [98] "CURATED_N_STAGE_AT_DIAGNOSIS_SIMPLE"                                                     
##  [99] "CURATED_M_STAGE_AT_DIAGNOSIS_COMPLEX"                                                    
## [100] "CURATED_M_STAGE_AT_DIAGNOSIS_SIMPLE"                                                     
## [101] "CURATED_PATHOLOGIC_STAGE_AJCC7_AT_DIAGNOSIS_COMPLEX"                                     
## [102] "CURATED_PATHOLOGIC_STAGE_AJCC7_AT_DIAGNOSIS_COMPLEX_NP"                                  
## [103] "CURATED_PATHOLOGIC_STAGE_AJCC7_AT_DIAGNOSIS_SIMPLE"                                      
## [104] "CURATED_TCGA_SPECIMEN_SITE"                                                              
## [105] "CURATED_DISTANT_ANATOMIC_SITE"                                                           
## [106] "CURATED_VITAL_STATUS"                                                                    
## [107] "CURATED_DAYS_TO_DEATH_OR_LAST_FU"                                                        
## [108] "CURATED_TCGA_DAYS_TO_DEATH_OR_LAST_FU"                                                   
## [109] "CURATED_MELANOMA_SPECIFIC_VITAL_STATUS..0....ALIVE.OR.CENSORED...1....DEAD.OF.MELANOMA.."
## [110] "CURATED_TCGA_SPECIMEN_Distant"                                                           
## [111] "CC.TT.nTotal.Mut"                                                                        
## [112] "DIPYRIM.C.T.nTotal.Mut"                                                                  
## [113] "DIPYRIM.C.T.n.C.T..mut"                                                                  
## [114] "SHATTERSEEK_Chromothripsis_calls"
```

If we explore this data we can see that the age of TCGA specimen and age at diagnosis differs, conditioning the approaches that can be done with the data generated (all specimen information is from diagnostic time).

```
plot(publication_metadata$CURATED_AGE_AT_INITIAL_PATHOLOGIC_DIAGNOSIS,
     publication_metadata$CURATED_AGE_AT_TCGA_SPECIMEN, 
     xlab="age at initial pathologic diagnosis (years)", 
     ylab="age at TCGA specimen (years)")

abline(0,1, col="red")
```

Let’s explore how is the distribution of the time window between the initial pathologic diagnosis and sample collection

```
diff_dx_collection <- publication_metadata$CURATED_AGE_AT_TCGA_SPECIMEN - publication_metadata$CURATED_AGE_AT_INITIAL_PATHOLOGIC_DIAGNOSIS

hist(diff_dx_collection)
```

```
summary(diff_dx_collection)
```

```
##    Min. 1st Qu.  Median    Mean 3rd Qu.    Max.    NA's 
##   0.000   1.000   2.000   3.628   5.000  29.000      45
```

Based on the exploration we can see that the median time window between specimen date and diagnostic date is two years after diagnosis (N=165).
In order to be consistant we will work with those samples taken close to diagnosis (maximum 1 years between diagnosis and sample collection).

```
publication_metadata$Name2[which(diff_dx_collection<=1)]
```

```
##   [1] TCGA-BF-A1PV TCGA-BF-A1Q0 TCGA-D3-A1Q1 TCGA-D3-A1Q3 TCGA-D3-A1Q4
##   [6] TCGA-D3-A1Q6 TCGA-D3-A1Q8 TCGA-D3-A1Q9 TCGA-D3-A2J9 TCGA-D3-A2JC
##  [11] TCGA-D3-A2JD TCGA-D3-A2JH TCGA-D3-A2JK TCGA-D3-A2JN TCGA-D3-A2JO
##  [16] TCGA-D3-A2JP TCGA-D3-A3C7 TCGA-D3-A3C8 TCGA-D3-A3CF TCGA-D3-A3ML
##  [21] TCGA-D3-A3MO TCGA-D3-A3MR TCGA-D3-A3MU TCGA-D3-A51F TCGA-D3-A51K
##  [26] TCGA-D3-A51N TCGA-D3-A51T TCGA-D3-A5GS TCGA-D9-A1JW TCGA-D9-A1JX
##  [31] TCGA-D9-A3Z1 TCGA-D9-A3Z3 TCGA-D9-A3Z4 TCGA-D9-A4Z2 TCGA-D9-A4Z3
##  [36] TCGA-D9-A4Z5 TCGA-DA-A1HV TCGA-DA-A1HW TCGA-DA-A1I0 TCGA-DA-A1IB
##  [41] TCGA-DA-A3F3 TCGA-DA-A3F8 TCGA-EB-A1NK TCGA-EB-A24C TCGA-EB-A24D
##  [46] TCGA-EB-A299 TCGA-EB-A3HV TCGA-EB-A3XB TCGA-EB-A3XC TCGA-EB-A3XD
##  [51] TCGA-EB-A3XE TCGA-EB-A3XF TCGA-EB-A3Y6 TCGA-EB-A3Y7 TCGA-EB-A41A
##  [56] TCGA-EB-A41B TCGA-EB-A42Y TCGA-EB-A42Z TCGA-EB-A431 TCGA-EB-A44N
##  [61] TCGA-EB-A44O TCGA-EB-A44P TCGA-EB-A44Q TCGA-EB-A4IQ TCGA-EB-A4XL
##  [66] TCGA-EB-A553 TCGA-EB-A5FP TCGA-EE-A17Z TCGA-EE-A182 TCGA-EE-A185
##  [71] TCGA-EE-A20I TCGA-EE-A29C TCGA-EE-A29D TCGA-EE-A29E TCGA-EE-A29L
##  [76] TCGA-EE-A29R TCGA-EE-A29X TCGA-EE-A2A2 TCGA-EE-A2GB TCGA-EE-A2GP
##  [81] TCGA-EE-A2GR TCGA-EE-A2M5 TCGA-EE-A2M8 TCGA-EE-A2MH TCGA-EE-A3AA
##  [86] TCGA-EE-A3AB TCGA-EE-A3AC TCGA-EE-A3AD TCGA-EE-A3AF TCGA-EE-A3AG
##  [91] TCGA-ER-A193 TCGA-ER-A194 TCGA-ER-A196 TCGA-ER-A197 TCGA-ER-A198
##  [96] TCGA-ER-A199 TCGA-ER-A19A TCGA-ER-A19D TCGA-ER-A19E TCGA-ER-A19J
## [101] TCGA-ER-A19K TCGA-ER-A19N TCGA-ER-A19O TCGA-ER-A19S TCGA-ER-A1A1
## [106] TCGA-ER-A2NB TCGA-ER-A2NF TCGA-FR-A2OS TCGA-FR-A3R1 TCGA-FS-A1Z3
## [111] TCGA-FS-A1Z7 TCGA-FS-A1ZF TCGA-FS-A1ZG TCGA-FS-A1ZH TCGA-FS-A1ZK
## [116] TCGA-FS-A1ZN TCGA-FS-A1ZR TCGA-FS-A4F5 TCGA-FS-A4F9 TCGA-FW-A3I3
## [121] TCGA-FW-A3TV TCGA-FW-A5DX TCGA-GF-A2C7 TCGA-GF-A3OT TCGA-GF-A4EO
## [126] TCGA-GN-A263 TCGA-GN-A265 TCGA-GN-A266 TCGA-GN-A269 TCGA-GN-A26C
## [131] TCGA-IH-A3EA
## 331 Levels: TCGA-BF-A1PU TCGA-BF-A1PV TCGA-BF-A1PX TCGA-BF-A1PZ ... TCGA-IH-A3EA
```

Now, we must check if these 131 samples have omics data available.

```
target_data <- publication_metadata[which(diff_dx_collection<=1),]
sum(target_data$Name2%in%rownames(clinical_skcm))
```

```
## [1] 128
```

We have omics information for 128 samples from the 131.
Let’s see the characteristics of these 128 cases candidates for forward analysis.

```
target_data2 <- target_data[target_data$Name2%in%rownames(clinical_skcm),]
dim(target_data2)
```

```
## [1] 128 114
```

```
summary(target_data2$CURATED_TCGA_SPECIMEN_SITE)
```

```
##                            -              [Not Available] 
##                            0                            2 
##           Distant Metastasis                Primary Tumor 
##                            5                           41 
##          Regional Lymph Node Regional Skin or Soft Tissue 
##                           63                           17
```

As the remaining number of samples from distant metastasis and regional skin or soft tissue is very limited, we will work only with primary tumor and regional LN.

```
target_data3 <- target_data2[which(target_data2$CURATED_TCGA_SPECIMEN_SITE=="Primary Tumor" | target_data2$CURATED_TCGA_SPECIMEN_SITE=="Regional Lymph Node"),]

dim(target_data3)
```

```
## [1] 104 114
```

```
target_data3$CURATED_TCGA_SPECIMEN_SITE <- droplevels(target_data3$CURATED_TCGA_SPECIMEN_SITE)

#merge the two clinical data bases
clinical_skcm2 <- merge(clinical_skcm, target_data3, by.x=0, by.y="Name2", sort=FALSE)
rownames(clinical_skcm2) <- clinical_skcm2$Row.names

clinical_skcm2$stage <- clinical_skcm2[,"CURATED_PATHOLOGIC_STAGE_AJCC7_AT_DIAGNOSIS_COMPLEX_NP"]
clinical_skcm2$stage_simplified <- clinical_skcm2[,"CURATED_PATHOLOGIC_STAGE_AJCC7_AT_DIAGNOSIS_SIMPLE"]
```

Then, based on survival information we can calculate the survival curve

```
# Load require packages
library(survival)
library(ggplot2)
library(ggfortify)

clinical_skcm2$survivalTime <- pmin(clinical_skcm2$days_to_death,
                                  clinical_skcm2$days_to_last_followup, na.rm = TRUE)

clinical_skcm2$survivalOutcome <- Surv(time = clinical_skcm2$survivalTime,
                                     event = clinical_skcm2$vital_status == '1')

# Plot survival curve
my.fit <- survfit(clinical_skcm2$survivalOutcome ~ 1)
autoplot(my.fit, surv.colour = 'orange', censor.colour = 'red',
         main='Kaplan-Meier estimate with 95% confidence bounds\n
         Skin Cutaneous Melanoma', xlab='Time (days)')
```

Finally, we explore the association between the clinical variables using the previously defined `clinCorrelation` function.

```
clin_var <- c("years_to_birth",
              "days_to_submitted_specimen_dx",
              "primary_site_of_disease",
              "neoplasm_diseasestage",
              "pathology_T_stage",
              "pathology_N_stage",
              "pathology_M_stage",
              "melanoma_ulceration",
              "Breslow_thickness",
              "dcc_upload_date",
              "gender",
              "date_of_initial_pathologic_diagnosis",
              "radiations_radiation_regimenindication",
              "race",
              "ethnicity",
              "batch_number",
              "survivalOutcome")

clinCorrelation(clinical=clinical_skcm2, clin_var=clin_var, maxlogPvalue = 10)
```

As we can see in the representation there’s a significant association between `batch_number` and `dcc_upload_date`. The variable `days_to_submitted_specimen_dx` corresponds to the age in years of the case at the initial pathologic diagnosis of cancer. Moreover, `neoplasm_diseasestage` shows significant association with `pathology_T_stage`, `pathology_N_stage` and `pathology_M_stage`, a result that could be expected as all them are assessing the disease stage.

##### 2.1.2 Individual omics data-type exploration

Previous to any integrative approach we should know which type of data do we have. Depending on the initial data-type we should apply some normalization, correction or transformation process.

###### 2.1.2.1 Expression data (`rna_skcm`)

Expression data from SKCM is generated under RNA-Seq technology (Illumina HiSeq 2000), so we will have count data. As we are working with level 3 data, the data has been already normalized. According to TCGA information, gene expression was quantified for the transcript models corresponding to the TCGA GAF2.1, using RSEM (accurate transcript quantification tool from RNA-Seq data with or without a reference genome - Li and Dewey, 2011) and normalized within-sample to a fixed upper quartile (UQ: the total counts are replaced by the upper quartile of counts different from 0 in the computation of the normalization factors.

```
#let's explore the data to be sure what type of data do we have
summary(rna_skcm[,1:3])
```

```
##   TCGA-GN-A4U7       TCGA-W3-AA1V        TCGA-FS-A1Z0     
##  Min.   :     0.0   Min.   :     0.00   Min.   :     0.0  
##  1st Qu.:     2.3   1st Qu.:     1.88   1st Qu.:     2.4  
##  Median :   120.0   Median :   155.79   Median :   173.6  
##  Mean   :  1024.9   Mean   :   906.26   Mean   :   848.0  
##  3rd Qu.:   763.9   3rd Qu.:   802.70   3rd Qu.:   810.6  
##  Max.   :361073.8   Max.   :129348.46   Max.   :158442.1
```

First of all, we will remove the three replicated samples identified before.

```
rna_skcm <- rna_skcm[,-rep_samples]
```

Then, we explore the distribution of the UQ normalized data.

```
# Prepare data for ggplot
sampleNames = vector()
intensities = vector()
for (i in 1:ncol(rna_skcm)){
  sampleNames = c(sampleNames,rep(colnames(rna_skcm)[i],nrow(rna_skcm)))
  intensities = c(intensities,rna_skcm[,i])
}

arrayData <- data.frame(log2(intensities+1),sampleNames)
colnames(arrayData)[1] <- "intensities"

g = ggplot(arrayData, aes(intensities, color=sampleNames))

g + geom_density() + 
  theme(legend.position='none') +
  labs(title='Gene Expression Density Curve',
       x='Log2 UQ Normalized Counts',
       y = 'Density')
```

Log2-transformation of data is required for forward analyses and 306 features with missings in all samples will be removed.

```
sel_na <- which(rowSums(rna_skcm)==0)
length(sel_na)
```

```
## [1] 306
```

```
#log2 transformation
log2_rna_skcm <- log2(rna_skcm[-sel_na,]+1)
```

Finally, the input data for gene expression of SKCM consist in a UQ normalized log2-transformed matrix with 20,225 features per 425 samples.

Let’s explore a little bit more this omic data-type alone using a dimension reduction methodology such as Principal Component analysis (PCA) in order to see if there’s some batch-effect.

```
pca_rna <- prcomp(t(log2_rna_skcm))
var_prcomp <- pca_rna$sdev^2

# Plot PCs variance
pcvar <- data.frame(var=var_prcomp/sum(var_prcomp), pc=c(1:length(var_prcomp)))
ggplot(pcvar[1:15,], aes(x = pc)) + geom_line(aes(y=var)) + 
  labs(x='Principal Component', y='Explained Variance') + 
  geom_point(aes(y=var)) + 
  ggtitle('RNA: Principal Components variance')
```

```
# Plot PCs scores
df <- data.frame(PC1=pca_rna$x[,1], PC2=pca_rna$x[,2], 
                 batch_number= clinical_skcm[colnames(log2_rna_skcm),
                              'batch_number'])

ggplot(df) +
  geom_point(aes(x=PC1, y=PC2, color=factor(batch_number)),size=5,shape=20) +
  guides(color=guide_legend('batch_number'),fill=guide_legend('batch_number'))+
  labs(x='PC1 (9.9%)', y='PC2 (8.4%)') +
  ggtitle('Principal Components for RNA data by Batch') +
  theme(legend.position="bottom")
```

The association between the clinical variables and principal components was explored using the `pcaCorrelation` function.

```
pcaCorrelation(clinical=clinical_skcm[colnames(log2_rna_skcm),],
               clin_var=clin_var, pca_data=pca_rna$x, maxlogPvalue = 10)
```

A significant association is observed between `gender` and PC16. Moreover, significant association is identified between principal components and `survivalOutcome`, `primary_site_of_disease`, `batch_number` and `days_to_submitted_specimen_dx` (corresponding to the age in years of the case at the initial pathological diagnosis of disease or cancer).

The most important point from these results is that a batch effect has been identified, pointing to the need of a batch correction.

###### 2.1.2.2 miRNA data (`mirna_skcm`)

miRNA data from SKCM is generated under miRNA-Seq technology (Illumina HiSeq 2000 ), so we will have count data. As we are working with level 3 data, counts normalized to reads per million mapped reads (RPM) is our initial data.

First of all, we will remove the three repeted samples identified before.

```
mirna_skcm <- mirna_skcm[,-rep_samples]
```

Then, we explore the distribution of the RPM data.

```
summary(mirna_skcm[,1:3])
```

```
##   TCGA-GN-A4U7        TCGA-W3-AA1V        TCGA-FS-A1Z0      
##  Min.   :     0.00   Min.   :     0.00   Min.   :     0.00  
##  1st Qu.:     0.00   1st Qu.:     0.00   1st Qu.:     0.00  
##  Median :     0.00   Median :     0.00   Median :     0.18  
##  Mean   :   947.67   Mean   :   944.07   Mean   :   951.79  
##  3rd Qu.:     7.19   3rd Qu.:     6.23   3rd Qu.:     5.59  
##  Max.   :192491.91   Max.   :137164.20   Max.   :141427.20  
##  NA's   :1           NA's   :1           NA's   :1
```

```
# Prepare data for ggplot
sampleNames = vector()
intensities = vector()
for (i in 1:ncol(mirna_skcm)){
  sampleNames = c(sampleNames,rep(colnames(mirna_skcm)[i],nrow(mirna_skcm)))
  intensities = c(intensities,mirna_skcm[,i])
}

arrayData <- data.frame(log2(intensities+1),sampleNames)
colnames(arrayData)[1] <- "intensities"

g = ggplot(arrayData, aes(intensities, color=sampleNames))
g + geom_density() + theme(legend.position='none') +
  labs(title='miRNA Density Curve', 
       x='Log2 RPM', 
       y='Density')
```

```
## Warning: Removed 425 rows containing non-finite values (stat_density).
```

Within this data set we found 1 missing feature and 147 features with 0 counts in all sampels. We decide to remove it.

```
sel_na <- which(rowSums(mirna_skcm)==0 | is.na(mirna_skcm[,1]))

#log2-transofrmation
log2_mirna_skcm <- log2(mirna_skcm[-sel_na,]+1)
```

Finally, the input data for miRNA expression of SKCM consist in a log2-transformed RPM matrix with 898 features per 425 samples.

Let’s explore a little bit more this omic alone, using a dimension reduction methodology such as Principal Component analysis (PCA), in order to see if there’s some batch-effect.

```
pca_mirna <- prcomp(t(log2_mirna_skcm))
var_prcomp <- pca_mirna$sdev^2

# Plot PCs variance
pcvar <- data.frame(var=var_prcomp/sum(var_prcomp), pc=c(1:length(var_prcomp)))
ggplot(pcvar[1:15,], aes(x = pc)) + geom_line(aes(y=var)) + 
  labs(x='Principal Component', y='Explained Variance') + 
  geom_point(aes(y=var)) + 
  ggtitle('miRNA: Principal Components variance')
```

```
# Plot PCs scores
df <- data.frame(PC1=pca_mirna$x[,1], PC2=pca_mirna$x[,2],
                 batch_number=clinical_skcm[colnames(log2_mirna_skcm),
                                            'batch_number'])

ggplot(df) +
  geom_point(aes(x=PC1, y=PC2, color=factor(batch_number)),size=5,shape=20)+
  guides(color=guide_legend('batch_number'),fill=guide_legend('batch_number'))+
  labs(x='PC1 (16.8%)', y='PC2 (10.7%)') +
  ggtitle('Principal Components for miRNA data by Batch') +
  theme(legend.position="bottom")
```

The association between the clinical variables and principal components was explored using the `pcaCorrelation` function.

```
pcaCorrelation(clinical=clinical_skcm[colnames(log2_mirna_skcm),],
               clin_var=clin_var, pca_data=pca_mirna$x, maxlogPvalue = 10)
```

In this case, batch effect is also detected as significant with several principal components. The other variable that shows significant association with some of the principals components is `primary_site_of_disease` and `days_to_submitted_specimen_dx` (corresponding to the age in years of the case at the initial pathological diagnosis of disease or cancer).

###### 2.1.2.3 Methylation data (`met_skcm`)

Methylation data from SKCM is generated under microarray technology (Illumina Infinium Human Methylation 450K). As we are working with level 3 data, the data has been already processes and the available data corresponds to beta values.

First of all, we will remove the three repeted samples previously identified.

```
met_skcm <- met_skcm[,-rep_samples]

summary(met_skcm[,1:4])
```

```
##   TCGA-GN-A4U7    TCGA-W3-AA1V    TCGA-FS-A1Z0    TCGA-D9-A3Z1  
##  Min.   :0.01    Min.   :0.01    Min.   :0.01    Min.   :0.01   
##  1st Qu.:0.04    1st Qu.:0.06    1st Qu.:0.06    1st Qu.:0.07   
##  Median :0.24    Median :0.41    Median :0.46    Median :0.50   
##  Mean   :0.38    Mean   :0.44    Mean   :0.46    Mean   :0.46   
##  3rd Qu.:0.77    3rd Qu.:0.81    3rd Qu.:0.84    3rd Qu.:0.79   
##  Max.   :0.99    Max.   :0.99    Max.   :0.99    Max.   :0.99   
##  NA's   :90563   NA's   :90727   NA's   :89939   NA's   :89594
```

As we can see in the summary of the first 4 samples there are a large number of missings

```
number_NAs <- rowSums(is.na(met_skcm))
sum(number_NAs!=0) #111,259 festures without NAs
```

```
## [1] 111259
```

```
sum(number_NAs==ncol(met_skcm)) #89,512 features with all NAs
```

```
## [1] 89512
```

There are 89,512 features with missing values in all cases, moreover there are 21,747 features with missings in at least one case. We decide to remove all samples that have at least one missing value.

```
met_skcm <- met_skcm[number_NAs==0,] #New dimension: 374318 x 425
```

Moreover, additional filters are applied based on probes annotation. Gender related probes (chr Y and X) and probes with SNPs, CpG or SBE in target sequence are removed.

```
# Load require packages
library(IlluminaHumanMethylation450kanno.ilmn12.hg19)
```

```
# Get annotation data from methylation
ann450k <- getAnnotation(IlluminaHumanMethylation450kanno.ilmn12.hg19)
ann450k[1:4,1:6]
```

```
## DataFrame with 4 rows and 6 columns
##                    chr       pos      strand        Name    AddressA
##            <character> <integer> <character> <character> <character>
## cg00050873        chrY   9363356           -  cg00050873    32735311
## cg00212031        chrY  21239348           -  cg00212031    29674443
## cg00213748        chrY   8148233           -  cg00213748    30703409
## cg00214611        chrY  15815688           -  cg00214611    69792329
##               AddressB
##            <character>
## cg00050873    31717405
## cg00212031    38703326
## cg00213748    36767301
## cg00214611    46723459
```

```
# Gender related probes
xy_linked <- which(ann450k$chr=='chrY' | ann450k$chr=='chrX') #11648 features

# SNPs, CpG and SBE in target sequence
target_snp <- which(!is.na(ann450k$Probe_rs)) #87,018 features
target_cpg <- which(!is.na(ann450k$CpG_rs)) #16,998 features
target_sbe <- which(!is.na(ann450k$SBE_rs)) #7,876 features

# Filter dataset
rm_probes <- unique(c(xy_linked,target_snp, target_sbe, target_cpg)) 
#114,614 features

met_skcm <- met_skcm[rownames(met_skcm)%in%rownames(ann450k)[-rm_probes],] 
#New dimension: 305,180 x 425
```

Finally, we transform the beta values to M values.

```
# Load require packages
library(lumi)

# Transform to M values
logit_met_skcm <- beta2m(met_skcm) 

summary(logit_met_skcm[,1:4])
```

```
##   TCGA-GN-A4U7     TCGA-W3-AA1V      TCGA-FS-A1Z0      TCGA-D9-A3Z1      
##  Min.   :-7.153   Min.   :-7.1649   Min.   :-7.2767   Min.   :-7.321959  
##  1st Qu.:-4.532   1st Qu.:-3.9062   1st Qu.:-4.0524   1st Qu.:-3.728057  
##  Median :-1.777   Median :-0.5127   Median :-0.3085   Median : 0.006373  
##  Mean   :-1.352   Mean   :-0.7285   Mean   :-0.6494   Mean   :-0.592811  
##  3rd Qu.: 1.841   3rd Qu.: 2.1708   3rd Qu.: 2.4274   3rd Qu.: 1.896938  
##  Max.   : 7.434   Max.   : 7.3051   Max.   : 7.1835   Max.   : 7.632917
```

The input data for methylation of SKCM consist in a M values matrix with 305,180 features per 425 samples.

Let’s explore a little bit more this omic alone, using a dimension reduction methodology such as Principal Component analysis (PCA), in order to see if there’s some batch-effect.

```
pca_met <- prcomp(t(logit_met_skcm))
var_prcomp <- pca_met$sdev^2

# Plot PCs variance
pcvar <- data.frame(var=var_prcomp/sum(var_prcomp), pc=c(1:length(var_prcomp)))
ggplot(pcvar[1:15,], aes(x = pc)) + geom_line(aes(y=var)) + 
  labs(x='Principal Component', y='Explained Variance') + 
  geom_point(aes(y=var)) + 
  ggtitle('Methylation: Principal Components variance')
```

```
# Plot PCs scores
df <- data.frame(PC1=pca_met$x[,1], PC2=pca_met$x[,2], 
                 batch_number=clinical_skcm[colnames(logit_met_skcm),
                              'batch_number'])

ggplot(df) +
  geom_point(aes(x=PC1, y=PC2, color=factor(batch_number)), size=5, shape=20) +
  guides(color=guide_legend('batch_number'),fill=guide_legend('batch_number')) +
  labs(x='PC1 (15.8%)', y='PC2 (11.2%)') +
  ggtitle('Principal Components for Methylation data by Batch') +
  theme(legend.position="bottom")
```

A function to explore the association between the clinical variables and principal components called `pcaCorrelation` is defined.

```
pcaCorrelation(clinical=clinical_skcm[colnames(logit_met_skcm),], 
               clin_var=clin_var, pca_data=pca_met$x, maxlogPvalue = 10)
```

In this case, `batch_number` is also detected as significantly associated with several principal components. The other variables that show significant association with some of the principals components are `primary_site_of_disease`, `days_to_submitted_specimen_dx` and `date_of_initial_pathologic_diagnosis`.

In general, although not in first components, these results highlight a batch effect in methylation data-type that must be corrected.

##### 2.1.3 Joint omics exploration

###### 2.1.3.1 Common samples

We have the same number of samples between omics, but let’s see if these are share between omics.

```
# Load require packages
library(VennDiagram)
library(plotrix)
```

```
m <- list(rna=colnames(log2_rna_skcm), 
          mirna=colnames(log2_mirna_skcm), 
          met=colnames(logit_met_skcm))

vennPlot <- venn.diagram(m, NULL, fill=c('red','green','blue'), 
                         cat.fontface=4, lty=2, cex=1.2, cat.cex=1.2)
grid.newpage()
grid.draw(vennPlot)
```

All samples are share between the three different data-types (mRNA, miRNA and methylation).

###### 2.1.3.2 OmicsPCA exploration

The exploration of each omic data-type independently have been shown a batch effect that must be corrected, but let’s explore what happens if we perform a joint exploration using the `omicsPCA` function from `STATegRa` package.

```
library(STATegRa)
library(MASS)
library(gridExtra)
```

####Data scaling

Before start the multi-omic approach, mean centred of each feature and the data-type normalization to unit sum of squares (forbenious normalization) is recommended to make more comparabable the different omics data-types.

```
#Calculate Forbenious normalization
frobenius_rna <- norm(as.matrix(log2_rna_skcm), type="F")
frobenius_mirna <- norm(as.matrix(log2_mirna_skcm), type="F")
frobenius_met <- norm(as.matrix(logit_met_skcm), type="F")

#Mean-centring and division by Frobenius normalization factor
rna_skcm_norm <- t(scale(t(log2_rna_skcm), scale=FALSE))/frobenius_rna
mirna_skcm_norm <- t(scale(t(log2_mirna_skcm), scale=FALSE))/frobenius_mirna
met_skcm_norm <- t(scale(t(logit_met_skcm), scale=FALSE))/frobenius_met

##Mean-centring without Frobenius normalization
rna_skcm_norm2 <- t(scale(t(log2_rna_skcm), scale=FALSE))
mirna_skcm_norm2 <- t(scale(t(log2_mirna_skcm), scale=FALSE))
met_skcm_norm2 <- t(scale(t(logit_met_skcm), scale=FALSE))
```

###### 2.1.3.3 Model selection

Now, we are ready to explore if there are share components between data-types.

The first step is to determine the number of share/common and individual components between omics.

How to determine these numbers is still an open question nowadays, so the user can explore different methods like jive, pca-gca and pESCA, between others. Following, we show how to perform the model selection based on these three methods.

###### 2.1.3.3.1 JIVE

```
# Load require packages
library(r.jive)
```

```
#Using r.jive

####################################################
## mRNA + miRNA (already centered and normalized) ##
####################################################
Data <- list(mRNA=rna_skcm_norm,
             miRNA=mirna_skcm_norm)
rjive_rna_mirna <- jive(Data, scale = FALSE, center = FALSE) 


##########################################################
## mRNA + methylation (already centered and normalized) ##
##########################################################
Data <- list(mRNA=rna_skcm_norm,
              met=met_skcm_norm)
rjive_rna_met <- jive(Data, scale = FALSE, center = FALSE)
```

###### 2.1.3.3.2 PCA-GCA

```
# Load require packages
library(RegularizedSCA)

#Using pca-gca

####################################################
## mRNA + miRNA (already centered and normalized) ##
####################################################
data <- cbind(t(rna_skcm_norm), 
              t(mirna_skcm_norm))

# Number of variables
Jk <- c(nrow(rna_skcm_norm), nrow(mirna_skcm_norm))

# Select components
pca_gca(data, Jk, cor_min = .7)

##########################################################
## mRNA + methylation (already centered and normalized) ##
##########################################################
data <- cbind(t(rna_skcm_norm), 
              t(met_skcm_norm))

# Number of variables
Jk <- c(nrow(rna_skcm_norm), nrow(met_skcm_norm))

# Select components
pca_gca(data, Jk, cor_min = .7)
```

###### 2.1.3.3.3 pESCA

```
# Load require packages
library(RpESCA)
library(RSpectra)

#Using RpESCA

################################################
## mRNA + miRNA (already centered)            ##
################################################

dataSets <- list(rna=t(rna_skcm_norm2), 
                 mirna=t(mirna_skcm_norm2))

dataTypes <- 'GG'

# used parameters
opts <- list()
opts$tol_obj <- 1E-4
opts$quiet <- 1

# save the estimated alphas, selected number of PCs and cvErrors
alphas <- rep(NA,2)
R_selected_list <- as.list(1:2)
cvErrors_list <- as.list(1:2)

# alpha estimation for each data set
for (i in 1:length(dataSets)){
  # index ith data set
  X <- dataSets[[i]]
  
  # alpha estimation procedure
  alpha_est <- alpha_estimation(X,K=3,Rs = 5:20,opts=opts)
  alphas[i] <- alpha_est$alphas_mean
  R_selected_list[[i]] <- alpha_est$R_CV
  cvErrors_list[[i]] <- alpha_est$cvErrors
}
names(alphas) <- paste0('alpha', 1:2)

## Setting the parameters for the pESCA model

# concave function and its hyper-parameter
fun_concave <- 'gdp'; gamma <- 1;
penalty = 'L2' # concave L2 norm penalty

# Parameters of a pESCA with concave L2norm penalty model
opts <- list()
opts$gamma <- gamma  # hyper-parameter for the used penalty
opts$rand_start <- 0
opts$tol_obj <- 1e-6 # stopping criteria
opts$maxit   <- 500
opts$alphas  <- alphas
#opts$R # components used Default 0.5*min(I,J) - Minimum number of variables.
opts$thr_path <- 0 # generaint thresholding path or not
opts$quiet <- 1

# Model selection pESCA conave L2 norm penalty
nTries <- 15
lambdas_CV <- log10_seq(from=1, to=500, length.out=nTries)

result_CV <- pESCA_CV(dataSets, dataTypes,
                      lambdas_CV, 
                      penalty=penalty, 
                      fun_concave=fun_concave, 
                      opts=opts)


# select the model with minimum CV error
index_min_cv <- which.min(result_CV$cvErrors_mat[,1])

# cvErrors during the model selection process
result_CV$cvErrors_mat

# selected value of lambda
lambdas_CV[index_min_cv]

# fit the final model
lambdas_opt <- rep(lambdas_CV[index_min_cv],length(dataSets))
opts_opt <- result_CV$inits[[index_min_cv]]
opts_opt$tol_obj <- 1E-8 # using high precision model


pESCA_L2 <- pESCA(dataSets = dataSets,
                          dataTypes = dataTypes,
                          lambdas = lambdas_opt,
                          penalty = penalty,
                          fun_concave= fun_concave,
                          opts = opts_opt)


mu <- pESCA_L2$mu
A <- pESCA_L2$A
B <- pESCA_L2$B
S <- pESCA_L2$S

# estimated variation explained ratios for each data set or the full data set
pESCA_L2$varExpTotals

# estimated variation explained ratios of each PC for each data set or the full data set
pESCA_L2$varExpPCs

#Cut-off 1% 
sel1 <- which(pESCA_L2$varExpPCs[1,]>1) #14
sel2 <- which(pESCA_L2$varExpPCs[2,]>1) #20
intersect(sel1,sel2)
# 10 common components
# 4 dist mRNA + 10 miRNA
# Total: 24 components

#Cut-off 5%
sel1 <- which(pESCA_L2$varExpPCs[1,]>5) #3
sel2 <- which(pESCA_L2$varExpPCs[2,]>5) #2
intersect(sel1,sel2)
# 1 common components
# 2 dist mRNA + 1 miRNA
# Total: 4 components


################################################
## mRNA + Methylation (already centered)      ##
################################################

dataSets <- list(rna=t(rna_skcm_norm2), 
                 met=t(met_skcm_norm2))

dataTypes <- 'GG'

# used parameters
opts <- list()
opts$tol_obj <- 1E-4
opts$quiet <- 1

# save the estimated alphas, selected number of PCs and cvErrors
alphas <- rep(NA,2)
R_selected_list <- as.list(1:2)
cvErrors_list <- as.list(1:2)

# alpha estimation for each data set
for (i in 1:length(dataSets)){
  # index ith data set
  X <- dataSets[[i]]
  
  # alpha estimation procedure
  alpha_est <- alpha_estimation(X,K=3,Rs = 5:20,opts=opts)
  alphas[i] <- alpha_est$alphas_mean
  R_selected_list[[i]] <- alpha_est$R_CV
  cvErrors_list[[i]] <- alpha_est$cvErrors
}
names(alphas) <- paste0('alpha', 1:2)

## Setting the parameters for the pESCA model

# concave function and its hyper-parameter
fun_concave <- 'gdp'; gamma <- 1;
penalty = 'L2' # concave L2 norm penalty

# Parameters of a pESCA with concave L2norm penalty model
opts <- list()
opts$gamma <- gamma  # hyper-parameter for the used penalty
opts$rand_start <- 0
opts$tol_obj <- 1e-6 # stopping criteria
opts$maxit   <- 500
opts$alphas  <- alphas
#opts$R  # components used Default 0.5*min(I,J) - Minimum number of variables.
opts$thr_path <- 0 # generaint thresholding path or not
opts$quiet <- 1

# Model selection pESCA conave L2 norm penalty
nTries <- 15
lambdas_CV <- log10_seq(from=1, to=500, length.out=nTries)

result_CV <- pESCA_CV(dataSets, dataTypes,
                      lambdas_CV, 
                      penalty=penalty, 
                      fun_concave=fun_concave, 
                      opts=opts)


# select the model with minimum CV error
index_min_cv <- which.min(result_CV$cvErrors_mat[,1])

# cvErrors during the model selection process
result_CV$cvErrors_mat

# selected value of lambda
lambdas_CV[index_min_cv]

# fit the final model
lambdas_opt <- rep(lambdas_CV[index_min_cv],length(dataSets))
opts_opt <- result_CV$inits[[index_min_cv]]
opts_opt$tol_obj <- 1E-8 # using high precision model


pESCA_L2 <- pESCA(dataSets = dataSets,
                          dataTypes = dataTypes,
                          lambdas = lambdas_opt,
                          penalty = penalty,
                          fun_concave= fun_concave,
                          opts = opts_opt)

mu <- pESCA_L2$mu
A <- pESCA_L2$A
B <- pESCA_L2$B
S <- pESCA_L2$S

# estimated variation explained ratios for each data set or the full data set
pESCA_L2$varExpTotals

# estimated variation explained ratios of each PC for each data set or the full data set
pESCA_L2$varExpPCs

#Cut-off 1%
sel1 <- which(pESCA_L2$varExpPCs[1,]>1) #14
sel2 <- which(pESCA_L2$varExpPCs[2,]>1) #10
intersect(sel1,sel2)
# 7 common components
# 7 dist mRNA + 3 methylation
# Total: 17 components

#Cut-off 5%
sel1 <- which(pESCA_L2$varExpPCs[1,]>5) #3
sel2 <- which(pESCA_L2$varExpPCs[2,]>5) #3
intersect(sel1,sel2)
# 1 common components
# 2 dist mRNA + 2 methylation
# Total: 5 components
```

Here, there is the table summarizing the output of the different methods.

|  | Component | r.JIVE | PCA-GCA (1%) | PCA-GCA (5%) | pESCA(1%) | pESCA (5%) |
| --- | --- | --- | --- | --- | --- | --- |
| mRNA + miRNA |  |  |  |  |  |  |
|  | Common | 6 | 26 | 0 | 10 | 1 |
|  | Dist. mRNA | 56 | 14 | 1 | 4 | 2 |
|  | Dist. miRNA | 33 | 41 | 4 | 10 | 1 |
| mRNA + met |  |  |  |  |  |  |
|  | Common | 6 | 38 | 1 | 7 | 1 |
|  | Dist. mRNA | 55 | 2 | 0 | 7 | 2 |
|  | Dist. miRNA | 110 | 184 | 4 | 3 | 2 |

As we can observe, the results from the different methods diverge. We are not able to recommend a method but, in general, we can say that it seems that there’s a small part of data variation that is share between omics. To follow with the analysis, we will explore the omicsPCA based on the output from pESCA (1%).

###### 2.1.3.4 Sub-space recovery

Based on the number of components identified (pESCA) we can recover the subspace.

It is interesting to take a look on the common and distinctive components identified through this method.

```
################################################
## mRNA + miRNA (already centered)            ##
################################################

# Explore the common and distinctive components and those variation explained
# Common components
cc <- intersect(sel1,sel2)
pESCA_L2$varExpPCs[,cc]
```

```
##             PC1      PC2      PC3      PC4      PC5      PC6      PC7      PC8
## X_1    9.708634 8.278301 6.631362 3.918253 3.498610 2.732839 2.058715 2.069639
## X_2    8.247317 3.685949 3.936646 5.073806 1.374215 2.053595 4.706448 1.305732
## X_full 9.612973 7.977675 6.454959 3.993898 3.359543 2.688374 2.232041 2.019632
##            PC11     PC12
## X_1    1.246845 1.088903
## X_2    3.533955 3.324827
## X_full 1.396564 1.235271
```

```
# Distinctive components
dist_rna <- sel1[!sel1%in%cc] #rna
pESCA_L2$varExpPCs[,dist_rna]
```

```
##              PC9      PC10      PC13      PC15
## X_1    1.9897776 1.3713951 1.1502629 1.0396666
## X_2    0.7312919 0.7382829 0.6499185 0.0000000
## X_full 1.9073943 1.3299502 1.1175092 0.9716076
```

```
dist_mirna <- sel2[!sel2%in%cc] #mirna
pESCA_L2$varExpPCs[,dist_mirna]
```

```
##             PC22      PC28      PC35      PC39      PC64      PC72      PC82
## X_1    0.5206619 0.5367749 0.4108676 0.2857387 0.0000000 0.0000000 0.0000000
## X_2    1.0389138 2.7831835 1.2591479 3.6434042 2.5259777 1.8815695 2.8489401
## X_full 0.5545879 0.6838299 0.4663980 0.5055392 0.1653563 0.1231718 0.1864981
##            PC86       PC87      PC91
## X_1    0.000000 0.00000000 0.0000000
## X_2    4.633309 1.19060667 2.7792485
## X_full 0.303307 0.07793983 0.1819359
```

```
# Keep de common and distinctive components
common_pc <- pESCA_L2$A[,cc]
colnames(common_pc) <- colnames(pESCA_L2$varExpPCs[,cc])

dist_rna_pc <- pESCA_L2$A[,dist_rna]
colnames(dist_rna_pc) <- names(dist_rna)

dist_mirna_pc <- pESCA_L2$A[,dist_mirna]
colnames(dist_mirna_pc) <- names(dist_mirna)


# metadata for common samples
clin <- clinical_skcm

# PCA plot
require(ggplot2)

df <- as.data.frame(common_pc)
df$group <- clin$primary_site_of_disease
df$batch <- clin$batch_number

#plot of first common components by gene expression subtype
p <- ggplot(df,aes_string("PC1","PC2",color="group"))
p + geom_point(size=5, shape=20) + ggtitle("PCA by Primary site of tumor") + theme(legend.position="bottom")
```

```
#plot of first common components by batch
p <- ggplot(df,aes_string("PC1","PC2",color="batch"))
p + geom_point(size=5, shape=20) + ggtitle("PCA by batch") + theme(legend.position="bottom")
```

Let’s explore how relate these common and distinctive components with the clinical variables.

```
df <- data.frame(common_pc,dist_rna_pc,dist_mirna_pc)
colnames(df) <- c(paste("common_",colnames(common_pc),sep=""),
                  paste("dist_rna_",colnames(dist_rna_pc),sep=""),
                  paste("dist_mirna_",colnames(dist_mirna_pc),sep=""))

pcaCorrelation(clinical=clin, clin_var, df, maxlogPvalue =10, maxPC=ncol(df), pc.names=TRUE)
```

The main common variability is associated with `primary_site_of_disease`. Remarkably, although not shared in the first common components, there is a strong batch effect in both, common and distinctive components.

Now, let’s explore what happens with the joint exploration of RNA and DNA methylation.

```
################################################
## mRNA + methylation (already centered)      ##
################################################

# Explore the common and distinctive components and those variation explained
# Common components
cc <- intersect(sel1,sel2)
pESCA_L2$varExpPCs[,cc]
```

```
##              PC1       PC2      PC3      PC4      PC6      PC8      PC9
## X_1     1.750222  7.235064 2.633154 2.093371 5.913507 1.394186 3.008767
## X_2    15.777109 11.048408 6.581409 2.749514 1.900028 1.488947 1.096054
## X_full 14.924348 10.816577 6.341376 2.709624 2.144026 1.483186 1.212337
```

```
# Distinctive components
dist_rna <- sel1[!sel1%in%cc] #rna
pESCA_L2$varExpPCs[,dist_rna]
```

```
##             PC15      PC22      PC23      PC27      PC44       PC66      PC84
## X_1    3.6949149 2.0682024 1.2743320 5.0552575 1.7474906 1.28043731 2.7846385
## X_2    0.5647188 0.6765279 0.4627220 0.0000000 0.2356691 0.00000000 0.0000000
## X_full 0.7550183 0.7611344 0.5120636 0.3073331 0.3275799 0.07784387 0.1692914
```

```
dist_met <- sel2[!sel2%in%cc] #met
pESCA_L2$varExpPCs[,dist_met]
```

```
##             PC5       PC7     PC16
## X_1    0.000000 0.9350582 0.000000
## X_2    2.284182 1.2876516 1.127067
## X_full 2.145316 1.2662158 1.058547
```

```
# Keep de common and distinctive components
common_pc <- pESCA_L2$A[,cc]
colnames(common_pc) <- colnames(pESCA_L2$varExpPCs[,cc])

dist_rna_pc <- data.frame(pESCA_L2$A[,dist_rna])
colnames(dist_rna_pc) <- names(dist_rna)

dist_met_pc <- pESCA_L2$A[,dist_met]
colnames(dist_met_pc) <- names(dist_met)


# metadata for common samples
clin <- clinical_skcm

# PCA plot
require(ggplot2)

df <- as.data.frame(common_pc)
df$group <- clin$primary_site_of_disease
df$batch <- clin$batch_number

#plot of first common components by gene expression subtype
p <- ggplot(df,aes_string("PC1","PC2",color="group"))
p + geom_point(size=5, shape=20) + ggtitle("PCA by Primary site of tumor") + theme(legend.position="bottom")
```

```
#plot of first common components by batch
p <- ggplot(df,aes_string("PC1","PC2",color="batch"))
p + geom_point(size=5, shape=20) + ggtitle("PCA by batch") + theme(legend.position="bottom")
```

Let’s explore how relate these common and distinctive components with the clinical variables.

```
df <- data.frame(common_pc,dist_rna_pc,dist_met_pc)
colnames(df) <- c(paste("common_",colnames(common_pc),sep=""),
                  paste("dist_rna_",colnames(dist_rna_pc),sep=""),
                  paste("dist_met_",colnames(dist_met_pc),sep=""))

pcaCorrelation(clinical=clin, clin_var, df, maxlogPvalue =10, maxPC=ncol(df), pc.names=TRUE)
```

The association study shows that there’s no clear shared variable explaining the common components. The most relevant ones seems to be `primary_site_of_disease` and `days_to_submitted_specimen_dx`. Nonethless, significant batch effect is detected in the first distinctive components.

##### 2.1.4 Batch correction

In order to remove batch effect from the different omics data we applied the ComBat method

```
library(sva)
```

Before to apply ComBat, it’s important to note that there is one batch with only one sample

```
summary(as.factor(as.character(clinical_skcm[,'batch_number'])))
```

```
## 180.50.0 180.65.0 180.66.0 198.62.0 198.63.0 206.64.0 240.58.0 262.54.0 
##        1       12      204        5       16       13       20        9 
## 277.56.0 291.49.0 316.46.0 332.37.0 358.31.0 388.28.0 393.26.0 408.20.0 
##       28       13       15       15       11       33       17        7 
## 416.18.0 
##        6
```

```
clinical_skcm[which(clinical_skcm$batch_number=='180.50.0'),20:22]
```

```
##               race              ethnicity batch_number
## TCGA-FS-A1ZE white not hispanic or latino     180.50.0
```

```
clinical_skcm <- clinical_skcm[rownames(clinical_skcm)!='TCGA-FS-A1ZE',]
```

This sample must be removed from all data sets to apply the batch correction.

```
log2_rna_skcm_corrected <- log2_rna_skcm[,colnames(log2_rna_skcm)!='TCGA-FS-A1ZE'] 
#New dimension: 20225 x 424

log2_mirna_skcm_corrected <- log2_mirna_skcm[,colnames(log2_mirna_skcm)!='TCGA-FS-A1ZE'] #New dimension: 898 x 424

logit_met_skcm_corrected <- logit_met_skcm[,colnames(logit_met_skcm)!='TCGA-FS-A1ZE'] #New dimension: 305180 x 424
```

Now, we are ready to apply the batch correction for each omic data.

###### 2.1.4.1 Expression data (`log2_rna_skcm_corrected`)

```
mod <- model.matrix(~gender, 
                    data=clinical_skcm)

batch <- as.factor(as.character(clinical_skcm$batch_number))

log2_rna_skcm_corrected <- ComBat(log2_rna_skcm_corrected,
                            batch=batch,
                            mod=mod, 
                            prior.plot=TRUE)
```

```
## Found 2198 genes with uniform expression within a single batch (all zeros); these will not be adjusted for batch.
```

Exploring the PCA from corrected data we can see that the batch effect have been removed.

```
pca_rna <- prcomp(t(log2_rna_skcm_corrected))
var_prcomp <- pca_rna$sdev^2

# Plot PCs variance
pcvar <- data.frame(var=var_prcomp/sum(var_prcomp), pc=c(1:length(var_prcomp)))
ggplot(pcvar[1:15,], aes(x = pc)) + geom_line(aes(y=var)) + 
  labs(x='Principal Component', y='Explained Variance') + 
  geom_point(aes(y=var)) + 
  ggtitle('RNA: Principal Components variance')
```

```
# Plot PCs scores
df <- data.frame(PC1=pca_rna$x[,1], PC2=pca_rna$x[,2],
                 batch_number=clinical_skcm$batch_number)

ggplot(df) +
  geom_point(aes(x=PC1, y=PC2, color=factor(batch_number)), size=5, shape=20) +
  guides(color=guide_legend("batch_number"),fill=guide_legend("batch_number")) +
  ggtitle("Principal Components from batch-adjusted RNA data") +
  theme(legend.position="bottom")
```

Now, the association analysis shows how the batch effect is clearly removed from our data and the main variability is related with gene expression subtype.

```
pcaCorrelation(clinical=clinical_skcm,
               clin_var=clin_var, pca_data=pca_rna$x, maxlogPvalue = 10)
```

###### 2.1.4.2 miRNA data (`log2_mirna_skcm_corrected`)

```
log2_mirna_skcm_corrected <- ComBat(log2_mirna_skcm_corrected,
                              batch=batch,
                              mod=mod, 
                              prior.plot=TRUE)
```

```
## Found 318 genes with uniform expression within a single batch (all zeros); these will not be adjusted for batch.
```

Exploring the PCA from corrected data we can see that the batch effect have been removed.

```
#miRNA
pca_mirna <- prcomp(t(log2_mirna_skcm_corrected))
var_prcomp <- pca_mirna$sdev^2

# Plot PCs variance
pcvar <- data.frame(var=var_prcomp/sum(var_prcomp), pc=c(1:length(var_prcomp)))
ggplot(pcvar[1:15,], aes(x = pc)) + geom_line(aes(y=var)) + 
  labs(x='Principal Component', y='Explained Variance') + 
  geom_point(aes(y=var)) + 
  ggtitle('miRNA: Principal Components variance')
```

```
# Plot PCs scores
df <- data.frame(PC1=pca_mirna$x[,1], PC2=pca_mirna$x[,2],
                 batch_number=clinical_skcm$batch_number)

ggplot(df) +
  geom_point(aes(x=PC1, y=PC2, color=factor(batch_number)), size=5, shape=20) +
  guides(color=guide_legend("batch_number"),fill=guide_legend("batch_number")) +
  ggtitle("Principal Components from batch-adjusted miRNA data") +
  theme(legend.position="bottom")
```

```
pcaCorrelation(clinical=clinical_skcm,
               clin_var=clin_var, pca_data=pca_mirna$x, maxlogPvalue = 10)
```

###### 2.1.4.3 Methylation data (`logit_met_skcm`)

As methylation data no follows a gausian distribution, a non-parametric ComBat is applied.

```
logit_met_skcm_corrected <- ComBat(logit_met_skcm_corrected,
                                  batch=batch, 
                                  mod=mod, 
                                  prior.plot=TRUE, 
                                  par.prior=FALSE, 
                                  BPPARAM=MulticoreParam())
```

Note: the non-parametric ComBat spends long time (>4days) to compute the corrected matrix.

Exploring the PCA from corrected data we can see that the batch effect have been removed.

```
pca_met <- prcomp(t(logit_met_skcm_corrected))
var_prcomp <- pca_met$sdev^2

# Plot PCs variance
pcvar <- data.frame(var=var_prcomp/sum(var_prcomp), pc=c(1:length(var_prcomp)))
ggplot(pcvar[1:15,], aes(x = pc)) + geom_line(aes(y=var)) + 
  labs(x='Principal Component', y='Explained Variance') + 
  geom_point(aes(y=var)) + 
  ggtitle('Methylation: Principal Components variance')
```

```
# Plot PCs scores
df <- data.frame(PC1=pca_met$x[,1], PC2=pca_met$x[,2],
                 batch_number=clinical_skcm$batch_number)

ggplot(df) +
  geom_point(aes(x=PC1, y=PC2, color=factor(batch_number)), size=5, shape=20) +
  guides(color=guide_legend("batch_number"),fill=guide_legend("batch_number")) +
  ggtitle("Principal Components from batch-adjusted Methylation data") +
  theme(legend.position="bottom")
```

Now, the association analysis shows how the batch effect is clearly removed from our data.

```
pcaCorrelation(clinical=clinical_skcm[colnames(logit_met_skcm_corrected),],
               clin_var=clin_var, pca_data=pca_met$x, maxlogPvalue = 10)
```

#### 2.2 Characterization of the data: Individual exploration.

After data pre-proces (filter, normalization and batch correction) the exploration of target cases should be done.
First of all we will generate the data matrices with the data of study. We will work with just the 104 samples with curated metadata identified before.

```
subset_log2_rna_skcm_corrected <- log2_rna_skcm_corrected[,rownames(clinical_skcm2)]
subset_log2_mirna_skcm_corrected <- log2_mirna_skcm_corrected[,rownames(clinical_skcm2)]
subset_logit_met_skcm_corrected <- logit_met_skcm_corrected[,rownames(clinical_skcm2)]
```

Let’s see the dimension of the final data:

```
dim(subset_log2_rna_skcm_corrected)
```

```
## [1] 20225   104
```

```
dim(subset_log2_mirna_skcm_corrected)
```

```
## [1] 898 104
```

```
dim(subset_logit_met_skcm_corrected)
```

```
## [1] 305180    104
```

##### 2.2.1 Expression data (`subset_log2_rna_skcm_corrected`)

Exploring the PCA from mRNA data (20,225 x 104)

```
pca_rna <- prcomp(t(subset_log2_rna_skcm_corrected))
var_prcomp <- pca_rna$sdev^2

# Plot PCs variance
pcvar <- data.frame(var=var_prcomp/sum(var_prcomp), pc=c(1:length(var_prcomp)))
ggplot(pcvar[1:15,], aes(x = pc)) + geom_line(aes(y=var)) + 
  labs(x='Principal Component', y='Explained Variance') + 
  geom_point(aes(y=var)) + 
  ggtitle('mRNA: Principal Components variance')
```

```
# Plot PCs scores
df <- data.frame(PC1=pca_rna$x[,1], PC2=pca_rna$x[,2],
                 primary_site_of_disease=clinical_skcm2$primary_site_of_disease)

ggplot(df) +
  geom_point(aes(x=PC1, y=PC2, color=factor(primary_site_of_disease)), size=5, shape=20) +
  guides(color=guide_legend("primary_site_of_disease"),fill=guide_legend("primary_site_of_disease")) +
  labs(x='PC1 (11.88%)', y='PC2 (10.7%)') +
  ggtitle("Principal Components from batch-adjusted mRNA data") + 
  theme(legend.position="bottom")
```

The main variables explaining the highest variability are “primary site of disease” and “neoplasm disease stage”.

```
pcaCorrelation(clinical=clinical_skcm2[colnames(subset_log2_rna_skcm_corrected),],
               clin_var=clin_var, pca_data=pca_rna$x, maxlogPvalue=10)
```

##### 2.2.2 miRNA data (`subset_log2_mirna_skcm_corrected`)

Exploring the PCA from miRNA data (898 x 104)

```
pca_mirna <- prcomp(t(subset_log2_mirna_skcm_corrected))
var_prcomp <- pca_mirna$sdev^2

# Plot PCs variance
pcvar <- data.frame(var=var_prcomp/sum(var_prcomp), pc=c(1:length(var_prcomp)))
ggplot(pcvar[1:15,], aes(x = pc)) + geom_line(aes(y=var)) + 
  labs(x='Principal Component', y='Explained Variance') + 
  geom_point(aes(y=var)) + 
  ggtitle('miRNA: Principal Components variance')
```

```
# Plot PCs scores
df <- data.frame(PC1=pca_mirna$x[,1], PC2=pca_mirna$x[,2], PC3=pca_mirna$x[,3],
                 primary_site_of_disease=clinical_skcm2$primary_site_of_disease)

ggplot(df) +
  geom_point(aes(x=PC1, y=PC2, color=factor(primary_site_of_disease)), size=5, shape=20) +
  guides(color=guide_legend("primary_site_of_disease"),fill=guide_legend("primary_site_of_disease")) +
  labs(x='PC1 (16.9%)', y='PC2 (10.4%)') +
  ggtitle("Principal Components from batch-adjusted miRNA data") +
  theme(legend.position="bottom")
```

Again, “primary site of disease” appears as one of the most relevant variables to explain most of the data variablity.

```
pcaCorrelation(clinical=clinical_skcm2[colnames(subset_log2_mirna_skcm_corrected),],
               clin_var=clin_var, pca_data=pca_mirna$x, maxlogPvalue=10)
```

##### 2.2.3 Methylation data (`subset_logit_met_skcm_corrected`)

Exploring the PCA from Methylation data (305,180 x 104)

```
pca_met <- prcomp(t(subset_logit_met_skcm_corrected))
var_prcomp <- pca_met$sdev^2

# Plot PCs variance
pcvar <- data.frame(var=var_prcomp/sum(var_prcomp), pc=c(1:length(var_prcomp)))
ggplot(pcvar[1:15,], aes(x = pc)) + geom_line(aes(y=var)) + 
  labs(x='Principal Component', y='Explained Variance') + 
  geom_point(aes(y=var)) + 
  ggtitle('Methylation: Principal Components variance')
```

```
# Plot PCs scores
df <- data.frame(PC1=pca_met$x[,1], PC2=pca_met$x[,2], PC3=pca_met$x[,3],
                 primary_site_of_disease=clinical_skcm2$primary_site_of_disease,
                 ulceration=clinical_skcm2$melanoma_ulceration)

ggplot(df) +
  geom_point(aes(x=PC1, y=PC2, color=factor(primary_site_of_disease)), size=5, shape=20) +
  guides(color=guide_legend("primary_site_of_disease"),fill=guide_legend("primary_site_of_disease")) +
  labs(x='PC1 (17.1%)', y='PC2 (11.6%)') +
  ggtitle("Principal Components from batch-adjusted Methylation data") +
  theme(legend.position="bottom")
```

For methylation the data seems more heterogenious and more related with age, survival outcome and primary site of disease.

```
pcaCorrelation(clinical=clinical_skcm2[colnames(subset_logit_met_skcm_corrected),],
               clin_var=clin_var, pca_data=pca_met$x, maxlogPvalue = 10)
```

#### 2.3 Characterization of the data: Joint exploration.

Based on first data exploration and pre-process, we know that the most rellevant variables in our omics data sets after batch correction are the primary site of disease, neoplasm disease stage and the survival outcome.

But, how much the different data-types (e.g. different omics) are “explaining the samples in the same way”? To be more specific: are there shared components or not? How many?

The joint exploration will be perofrmed for:

- mRNA + miRNA
- mRNA + methylation

##### 2.3.1 Data scaling

Before start the multi-omic approach, mean centred of each feature and the data set normalization to unit sum of squares (forbenious normalization) is recommended to make more comparabable the different omics data-types.

```
#Calculate Forbenious normalization
frobenius_rna <- norm(as.matrix(subset_log2_rna_skcm_corrected), type="F")
frobenius_mirna <- norm(as.matrix(subset_log2_mirna_skcm_corrected), type="F")
frobenius_met <- norm(as.matrix(subset_logit_met_skcm_corrected), type="F")

#Mean-centring and division by Frobenius normalization factor
rna_skcm_norm <- t(scale(t(subset_log2_rna_skcm_corrected), scale=FALSE))/frobenius_rna
mirna_skcm_norm <- t(scale(t(subset_log2_mirna_skcm_corrected), scale=FALSE))/frobenius_mirna
met_skcm_norm <- t(scale(t(subset_logit_met_skcm_corrected), scale=FALSE))/frobenius_met

##Mean-centring without Frobenius normalization
rna_skcm_norm2 <- t(scale(t(subset_log2_rna_skcm_corrected), scale=FALSE))
mirna_skcm_norm2 <- t(scale(t(subset_log2_mirna_skcm_corrected), scale=FALSE))
met_skcm_norm2 <- t(scale(t(subset_logit_met_skcm_corrected), scale=FALSE))
```

##### 2.3.2 Model selection

Now, we are ready to explore if there are share components between data-types.

The first step is to determine the number of shared/common and individual components between omics.

How to determine these numbers is still an open question nowadays, so the user can explore different methods like jive, pca-gca and pESCA, between others. Following, we show how to perform the model selection based on these three methods.

###### 2.3.2.1 JIVE

```
# Load require packages
library(r.jive)
```

```
#Using r.jive

####################################################
## mRNA + miRNA (already centered and normalized) ##
####################################################
Data <- list(mRNA=rna_skcm_norm,
             miRNA=mirna_skcm_norm)
rjive_rna_mirna <- jive(Data, scale = FALSE, center = FALSE) 


##########################################################
## mRNA + methylation (already centered and normalized) ##
##########################################################
Data <- list(mRNA=rna_skcm_norm,
              met=met_skcm_norm)
rjive_rna_met <- jive(Data, scale = FALSE, center = FALSE)
```

###### 2.3.2.2 PCA-GCA

```
# Load require packages
library(RegularizedSCA)

#Using pca-gca

####################################################
## mRNA + miRNA (already centered and normalized) ##
####################################################
data <- cbind(t(rna_skcm_norm), 
              t(mirna_skcm_norm))

# Number of variables
Jk <- c(nrow(rna_skcm_norm), nrow(mirna_skcm_norm))

# Select components
pca_gca(data, Jk, cor_min = .7)

##########################################################
## mRNA + methylation (already centered and normalized) ##
##########################################################
data <- cbind(t(rna_skcm_norm), 
              t(met_skcm_norm))

# Number of variables
Jk <- c(nrow(rna_skcm_norm), nrow(met_skcm_norm))

# Select components
pca_gca(data, Jk, cor_min = .7)
```

###### 2.3.2.3 pESCA

```
# Load require packages
library(RpESCA)
library(RSpectra)

#Using RpESCA

################################################
## mRNA + miRNA (already centered)            ##
################################################

dataSets <- list(rna=t(rna_skcm_norm2), 
                 mirna=t(mirna_skcm_norm2))

dataTypes <- 'GG'

# used parameters
opts <- list()
opts$tol_obj <- 1E-4
opts$quiet <- 1

# save the estimated alphas, selected number of PCs and cvErrors
alphas <- rep(NA,2)
R_selected_list <- as.list(1:2)
cvErrors_list <- as.list(1:2)

# alpha estimation for each data set
for (i in 1:length(dataSets)){
  # index ith data set
  X <- dataSets[[i]]
  
  # alpha estimation procedure
  alpha_est <- alpha_estimation(X,K=3,Rs = 5:20,opts=opts)
  alphas[i] <- alpha_est$alphas_mean
  R_selected_list[[i]] <- alpha_est$R_CV
  cvErrors_list[[i]] <- alpha_est$cvErrors
}
names(alphas) <- paste0('alpha', 1:2)

## Setting the parameters for the pESCA model

# concave function and its hyper-parameter
fun_concave <- 'gdp'; gamma <- 1;
penalty = 'L2' # concave L2 norm penalty

# Parameters of a pESCA with concave L2norm penalty model
opts <- list()
opts$gamma <- gamma  # hyper-parameter for the used penalty
opts$rand_start <- 0
opts$tol_obj <- 1e-6 # stopping criteria
opts$maxit   <- 500
opts$alphas  <- alphas
#opts$R # components used Default 0.5*min(I,J) - Minimum number of variables.
opts$thr_path <- 0 # generaint thresholding path or not
opts$quiet <- 1

# Model selection pESCA conave L2 norm penalty
nTries <- 15
lambdas_CV <- log10_seq(from=1, to=500, length.out=nTries)

result_CV <- pESCA_CV(dataSets, dataTypes,
                      lambdas_CV, 
                      penalty=penalty, 
                      fun_concave=fun_concave, 
                      opts=opts)


# select the model with minimum CV error
index_min_cv <- which.min(result_CV$cvErrors_mat[,1])

# cvErrors during the model selection process
result_CV$cvErrors_mat

# selected value of lambda
lambdas_CV[index_min_cv]

# fit the final model
lambdas_opt <- rep(lambdas_CV[index_min_cv],length(dataSets))
opts_opt <- result_CV$inits[[index_min_cv]]
opts_opt$tol_obj <- 1E-8 # using high precision model


pESCA_L2 <- pESCA(dataSets = dataSets,
                          dataTypes = dataTypes,
                          lambdas = lambdas_opt,
                          penalty = penalty,
                          fun_concave= fun_concave,
                          opts = opts_opt)


mu <- pESCA_L2$mu
A <- pESCA_L2$A
B <- pESCA_L2$B
S <- pESCA_L2$S

# estimated variation explained ratios for each data set or the full data set
pESCA_L2$varExpTotals

# estimated variation explained ratios of each PC for each data set or the full data set
pESCA_L2$varExpPCs

#Cut-off 1% 
sel1 <- which(pESCA_L2$varExpPCs[1,]>1) #19
sel2 <- which(pESCA_L2$varExpPCs[2,]>1) #8
intersect(sel1,sel2)
# 7 common components
# 12 dist mRNA + 1 miRNA
# Total: 20 components

#Cut-off 5%
sel1 <- which(pESCA_L2$varExpPCs[1,]>5) #4
sel2 <- which(pESCA_L2$varExpPCs[2,]>5) #5
intersect(sel1,sel2)
# 2 common components
# 2 dist mRNA + 3 miRNA
# Total: 7 components


################################################
## mRNA + Methylation (already centered)      ##
################################################

dataSets <- list(rna=t(rna_skcm_norm2), 
                 met=t(met_skcm_norm2))

dataTypes <- 'GG'

# used parameters
opts <- list()
opts$tol_obj <- 1E-4
opts$quiet <- 1

# save the estimated alphas, selected number of PCs and cvErrors
alphas <- rep(NA,2)
R_selected_list <- as.list(1:2)
cvErrors_list <- as.list(1:2)

# alpha estimation for each data set
for (i in 1:length(dataSets)){
  # index ith data set
  X <- dataSets[[i]]
  
  # alpha estimation procedure
  alpha_est <- alpha_estimation(X,K=3,Rs = 5:20,opts=opts)
  alphas[i] <- alpha_est$alphas_mean
  R_selected_list[[i]] <- alpha_est$R_CV
  cvErrors_list[[i]] <- alpha_est$cvErrors
}
names(alphas) <- paste0('alpha', 1:2)

## Setting the parameters for the pESCA model

# concave function and its hyper-parameter
fun_concave <- 'gdp'; gamma <- 1;
penalty = 'L2' # concave L2 norm penalty

# Parameters of a pESCA with concave L2norm penalty model
opts <- list()
opts$gamma <- gamma  # hyper-parameter for the used penalty
opts$rand_start <- 0
opts$tol_obj <- 1e-6 # stopping criteria
opts$maxit   <- 500
opts$alphas  <- alphas
#opts$R  # components used Default 0.5*min(I,J) - Minimum number of variables.
opts$thr_path <- 0 # generaint thresholding path or not
opts$quiet <- 1

# Model selection pESCA conave L2 norm penalty
nTries <- 15
lambdas_CV <- log10_seq(from=1, to=500, length.out=nTries)

result_CV <- pESCA_CV(dataSets, dataTypes,
                      lambdas_CV, 
                      penalty=penalty, 
                      fun_concave=fun_concave, 
                      opts=opts)


# select the model with minimum CV error
index_min_cv <- which.min(result_CV$cvErrors_mat[,1])

# cvErrors during the model selection process
result_CV$cvErrors_mat

# selected value of lambda
lambdas_CV[index_min_cv]

# fit the final model
lambdas_opt <- rep(lambdas_CV[index_min_cv],length(dataSets))
opts_opt <- result_CV$inits[[index_min_cv]]
opts_opt$tol_obj <- 1E-8 # using high precision model


pESCA_L2 <- pESCA(dataSets = dataSets,
                          dataTypes = dataTypes,
                          lambdas = lambdas_opt,
                          penalty = penalty,
                          fun_concave= fun_concave,
                          opts = opts_opt)

mu <- pESCA_L2$mu
A <- pESCA_L2$A
B <- pESCA_L2$B
S <- pESCA_L2$S

# estimated variation explained ratios for each data set or the full data set
pESCA_L2$varExpTotals

# estimated variation explained ratios of each PC for each data set or the full data set
pESCA_L2$varExpPCs

#Cut-off 1%
sel1 <- which(pESCA_L2$varExpPCs[1,]>1) #5
sel2 <- which(pESCA_L2$varExpPCs[2,]>1) #17
intersect(sel1,sel2)
# 4 common components
# 1 dist mRNA + 13 methylation
# Total: 18 components

#Cut-off 5%
sel1 <- which(pESCA_L2$varExpPCs[1,]>5) #4
sel2 <- which(pESCA_L2$varExpPCs[2,]>5) #3
intersect(sel1,sel2)
# 1 common components
# 3 dist mRNA + 2 methylation
# Total: 6 components
```

Here, there is the table summarizing the output of the different methods.

|  | Component | r.JIVE | PCA-GCA (1%) | PCA-GCA (5%) | pESCA(1%) | pESCA (5%) |
| --- | --- | --- | --- | --- | --- | --- |
| mRNA + miRNA |  |  |  |  |  |  |
|  | Common | 3 | 50 | 1 | 7 | 2 |
|  | Dist. mRNA | 20 | 15 | 1 | 12 | 2 |
|  | Dist. miRNA | 17 | 13 | 2 | 1 | 3 |
| mRNA + met |  |  |  |  |  |  |
|  | Common | 4 | 65 | 1 | 4 | 1 |
|  | Dist. mRNA | 19 | 0 | 1 | 1 | 3 |
|  | Dist. miRNA | 36 | 38 | 3 | 13 | 2 |

As we can observe, the results from the different methods diverge. We are not able to recommend a method but, in general, we can say that it seems that there’s a small part of data variation that is shared between omics. To follow with the analysis, we will explore the omicsPCA based on the output from pESCA (1%).

##### 2.3.3 Sub-space recovery

Based on the number of components identified (pESCA) we can recover the subspace.

It is interesting to take a look on the common and distinctive components identified thorugh this method.

```
################################################
## mRNA + miRNA (already centered)            ##
################################################

# Explore the common and distinctive components and those variation explained
# Common components
cc <- intersect(sel1,sel2)
pESCA_L2$varExpPCs[,cc]
```

```
##              PC1       PC2      PC4       PC5      PC6      PC8     PC16
## X_1    11.579509 10.533584 5.419378  3.820993 3.025871 1.957098 1.025214
## X_2     8.374504  5.984612 4.326859 10.239672 2.992696 5.973369 5.732144
## X_full 11.408189 10.290424 5.360978  4.164097 3.024098 2.171783 1.276817
```

```
# Distinctive components
dist_rna <- sel1[!sel1%in%cc] #rna
pESCA_L2$varExpPCs[,dist_rna]
```

```
##             PC3      PC7      PC9     PC10     PC11     PC12     PC13     PC14
## X_1    6.596958 2.900797 1.739578 1.769607 1.877849 1.629797 1.427576 1.306367
## X_2    0.000000 0.000000 0.000000 0.000000 0.000000 0.000000 0.000000 0.000000
## X_full 6.244325 2.745738 1.646591 1.675015 1.777470 1.542678 1.351267 1.236537
##            PC15    PC17     PC18      PC20
## X_1    1.244135 1.18888 1.103844 1.0237636
## X_2    0.000000 0.00000 0.000000 0.0000000
## X_full 1.177631 1.12533 1.044839 0.9690394
```

```
dist_mirna <- sel2[!sel2%in%cc] #mirna
pESCA_L2$varExpPCs[,dist_mirna]
```

```
##      X_1      X_2   X_full 
## 0.000000 3.666469 0.195987
```

```
# Keep de common and distinctive components
common_pc <- pESCA_L2$A[,cc]
colnames(common_pc) <- colnames(pESCA_L2$varExpPCs[,cc])

dist_rna_pc <- pESCA_L2$A[,dist_rna]
colnames(dist_rna_pc) <- names(dist_rna)

dist_mirna_pc <- data.frame(pESCA_L2$A[,dist_mirna])
colnames(dist_mirna_pc) <- names(dist_mirna)


# metadata for common samples
clin <- clinical_skcm2

# PCA plot
require(ggplot2)

df <- as.data.frame(common_pc)
df$group <- clin$primary_site_of_disease
df$batch <- clin$batch_number

#plot of first common components by gene expression subtype
p <- ggplot(df,aes_string("PC1","PC2",color="group"))
p + geom_point(size=5, shape=20) + ggtitle("PCA by Primary site of tumor") + theme(legend.position="bottom")
```

```
#plot of first common components by batch
p <- ggplot(df,aes_string("PC1","PC2",color="batch"))
p + geom_point(size=5, shape=20) + ggtitle("PCA by batch") + theme(legend.position="bottom")
```

Let’s explore how relate these common and distinctive components with the clinical variables.

```
df <- data.frame(common_pc,dist_rna_pc,dist_mirna_pc)
colnames(df) <- c(paste("common_",colnames(common_pc),sep=""),
                  paste("dist_rna_",colnames(dist_rna_pc),sep=""),
                  paste("dist_mirna_",colnames(dist_mirna_pc),sep=""))

pcaCorrelation(clinical=clin, clin_var, df, maxlogPvalue =10, maxPC=ncol(df), pc.names=TRUE)
```

Based on these results, it seems that there are some information shared between mRNA and miRNA related with `primary_site_of_disease`.

```
################################################
## mRNA + methylation (already centered)      ##
################################################

# Explore the common and distinctive components and those variation explained
# Common components
cc <- intersect(sel1,sel2)
pESCA_L2$varExpPCs[,cc]
```

```
##              PC2      PC5      PC7      PC8
## X_1     7.558119 6.496145 3.531933 9.899786
## X_2    11.129553 3.020236 1.436063 1.080839
## X_full 10.869422 3.273409 1.588719 1.723182
```

```
# Distinctive components
dist_rna <- sel1[!sel1%in%cc] #rna
pESCA_L2$varExpPCs[,dist_rna]
```

```
##       X_1       X_2    X_full 
## 6.4767897 0.0000000 0.4717474
```

```
dist_met <- sel2[!sel2%in%cc] #met
pESCA_L2$varExpPCs[,dist_met]
```

```
##             PC1      PC3      PC4      PC6      PC9     PC10     PC11     PC12
## X_1     0.00000 0.000000 0.000000 0.000000 0.000000 0.000000 0.000000 0.000000
## X_2    16.99856 7.476659 3.274144 2.210317 1.807941 1.563034 1.459798 1.348897
## X_full 15.76044 6.932084 3.035667 2.049325 1.676257 1.449188 1.353472 1.250648
##            PC13     PC14     PC15     PC16      PC17
## X_1    0.000000 0.000000 0.000000 0.000000 0.0000000
## X_2    1.275617 1.187714 1.131896 1.080992 1.0539083
## X_full 1.182705 1.101205 1.049452 1.002256 0.9771452
```

```
# Keep de common and distinctive components
common_pc <- pESCA_L2$A[,cc]
colnames(common_pc) <- colnames(pESCA_L2$varExpPCs[,cc])

dist_rna_pc <- data.frame(pESCA_L2$A[,dist_rna])
colnames(dist_rna_pc) <- names(dist_rna)

dist_met_pc <- pESCA_L2$A[,dist_met]
colnames(dist_met_pc) <- names(dist_met)


# metadata for common samples
clin <- clinical_skcm2

# PCA plot
require(ggplot2)

df <- as.data.frame(common_pc)
df$group <- clin$primary_site_of_disease
df$batch <- clin$batch_number


#plot of first common components by gene expression subtype
p <- ggplot(df,aes_string("PC2","PC5",color="group"))
p + geom_point(size=5, shape=20) + ggtitle("PCA by Primary site of tumor") + theme(legend.position="bottom")
```

```
#plot of first common components by batch
p <- ggplot(df,aes_string("PC2","PC5",color="batch"))
p + geom_point(size=5, shape=20) + ggtitle("PCA by batch") + theme(legend.position="bottom")
```

Let’s explore how relate these common and distinctive components with the clinical variables.

```
df <- data.frame(common_pc,dist_rna_pc,dist_met_pc)
colnames(df) <- c(paste("common_",colnames(common_pc),sep=""),
                  paste("dist_rna_",colnames(dist_rna_pc),sep=""),
                  paste("dist_met_",colnames(dist_met_pc),sep=""))

pcaCorrelation(clinical=clin, clin_var, df, maxlogPvalue =10, maxPC=ncol(df), pc.names=TRUE)
```

Within the common components we can identify a significant association with the `primary site of disease` and `survivalOutcome`.

#### 2.4 Integrative differential analysis by NPC

Now, we want to identify genes that, according to all modalities considered as a whole (mRNA, miRNA, methylation, etc.), are either deregulated or associated to an outcome of interest.

From the previous steps (individual omics exploration and joint exploration) we will be considering (1) which co-variates are necessary to include and (2) which analysis do we want to perform.

Using omicsNPC we can explore the combination of two omics. The NPC will provide two outcomes:

- The features that are differentiated in both omics, therefore we are looking into coordinated effects.
- The features that are differentiated in only one of the omics.

##### 2.4.1 mRNA + miRNA

First of all we need to have a mapping file mathcing omics.

###### 2.4.1.1 Mapping file

```
#Load the mapping info
#load('hsa-vtm-gene.Rdata')
#the mapping can be obtained using SpidermiR package

#Identify the miRBase version of mapping file and data
library(miRNAmeConverter)

miRNAs <- rownames(subset_log2_mirna_skcm_corrected)
miRNAs_mapping <- names(id)
nc <- MiRNANameConverter()
assessVersion(nc, miRNAs, verbose = FALSE) #version 16
```

```
##    version frequency
## 1     16.0       638
## 2     17.0       606
## 3     15.0       554
## 4     14.0       436
## 5     13.0       421
## 6     18.0       420
## 7     12.0       416
## 8     11.0       402
## 9     19.0       394
## 10    20.0       331
## 11    21.0       328
## 12    10.1       325
## 13    22.0       314
## 14    10.0       313
## 15     9.2       310
## 16     9.1       309
## 17     9.0       308
## 18     8.2       303
## 19     8.1       303
## 20     8.0       200
## 21     7.1       197
## 22     6.0       130
```

```
assessVersion(nc, miRNAs_mapping, verbose = FALSE) #version 20
```

```
##    version frequency
## 1     20.0       573
## 2     21.0       569
## 3     22.0       560
## 4     19.0       546
## 5     18.0       533
## 6     17.0       380
## 7     16.0       372
## 8     15.0       359
## 9     14.0       341
## 10    13.0       339
## 11    12.0       338
## 12    11.0       337
## 13    10.1       321
## 14    10.0       314
## 15     9.2       228
## 16     9.1       228
## 17     9.0       227
## 18     8.2       220
## 19     8.1       220
## 20     8.0       181
## 21     7.1       178
## 22     6.0       145
```

```
#convert miRNA annotation from our miRNA matrix from v.16 to v.20
library(anamiR)
new_mirna_matrix <- miR_converter(subset_log2_mirna_skcm_corrected, remove_old = TRUE, original_version=16, latest_version = 20)
#Original data 898 miRNAs, now 686 miRNAs

#restricting mapping info to measured miRNA
id_mapping <- id[intersect(names(id), rownames(new_mirna_matrix))] 
#from the initial 686 miRNA features we get 239 miRNA shared with mapping info

#creating the miRNA association tables
mirna2GeneMap <- stack(id_mapping) #convert in a data.frame. Each line has a gene and one mirna. 
names(mirna2GeneMap) <- c('Gene', 'miRNA')
#Dimension: 22,471 x 2

head(mirna2GeneMap)
```

```
##     Gene         miRNA
## 1 AARSD1 hsa-let-7b-5p
## 2   AATF hsa-let-7b-5p
## 3 ABCB10 hsa-let-7b-5p
## 4  ABCB8 hsa-let-7b-5p
## 5  ABCC1 hsa-let-7b-5p
## 6  ABCF1 hsa-let-7b-5p
```

```
#creating the mRNA association tables
gene_names_rna <- unlist(lapply(strsplit(rownames(subset_log2_rna_skcm_corrected),"\\|"),
                                FUN=function(x){return(x[1])}))
rownames(subset_log2_rna_skcm_corrected) <- gene_names_rna

expr2GeneMap <- data.frame(measurement = gene_names_rna, 
                           Gene = gene_names_rna)
#Dimension: 20,225 x 2

#creating the data mapping
dataMappingExprMirna <- combiningMappings(mappings = list(expr = expr2GeneMap, 
                                                          mirna = mirna2GeneMap),
                                          retainAll = TRUE, reference = 'Gene'); 
#Dimension: 21,321 x 3

head(dataMappingExprMirna)
```

```
##      expr           mirna   Gene
## 8     A2M   hsa-miR-98-5p    A2M
## 9     A2M  hsa-miR-122-5p    A2M
## 11 A4GALT hsa-miR-193b-3p A4GALT
## 14   AAAS   hsa-miR-28-5p   AAAS
## 15   AAAS hsa-miR-1296-5p   AAAS
## 16   AAAS hsa-miR-193b-3p   AAAS
```

We can see that based on our mRNA and miRNA analysis, we have 21,321 pairs in total, it corresponds to 9,491 unique genes (46.9% from the whole mRNA dataset) and 239 unique miRNAs (26.6% from the whole miRNA dataset).

Then, we define the relevant clinical variables in each escenario. The variable of interest in the case of SKCM is the `primary_site_of_disease`. Moreover, `years_to_birth` is introduced in the model as a confounder.

```
clinicalVariables <- list(expr = c('years_to_birth','primary_site_of_disease'), 
                          met = c('years_to_birth','primary_site_of_disease'),
                          mirna = c('years_to_birth','primary_site_of_disease'));
```

```
#restrict the datasets to the elements of the data mapping
exprTMP <- subset_log2_rna_skcm_corrected[unique(dataMappingExprMirna$expr), ]; #9491 x 104

mirnaTMP <- new_mirna_matrix[unique(dataMappingExprMirna$mirna), ]; #239 x 104

#specifying the data types.
dataTypesExprMirna <- c(expr = 'continuous', mirna = 'continuous')

#preparing the datasets
mexpr <- as.matrix(clinical_skcm2[,clinicalVariables[['expr']]])
rownames(mexpr) <- rownames(clinical_skcm2)
colnames(mexpr) <- clinicalVariables[['expr']]

exprTMP <- createOmicsExpressionSet(Data = as.matrix(exprTMP), 
                                    pData = mexpr)

mmirn <- as.matrix(clinical_skcm2[,clinicalVariables[['mirna']]])
rownames(mmirn) <- rownames(clinical_skcm2)
colnames(mmirn) <- clinicalVariables[['mirna']]

mirnaTMP <- createOmicsExpressionSet(Data = as.matrix(mirnaTMP), 
                                     pData = mmirn)

dataInputExprMirna <- list(expr = exprTMP, mirna = mirnaTMP)
```

###### 2.4.1.2 Parametric Combination

```
set.seed(12345)
omicsPCRes_cPC <- omicsPC(dataInput = dataInputExprMirna,
                       dataMapping = dataMappingExprMirna,
                       dataTypes = dataTypesExprMirna,
                       verbose = TRUE)
```

The output of `omicsPC` provides information regarding p-value and adjusted p-value from both individual and combined analysis.

```
res <- omicsPCRes_cPC
str(res)
```

```
## List of 4
##  $ expr     :'data.frame':   9491 obs. of  6 variables:
##   ..$ logFC    : num [1:9491] -3.356 0.445 0.125 0.252 -2.491 ...
##   ..$ AveExpr  : num [1:9491] 3.58 6.53 9.79 1.11 1.01 ...
##   ..$ t        : num [1:9491] -3.74 1.394 0.928 0.499 -5.206 ...
##   ..$ P.Value  : num [1:9491] 4.35e-04 1.69e-01 3.57e-01 6.19e-01 2.85e-06 ...
##   ..$ adj.P.Val: num [1:9491] 0.02611 0.7101 0.86608 0.95332 0.00075 ...
##   ..$ B        : num [1:9491] -0.13 -5.27 -5.78 -6.08 4.5 ...
##  $ mirna    :'data.frame':   239 obs. of  6 variables:
##   ..$ logFC    : num [1:239] -6.34e-01 2.48e-01 3.76e-02 -1.81e-03 5.27e-05 ...
##   ..$ AveExpr  : num [1:239] 7.446734 0.309937 5.859169 0.008503 0.000168 ...
##   ..$ t        : num [1:239] -1.75156 1.12416 0.12963 -0.10786 0.00736 ...
##   ..$ P.Value  : num [1:239] 0.0854 0.2659 0.8973 0.9145 0.9942 ...
##   ..$ adj.P.Val: num [1:239] 0.435 0.702 0.995 0.995 0.997 ...
##   ..$ B        : num [1:239] -5.67 -6.54 -7.17 -7.17 -7.18 ...
##  $ pvaluesPC:'data.frame':   21321 obs. of  8 variables:
##   ..$ expr     : chr [1:21321] "A2M" "A2M" "A4GALT" "AAAS" ...
##   ..$ mirna    : chr [1:21321] "hsa-miR-98-5p" "hsa-miR-122-5p" "hsa-miR-193b-3p" "hsa-miR-28-5p" ...
##   ..$ Fisher   : num [1:21321] 0.000416 0.001163 0.437606 0.692071 0.7227 ...
##   ..$ Liptak   : num [1:21321] 0.000446 0.002582 0.586256 0.760884 0.936259 ...
##   ..$ Tippett  : num [1:21321] 0.000869 0.000869 0.337903 0.714267 0.714267 ...
##   ..$ Benjamini: num [1:21321] 0.000869 0.000869 0.337903 0.714267 0.714267 ...
##   ..$ Simes    : num [1:21321] 0.000869 0.000869 0.337903 0.714267 0.714267 ...
##   ..$ Sidak    : num [1:21321] 0.000869 0.000869 0.309358 0.586723 0.586723 ...
##  $ qvaluesPC:'data.frame':   21321 obs. of  8 variables:
##   ..$ expr     : chr [1:21321] "A2M" "A2M" "A4GALT" "AAAS" ...
##   ..$ mirna    : chr [1:21321] "hsa-miR-98-5p" "hsa-miR-122-5p" "hsa-miR-193b-3p" "hsa-miR-28-5p" ...
##   ..$ Fisher   : num [1:21321] 0.0223 0.0478 0.7435 0.8979 0.9167 ...
##   ..$ Liptak   : num [1:21321] 0.0277 0.0861 0.8814 0.9611 1 ...
##   ..$ Tippett  : num [1:21321] 0.0405 0.0405 0.6694 0.9856 0.9856 ...
##   ..$ Benjamini: num [1:21321] 0.0405 0.0405 0.6358 0.9297 0.9297 ...
##   ..$ Simes    : num [1:21321] 0.0405 0.0405 0.6358 0.9297 0.9297 ...
##   ..$ Sidak    : num [1:21321] 0.0405 0.0405 0.6129 0.8096 0.8096 ...
```

```
#Nominal p-values plot individual omics
data_plot <- data.frame(pval=c(res$expr[,"P.Value"],res$mirna[,"P.Value"]), 
                        group=c(rep(names(res)[1], nrow(res$expr)), 
                                rep(names(res)[2], nrow(res$mirna))))

p1 <- ggplot(data_plot, aes(x=group, y=pval, fill=group)) +
  geom_violin(trim=FALSE)+
  labs(title='Plot of individual omics p-value from overlapping samples')

#FDR plot individual omics
data_plot <- data.frame(pval=c(res$expr[,"adj.P.Val"],res$mirna[,"adj.P.Val"]), 
                        group=c(rep(names(res)[1], nrow(res$expr)),
                                rep(names(res)[2], nrow(res$mirna))))

p2 <- ggplot(data_plot, aes(x=group, y=pval, fill=group)) +
  geom_violin(trim=FALSE)+
  labs(title='Plot of individual omics FDR from overlapping samples')


#Nominal p-value plot PC
data_plot <- data.frame(pval=unlist(res$pvaluesPC[,-c(1,2)]), 
                        combination_method=rep(colnames(res$pvaluesPC[,-c(1,2)]),
                                               each=nrow(res$pvaluesPC)))

p3 <- ggplot(data_plot, aes(x=combination_method, y=pval, 
                            fill=combination_method)) + 
  geom_violin(trim=FALSE)+
  labs(title='Plot of p-values in overlapping omicsPC')


#FDR plot PC
data_plot <- data.frame(pval=unlist(res$qvaluesPC[,-c(1,2)]), 
                        combination_method=rep(colnames(res$qvaluesPC[,-c(1,2)]),
                                               each=nrow(res$qvaluesPC)))

p4 <- ggplot(data_plot, aes(x=combination_method, y=pval,
                            fill=combination_method)) +
  geom_violin(trim=FALSE)+
  labs(title='Plot of FDR in overlapping omicsPC')


grid.arrange(p1, p2, p3, p4, ncol=2, nrow=2)
```

Now, based on FDR results we can check if there are an increase of statistical power when both omics are jointly analyzed.

Note: here we consider significant an FDR<0.05.

```
#Significant genes or miRNA from individual omics (FDR<0.05)
sig_expr_cPC <- rownames(res$expr)[which(res$expr[,"adj.P.Val"]<0.05)]
# 216 genes

#significant miRNA
(sig_mirna_cPC <- rownames(res$mirna)[which(res$mirna[,"adj.P.Val"]<0.05)])
```

```
## [1] "hsa-miR-1227-3p" "hsa-miR-204-5p"  "hsa-miR-200b-3p" "hsa-miR-200a-3p"
## [5] "hsa-miR-140-5p"  "hsa-miR-202-3p"
```

```
# 6 miRNA 

#Significant genes from mRNA + miRNA parametric combination (Fisher, FDR<0.05)
sig_expr_mirna_cPC <- res$qvaluesPC[which(res$qvaluesPC$Fisher<0.05),1:2] 
# 529 pairs


#venn diagram
m1 <- list(rna=sig_expr_cPC, rna_fisher=unique(sig_expr_mirna_cPC$expr))

vennPlot1 <- venn.diagram(m1, NULL, fill=c('red','blue'), 
                         cat.fontface=4, lty=2, cex=1.2, cat.cex=1.2, main='Significant genes identified with individual (rna) or integration (rna_fisher)\n omics analysis: PC')
grid.newpage()
grid.draw(vennPlot1)
```

```
m2 <- list(mirna=sig_mirna_cPC, mirna_fisher=unique(sig_expr_mirna_cPC$mirna))

vennPlot2 <- venn.diagram(m2, NULL, fill=c('green','blue'), 
                         cat.fontface=4, lty=2, cex=1.2, cat.cex=1.2, main='Significant miRNAs identified with individual (mirna) or integration (mirna_fisher)\n omics analysis: PC')

grid.newpage()
grid.draw(vennPlot2)
```

We identified new 190 significant genes and 77 new mirnas with the joint exploration.

Here we can see the list of some of the new genes (n=190) identified based on Fisher combination.

```
setdiff(sig_expr_mirna_cPC$expr,sig_expr_cPC)[1:50]
```

```
##  [1] "ACPP"     "ADAM9"    "AKAP1"    "ALDH3A1"  "ALPL"     "ANAPC5"  
##  [7] "AP1S1"    "AP1S2"    "ARHGAP26" "ARHGAP29" "ATP2B1"   "BAP1"    
## [13] "BCL2"     "BCL2L2"   "BEX2"     "BIRC2"    "BPTF"     "BRD2"    
## [19] "C11orf52" "CASD1"    "CCDC121"  "CCL28"    "CCNE2"    "CCNT2"   
## [25] "CDH2"     "CDX2"     "CIDEA"    "CLUAP1"   "COL5A3"   "CPA4"    
## [31] "CREB1"    "CREB5"    "CSDE1"    "CSK"      "CTNNB1"   "CTNNBIP1"
## [37] "CXCL3"    "DLX5"     "DNMT1"    "DNPEP"    "E2F3"     "EDEM1"   
## [43] "EFNB2"    "ELMO2"    "ELOVL6"   "ENO1"     "EP300"    "ERBB2IP" 
## [49] "ETS1"     "EZR"
```

And part of the new miRNAs (n=77) identified based on Fisher combination.

```
setdiff(sig_expr_mirna_cPC$mirna,sig_mirna_cPC)[1:50]
```

```
##  [1] "hsa-miR-98-5p"   "hsa-miR-122-5p"  "hsa-miR-185-5p"  "hsa-miR-148b-3p"
##  [5] "hsa-let-7b-5p"   "hsa-miR-23a-3p"  "hsa-miR-27b-3p"  "hsa-miR-25-3p"  
##  [9] "hsa-miR-21-5p"   "hsa-miR-34a-5p"  "hsa-miR-1226-3p" "hsa-miR-145-5p" 
## [13] "hsa-miR-26b-5p"  "hsa-miR-335-5p"  "hsa-miR-331-3p"  "hsa-miR-373-3p" 
## [17] "hsa-miR-744-5p"  "hsa-miR-766-3p"  "hsa-miR-186-5p"  "hsa-miR-146a-5p"
## [21] "hsa-miR-203a"    "hsa-miR-215-5p"  "hsa-miR-671-5p"  "hsa-miR-10b-5p" 
## [25] "hsa-miR-193b-3p" "hsa-miR-339-5p"  "hsa-miR-224-5p"  "hsa-miR-150-5p" 
## [29] "hsa-miR-93-5p"   "hsa-miR-32-5p"   "hsa-let-7d-5p"   "hsa-miR-33a-5p" 
## [33] "hsa-miR-17-5p"   "hsa-miR-192-5p"  "hsa-miR-10a-5p"  "hsa-miR-27a-3p" 
## [37] "hsa-miR-155-5p"  "hsa-miR-30a-5p"  "hsa-miR-1180-3p" "hsa-miR-340-3p" 
## [41] "hsa-miR-148a-3p" "hsa-miR-223-3p"  "hsa-miR-149-5p"  "hsa-miR-451a"   
## [45] "hsa-miR-505-3p"  "hsa-miR-31-5p"   "hsa-miR-15a-5p"  "hsa-miR-125a-5p"
## [49] "hsa-miR-342-3p"  "hsa-miR-504-5p"
```

###### 2.4.1.3 Non Parametric Combination

```
set.seed(12345)

# Setting methods to combine pvalues
combMethods <- c("Fisher", "Liptak", "Tippett")
# Setting number of permutations
numPerms <- 1000
# Setting number of cores
numCores <- 1
# Setting omicsNPC to print out the steps that it performs.
verbose <- TRUE

omicsNPCRes_cNPC_rna_mirna <- omicsNPC(dataInput = dataInputExprMirna,
                         dataMapping = dataMappingExprMirna,
                         dataTypes = dataTypesExprMirna,
                         combMethods = combMethods, 
                         numPerms = numPerms,
                         numCores = numCores,
                         verbose = verbose)
```

The output of `omicsNPC` provides information regarding p-value and adjusted p-value from both individual and combined analysis.

```
res <- omicsNPCRes_cNPC_rna_mirna
str(res)
```

```
## List of 4
##  $ expr      :'data.frame':  9491 obs. of  6 variables:
##   ..$ logFC    : num [1:9491] -3.356 0.445 0.125 0.252 -2.491 ...
##   ..$ AveExpr  : num [1:9491] 3.58 6.53 9.79 1.11 1.01 ...
##   ..$ t        : num [1:9491] -3.74 1.394 0.928 0.499 -5.206 ...
##   ..$ P.Value  : num [1:9491] 4.35e-04 1.69e-01 3.57e-01 6.19e-01 2.85e-06 ...
##   ..$ adj.P.Val: num [1:9491] 0.02611 0.7101 0.86608 0.95332 0.00075 ...
##   ..$ B        : num [1:9491] -0.13 -5.27 -5.78 -6.08 4.5 ...
##  $ mirna     :'data.frame':  239 obs. of  6 variables:
##   ..$ logFC    : num [1:239] -6.34e-01 2.48e-01 3.76e-02 -1.81e-03 5.27e-05 ...
##   ..$ AveExpr  : num [1:239] 7.446734 0.309937 5.859169 0.008503 0.000168 ...
##   ..$ t        : num [1:239] -1.75156 1.12416 0.12963 -0.10786 0.00736 ...
##   ..$ P.Value  : num [1:239] 0.0854 0.2659 0.8973 0.9145 0.9942 ...
##   ..$ adj.P.Val: num [1:239] 0.435 0.702 0.995 0.995 0.997 ...
##   ..$ B        : num [1:239] -5.67 -6.54 -7.17 -7.17 -7.18 ...
##  $ pvaluesNPC:'data.frame':  21321 obs. of  7 variables:
##   ..$ expr             : chr [1:21321] "A2M" "A2M" "A4GALT" "AAAS" ...
##   ..$ mirna            : chr [1:21321] "hsa-miR-98-5p" "hsa-miR-122-5p" "hsa-miR-193b-3p" "hsa-miR-28-5p" ...
##   ..$ perm_pvalue_expr : num [1:21321] 0.000999 0.000999 0.164835 0.348651 0.348651 ...
##   ..$ perm_pvalue_mirna: num [1:21321] 0.0869 0.2937 0.8821 0.7293 0.6843 ...
##   ..$ Fisher           : num [1:21321] 0.002 0.004 0.415 0.589 0.592 ...
##   ..$ Liptak           : num [1:21321] 0.003 0.00799 0.55644 0.55145 0.53746 ...
##   ..$ Tippett          : num [1:21321] 0.002 0.002 0.297 0.575 0.578 ...
##  $ qvaluesNPC:'data.frame':  21321 obs. of  7 variables:
##   ..$ expr             : chr [1:21321] "A2M" "A2M" "A4GALT" "AAAS" ...
##   ..$ mirna            : chr [1:21321] "hsa-miR-98-5p" "hsa-miR-122-5p" "hsa-miR-193b-3p" "hsa-miR-28-5p" ...
##   ..$ perm_qvalue_expr : num [1:21321] 0.0298 0.0298 0.7446 0.8665 0.8665 ...
##   ..$ perm_qvalue_mirna: num [1:21321] 0.325 0.435 0.734 0.719 0.758 ...
##   ..$ Fisher           : num [1:21321] 0.0637 0.073 0.5362 0.6217 0.6157 ...
##   ..$ Liptak           : num [1:21321] 0.0853 0.1233 0.6737 0.6752 0.6571 ...
##   ..$ Tippett          : num [1:21321] 0 0 0.396 0.54 0.54 ...
```

```
#Nominal p-values plot individual omics
data_plot <- data.frame(pval=c(res$expr[,"P.Value"],res$mirna[,"P.Value"]), 
                        group=c(rep(names(res)[1], nrow(res$expr)), 
                                rep(names(res)[2], nrow(res$mirna))))

p1 <- ggplot(data_plot, aes(x=group, y=pval, fill=group)) +
  geom_violin(trim=FALSE)+
  labs(title='Plot of individual omics p-value from overlapping samples')

#FDR plot individual omics
data_plot <- data.frame(pval=c(res$expr[,"adj.P.Val"],res$mirna[,"adj.P.Val"]),
                        group=c(rep(names(res)[1], nrow(res$expr)),
                                rep(names(res)[2], nrow(res$mirna))))

p2 <- ggplot(data_plot, aes(x=group, y=pval, fill=group)) +
  geom_violin(trim=FALSE)+
  labs(title='Plot of individual omics FDR from overlapping samples')


#Nominal p-value plot NPC
data_plot <- data.frame(pval=unlist(res$pvaluesNPC[,-c(1:4)]), 
                        combination_method=rep(colnames(res$pvaluesNPC[,-c(1:4)]),
                                               each=nrow(res$pvaluesNPC)))

p3 <- ggplot(data_plot, aes(x=combination_method, y=pval, fill=combination_method)) +
  geom_violin(trim=FALSE)+
  labs(title='Plot of p-values in overlapping omicsNPC')


#FDR plot NPC
data_plot <- data.frame(pval=unlist(res$qvaluesNPC[,-c(1:4)]), 
                        combination_method=rep(colnames(res$qvaluesNPC[,-c(1:4)]),
                                               each=nrow(res$qvaluesNPC)))

p4 <- ggplot(data_plot, aes(x=combination_method, y=pval, fill=combination_method)) +
  geom_violin(trim=FALSE)+
  labs(title='Plot of FDR in overlapping omicsNPC')


grid.arrange(p1, p2, p3, p4, ncol=2, nrow=2)
```

Now, based on FDR results we can check if there are an increase of statistical power when both omics are jointly analyzed.

Note: here we consider significant a FDR<0.05.

```
#Significant genes or miRNA from individual omics (FDR<0.05)

#significant genes
sig_expr_cNPC <- rownames(res$expr)[which(res$expr[,"adj.P.Val"]<0.05)]
#216 gene

#significant miRNA
(sig_mirna_cNPC <- rownames(res$mirna)[which(res$mirna[,"adj.P.Val"]<0.05)])
```

```
## [1] "hsa-miR-1227-3p" "hsa-miR-204-5p"  "hsa-miR-200b-3p" "hsa-miR-200a-3p"
## [5] "hsa-miR-140-5p"  "hsa-miR-202-3p"
```

```
#6 mirna

#Significant genes from mRNA + miRNA non parametric combination (Fisher, FDR<0.05)
sig_expr_mirna_cNPC <- res$qvaluesNPC[which(res$qvaluesNPC[,'Fisher']<0.05),1:2]
#114 pairs


#venn diagram
m1 <- list(rna=sig_expr_cNPC, rna_fisher=unique(sig_expr_mirna_cNPC$expr))

vennPlot1 <- venn.diagram(m1, NULL, fill=c('red','blue'), 
                         cat.fontface=4, lty=2, cex=1.2, cat.cex=1.2, main='Significant genes identified with individual (rna) or integration (rna_fisher)\n omics analysis: NPC')
grid.newpage()
grid.draw(vennPlot1)
```

```
m2 <- list(mirna=sig_mirna_cNPC, mirna_fisher=unique(sig_expr_mirna_cNPC$mirna))

vennPlot2 <- venn.diagram(m2, NULL, fill=c('green','blue'), 
                         cat.fontface=4, lty=2, cex=1.2, cat.cex=1.2, main='Significant miRNAs identified with individual (mirna) or integration (mirna_fisher)\n omics analysis: NPC')

grid.newpage()
grid.draw(vennPlot2)
```

For the Non-Parametric approach we get 48 new genes and 14 new miRNAs.

Here we can see the list of new genes (n=48) identified based on Fisher combination.

```
setdiff(sig_expr_mirna_cNPC$expr,sig_expr_cNPC)
```

```
##  [1] "ADAM9"    "ALPL"     "ATP2B1"   "BAP1"     "CAMK4"    "CCDC88C" 
##  [7] "CCL28"    "CREB5"    "CTNNBIP1" "ERBB2IP"  "FLT1"     "FZD1"    
## [13] "GIMAP4"   "GPHB5"    "HACL1"    "HDAC4"    "HEMGN"    "HSPA2"   
## [19] "ITGB2"    "KLF11"    "KLHL20"   "LBR"      "LIN28B"   "MMP13"   
## [25] "POM121"   "POP1"     "PSD3"     "PTPRD"    "RAB38"    "RAB40B"  
## [31] "RELA"     "RPTOR"    "SERPINE1" "SNAI2"    "SNAP23"   "SOST"    
## [37] "SPNS1"    "TBC1D15"  "TCOF1"    "TERF2IP"  "TFRC"     "TGFBR2"  
## [43] "TRAF6"    "UBE2I"    "USP14"    "WASF3"    "XIAP"     "ZEB2"
```

And the new miRNAs (n=14) identified based on Fisher combination.

```
setdiff(sig_expr_mirna_cNPC$mirna,sig_mirna_cNPC)
```

```
##  [1] "hsa-miR-148b-3p" "hsa-miR-23a-3p"  "hsa-let-7b-5p"   "hsa-miR-744-5p" 
##  [5] "hsa-miR-146a-5p" "hsa-miR-215-5p"  "hsa-miR-20a-5p"  "hsa-miR-27a-3p" 
##  [9] "hsa-miR-340-3p"  "hsa-miR-148a-3p" "hsa-miR-31-5p"   "hsa-miR-30e-5p" 
## [13] "hsa-miR-18a-5p"  "hsa-miR-149-5p"
```

###### 2.4.1.4 Integration strategies comparison

Finally, we can compare the divergences between parametric and non-parametric approaches.

```
#Significant pairs from different approaches

#parmetric combination of significant pairs
PC <- paste(sig_expr_mirna_cPC$expr,sig_expr_mirna_cPC$mirna, sep=":") #529
#non parmetric combination of significant pairs
NPC <- paste(sig_expr_mirna_cNPC$expr,sig_expr_mirna_cNPC$mirna, sep=":") #114

res_all <- list(PC=PC, NPC=NPC)

vennPlot_m1 <- venn.diagram(res_all, NULL, fill=c("green","blue"), 
                            cat.fontface=4, lty=2, cex=1.2, cat.cex=1.2, 
                          main="Significant pairs for the different escenarios \n FDR (cut-off=0.05)")

grid.newpage()
grid.draw(vennPlot_m1)
```

If we look in the significant pairs identified by each approach (FDR<0.05), we see that almost all the pairs identified by NPC are also identified by PC, but PC identifies a large number of additonal pairs as significant (n=423).

##### 2.4.2 mRNA + Methylation

First of all we need to have a mapping file mathcing omics.

###### 2.4.2.1 Mapping file

```
#Load the mapping info
#load('methy2GeneMap.RData')
methy2GeneMap <- methy2GeneMap[-40902, ]; #repetition

#restricting mapping info to methylation info
methy2GeneMap_1 <- methy2GeneMap[methy2GeneMap[,1]%in%rownames(subset_logit_met_skcm_corrected),] #from the initial 305,180 methyl features we get 55,774 in common

head(methy2GeneMap_1)
```

```
##          name   Gene
## 15 cg02725885 PPAP2C
## 16 cg09908879 PPAP2C
## 17 cg16691033 PPAP2C
## 18 cg02935627 PPAP2C
## 19 cg12272837 PPAP2C
## 20 cg25697727 PPAP2C
```

```
#creating the mRNA association tables has been previously created
tail(expr2GeneMap)
```

```
##       measurement      Gene
## 20220      ZYG11B    ZYG11B
## 20221         ZYX       ZYX
## 20222       ZZEF1     ZZEF1
## 20223        ZZZ3      ZZZ3
## 20224   psiTPTE22 psiTPTE22
## 20225        tAKR      tAKR
```

```
#creating the data mapping (55,729 x 3)
dataMappingExprMet <- combiningMappings(mappings = list(expr = expr2GeneMap, 
                                                        met = methy2GeneMap_1), 
                                          retainAll = TRUE, reference = 'Gene'); 

head(dataMappingExprMet)
```

```
##      expr        met   Gene
## 6     A2M cg00134295    A2M
## 8  A4GALT cg07393322 A4GALT
## 9  A4GALT cg01268859 A4GALT
## 10 A4GALT cg20706071 A4GALT
## 11 A4GALT cg26705472 A4GALT
## 12  A4GNT cg23669440  A4GNT
```

We can see that based on our mRNA and methylation analysis, we have 55,729 pairs in total, corresponding to 9,564 unique genes (47.3% from the whole mRNA dataset) and 55,729 unique methylation sites (18% from teh whole methylation dataset).

Then, we define the relevant clinical variables in each escenario. The variable of interest in the case of SKCM is the `primary_site_of_disease`. Moreover, `years_to_birth` is introduced in the model as a confounder.

```
clinicalVariables <- list(expr = c('years_to_birth','primary_site_of_disease'), 
                          met = c('years_to_birth','primary_site_of_disease'),
                          mirna = c('years_to_birth','primary_site_of_disease'));
```

```
#restrict the datasets to the elements of the data mapping
exprTMP <- subset_log2_rna_skcm_corrected[unique(dataMappingExprMet$expr),]
#Dimension: 9564 x 104

metTMP <- subset_logit_met_skcm_corrected[unique(dataMappingExprMet$met),]
#Dimension: 55729 x 104

#specifying the data types.
dataTypesExprMet <- c(expr = 'continuous', met = 'continuous')


#preparing the datasets
mexpr <- as.matrix(clinical_skcm2[,clinicalVariables[['expr']]])
rownames(mexpr) <- rownames(clinical_skcm2)
colnames(mexpr) <- clinicalVariables[['expr']]

exprTMP <- createOmicsExpressionSet(Data = as.matrix(exprTMP), 
                                    pData = mexpr)

mmet <- as.matrix(clinical_skcm2[,clinicalVariables[['met']]])
rownames(mmet) <- rownames(clinical_skcm2)
colnames(mmet) <- clinicalVariables[['met']]

metTMP <- createOmicsExpressionSet(Data = as.matrix(metTMP), 
                                     pData = mmet)

dataInputExprMet <- list(expr = exprTMP, met = metTMP)
```

###### 2.4.2.2 Parametric Combination

```
set.seed(12345)
omicsPCRes_cPC <- omicsPC(dataInput = dataInputExprMet,
                       dataMapping = dataMappingExprMet,
                       dataTypes = dataTypesExprMet,
                       verbose = TRUE)
```

The output of `omicsPC` provides information regarding p-value and adjusted p-value from both individual and combined analysis.

```
res <- omicsPCRes_cPC
str(res)
```

```
## List of 4
##  $ expr     :'data.frame':   9564 obs. of  6 variables:
##   ..$ logFC    : num [1:9564] -3.356 0.445 0.119 0.125 0.252 ...
##   ..$ AveExpr  : num [1:9564] 3.579 6.529 0.686 9.792 1.109 ...
##   ..$ t        : num [1:9564] -3.737 1.393 0.784 0.928 0.499 ...
##   ..$ P.Value  : num [1:9564] 0.000439 0.169274 0.436072 0.357572 0.619693 ...
##   ..$ adj.P.Val: num [1:9564] 0.0213 0.6734 0.8876 0.8515 0.9487 ...
##   ..$ B        : num [1:9564] -0.15 -5.29 -5.92 -5.8 -6.1 ...
##  $ met      :'data.frame':   55729 obs. of  6 variables:
##   ..$ logFC    : num [1:55729] -0.381 -0.588 -0.67 -0.272 -0.151 ...
##   ..$ AveExpr  : num [1:55729] -1.84 -0.979 -1.33 -2.729 -4.22 ...
##   ..$ t        : num [1:55729] -1.157 -1.562 -1.212 -0.443 -0.459 ...
##   ..$ P.Value  : num [1:55729] 0.252 0.124 0.231 0.659 0.648 ...
##   ..$ adj.P.Val: num [1:55729] 0.754 0.636 0.738 0.928 0.925 ...
##   ..$ B        : num [1:55729] -5.07 -4.6 -5.02 -5.58 -5.57 ...
##  $ pvaluesPC:'data.frame':   55729 obs. of  8 variables:
##   ..$ expr     : chr [1:55729] "A2M" "A4GALT" "A4GALT" "A4GALT" ...
##   ..$ met      : chr [1:55729] "cg00134295" "cg07393322" "cg01268859" "cg20706071" ...
##   ..$ Fisher   : num [1:55729] 0.00112 0.10199 0.16559 0.3563 0.35205 ...
##   ..$ Liptak   : num [1:55729] 0.00237 0.06757 0.11546 0.34954 0.34148 ...
##   ..$ Tippett  : num [1:55729] 0.000878 0.247688 0.338549 0.338549 0.338549 ...
##   ..$ Benjamini: num [1:55729] 0.000878 0.169274 0.230509 0.338549 0.338549 ...
##   ..$ Simes    : num [1:55729] 0.000878 0.169274 0.230509 0.338549 0.338549 ...
##   ..$ Sidak    : num [1:55729] 0.000878 0.23235 0.309895 0.309895 0.309895 ...
##  $ qvaluesPC:'data.frame':   55729 obs. of  8 variables:
##   ..$ expr     : chr [1:55729] "A2M" "A4GALT" "A4GALT" "A4GALT" ...
##   ..$ met      : chr [1:55729] "cg00134295" "cg07393322" "cg01268859" "cg20706071" ...
##   ..$ Fisher   : num [1:55729] 0.0467 0.4763 0.5752 0.7663 0.7632 ...
##   ..$ Liptak   : num [1:55729] 0.085 0.425 0.521 0.774 0.769 ...
##   ..$ Tippett  : num [1:55729] 0.043 0.71 0.797 0.797 0.797 ...
##   ..$ Benjamini: num [1:55729] 0.043 0.589 0.66 0.749 0.749 ...
##   ..$ Simes    : num [1:55729] 0.043 0.589 0.66 0.749 0.749 ...
##   ..$ Sidak    : num [1:55729] 0.043 0.666 0.729 0.729 0.729 ...
```

```
#Nominal p-values plot individual omics
data_plot <- data.frame(pval=c(res$expr[,"P.Value"],res$met[,"P.Value"]), 
                        group=c(rep(names(res)[1], nrow(res$expr)), 
                                rep(names(res)[2], nrow(res$met))))

p1 <- ggplot(data_plot, aes(x=group, y=pval, fill=group)) +
  geom_violin(trim=FALSE)+
  labs(title='Plot of individual omics p-value from overlapping samples')

#FDR plot individual omics
data_plot <- data.frame(pval=c(res$expr[,"adj.P.Val"],res$met[,"adj.P.Val"]), 
                        group=c(rep(names(res)[1], nrow(res$expr)),
                                rep(names(res)[2], nrow(res$met))))

p2 <- ggplot(data_plot, aes(x=group, y=pval, fill=group)) +
  geom_violin(trim=FALSE)+
  labs(title='Plot of individual omics FDR from overlapping samples')


#Nominal p-value plot PC
data_plot <- data.frame(pval=unlist(res$pvaluesPC[,-c(1,2)]), 
                        combination_method=rep(colnames(res$pvaluesPC[,-c(1,2)]),
                                               each=nrow(res$pvaluesPC)))

p3 <- ggplot(data_plot, aes(x=combination_method, y=pval, 
                            fill=combination_method)) + 
  geom_violin(trim=FALSE)+
  labs(title='Plot of p-values in overlapping omicsPC')


#FDR plot PC
data_plot <- data.frame(pval=unlist(res$qvaluesPC[,-c(1,2)]), 
                        combination_method=rep(colnames(res$qvaluesPC[,-c(1,2)]),
                                               each=nrow(res$qvaluesPC)))

p4 <- ggplot(data_plot, aes(x=combination_method, y=pval,
                            fill=combination_method)) +
  geom_violin(trim=FALSE)+
  labs(title='Plot of FDR in overlapping omicsPC')


grid.arrange(p1, p2, p3, p4, ncol=2, nrow=2)
```

Now, based on FDR results we can check if there are an increase of statistical power when both omics are jointly analyzed.

Note: here we consider significant an FDR<0.05.

```
#Significant genes or methyl from individual omics (FDR<0.05)

#significant genes
sig_expr_cPC <- rownames(res$expr)[which(res$expr[,"adj.P.Val"]<0.05)]
# 216 gene

#significant methyl
(sig_met_cPC <- rownames(res$met)[which(res$met[,"adj.P.Val"]<0.05)])
```

```
##  [1] "cg11740099" "cg07286123" "cg18267489" "cg10578007" "cg10018632"
##  [6] "cg21023001" "cg03662648" "cg21037265" "cg00862588" "cg18827221"
## [11] "cg03053826" "cg08299521"
```

```
# 12 methylation sites

#Significant genes from mRNA + miRNA non parametric combination (Fisher, FDR<0.05)
sig_expr_met_cPC <- res$qvaluesPC[which(res$qvaluesPC[,'Fisher']<0.05),1:2]
# 1,418 pairs

#venn diagram
m1 <- list(rna=sig_expr_cPC, rna_fisher=unique(sig_expr_met_cPC$expr))

vennPlot1 <- venn.diagram(m1, NULL, fill=c('red','blue'), 
                         cat.fontface=4, lty=2, cex=1.2, cat.cex=1.2, main='Significant genes identified with individual (rna) or integration (rna_fisher)\n omics analysis: NPC (all samples)')
grid.newpage()
grid.draw(vennPlot1)
```

```
m2 <- list(met=sig_met_cPC, met_fisher=unique(sig_expr_met_cPC$met))

vennPlot2 <- venn.diagram(m2, NULL, fill=c('green','blue'), 
                         cat.fontface=4, lty=2, cex=1.2, cat.cex=1.2, main='Significant methylation sites identified with individual (met) or integration (met_fisher)\n omics analysis: NPC (all samples)')

grid.newpage()
grid.draw(vennPlot2)
```

We identified new 245 significant genes and 1,406 new mathylation sites with the joint exploration.

Here we can see the list of some of the new genes (n=245) identified based on Fisher combination.

```
setdiff(sig_expr_met_cPC$expr,sig_expr_cPC)[1:50]
```

```
##  [1] "ACR"       "ADORA1"    "AHNAK2"    "ALDH3A1"   "ALLC"      "ALOX5AP"  
##  [7] "ALPK3"     "ALX4"      "AMIGO2"    "APOBEC3F"  "APOF"      "ARHGAP25" 
## [13] "ARHGDIA"   "ARID5A"    "ATP2C2"    "AVPR1B"    "C20orf195" "C9orf156" 
## [19] "CA7"       "CALCA"     "CCBP2"     "CCDC88C"   "CCL22"     "CCL24"    
## [25] "CCNA1"     "CCR1"      "CCR5"      "CCR7"      "CCT6B"     "CD22"     
## [31] "CD37"      "CD6"       "CD63"      "CD79B"     "CD80"      "CD93"     
## [37] "CEACAM21"  "CECR5"     "CHRND"     "CLDN4"     "CLIC3"     "CMKLR1"   
## [43] "CNR2"      "CNTFR"     "COCH"      "CROCC"     "CRYBA4"    "CRYGB"    
## [49] "CYFIP2"    "CYTL1"
```

And part of the new methylation sites (n=1,406) identified based on Fisher combination.

```
setdiff(sig_expr_met_cPC$met,sig_met_cPC)[1:50]
```

```
##  [1] "cg00134295" "cg24822001" "cg12832630" "cg21475256" "cg14121103"
##  [6] "cg19150852" "cg23239109" "cg03626746" "cg02059996" "cg13606015"
## [11] "cg17200204" "cg25232725" "cg26034629" "cg05615487" "cg24287438"
## [16] "cg11773720" "cg19962990" "cg16457786" "cg22388954" "cg21438101"
## [21] "cg15458338" "cg18319029" "cg08132931" "cg11220663" "cg11341981"
## [26] "cg01133446" "cg06960824" "cg20959872" "cg22087450" "cg00443276"
## [31] "cg26095296" "cg23413349" "cg06025938" "cg18405900" "cg12448860"
## [36] "cg25263140" "cg12674357" "cg19343088" "cg27632402" "cg00974935"
## [41] "cg21079345" "cg02215070" "cg14629509" "cg06864853" "cg04663564"
## [46] "cg13801416" "cg09957386" "cg18957070" "cg24482246" "cg19188632"
```

###### 2.4.2.3 Non Parametric Combination

```
set.seed(12345)

# Setting methods to combine pvalues
combMethods <- c("Fisher", "Liptak", "Tippett")
# Setting number of permutations
numPerms <- 1000
# Setting number of cores
numCores <- 1
# Setting omicsNPC to print out the steps that it performs.
verbose <- TRUE

omicsNPCRes_cNPC <- omicsNPC(dataInput = dataInputExprMet,
                         dataMapping = dataMappingExprMet,
                         dataTypes = dataTypesExprMet,
                         combMethods = combMethods, 
                         numPerms = numPerms,
                         numCores = numCores,
                         verbose = verbose)
```

The output of `omicsNPC` provides information regarding p-value and adjusted p-value from both individual and combined analysis.

```
res <- omicsNPCRes_cNPC_rna_met

#Nominal p-values plot individual omics
data_plot <- data.frame(pval=c(res$expr[,"P.Value"],res$met[,"P.Value"]), 
                        group=c(rep(names(res)[1], nrow(res$expr)), 
                                rep(names(res)[2], nrow(res$met))))

p1 <- ggplot(data_plot, aes(x=group, y=pval, fill=group)) +
  geom_violin(trim=FALSE)+
  labs(title='Plot of individual omics p-value from overlapping samples')

#FDR plot individual omics
data_plot <- data.frame(pval=c(res$expr[,"adj.P.Val"],res$met[,"adj.P.Val"]),
                        group=c(rep(names(res)[1], nrow(res$expr)),
                                rep(names(res)[2], nrow(res$met))))

p2 <- ggplot(data_plot, aes(x=group, y=pval, fill=group)) + 
  geom_violin(trim=FALSE)+
  labs(title='Plot of individual omics FDR from overlapping samples')


#Nominal p-value plot NPC
data_plot <- data.frame(pval=unlist(res$pvaluesNPC[,-c(1:4)]), 
                        combination_method=rep(colnames(res$pvaluesNPC[,-c(1:4)])
                                               , each=nrow(res$pvaluesNPC)))

p3 <- ggplot(data_plot, aes(x=combination_method, y=pval,
                            fill=combination_method)) +
  geom_violin(trim=FALSE)+
  labs(title='Plot of p-values in overlapping omicsNPC')


#FDR plot NPC
data_plot <- data.frame(pval=unlist(res$qvaluesNPC[,-c(1:4)]), 
                        combination_method=rep(colnames(res$qvaluesNPC[,-c(1:4)])
                                               , each=nrow(res$qvaluesNPC)))

p4 <- ggplot(data_plot, aes(x=combination_method, y=pval,
                            fill=combination_method)) +
  geom_violin(trim=FALSE)+
  labs(title='Plot of FDR in overlapping omicsNPC')


grid.arrange(p1, p2, p3, p4, ncol=2, nrow=2)
```

Now, based on FDR results we can check if there are an increase of statistical power when both omics are jointly analyzed.

Note: here we consider significant an FDR<0.05.

```
#Significant genes or miRNA from individual omics (FDR<0.05)

#significant genes
sig_expr_cNPC <- rownames(res$expr)[which(res$expr[,"adj.P.Val"]<0.05)]
# 277 genes

#significant miRNA
(sig_met_cNPC <- rownames(res$met)[which(res$met[,"adj.P.Val"]<0.05)])
```

```
##  [1] "cg11740099" "cg07286123" "cg18267489" "cg10578007" "cg10018632"
##  [6] "cg21023001" "cg03662648" "cg21037265" "cg00862588" "cg18827221"
## [11] "cg03053826" "cg08299521"
```

```
# 12 methylation sites

#Significant genes from mRNA + methylation non parametric combination (Fisher, FDR<0.05)
sig_expr_met_cNPC <- res$qvaluesNPC[which(res$qvaluesNPC[,'Fisher']<0.05),1:2]
# 432 pairs found

#venn diagram
m1 <- list(rna=sig_expr_cNPC, rna_fisher=unique(sig_expr_met_cNPC$expr))

vennPlot1 <- venn.diagram(m1, NULL, fill=c('red','blue'), 
                         cat.fontface=4, lty=2, cex=1.2, cat.cex=1.2, 
                         main='Significant genes identified with individual (rna) or integration (rna_fisher)\n omics analysis: NPC (all samples)')
grid.newpage()
grid.draw(vennPlot1)
```

```
m2 <- list(met=sig_met_cNPC, met_fisher=unique(sig_expr_met_cNPC$met))

vennPlot2 <- venn.diagram(m2, NULL, fill=c('green','blue'), 
                         cat.fontface=4, lty=2, cex=1.2, cat.cex=1.2, 
                         main='Significant methylation sites identified with individual (met) or integration (met_fisher)\n omics analysis: NPC (all samples)')

grid.newpage()
grid.draw(vennPlot2)
```

For the Non-Parametric approach we get 116 new genes and 428 new methylation sites.

Here we can see part of the list of new genes (n=116) identified based on Fisher combination.

```
setdiff(sig_expr_met_cNPC$expr,sig_expr_cNPC)[1:50]
```

```
##  [1] "ALDH3A1"  "ALOX5AP"  "AMIGO2"   "AQP1"     "ATP2C2"   "C9orf156"
##  [7] "CCDC88C"  "CCL5"     "CCR1"     "CCR7"     "CD22"     "CD37"    
## [13] "CD63"     "CD79B"    "CEACAM21" "CLDN4"    "CMKLR1"   "CNTFR"   
## [19] "COCH"     "CRYBA4"   "CRYGB"    "CTLA4"    "CYTL1"    "DENND1C" 
## [25] "DENND3"   "DHRS3"    "DLG5"     "EBF2"     "ECEL1"    "EFNA1"   
## [31] "ELF3"     "EPHA3"    "EVI2B"    "FCER1G"   "FCHO1"    "FCHSD2"  
## [37] "FGD2"     "FGFR2"    "FGL2"     "GABRA5"   "GAST"     "GLUL"    
## [43] "GMFG"     "GMIP"     "HCLS1"    "HDAC11"   "HIC1"     "HLA-DMB" 
## [49] "HLF"      "HS3ST3A1"
```

And part of the new methylation sites (n=428) identified based on Fisher combination.

```
setdiff(sig_expr_met_cNPC$met,sig_met_cNPC)[1:50]
```

```
##  [1] "cg24822001" "cg16457786" "cg18957070" "cg24482246" "cg20103124"
##  [6] "cg12199378" "cg12210770" "cg24603490" "cg07473175" "cg19327844"
## [11] "cg15373767" "cg24762359" "cg16634108" "cg06786050" "cg00479411"
## [16] "cg08566293" "cg11635839" "cg19008597" "cg21500300" "cg15742700"
## [21] "cg17124224" "cg00768409" "cg26429925" "cg06523224" "cg20941110"
## [26] "cg27304406" "cg07016258" "cg27169020" "cg26654798" "cg16254374"
## [31] "cg23741520" "cg16049391" "cg14385245" "cg09180848" "cg10275315"
## [36] "cg02713760" "cg14122696" "cg12498409" "cg13486755" "cg01331191"
## [41] "cg03469122" "cg16438210" "cg19411729" "cg07115110" "cg13504059"
## [46] "cg10475172" "cg09824255" "cg05433111" "cg27565966" "cg05981394"
```

###### 2.4.2.4 Integration strategies comparison

Finally, we can compare the divergences between parametric and non-parametric approaches.

```
#Significant pairs from different approaches

#parmetric combination significant pairs
PC <- paste(sig_expr_met_cPC$expr,sig_expr_met_cPC$met, sep=":") #3491
#non parmetric combination significant pairs
NPC <- paste(sig_expr_met_cNPC$expr,sig_expr_met_cNPC$met, sep=":") #2885

res_all <- list(PC=PC, NPC=NPC)

vennPlot_m1 <- venn.diagram(res_all, NULL, fill=c("green","blue"), 
                            cat.fontface=4, lty=2, cex=1.2, cat.cex=1.2, 
                          main="Significant pairs for the different escenarios \n FDR (cut-off=0.05)")

grid.newpage()
grid.draw(vennPlot_m1)
```

If we look in the significant pairs identified in each approach (FDR<0.05), we see that both strategies identify a large number of common pairs (68%). Almost all pairs identified by NPC are also identified by PC, but PC identifies a significant number of additonal pairs as significant (n=423).

Finally, you can see the R session information used to run all the code.

```
sessionInfo()
```

```
## R version 4.0.3 (2020-10-10)
## Platform: x86_64-w64-mingw32/x64 (64-bit)
## Running under: Windows 10 x64 (build 16299)
## 
## Matrix products: default
## 
## Random number generation:
##  RNG:     Mersenne-Twister 
##  Normal:  Inversion 
##  Sample:  Rounding 
##  
## locale:
## [1] LC_COLLATE=Spanish_Spain.1252  LC_CTYPE=Spanish_Spain.1252   
## [3] LC_MONETARY=Spanish_Spain.1252 LC_NUMERIC=C                  
## [5] LC_TIME=Spanish_Spain.1252    
## 
## attached base packages:
##  [1] grid      stats4    parallel  stats     graphics  grDevices utils    
##  [8] datasets  methods   base     
## 
## other attached packages:
##  [1] edgeR_3.30.3                                      
##  [2] limma_3.44.3                                      
##  [3] data.table_1.13.0                                 
##  [4] anamiR_1.13.0                                     
##  [5] miRNAmeConverter_1.16.0                           
##  [6] miRBaseVersions.db_1.1.0                          
##  [7] sva_3.36.0                                        
##  [8] BiocParallel_1.22.0                               
##  [9] genefilter_1.70.0                                 
## [10] mgcv_1.8-33                                       
## [11] nlme_3.1-149                                      
## [12] r.jive_2.1                                        
## [13] gridExtra_2.3                                     
## [14] MASS_7.3-53                                       
## [15] STATegRa_1.17.1                                   
## [16] plotrix_3.7-8                                     
## [17] VennDiagram_1.6.20                                
## [18] futile.logger_1.4.3                               
## [19] lumi_2.40.0                                       
## [20] IlluminaHumanMethylation450kanno.ilmn12.hg19_0.6.0
## [21] minfi_1.34.0                                      
## [22] bumphunter_1.30.0                                 
## [23] locfit_1.5-9.4                                    
## [24] iterators_1.0.13                                  
## [25] foreach_1.5.1                                     
## [26] Biostrings_2.56.0                                 
## [27] XVector_0.28.0                                    
## [28] SummarizedExperiment_1.18.2                       
## [29] DelayedArray_0.14.1                               
## [30] matrixStats_0.57.0                                
## [31] Biobase_2.48.0                                    
## [32] GenomicRanges_1.40.0                              
## [33] GenomeInfoDb_1.24.2                               
## [34] IRanges_2.22.2                                    
## [35] S4Vectors_0.26.1                                  
## [36] BiocGenerics_0.34.0                               
## [37] ggfortify_0.4.11                                  
## [38] ggplot2_3.3.2                                     
## [39] survival_3.2-7                                    
## [40] reshape_0.8.8                                     
## [41] BiocStyle_2.16.1                                  
## 
## loaded via a namespace (and not attached):
##   [1] questionr_0.7.3           tidyselect_1.1.0         
##   [3] RSQLite_2.2.1             AnnotationDbi_1.50.3     
##   [5] combinat_0.0-8            munsell_0.5.0            
##   [7] codetools_0.2-16          preprocessCore_1.50.0    
##   [9] nleqslv_3.3.2             miniUI_0.1.1.1           
##  [11] withr_2.3.0               colorspace_1.4-1         
##  [13] AlgDesign_1.2.0           highr_0.8                
##  [15] knitr_1.30                rstudioapi_0.11          
##  [17] labeling_0.4.2            GenomeInfoDbData_1.2.3   
##  [19] bit64_4.0.5               farver_2.0.3             
##  [21] rhdf5_2.32.4              vctrs_0.3.4              
##  [23] generics_0.0.2            lambda.r_1.2.4           
##  [25] xfun_0.18                 BiocFileCache_1.12.1     
##  [27] R6_2.4.1                  illuminaio_0.30.0        
##  [29] bitops_1.0-6              assertthat_0.2.1         
##  [31] promises_1.1.1            scales_1.1.1             
##  [33] gtable_0.3.0              affy_1.66.0              
##  [35] methylumi_2.34.0          rlang_0.4.8              
##  [37] calibrate_1.7.7           splines_4.0.3            
##  [39] rtracklayer_1.48.0        GEOquery_2.56.0          
##  [41] BiocManager_1.30.10       yaml_2.2.1               
##  [43] abind_1.4-5               GenomicFeatures_1.40.1   
##  [45] httpuv_1.5.4              RMySQL_0.10.20           
##  [47] SpatioTemporal_1.1.9.1    tools_4.0.3              
##  [49] bookdown_0.21             nor1mix_1.3-0            
##  [51] affyio_1.58.0             ellipsis_0.3.1           
##  [53] gplots_3.1.0              RColorBrewer_1.1-2       
##  [55] siggenes_1.62.0           Rcpp_1.0.5               
##  [57] plyr_1.8.6                progress_1.2.2           
##  [59] zlibbioc_1.34.0           purrr_0.3.4              
##  [61] RCurl_1.98-1.2            prettyunits_1.1.1        
##  [63] openssl_1.4.3             cluster_2.1.0            
##  [65] haven_2.3.1               magrittr_1.5             
##  [67] futile.options_1.0.1      hms_0.5.3                
##  [69] mime_0.9                  evaluate_0.14            
##  [71] xtable_1.8-4              klaR_0.6-15              
##  [73] XML_3.99-0.5              mclust_5.4.6             
##  [75] compiler_4.0.3            biomaRt_2.44.4           
##  [77] gage_2.38.3               tibble_3.0.4             
##  [79] KernSmooth_2.23-17        crayon_1.3.4             
##  [81] agricolae_1.3-3           htmltools_0.5.0          
##  [83] later_1.1.0.1             geneplotter_1.66.0       
##  [85] tidyr_1.1.2               DBI_1.1.0                
##  [87] formatR_1.7               dbplyr_1.4.4             
##  [89] rappdirs_0.3.1            Matrix_1.2-18            
##  [91] readr_1.4.0               quadprog_1.5-8           
##  [93] forcats_0.5.0             pkgconfig_2.0.3          
##  [95] GenomicAlignments_1.24.0  xml2_1.3.2               
##  [97] annotate_1.66.0           rngtools_1.5             
##  [99] multtest_2.44.0           beanplot_1.2             
## [101] doRNG_1.8.2               scrime_1.3.5             
## [103] stringr_1.4.0             digest_0.6.25            
## [105] graph_1.66.0              rmarkdown_2.4            
## [107] base64_2.0                DelayedMatrixStats_1.10.1
## [109] curl_4.3                  shiny_1.5.0              
## [111] Rsamtools_2.4.0           gtools_3.8.2             
## [113] lifecycle_0.2.0           Rhdf5lib_1.10.1          
## [115] askpass_1.1               labelled_2.7.0           
## [117] pillar_1.4.6              lattice_0.20-41          
## [119] KEGGREST_1.28.0           GO.db_3.11.4             
## [121] fastmap_1.0.1             httr_1.4.2               
## [123] glue_1.4.2                png_0.1-7                
## [125] bit_4.0.4                 stringi_1.5.3            
## [127] HDF5Array_1.16.1          blob_1.2.1               
## [129] DESeq2_1.28.1             caTools_1.18.0           
## [131] memoise_1.1.0             dplyr_1.0.2
```
